# Intrathecal sc-AAV9-CB-GFP: Systemic Distribution Predominates Following Single-Dose Administration in Cynomolgus Macaques

**DOI:** 10.1101/2021.11.28.470258

**Authors:** Emily K. Meseck, Ghiabe Guibinga, Stephen Wang, Cameron McElroy, Eloise Hudry, Keith Mansfield

## Abstract

Biodistribution of self-complementary adeno-associated virus-9 (scAAV9)–chicken β promoter–green fluorescent protein (GFP) was assessed in juvenile cynomolgus macaques infused intrathecally via lumbar puncture or the intracisterna magna (1.0×1013 or 3.0×1013 vg/animal), with necropsy 28 days later. Our results characterized central nervous system biodistribution compared with systemic organs/tissues by droplet digital polymerase chain reaction for DNA and in situ hybridization. GFP expression was characterized by Meso Scale Discovery electrochemiluminescence immunosorbent assay and immunohistochemistry (IHC). Biodistribution was widespread but variable, with vector DNA and GFP expression greatest in the spinal cord, dorsal root ganglia (DRG), and certain systemic tissues (e.g., liver), with low concentrations in many brain regions despite direct cerebrospinal fluid administration. Transduction and expression were observed primarily in perivascular astrocytes in the brain, with a paucity in neurons. Greater GFP expression was observed in hepatocytes, striated myocytes, cardiomyocytes, spinal cord lower motor neurons, and DRG sensory neurons by IHC. These results suggest caution for use of scAAV9-based intrathecal delivery with the current expression cassette as a modality for neurologic diseases that require widespread brain neuronal expression. This capsid/expression cassette combination may be better suited for diseases that express a secreted protein and/or do not require widespread brain neuronal transduction.

## INTRODUCTION

Adeno-associated virus (AAV) vectors have emerged over the past several years as the gene replacement therapy tool of choice for myriad diseases, including neurologic diseases, and hold tremendous promise for patients across a range of indications. Following the approval of intravenous onasemnogene abeparvovec (Zolgensma^®^, Novartis Gene Therapies, Inc.),^1^ we investigated the potential to use self-complementary AAV serotype 9 (scAAV9) with alternate transgenes in the expression cassette for the treatment of other monogenic, seriously debilitating, life-threatening neurodegenerative diseases that, in contrast to spinal muscular atrophy, would require widespread brain neuronal transduction rather than requiring transduction of other aspects of the nervous system (e.g., spinal cord or peripheral nervous system).

Intrathecal administration has been investigated for AAV vector delivery because it was theorized that this route would provide direct access to central nervous system (CNS) tissues and thereby reduces vector dosage compared with intravenous delivery, potentially reducing or limiting systemic adverse events. Thus, a biodistribution study with safety endpoints was conducted in juvenile, female cynomolgus macaques of Asian geographic ancestry (*Macaca fascicularis*) to characterize the biodistribution (transduction and expression) of self-complementary adeno-associated virus-9–chicken β fluorescent protein (scAAV9-CB-GFP) in the CNS and peripheral tissues following a single intrathecal infusion and 28 days of observation. This study was conducted under a study protocol at an Association for Assessment and Accreditation of Laboratory Animal Care International (AAALAC-International)–accredited, independent contract research organization under non-Good Laboratory Practice Conditions with high scientific quality. The scAAV9-CB-GFP test article was infused into the cerebrospinal fluid (CSF) at 1.0×10^13^ or 3.0×10^13^ vg/animal by lumbar puncture (intrathecal-LP) or intracisterna magna (ICM) administration to directly compare these routes of intrathecal administration. The use of a surrogate reporter gene (e.g., green fluorescent protein [GFP]) was driven in part by the 3Rs principle to reduce the overall number of nonhuman primates (NHPs) used across distinct disease indications and development programs. The use of GFP enables a detailed transgene expression “map” of the CNS and peripheral tissues and is readily detectable by many molecular techniques. Subsequently, GFP could be replaced by a clinically relevant transgene in an identical capsid and expression cassette for a program- and/or indication-specific GLP-compliant, investigational new drug–enabling toxicity study. Use of a non-native reporter transgene, such as GFP, for a detailed biodistribution analysis avoids instances in which endogenous gene expression may be indistinguishable from transgene expression in cases when there is a high homology between the animal and human transgene or gene product.

The results provide insight into vector genome biodistribution, consisting of quantitative DNA transduction and GFP protein expression in the CNS and peripheral tissues as determined by droplet digital polymerase chain reaction (ddPCR) and Meso Scale Discovery (MSD) electrochemiluminescence immunosorbent assay (ECLIA), respectively. Molecular pathology techniques of immunohistochemistry (IHC) with a monoclonal antibody for GFP and *in situ* hybridization (ISH) for the transgene transcript conducted on formalin-fixed, paraffin-embedded (FFPE) tissues are also presented. These molecular techniques demonstrated transgene protein and/or mRNA expression in the context of the FFPE tissue morphology at a cellular level to provide greater context to the tissue homogenate analyses from ddPCR and MSD techniques. No consistent, discernibly different patterns of distribution were detected in transduction or transgene expression with administration of the scAAV9-CB-GFP vector by ICM or intrathecal- LP routes in the CNS or in the examined peripheral tissues (liver, lung, kidney, ovary, spleen, heart, mandibular lymph node, skeletal muscle, and pancreas). In the nervous system of these juvenile NHPs (aged 13–17 months), transduction and expression in lower motor neurons of the spinal cord and sensory neurons in the dorsal root (DRG) and trigeminal (TG) ganglia were strong. In contrast, transduction and expression in the brain were relatively low, generally favoring astrocytes over neurons, and had a distribution pattern that was patchy, perivascular, and/or adjacent to the meningeal surfaces of the brain.

## MATERIALS AND METHODS

### Animals

Female cynomolgus macaques (*Macaca fascicularis*) of mainland Asian geographic ancestry sourced from Worldwide Primates, Inc. (Miami, FL, USA), and were acclimated to the test facility for at least 4 weeks prior to dosing. NHPs were juveniles, ages 13 to 17 months (weight, 1.2–1.8 kg) at dosing initiation.

Pre-dose collection and analysis for anti-AAV9 antibodies were undertaken in serum samples collected at least once prior to dosing and on Day 15, Day 22, and the day of the scheduled euthanasia. In addition, CSF samples were collected on Day 1 (prior to dosing), Day 15, and on the day of the scheduled euthanasia. Analysis of serum samples and CSF samples was conducted at Labcorp-Chantilly using electrochemiluminescence assays validated for the detection of anti- AAV9 antibodies in cynomolgus monkeys for both matrices.

Novartis is committed to the highest animal welfare standards (https://www.reporting.novartis.com/novartis-in-society/strategic-areas/being-a-responsible-citizen/conducting-animal-research-responsibly.html). The study protocol described in this manuscript and all associated procedures were reviewed and approved by the contract research organization’s Institutional Animal Care and Use Committee and were compliant with applicable animal welfare regulations, including the Animal Welfare Act, the Guide for the Care and Use of Laboratory Animals, and the Office of Laboratory Animal Welfare. Labcorp Madison Preclinical site has full AAALAC-Internation accreditation. This study did not unnecessarily duplicate any previous work, and no other model fulfilled study requirements.

### Test Article and Vehicle Control Article

The test article was scAAV9-CB-GFP encoding the enhanced *GFP* gene and was used at targeted concentrations of 1.0×10^13^ and 3.0×10^13^ vg/mL. The CB promoter (CMV early enhancer and a hybrid CMV enhancer/chicken beta-actin promoter) is a ubiquitously active promoter expected to allow for transcription of vector DNA in any transduced tissue.^2^ Other vector elements included a simian virus 40–derived intron and bovine growth hormone polyadenylation signal. The vehicle control article was 20 mM Tris buffer with 1 mM MgCl_2_, 200 mM NaCl, and 0.005% poloxamer 188, pH 8.1±0.1 at 20°C.

### Experimental Design

On Study Day 1, a single dose of 1.0×10^13^ vg/animal was administered to animals in Group 3 (intrathecal-LP) or Group 5 (ICM), and 3.0×10^13^ vg/animal was administered to animals in Group 4 (intrathecal-LP) or Group 6 (ICM). Each group was comprised of n=4 animals/group of female cynomolgus macaques (NHPs) (**Supplemental Table 1**). Two additional groups, also composed of four female animals per group, were administered vehicle control article and served as controls (intrathecal-LP, Group 1 or ICM, Group 2). Following 28 days of observation, animals were euthanized, and samples were collected for analysis.

### Dose Administration

Animals were anesthetized at the time of dosing with 10.0 mg/kg of ketamine and dexmedetomidine (0.02 mg/kg for intrathecal-LP dosing, 0.01 mg/kg for ICM dosing). Animals were also administered buprenorphine (0.02 mg/kg) after the ketamine and dexmedetomidine but before the dosing procedure for intrathecal-LP or before anesthesia for the ICM intrathecal dosing. In addition, animals dosed at the ICM location were also administered meloxicam SR (0.6 mg/kg) by subcutaneous injection. Atipamezole (0.2 mg/kg for intrathecal-LP dosing and 0.1 mg/kg for ICM dosing) was administered as a reversal agent for the anesthesia shortly after the dosing procedure. As much as 1.0 mL of CSF was collected prior to dose administration at either location (LP or ICM). The scAAV9-CB-GFP and vehicle control article formulations were administered at a dose volume of 1.0 mL for each animal followed by a flush with 0.25 mL of artificial CSF. Doses were infused via intrathecal injection into the intervertebral space of L5 to L6 (for at least 1 minute) or the cisterna magna (for at least 2 minutes). Following intrathecal-LP dosing, NHPs were maintained in dorsal recumbence with hind legs elevated (Trendelenburg- like position) for at least 10 minutes following dose administration.

### Clinical Assessments

Clinical observations included (but were not limited to) twice-daily general observations, once-daily cage-side observations, and detailed observations (at least weekly), weekly body weights, and qualitative daily food consumption determinations for all groups prior to dosing and during the observation period as indicated by the study protocol. Neurologic exams were conducted on Days 1 and 2 (approximately 4- and 24-hours post-dose, respectively), once during Week 2, once during Week 4 of the observation period, and twice during the pre-dose phase. Clinical laboratory evaluations were performed at least once during the pre-dose phase (hematology, coagulation, and clinical chemistry), on Day 2 (hematology only), and on Day 15 (hematology and clinical chemistry) of the observation period, as well as on the day of scheduled euthanasia, Day 28 (hematology, coagulation, and clinical chemistry).

### Sample Collection for Immunogenicity, Biodistribution, and Histopathology

Blood samples (serum) for anti-AAV9 or anti-GFP antibodies were collected once during the pre-dose phase, once on Days 15 and 22, and on the day of scheduled euthanasia (Day 28). CSF was collected for assessment of antibodies against AAV9 or GFP on Day 1 of the dosing phase, once for all animals on Day 15 from the ICM, and on the day of scheduled euthanasia for respective cohorts from the ICM. On Study Day 28, the day of scheduled euthanasia, protocol-specified tissue samples were frozen for biodistribution analysis and/or GFP protein quantification or were collected for formalin fixation and paraffin embedding for microscopic evaluation and/or molecular pathology, as presented in **Supplemental Table 2**.

### Droplet Digital Polymerase Chain Reaction Analysis

Specified tissues were collected using instruments cleaned with sterile saline and/or sterile instruments to avoid contamination (see **Supplemental Table 2**). Samples for ddPCR or MSD protein analysis were collected, flash frozen in liquid nitrogen, and stored between –60°C to – 80°C until packed on dry ice and shipped for analysis. From each study animal, 38 tissue samples were collected for analysis of scAAV9-CB-GFP vector genome content using ddPCR quantitation with a C1000 Thermal Cycler (Bio-Rad). Specific primers and probes used for the ddPCR analysis are described in the Supplementary Methods. In parallel with quantifying scAAV9-CB-GFP vector genomes, a two-copy reference gene (*CFTR*) was also quantified for normalizing purposes. *CFTR* primers and probe were added in the master mix along with scAAV9-CB-GFP primers and probe, and scAAV9-CB-GFP genomes and *CFTR* genes were quantified in the same reaction using multiplex ddPCR. The resulting values of scAAV9-CB-GFP were presented as vg/diploid genome, thus normalizing vector genomes to the *CFTR* reference gene. The limit of quantitation determined for onasemnogene abeparvovec vector genome in 100× diluted tissue homogenates, 26 copies/reaction, was used in this study because scAAV9-CB-GFP shares the same non-transgene components of the expression cassette. It is not possible to express the lower limit of quantitation as viral genome copies/diploid genome, but only as a limit of copy numbers per reaction.

### Meso Scale Discovery Electrochemiluminescence Immunosorbent Assay for GFP Protein Quantification

Green fluorescent protein expression was quantified using MSD ECLIA. For each tissue sample, 250 µL of pre-chilled complete protein extraction buffer (pre-chilled T-PER tissue extraction reagent [ThermoFisher Scientific] with 1×Halt protease inhibitor [ThermoFisher Scientific]) was added per sample tube. A 5-mm bead was added into each tube, and the tissues were homogenized using a TissueLyser (Qiagen) at 30 Hz for 2 minutes. After centrifugation at 10,000×g and 4°C for 20 minutes, protein lysates were transferred to a new tube and stored at –20°C until analysis. Protein was quantified with the Rapid Gold BCA Protein Assay Kit (ThermoFischer Scientific) according to manufacturer’s protocol.

For GFP quantification, wells were coated by adding 25 µL of mouse anti-GFP monoclonal capture antibody (diluted 10 µg/mL in phosphate-buffered saline [PBS]) overnight at 4°C. After washing three times with 250 µL of PBS-T wash buffer (0.5% Tween-20/PBS), 150 µL of MSD Blocker A was added in each well and incubated on a shaker set to 500 rpm. After shaking at 500 rpm for 1 to 2 hours at room temperature, 25 µL of prepared standards (serial dilutions of GFP recombinant protein) or samples were added to each well. After being shaken for 1 to 2 hours at room temperature, the wells were washed three times with PBS-T and incubated with 25 µL of rabbit anti-GFP polyclonal antibody detection antibody diluted 1:10,000 in antibody dilution buffer (1.5 mg/mL mouse gamma globulin, 2.5% bovine serum albumin, 0.025% Tween- 20/PBS) and incubated shaking for 1 to 2 hours at room temperature. The wells were washed three times with PBS-T and then incubated in MSD SULFO-TAG anti-rabbit secondary antibody (Meso Scale Diagnostics) diluted to 0.5 µg/mL in antibody dilution buffer for 1 hour while shaking at room temperature. After washing three times with PBS-T, 150 µL of read buffer was added to each well, and the plates were analyzed using a MESO QuickPlex SQ 120 (Meso Scale Diagnostics). Recombinant GFP was serially diluted prior to assaying by MSD to determine the limit of quantitation and to plot a standard curve across all plates. The limit of quantification was determined to be 0.024 ng/mL GFP protein in the assay.

### Microscopic Pathology Evaluation

Selected tissues collected at necropsy (see **Supplemental Table 2**) were fixed in 10% neutral buffered formalin for at least two but not more than three overnights following collection (up to 72 hours). Tissues were processed routinely to paraffin block and sectioned at a nominal thickness of approximately 4 µm, stained with hematoxylin and eosin, and evaluated microscopically by an American College of Veterinary Pathologists (ACVP) board-certified anatomic veterinary pathologist experienced in toxicologic pathology and neuropathology. The anatomic pathology results, including macroscopic and microscopic observations, were reviewed by a second ACVP board-certified, similarly qualified and experienced anatomic pathologist, and consensus was reached between the study pathologist and the reviewing pathologist on nomenclature, severity grades of microscopic findings, and interpretation of the findings.

### Molecular Pathology

Immunohistochemistry staining for GFP, including the deparaffinization and antigen retrieval steps, was performed on a Ventana Discovery XT autostainer using standard Ventana Discovery XT reagents (Ventana Medical Systems). Slides were deparaffinized, then submitted to heat-induced antigen retrieval by covering them with cell conditioning 1 (pH 8) solution according to the standard Ventana retrieval protocol. Slides were incubated with the primary rabbit monoclonal anti-GFP antibody (clone EPR14104-89 [Abcam]; 0.372 µg/mL) or a non-immune isotype-matched control (rabbit monoclonal immunoglobulin G [IgG] clone DA1E [Cell Signaling Technology]; 0.372 µg/mL) for 1 hour. Visualization was obtained by incubation with the appropriate Discovery OmniMap horseradish peroxidase (HRP; Ventana Medical Systems) reagent followed by Discovery ChromoMap DAB (Ventana Medical Systems). Counterstaining was performed using hematoxylin (Ventana Medical Systems) and bluing reagent (Ventana Medical Systems) for 4 minutes each. Slides were dehydrated, cleared, and coverslips applied with a synthetic mounting medium. Slides were evaluated by the protocol-specified contributing scientist for Molecular Localization, an ACVP board-certified anatomic veterinary pathologist (KGM) experienced in toxicologic and molecular pathology. The slides were assigned negative, minimal, mild, moderate, or strong reactivity scores based on absence of staining, <1% of the area stained, 1% to 10% of the tissue stained, 11% to 50% of the tissue stained, or >50% of the tissue stained, respectively. The cell types stained were reported by the evaluating pathologist as a qualitative evaluation in addition to the semiquantitative reactivity score described.

Image analysis was performed using the HALO platform (Indica Labs; v3.0.311.149) on 20× digital magnification whole slide images (WSI) scanned on an Aperio AT2 scanner (Leica Biosystems, Inc.). The tissue section in the WSI was manually annotated to remove non-specific background staining. Area quantification algorithm v2.1.3 using pixel-based deconvolution was optimized to positive GFP IHC signal and run-on annotated images. Results were based on positive signal normalized to total tissue area, resulting in percentage positive signal/total area. Analysis and graphs were performed on GraphPad Prism version 8.1.2.

To detect GFP antisense (AS) and sense (S) sequences specific to the AAV vector genome and GFP transcripts, as well as in *Macaca fascicularis*-peptidylprolyl isomerase B (PPIB) (AS) (included as routine positive control and tissue quality control) and dihydrodipicolinate reductase (DapB) (AS) (included as routine negative control) genes, ISH was performed on tissue sections from select FFPE tissue blocks using reagents and equipment supplied by Advanced Cell Diagnostics, Inc., and Ventana Medical Systems, Inc. The ISH RNAscope^®^ probes were designed by Advanced Cell Diagnostics, Inc. Positive PPIB and negative DapB control probe sets were included to ensure mRNA quality and specificity, respectively. The hybridization method followed protocols established by Advanced Cell Diagnostics, Inc., and Ventana Medical Systems, Inc., using mRNA RED chromogens. Briefly, 5-µm tissue sections were baked at 60°C for 60 minutes and used for hybridization. The deparaffinization and rehydration protocol was performed using a Tissue-Tek DR5 stainer (Sakura) with the following steps: three times xylene for 5 minutes each; two times 100% alcohol for 2 minutes; and air dried for 5 minutes. Off-line manual pretreatment was carried out in 1× retrieval buffer at 98°C to 104°C for 15 minutes. Optimization was performed by first evaluating PPIB and DapB hybridization signal and subsequently using the same conditions for all slides. Following pretreatment, the slides were transferred to an Ultra autostainer (Ventana Medical Systems, Inc.) to complete the ISH procedure, including protease pretreatment, hybridization at 43°C for 2 hours, amplification, and detection with HRP and hematoxylin counterstain.

The unique attributes of nonclinical studies allow application of a series of “best practices” for optimal IHC and ISH results, including 1) the use of optimized/controlled fixation in 10% neutral buffered formalin for a limited and specified period of time. In this study, immersion fixation was completed for at least two but not more than three overnights (or up to 72 hours) following necropsy. This kind of precision is generally not feasible for (human) autopsy material. 2) The use of paraffin embedding followed by nominal 4- to 5- μm thick sections (performed for NHP studies, but often not possible for autopsy material). 3) The use of optimized monoclonal antibodies (for IHC) using chromogenic instead of fluorescent detection, because this eliminates the challenge of autofluorescence from neurons in the CNS. For ISH, specific RNAscope^®^ (Advanced Cell Diagnostics, Inc.) probes were added with both positive and negative control (PPIB and DapB, respectively) probes during each run.

### Statistical Analysis

The small number of animals per group precluded the use of extensive statistics beyond median and mean with or without standard deviations for most quantitative endpoints. No statistical analysis was determined for the antibody data. Data are summarized on individual animal basis. For vector DNA concentration and GFP concentration data, mean, median, minimum, and maximum concentrations for each analyte were calculated for each tissue and dose group. For the purpose of calculation of these statistics and accompanying visualization (**Figures 1** and **2**, as well as **Supplemental Figures 1-6**), concentrations that were below the lower limit of quantification were set equal to 0. If the mean or median value was less than the lower limit of quantification, the result was reported as below limit of quantitation. For image analysis of GFP IHC staining, differences in percent pixel positive area between the intrathecal-LP and ICM groups were compared using multiple Mann-Whitney tests of group analysis.

**Figure 1.**
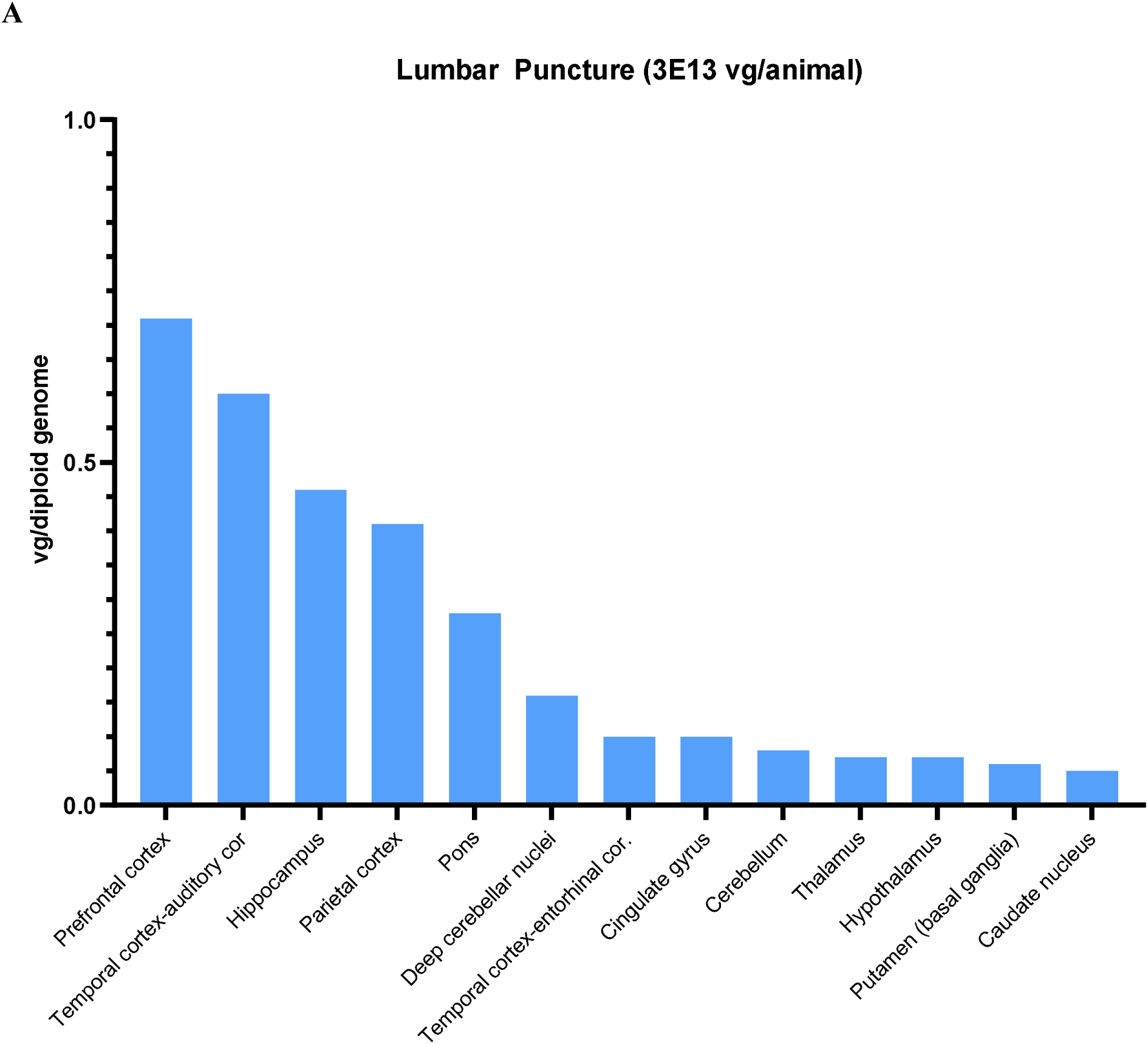
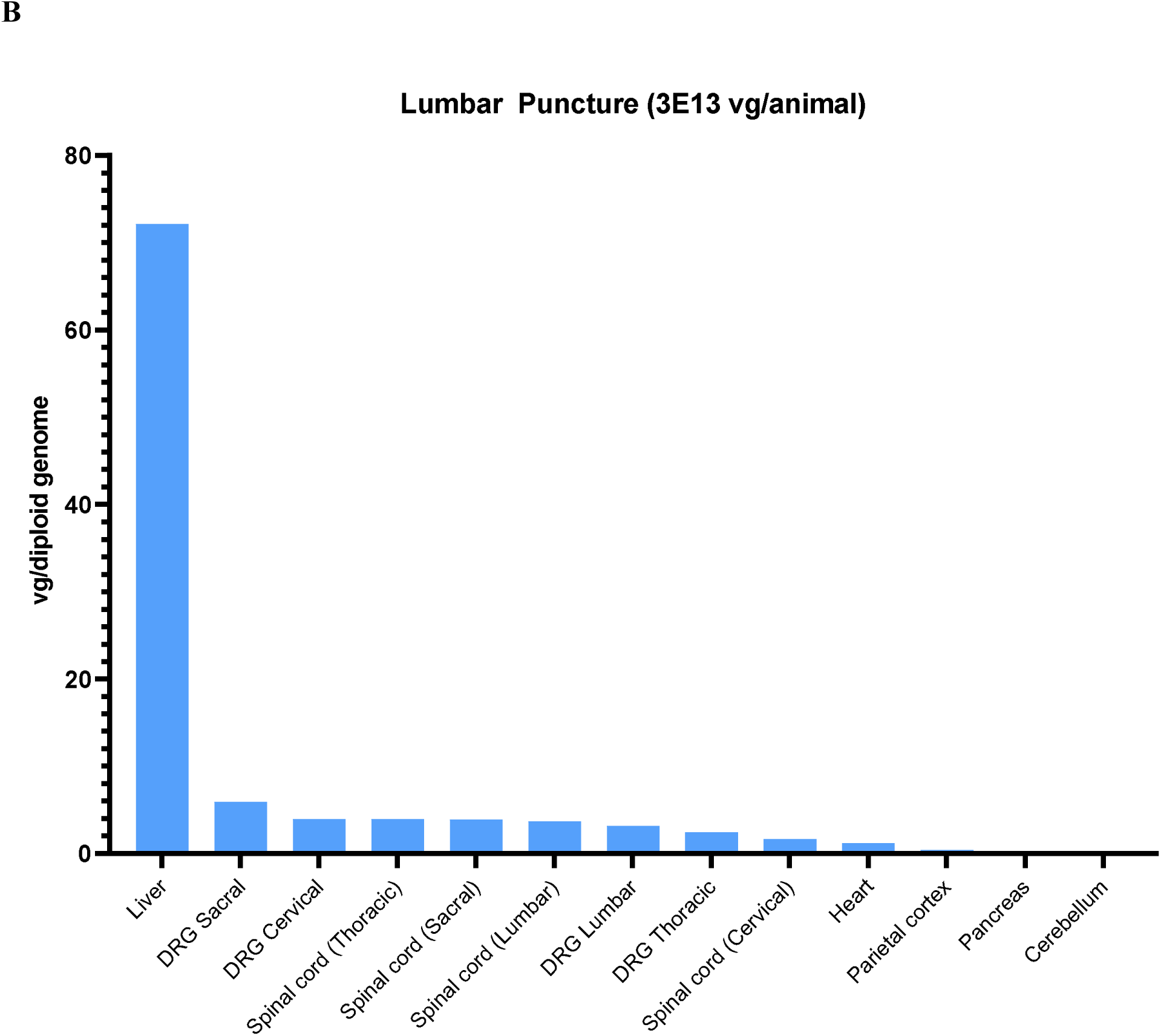
Biodistribution of scAAV9-CB-GFP vector genomes normalized to reference gene, *CFTR,* in tissues and red blood cell pellets of treated monkeys at 28 days post- injection via intrathecal-LP. NHPs were dosed at 3.0×10^13^ vg/animal. (A) Select brain regions and (B) systemic tissues. For each tissue in each animal, values were averages of three technical replicates and calculated as vg/diploid genome. The median values were reported from four replicate animals for all tissues. Individual data are presented in the supplemental data. Values below the limit of quantitation were entered as zero for the purposes of calculating the median. *CFTR*, cystic fibrosis transmembrane conductance regulator; DRG, dorsal root ganglia; ICM, intracisternal magna; intrathecal-LP, intrathecal infusion by lumbar puncture; NHP, nonhuman primate; scAAV9-CB-GFP, self-complementary adeno-associated virus-9–chicken β fluorescent protein; vg, vector genomes.

**Figure 2.**
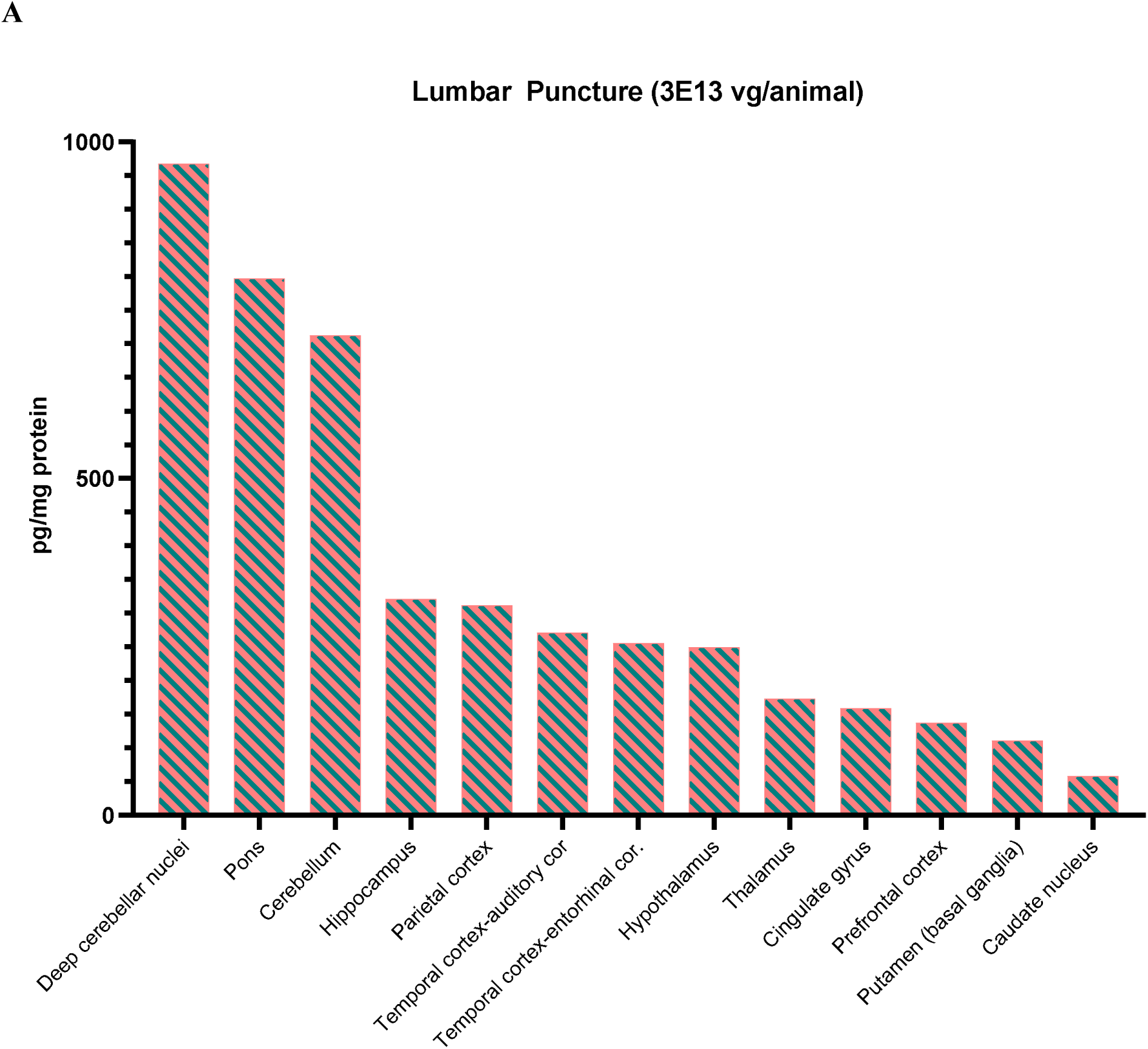
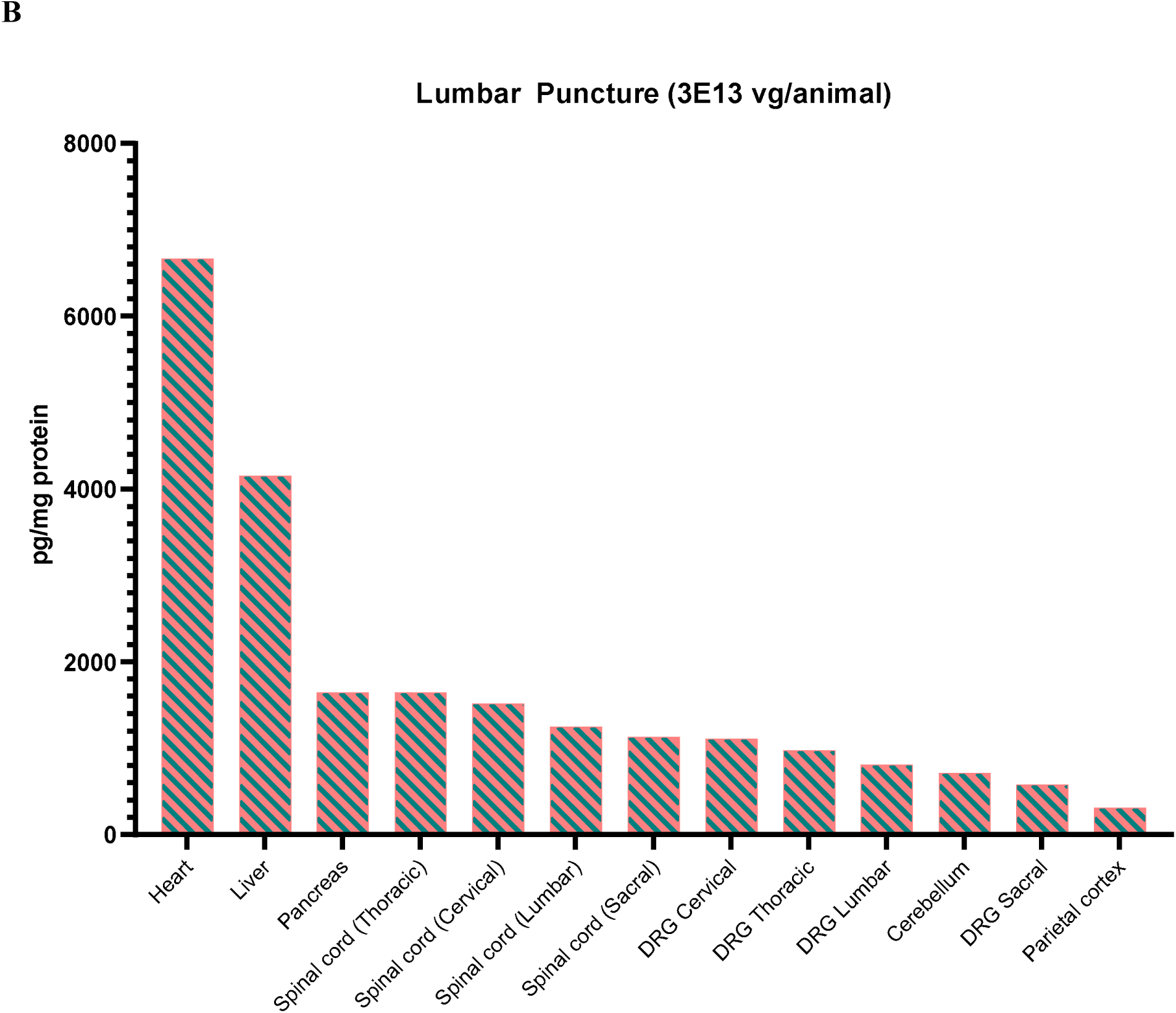
Biodistribution of GFP following intrathecal-LP of scAAV9-CB-GFP. NHPs received intrathecal-LP scAAV9-CB-GFP at 3.0×10^13^ vg/animal. (A) Select brain regions and (B) systemic tissues. GFP was quantified at 28 days post-injection as measured by ECLIA. For each tissue in each animal, values were averages of two technical replicates and calculated as picogram GFP per mg total protein. The median values were reported from four replicate animals for all tissues. Individual data are presented in the supplemental data. Values below the limit of quantitation were entered as zero for the purposes of calculating the median. DRG, dorsal root ganglia; ECLIA, electrochemiluminescence immunosorbent assay; ICM, intracisternal magna; intrathecal-LP, intrathecal infusion by lumbar puncture; NHP, nonhuman primate; scAAV9-CB-GFP, self-complementary adeno-associated virus-9–chicken β ctin–green fluorescent protein; vg, vector genomes.

## RESULTS

### Humoral Immune Response to AAV9 Capsid

Serum and CSF samples sent to Labcorp-Chantilly from all collected time points were evaluated for the presence of anti-AAV9 antibodies. Serum and CSF data are presented in **Supplemental Tables 3** and **4**, respectively. Anti-AAV9 antibodies were detected in the serum and CSF of a few animals. In the control group, the titers of anti-AAV9 antibodies did not meaningfully change after administration of vehicle (Groups 1 and 2). In all animals dosed with scAAV9-CB-GFP, there was a marked increase in anti-AAV9 antibody titer on Day 15 in both the serum and CSF compared to the pre-study sample. The titers of anti-AAV9 antibodies were lower in the CSF compared to the matching serum sample.

### Clinical Observations and Clinical Pathology

scAAV9-CB-GFP was well-tolerated when administered as a single dose of 1.0×10^13^ or 3.0×10^13^ vg/animal via intrathecal administration via LP or ICM injection to female cynomolgus monkeys 13 to 17 months of age. All animals survived to scheduled euthanasia (28 days post-dose). No scAAV9-CB-GFP–related changes in body weight, neurologic observations, or coagulation endpoints were observed for animals receiving up to 3.0×10^13^ vg/animal via either route of administration. For individual female animals administered 3.0×10^13^ vg/animal by either intrathecal-LP or ICM, scAAV9-CB-GFP was associated with general decreased activity, whole body tremors, hunched posture, or abnormal yellow periorbital color that correlated with clinical pathology changes that are outside the scope of the manuscript. Such observations were not present in animals administered intrathecal 1.0×10^13^ vg/animal scAAV9-CB-GFP (intrathecal-LP or ICM). The clinical chemistry findings were generally similar between the intrathecal-LP and ICM administration groups.

### Biodistribution (ddPCR)

Biodistribution of scAAV9-CB-GFP vector genomes was described from tissue samples collected from the scheduled euthanasia of cynomolgus monkeys following a single dose (1.0×10^13^ vg/animal or 3.0×10^13^ vg/animal) administered by intrathecal-LP or ICM route. Detailed individual animal scAAV9-CB-GFP DNA data (vg/diploid genome) and the associated summary statistics (mean, median, minimum, and maximum concentrations for cDNA vg/diploid genome for each tissue and dose group) are presented in the Supplemental Materials (**Supplemental Tables 5–18**). Graphs of sampled brain regions and of selected other nervous system tissues and systemic tissues are presented in **Figure 1**.

scAAV9-CB-GFP DNA was not detected in the tissues within the control (Groups 1 and 2) animals at 28 days post-dose (**Supplemental Tables 5, 6, 13, and 14**). In NHPs dosed via intrathecal-LP at 1.0×10^13^ vg/animal (Group 3), scAAV9-CB-GFP DNA was detected in all animals and tissues analyzed at 28 days post-dose except for selected brain regions consisting of the corpus callosum (all animals), the temporal-entorhinal cortex, occipital-primary visual cortex and cingulate gyrus (one of four animals), caudate nucleus (three of four animals), putamen (basal ganglia; two of four animals), temporal-auditory cortex of animal (two of four animals), parietal cortex (one of four animals), thalamus (two of four animals), hypothalamus (one of four animals), hippocampus (one of four animals), substantia nigra (three of four animals), cerebellum (two of four animals), deep cerebellar nuclei (two of four animals), and optic nerve (one of four animals) (**Supplemental Tables 7 and 15**). In animals administered scAAV9-CB-GFP via intrathecal-LP at 3.0×10^13^ vg/animal (Group 4), scAAV9-CB-GFP DNA was detected in all animals and tissues analyzed at 28 days post-dose except for selected brain regions of the temporal-entorhinal cortex, caudate nucleus, putamen (basal ganglia), hypothalamus and substantia nigra of one animal (**Supplemental Tables 8 and 16**).

In animals dosed via ICM at 1.0×10^13^ vg/animal (Group 5), scAAV9-CB-GFP DNA was detected in all animals and tissues analyzed at 28 days post-dose except for the corpus callosum and thalamus (all four animals), the temporal-entorhinal cortex (one of four animals), caudate nucleus (three of four animals), putamen (basal ganglia; two of four animals), cingulate gyrus (three of four animals), parietal cortex (one of four animals), hypothalamus (three of four animals), hippocampus (one of four animals), amygdala (two of four animals), substantia nigra (one of four animals), pons (two of four animals), occipital-primary visual cortex (one of four animals), deep cerebellar nuclei (two of four animals) (**Supplemental Tables 9 and 17**). In addition, scAAV9-CB-GFP DNA was not detected in the skeletal muscle of one female animal in this dose group. In animals administered 3.0×10^13^ vg/animal scAAV9-CB-GFP via ICM (Group 6), scAAV9-CB-GFP DNA was detected in all animals and tissues analyzed at 28 days post-dose except for the caudate nucleus in one animal (**Supplemental Tables 10 and 18**).

No marked, consistent differences in scAAV9-CB-GFP DNA concentrations were detected in tissues from animals administered scAAV9-CB-GFP into the CSF by intrathecal-LP or ICM injection at either 1.0×10^13^ or 3.0×10^13^ vg/animal, indicating no advantage for CNS biodistribution/transduction in the selection of one administration site over the other. Within the dose groups of 1.0×10^13^ or 3.0×10^13^ vg/animal, broad interanimal variability was noted in scAAV9-CB-GFP DNA concentrations observed at a single time point (28 days post-dose) for a particular tissue. In addition, variability in scAAV9-CB-GFP DNA concentrations was noted between different systemic and nervous system tissues and between different regions of the brain in the same animal (**Supplemental Tables 13–18**). Based on these differences, it was challenging to elucidate a clear relationship between tissue scAAV9-CB-GFP DNA concentrations and the administered number of vector genomes (total dose), particularly in a complex tissue like the brain. Broad themes can be gleaned from a comparison of median (and mean) concentration data (**Supplemental Tables 7–10**, and **12**), demonstrating that scAAV9- CB-GFP DNA concentrations tended to be greater for both routes of administration dosed at the 3.0×10^13^ vg/animal dose compared with the 1.0×10^13^ vg/animal dose.

**Figure 1** presents median scAAV9-CB-GFP DNA concentrations (vg/diploid genome, normalized with *CFTR* reporter gene) for animals administered 3.0×10^13^ vg/animal scAAV9- CB-GFP by intrathecal-LP. The highest concentration of scAAV9-CB-GFP DNA (vg/diploid genome) was in the liver of animals administered either 1.0×10^13^ or 3.0×10^13^ vg/animal scAAV9-CB-GFP by either intrathecal-LP or ICM and sampled (**Figure 1B; Supplemental Figures 1B, 2B, and 3B, Supplemental Tables 11 and 13–18**). Relatively high scAAV9-CB-GFP DNA concentrations were also observed in the DRG (cervical, thoracic, lumbar, and sacral) and TG, as well as the spinal cord (all regions), and mandibular lymph node. High scAAV9-CB-GFP DNA concentrations, or transduction, observed in the liver, spinal cord, and DRG was similar to published studies by other laboratories using intrathecally administered AAV9 gene therapies.^3,4^ In contrast, the lowest scAAV9-CB-GFP DNA concentrations in dosed animals, regardless of dose (1.0×10^13^ or 3.0×10^13^ vg/animal) or route of administration (intrathecal-LP or ICM) were in the brain, particularly in brain regions that tended to be located away from (or deep relative to) meningeal surfaces, including the caudate nucleus, deep cerebellar nuclei, putamen, and cingulate gyrus.

Notably, while scAAV9-CB-GFP DNA was detected above the limit of quantification in the ovaries of animals regardless of dose or route of administration, it was among the lowest concentration relative to all other evaluated tissues (**Supplemental Tables 11 and 13–18**). Homogenates prepared from the ovary were not indicative of results for germinal cells, as stromal and germinal cells were homogenized together and analyzed as a single tissue.

In summary, following administration directly into the CSF at doses of 1.0×10^13^ or 3.0×10^13^ vg/animal, scAAV9-CB-GFP distributed widely throughout the body, most notably to the liver, sensory ganglia (DRG and TG), and spinal cord in female cynomolgus monkeys that were observed for 28 days (**Figure 1 and Supplemental Figures 1-3**). scAAV9-CB-GFP DNA concentration (vg/diploid genome) was generally proportional to dose (**Supplemental Table 12**), although tremendous inter-animal variability was observed within each dose group. For both routes of administration, consistent median concentration ratios were observed (IT= 3.33; ICM= 2.79) between the low dose (1.0×10^13^ vg/animal) and high dose (3.0×10^13^ vg/animal). No marked, consistent differences in scAAV9-CB-GFP DNA concentrations were detected when dosed intrathecally by intrathecal-LP or ICM injection.

### Protein Analysis (MSD ECLIA) for GFP

The biodistribution of scAAV9-CB-GFP was also determined by MSD ECLIA for GFP quantification as a measure of transgene expression. Detailed individual animal GFP protein data (pg GFP/mg total protein) and the associated summary statistics (mean, median, minimum, and maximum concentration for GFP protein concentrations for each tissue and dose group) are presented in the Supplemental Materials (**Supplemental Tables 19–32**). GFP expression was detected in all tissues examined at both doses and routes of administration, with a dose-dependent response observed specifically in most brain regions (**Figure 2; Supplemental Table 26; Supplemental Figures 4 6**. A dose response was not discernible in most other tissues (**Figure 2; Supplemental Figures 4 6**). Similar to the pattern observed for scAAV9-CB-GFP DNA quantified by ddPCR, the route of administration did not affect transgene protein expression patterns. The tissues with greater concentrations of GFP protein included the heart, liver, pancreas, and skeletal muscle (both biceps femoris and diaphragm). In comparison, very low and highly variable GFP protein concentrations were noted in multiple regions of the brain parenchyma at both doses and routes of administration (**Figure 2A** and **Supplemental Figures 1A, 2A, and 3A**). Samples from vehicle-control animals had no detectable GFP (**Supplemental Tables 19, 20, 27**, and **28**).

### Microscopic Findings

scAAV9-CB-GFP–related microscopic findings were observed in the nervous system (brain, spinal cord, DRG, and peripheral nerves), as well as the liver and heart of animals at both doses and routes of intrathecal administration (intrathecal-LP and ICM). scAAV9-CB-GFP–related microscopic findings in the brain included gliosis of the white and gray matter, mononuclear or mixed-cell infiltrates, and axon and/or neuron degeneration (**Supplemental Table 33**). In the spinal cord, scAAV9-CB-GFP-related axon degeneration, gliosis of the white and/or gray matter, and/or mononuclear cell infiltrates were observed (**Supplemental Table 33**). scAAV9-CB-GFP– related microscopic findings in the DRG included neuron degeneration/necrosis and mononuclear cell inflammation, increased satellite glial cell/neuron cell loss, and/or mononuclear cell infiltrates (**Supplemental Table 33**). In the cauda equine, axon degeneration and gliosis were observed only at the intrathecal-LP injection site (**Supplemental Table 33**) and may have been related to the route of administration in addition to the test article. scAAV9-CB-GFP– related microscopic findings in peripheral nerves included axon degeneration in the sciatic, tibial, sural, radial, and ulnar nerves, and mononuclear cell infiltrates in the sciatic nerve. In the context of this study, differentiating whether these findings were related to scAAV9-CB-GFP or to expression of the foreign transgene GFP protein was not possible, although some microscopic findings, such as those in the DRG and TG, have been reported with other scAA9 gene therapy products administered by intravenous or intrathecal routes of administration in rhesus or cynomolgus monkeys.^3–6^

In the liver and heart, scAAV9-CB-GFP–related microscopic findings of mononuclear cell infiltrates and/or inflammation were observed in animals administered 1.0×10^13^ or 3.0×10^13^ vg/animal via intrathecal-LP or ICM injection (**Supplemental Table 33**). In the liver, scAAV9- CB-GFP–related hepatocyte necrosis was also noted only in animals administered 3.0×10^13^ vg/animal via intrathecal-LP and oval cell hyperplasia was observed in animals administered either 1.0×10^13^ or 3.0×10^13^ vg/animal via intrathecal-LP. Generally, scAAV9-CB-GFP–related microscopic findings in the nervous system, and other tissues occurred with similar incidences and severities in animals administered scAAV9-CB-GFP intrathecally via intrathecal-LP or ICM injection.

### Molecular Pathology (IHC and ISH)

Molecular pathology analysis was performed on tissues from animals administered 3.0×10^13^ vg/animal via intrathecal-LP or ICM routes across the evaluated regions.

#### Central nervous system

To characterize the distribution of GFP expression in the CNS, IHC was performed on blocks of brain, spinal cord, and lumbar DRG and scored for the degree of staining (**Supplemental Table 34**). Staining of control tissue produced no signal, and no non-specific signal was observed with rabbit IgG control antibody. Compared with lumbar DRG and spinal cord (**Figure 3**), overall lower and more variable GFP protein expression was detected in sections of brain from all evaluated animals administered scAAV9-CB-GFP (3.0×10^13^ vg/animal) by either the intrathecal-LP or ICM routes. In the cerebral cortex, GFP-positive cells were identified multifocally, often in a perivascular pattern (**Figure 4**). Most positive cells were morphologically consistent with astrocytes, and double-label experiments confirmed that the majority of GFP-positive cells were also positive for glial fibrillary acidic protein (**Figure 5**). In the cerebellum, multifocal expression in the molecular layer, consistent with Bergman glia, a diversified astrocyte subtype, was observed in cells (**Figure 6**). Only rare GFP-positive Purkinje neurons and granular cell layer neurons were observed. GFP expression was not detected in neurons of the deep cerebellar nuclei.

**Figure 3.**
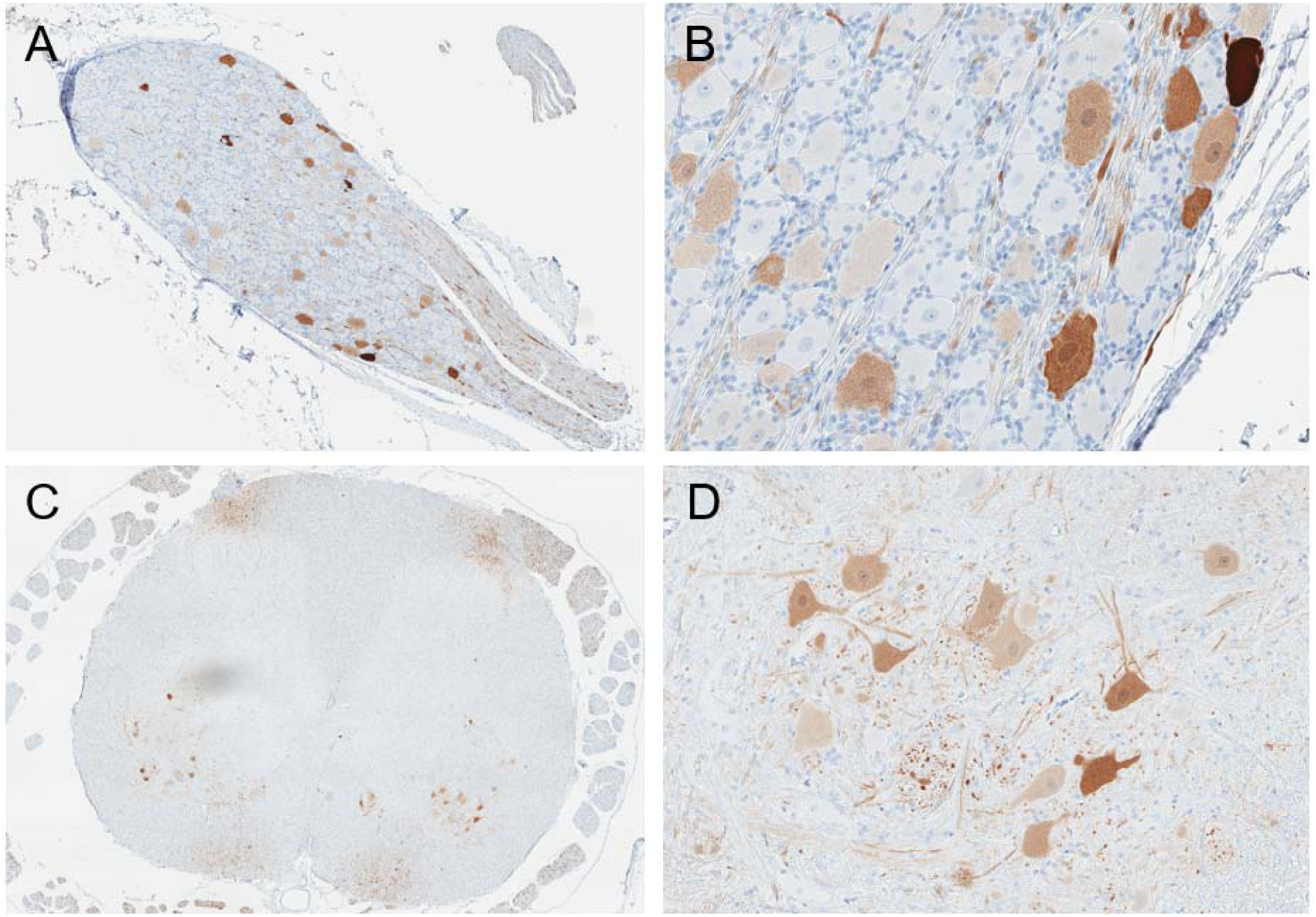
Immunohistochemistry for GFP expression in lumbar DRG (A and B) and lumbar spinal cord (C and D) in an NHP dosed by intrathecal-LP administration with 3.0×10^13^ vg/animal of scAAV9-CB-GFP. DRG, dorsal root ganglion; GFP, green fluorescent protein; LP, lumbar puncture; NHP, nonhuman primate; scAAV9-CB-GFP, self-complementary adeno-associated virus-9–chicken β actin–green fluorescent protein; vg, vector genomes.

**Figure 4.**
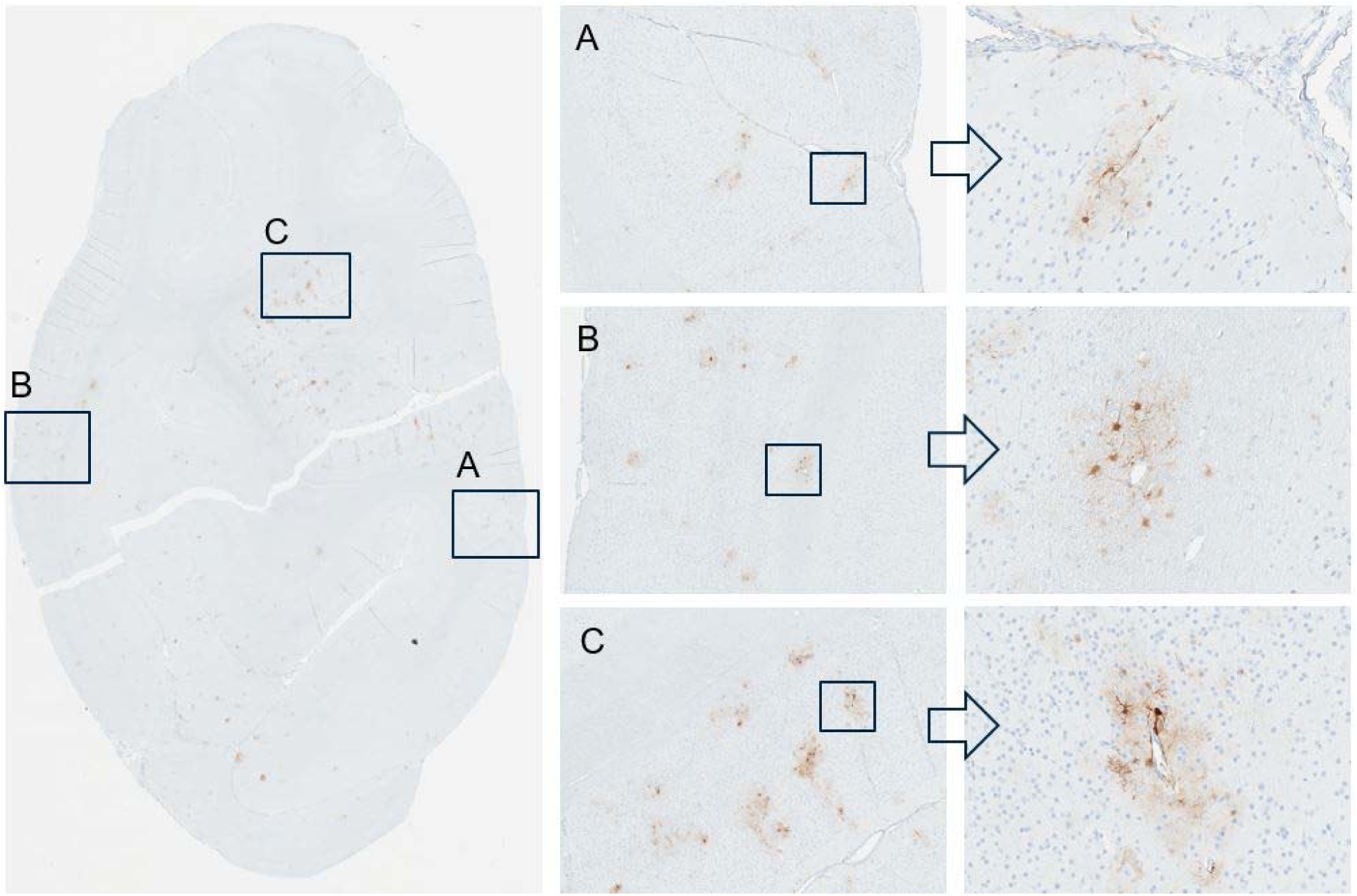
Immunohistochemistry for GFP performed on occipital cortex in an NHP dosed by intrathecal-LP administration with 3.0×10^13^ vg/animal of scAAV9-CB-GFP. Panels A, B and C represent greater magnifications demonstrating green fluorescent protein localization to perivascular astrocytes. Intrathecal-LP, intrathecal infusion by lumbar puncture; NHP, nonhuman primate; scAAV9-CB-GFP, self-complementary adeno-associated virus-9–chicken β-actin–green fluorescent protein; vg, vector genomes.

**Figure 5.**
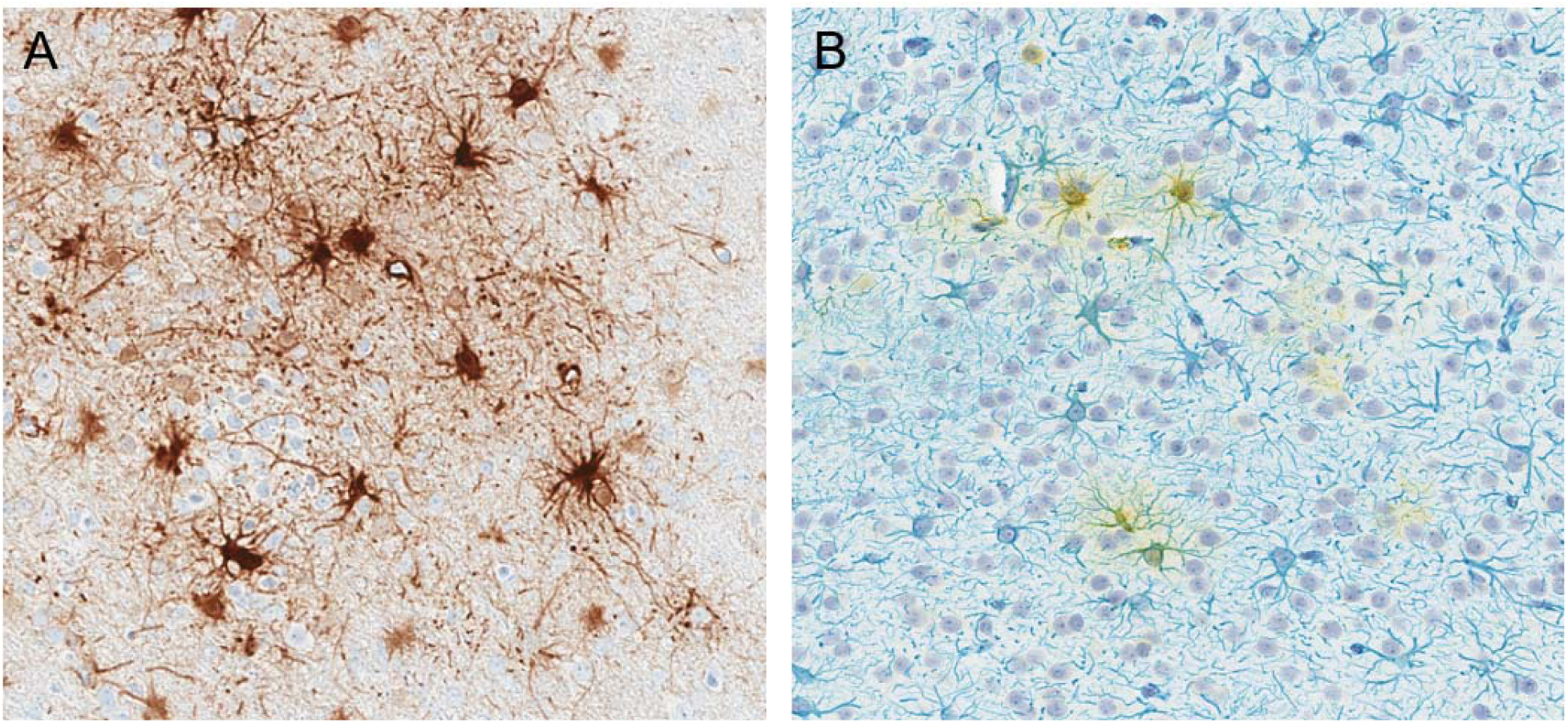
Astrocytic morphology of GFP-positive cells. Single-label GFP immunohistochemistry with DAB chromogen (A). Double immunohistochemical label for GFP (yellow) and the astrocytic marker GFAP (blue) (B). DAB, 3,3′ diaminobenzidine; GFAP, glial fibrillary acidic protein; GFP, green fluorescent protein.

**Figure 6.**
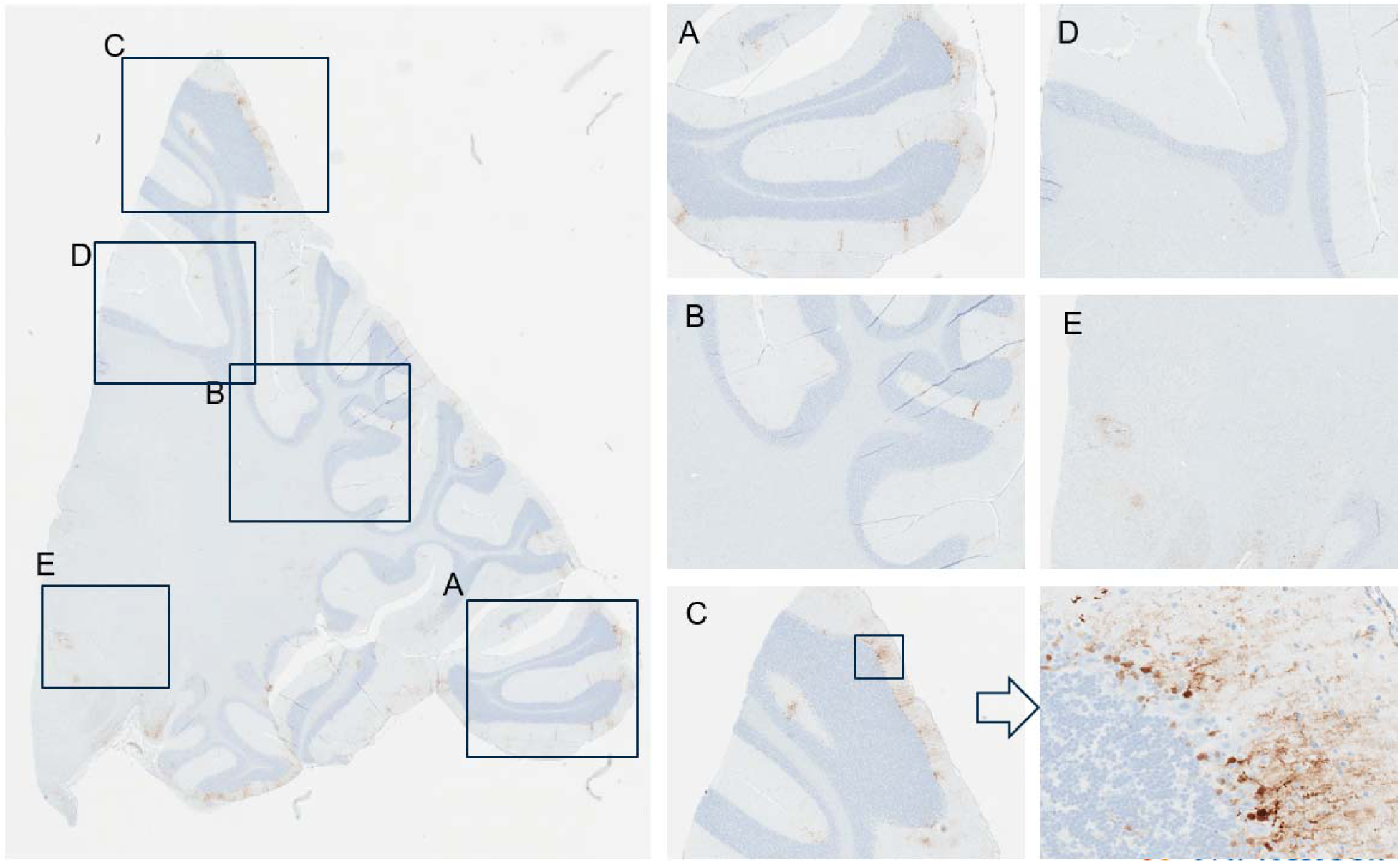
Immunohistochemistry for GFP performed on cerebellum and deep cerebellar nuclei from an NHP dosed by intrathecal administration with 3.0×10^13^ vg/animal of scAAV9-CB-GFP. Panels A, B, C, and D represent greater magnifications demonstrating multifocal green fluorescent protein localization and expression in cells consistent with Bergman glia cells. NHP, nonhuman primate; scAAV9-CB-GFP, self-complementary adeno-associated virus-9–chicken β-actin–green fluorescent protein; vg, vector genomes.

Quantitative image analysis was performed on sections of brain, spinal cord (lumbar), and DRG (sacral) stained for GFP by IHC and reported as percent GFP-positive pixels (**Supplemental Figure 7**). GFP expression was greatest in the spinal cord and DRG with lower staining intensity and therefore lower expression detected in the brain. No statistically significant differences were observed between animals administered 3.0×10^13^ vg/animal via intrathecal-LP or ICM routes across the evaluated regions.

Probes (S and AS) specific for GFP were utilized for ISH in order to identify vector genome sequences in selected regions of the brain. These probes detected similar patterns of vector localization compared with IHC for GFP and often revealed signal in a vascular and perivascular pattern (**Figure 7**). Differences were not observed between intrathecal-LP– and ICM–dosed animals.

**Figure 7.**
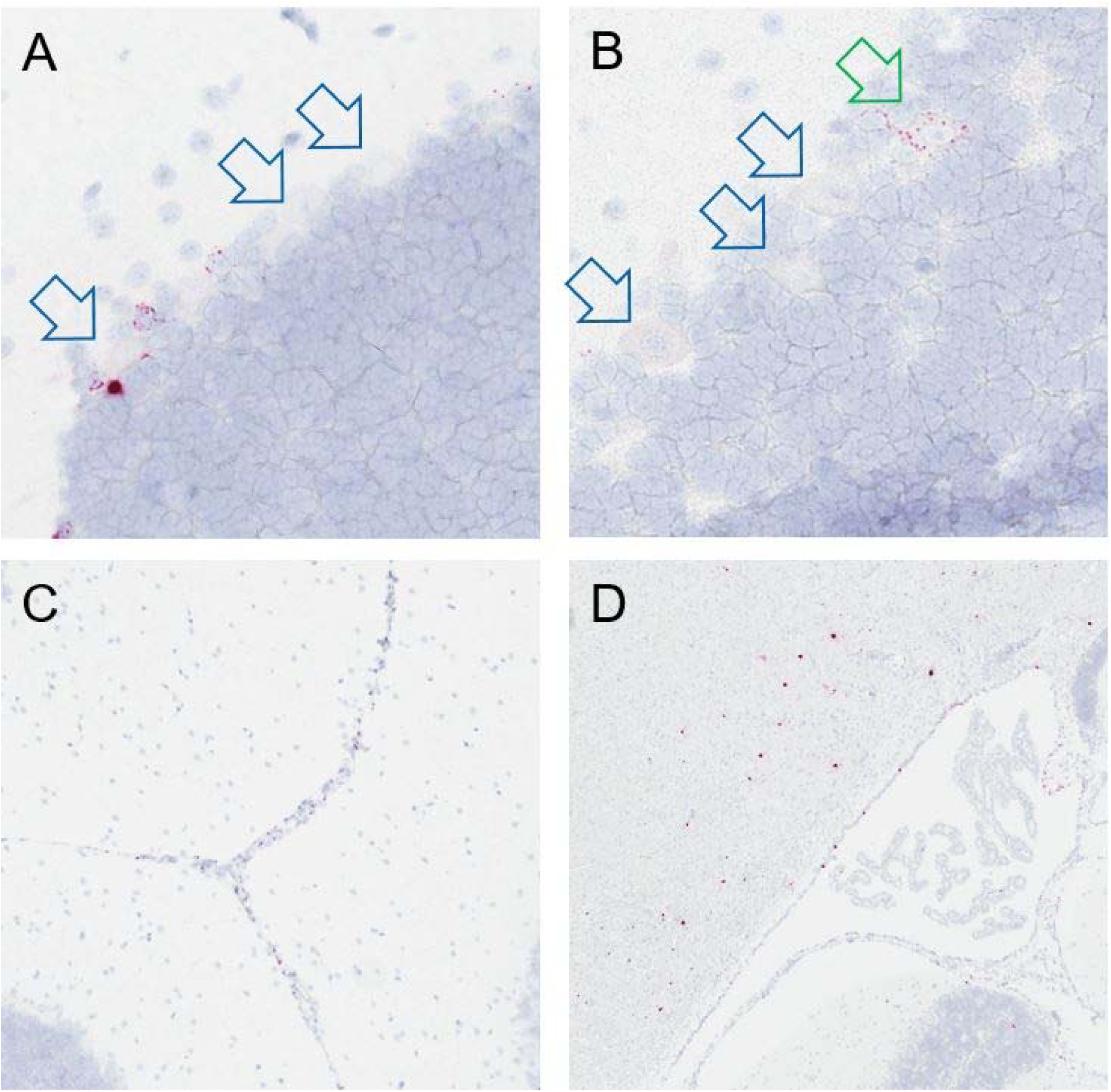
*In situ* hybridization for vector nucleic acid using antisense probes performed on cerebellum in an NHP dosed by intrathecal-LP administration with 3.0×10^13^ vg/animal of scAAV9-CB-GFP. The majority of Purkinje cell neurons (A and B) were negative (open blue arrows) and only rare cells demonstrated were positive (open green arrow). Perivascular and periventricular signal was also evident (C and D).

#### Peripheral tissues

GFP expression was analyzed by IHC in a range of peripheral tissues from animals dosed with 3.0×10^13^ vg/animal scAAV9-CB-GFP by the intrathecal-LP route. Greater staining intensity consistent with greater protein expression was detected in hepatocytes, striated myocytes (skeletal muscle), and cardiomyocytes (**Supplemental Figure 8**). IHC performed on tissues from a control animal produced no signal, which is the expected result, consistent with the high specificity of the assay (**Supplemental Figures 9–11**).

*In situ* hybridization using probes specific for GFP was performed on liver, heart, and skeletal muscle, and demonstrated greater concentrations of transgene mRNA in these tissues (**Supplemental Figure 12**), which is consistent with protein expression IHC data and the MSD ECLIA protein quantification data. Despite the intrathecal route of administration, the concentrations of GFP mRNA and protein expression were greater in these peripheral tissues compared with multiple regions of the brain and were concordant with GFP DNA and protein biodistribution data generated by ddPCR and MSD, respectively.

More tissues were examined by IHC from intrathecal-LP–dosed animals (3.0×10^13^ vg/animal) (**Supplemental Figures 13 and 14**) and are summarized in **Supplemental Table 35**. Signal was evident within germinal centers of the spleen and lymph nodes and to a lesser extent gut/bronchiolar-associated lymphoid tissue. The pattern of staining and the morphology of the transduced cells were consistent with follicular dendritic cells. Expression patterns were often variable within different cell types of a single organ. For example, greater degrees of expression in the kidney were observed in renal tubular epithelial cells associated with the juxtaglomerular apparatus, moderate expression was observed in renal medullary cells, and minimal expression was observed in the renal glomerulus and remaining renal cortical cells. In addition, moderate degrees of expression were detected in brown adipocytes, the parathyroid gland, adrenal cortical cells, and exocrine and endocrine pancreatic cells. Lesser degrees of expression were observed in the thyroid gland and stomach, and minimal expression was observed in the lung, aorta, adrenal medullary cells, esophageal mucosa, small intestine, large intestine, thymus, lymph node non-follicular cortex and medulla, hepatic portal and sinusoidal cells, white adipocytes, bone marrow, and intraocular structures.

## DISCUSSION

This nonclinical biodistribution and toxicity (safety) study in female cynomolgus monkeys (NHPs) 13 to 17 months of age was conducted to elucidate and characterize the biodistribution of scAAV9-CB-GFP to brain, spinal cord, and peripheral (non-CNS) tissues when administered directly into the CSF (intrathecal) by either intrathecal-LP or ICM routes.

In all animals dosed with scAAV9-CB-GFP, there was a marked increase in anti-AAV9 antibody titer on Day 15 in both the serum and CSF compared with the pre-study sample. The titers of anti-AAV9 antibodies were lower in the CSF compared to the matching serum sample.

Biodistribution of DNA by ddPCR and GFP protein by MSD ECLIA, as well as molecular localization on tissues from animals dosed by the intrathecal route (intrathecal-LP and ICM) with 1.0×10^13^ vg/animal or 3.0×10^13^ vg/animal scAAV9-CB-GFP, were generally concordant and convergent. This demonstrated robust GFP DNA, mRNA, and protein expression in several peripheral tissues, including liver, skeletal muscle, and heart, as well as in the spinal cord (lower motor neurons) and sensory ganglia neurons (DRG and TG). In marked contrast, only minimal expression of these endpoints was detected in diverse regions of the brain despite extensive sampling. These results indicate vector distribution from the CSF primarily to systemic tissues and spinal cord, as well as sensory ganglia neurons, with only limited distribution to the brain parenchyma. This is consistent with Pardridge,^7^ who emphasized the potential misconception of CSF administration via these approaches as a means to access the brain parenchyma. Similar to the data presented in the current study, Pardridge^7^ noted that drugs within the CSF preferentially distribute to the systemic blood circulation with little penetration into deeper regions of the brain.

The current study demonstrates notable interanimal variability for biodistribution endpoints, including DNA and protein expression in NHPs, that is concordant with molecular localization results using IHC for GFP and ISH for the transgene sense and antisense sequences. This variability, present between animals in the same dose group and between tissues in the same animal, complicates understanding and reporting of biodistribution, as well as the relationship between administered dose and resultant biodistribution, and contributes further efforts to predict a potential human clinical dose in a therapeutic program using a clinically relevant transgene. It should be noted that small aliquots of tissue (∼25 mg) were analyzed and hence may not represent the concentration of vector DNA or GFP throughout the whole tissue. This may be a contributing factor to the variability in vector DNA and GFP concentrations between animals for a particular tissue, dose, or route of administration. It is possible that the sampling of the tissues is a greater source of variability than intrinsic differences between the animals. It should also be noted that separate aliquots were analyzed for vector DNA and GFP concentrations, which may confound the interpretation of the relative concentrations of vector DNA and GFP in a particular animal. Use of homogenized tissue in polymerase chain reaction assays may be supplemented by other techniques that permit evaluation and analysis of cellular patterns of distribution to provide a better understanding of variability within a tissue and the cell-specific tropism of the viral particle, which may be influenced by age, dose, route of administration, serotype, promoter, anesthesia, physical positioning, dose volume, and myriad other factors we are only beginning to understand. In comparing the intrathecal-LP and ICM routes of administration, molecular localization techniques were critical to visualize and understand the patchy nature of transduction in the brain, which trended toward multifocal, inconsistent subpial distribution up to 1 mm into the cortex and/or a distribution that favored the astrocytes around vessels, with relatively less GFP IHC staining (protein expression) or ISH RNAscope^®^ for AAV9 GFP mRNA expression in neurons within areas of interest, including the deep cerebellar nuclei and neuron-rich cortical gray matter regions deeper than 1 mm from the pial surface. This astrocytic pattern in the brain was in stark contrast to the substantial staining observed by both IHC and ISH in the lower motor neurons of the spinal cord and the DRG sensory neurons as well as a number of peripheral tissues, including skeletal muscle, heart, and liver.

The mechanism by which AAV vectors may distribute within the brain parenchyma following intrathecal dosing in NHPs has not been defined, but dosing into the CSF is not a direct route to expose neurons behind the blood brain barrier in the parenchyma of the brain. The glymphatic system is a recently recognized system by which CSF is drawn into the deeper regions of the brain along periarterial spaces formed by vessel-adjacent astrocytes, in which CSF may exchange with the interstitial fluid prior to exiting the brain in an equivalent perivenous space.^8^ This system is thought to play a major role in the movement of fluid and removal of macromolecules from the brain parenchyma.^9^ Larger particles, such as lipoproteins,^10^ which are of approximately equivalent size to AAV vectors, move through the glymphatic system. The GFP brain distribution patterns observed in the present study are consistent with limited diffusion of vectors across membranes that line the surface of the brain and of vector entry occurring primarily through glymphatic influx resulting in multifocal transduction and expression within perivascular astrocytes.^7, 11^

Based on the pattern of GFP expression in this study, a model of scAAV9 vector CNS and systemic distribution following intrathecal administration may be proposed (**Figure 8**). CSF is constantly produced with a turnover of approximately 5 hours in cynomolgus macaques and humans before it drains from the intrathecal space through arachnoid granulations, cranial nerves, and nerve roots after which it enters meningeal lymphatics and subsequently the systemic circulation.^6^ Based on quantified vector DNA copy number and the estimated number of averaged cell densities in these tissues,^12–15^ only 0.01% of the total 3.0×10^13^ vector dose is detectable in the brain at 28 days post-dosing compared with 1.3% of the total vector dose detected in the liver at the same time following intrathecal delivery. This differential transduction and quantification is consistent with the vast majority of vector draining from the intrathecal space to the systemic circulation prior to exchange with interstitial fluid in the brain parenchyma through glymphatic influx, convection, and/or diffusion.^7, 16, 17^ By enhancing glymphatic influx at the time of intrathecal dosing, greater concentrations of vector in the brain interstitial fluid may be achieved, resulting in improved and more uniform transduction of targeted cell types. Moreover, reduction in vector distribution to systemic organs may reduce adverse events associated with these tissues.

**Figure 8.**
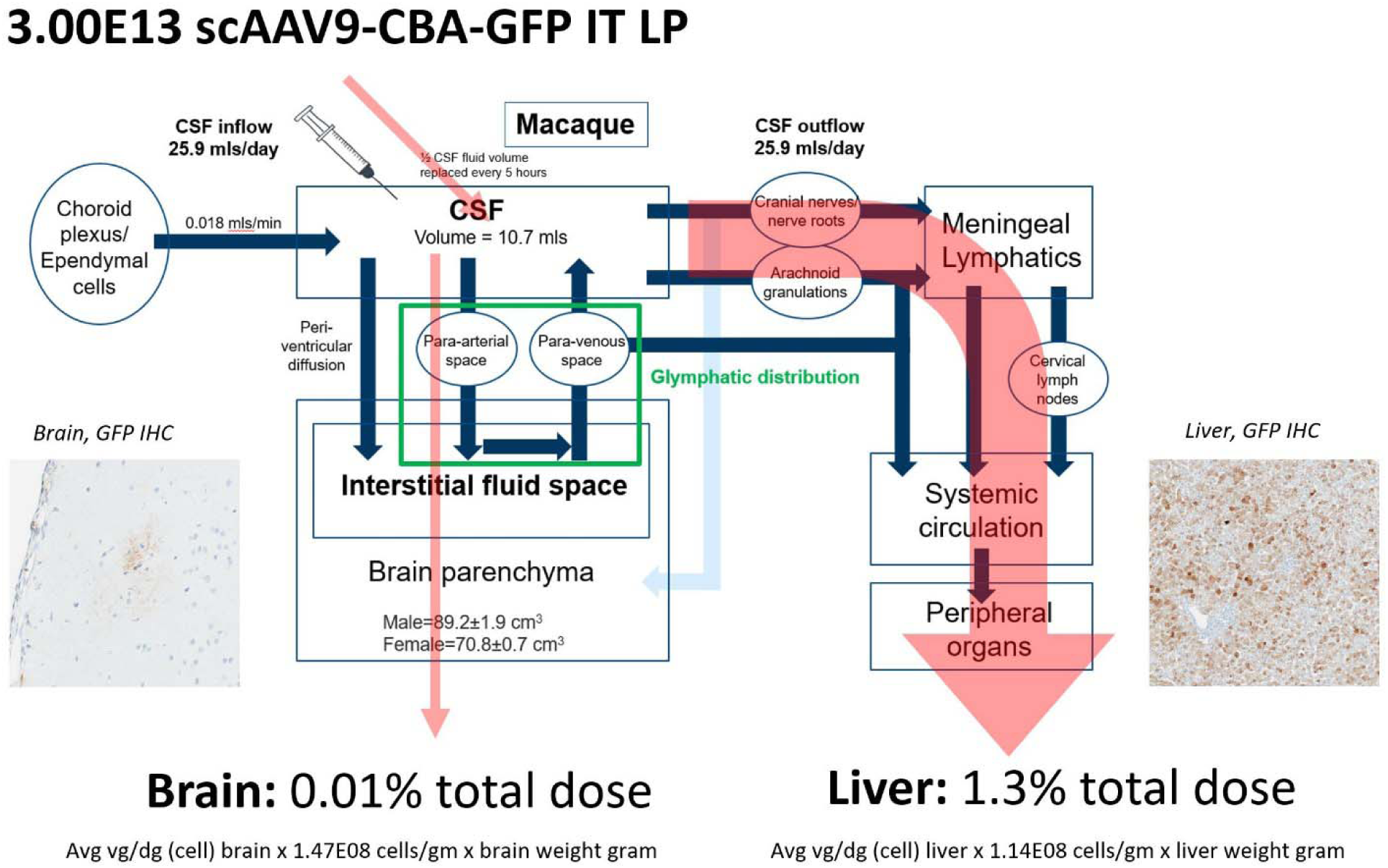
Model of AAV vector distribution following intrathecal administration by lumbar puncture to brain and systemic tissues in cynomolgus macaques. AAV vector transduction of brain requires distribution from the CSF to the interstitial fluid space of the brain parenchyma through glymphatic influx and limited periventricular diffusion. Because of rapid production and turnover of CSF, much of the intrathecal-delivered vector drains from the intrathecal space through arachnoid granulations, cranial nerves, and nerve roots to the systemic circulation in which transduction of non-CNS tissues may occur. The total amount of vector genomes in each organ was estimated for each animal based on 1) the average vg/diploid genome detected by ddPCR in each tissue of each NHP at necropsy (all the biodistribution values from the brain samples of each animal were averaged), 2) the reported number of cells per gram of liver or brain tissue^12–15;^ and 3) the total weight of liver and brain (in grams). The percentage of the total vector dose detectable in the brain or liver was then calculated as follows: (total amount of vector genomes detected in each tissue)/(initial dose administered) *100. AAV, adeno-associated virus; CNS, central nervous system; CSF, cerebral spinal fluid; GFP, green fluorescent protein; IT, intrathecal; LP, lumbar puncture; NHP, non-human primate.

In conclusion, widespread but variable biodistribution of scAAV9-CB-GFP was present in the spinal cord and sensory ganglia of the nervous system as well as the analyzed peripheral tissues (including liver, skeletal muscle, and heart) of animals administered up to 3.0×10^13^ vg/animal via intrathecal-LP or ICM injection. Poor biodistribution (i.e., low concentrations) in regions of the brain suggest that while vector genome readily transfers from the intrathecal CSF surrounding the spinal cord to the systemic circulation, there is only limited distribution to interstitial fluid of the brain parenchyma. Overall, these study data suggest the use of scAAV9-capsid gene therapy with these expression cassette elements may have limited diffuse neuronal gene expression in the brain when delivered intrathecally “downstream” in the vertebral column intrathecal space. This may be a limiting factor for its use as a clinical development tool for complex neurologic diseases requiring broad and diffused neuron transduction in the brain. This capsid/expression cassette combination may be better suited for diseases that are responsive to repair via a secreted protein or which do not require widespread brain neuronal transduction.

## Supporting information

Supplemental tables and data for Biodistribution, molecular pathology and pathology

## ACKNOWLEDGMENTS

The authors wish to acknowledge Thomas Larsen and Stephanie Cossette from Labcorp, of Chantilly, VA, and Madison, WI, USA, respectively, for their contributions to the study conduct. They also thank Maria Magnifico, Yanli Song, and Janet Do for their technical expertise; Andreas Kiessling and Fatih Ozsolak for their contributions to key scientific questions; Mark Milton for interpretation of immunogenicity and biodistribution data; and Kenard (Brian) Lee (all of Novartis, Novartis Institutes for BioMedical Research Biologics Center, or Novartis Gene Therapies) for image assistance and technical expertise.

## CONFLICTS OF INTEREST

**Emily K. Meseck** and **Cameron McElroy** are employees of Novartis Pharmaceuticals Corporation, East Hanover, NJ, USA. **Ghiabe Guibinga** is an employee of Novartis Institutes for BioMedical Research Biologics Center, San Diego, CA, USA. **Stephen Wang, Eloise Hudry,** and **Keith Mansfield** are employees of Novartis Institutes for BioMedical Research, Cambridge, MA, USA.

## FUNDING/SUPPORT

Writing and editing assistance, including preparation of a draft manuscript under the direction and guidance of the authors, incorporating author feedback, and manuscript submission, was provided by Marjet Heitzer, PhD, from Kay Square Scientific, Newtown Square, PA. This support was funded by Novartis Gene Therapies, Inc.

