## Supplemental tables and data for Biodistribution, molecular pathology and pathology for "Intrathecal sc-AAV9-CB-GFP: Systemic Distribution Predominates Following Single-Dose Administration in Cynomolgus Macaques"

**SUPPLEMENTAL MATERIAL**

**SUPPLEMENTAL METHODS**

**Droplet Digital Polymerase Chain Reaction Primers and Probes**

The following primers and probes were used for the droplet digital polymerase chain reaction (ddPCR) analysis: CB-GFP forward primer: 5′-GCAAATCAAAGAACTGCTCCTC-3′; CB-GFP reverse primer: 5′-GAACAGCTCCTCGCCCTTG-3′; CB-GFP probe: 5′-AGTGGATGTTGCCTTTACTTCTAGGCC-3′; monkey cystic fibrosis transmembrane conductance regulator (*CFTR*) forward primer: 5′-AGAGAGAAGGCTGTCCTTAGT-3’; monkey *CFTR* reverse primer: 5′-GAGTGTGTCATCAGGTTCAGA-3’; and monkey *CFTR* probe: 5′-TTCTGAGCAGGGAGAGGTGAT-3′.

**SUPPLEMENTAL TABLES**

**Supplemental Table 1.** **Study** **Design.**

| **Group^a^** | |  | **Number of Females** | **Test article** | **Route of administration** | **Dose concentration,** | **Targeted concentration,** |
| --- | --- | --- | --- | --- | --- | --- | --- |
|  |  |  |  |  |  | **vg/animal** | **vg/mL** |
| 1 (Control-Intrathecal-LP) | |  | 4 | VCA | Intrathecal-LP | 0 | 0 |
| 2 (Control-ICM) | |  | 4 | VCA | ICM | 0 | 0 |
| 3 (Low Intrathecal-LP) | |  | 4 | scAAV9-CB-GFP | Intrathecal-LP | 1.0×10^13^ | 1.0×10^13^ |
| 4 (High Intrathecal-LP) | |  | 4 | scAAV9-CB-GFP | Intrathecal-LP | 3.0×10^13^ | 3.0×10^13^ |
| 5 (Low ICM) | |  | 4 | scAAV9-CB-GFP | ICM | 1.0×10^13^ | 1.0×10^13^ |
| 6 (High ICM) | |  | 4 | scAAV9-CB-GFP | ICM | 3.0×10^13^ | 3.0×10^13^ |

Abbreviations: ICM, intracisterna magna; Intrathecal-LP, intrathecal lumbar puncture; scAAV9-CB-GF, self-complementary adeno-associated virus serotype 9–chicken β-actin promoter–green fluorescent protein; VCA, vehicle control article; vg, vector genomes.

^a^Groups 1 and 2 were administered vehicle control article only.

**Supplemental Table 2. Tissues Collected for Vector DNA, GFP Protein, and Microscopic Evaluation.**

| **Tissue** | **Frozen for vector DNA** | **Frozen for GFP protein** | **Fixed for tissue processing and microscopic evaluation** |
| --- | --- | --- | --- |
| Brain^a^, prefrontal cortex | X | X | X^b^ |
| Brain^a^ parietal cortex | X | X | X^b^ |
| Brain^a^ temporal cortex entorhinal cortex | X | X | X^b^ |
| Brain^a^ temporal cortex: auditory cortex | X | X | X^b^ |
| Brain^a^ occipital cortex: primary visual cortex | X | X | X^b^ |
| Brain^a^ cingulate gyrus | X | X | X^b^ |
| Brain^a^ corpus callosum | X | X | X^b^ |
| Brain^a^ cerebellum | X | X | X^b^ |
| Brain^a^ caudate nucleus (basal ganglia) | X | X | X^b^ |
| Brain^a^ putamen | X | X | X^b^ |
| Brain^a^ thalamus | X | X | X^b^ |
| Brain^a^ hypothalamus | X | X | X^b^ |
| Brain^a^ substantia nigra | X | X | X^b^ |
| Brain^a^ amygdala | X | X | X^b^ |
| Brain^a^ pons | X | X | X^b^ |
| Brain^a^ deep cerebellar nuclei | X | X | X^b^ |
| Brain^a^ hippocampus | X | X | X^b^ |
| DRG^c^ (spinal ganglia) to include dorsal spinal nerve root (cervical, thoracic, lumbar, sacral) | X | X | X^b^ |
| Trigeminal ganglia/nerve^a^ | X | X | X^b^ |
| Liver | X (left lateral lobe) | X (left lateral lobe) | X^b^ |
| Lung | X (accessory lobe) | X (accessory lobe) | X |
| Kidney | X (right pole) | X (right pole) | X |
| Ovary^a^ | X | X | X |
| Spleen | X | X | X |
| Heart | X (apex) | X (apex) | X |
| Lymph node, mandibular | X | X | X |
| GALT | N/A | N/A | X |
| Skeletal muscle (biceps femoris) | X | X | X |
| Pancreas | X | X | X |
| Spinal cord (cervical, thoracic, lumbar, sacral) to include injection sites | X | X | X |
| Eye^a^ | X | X | X^d^ |
| Optic nerve^a^ | X | X | X^d^ |
| Adrenal | N/A | N/A | X |
| Aorta | N/A | N/A | X |
| Bone, femur with marrow | N/A | N/A | X |
| Sternum with bone marrow | N/A | N/A | X |
| Cecum | N/A | N/A | X |
| Cranial (superior) cervical ganglion | N/A | N/A | X |
| Duodenum | N/A | N/A | X |
| Nerve, radial | N/A | N/A | X |
| Nerve, sciatic | N/A | N/A | X^b^ |
| Spinal nerve roots/cauda equina | N/A | N/A | X^b^ |
| Stomach | N/A | N/A | X |
| Nerve, sural | N/A | N/A | X^b^ |
| Thymus | N/A | N/A | X |
| Thyroid gland with parathyroid | N/A | N/A | X |
| Nerve, tibial | N/A | N/A | X^b^ |
| Nerve, ulnar | N/A | N/A | X^b^ |

Abbreviations: CNS, central nervous system; ddPCR, droplet digital polymerase chain reaction; DRG, dorsal root ganglion; GALT, gut-associated lymphoid tissue; GFP, green fluorescent protein; MSD, meso scale discovery; MSD ELISA, meso scale discovery enzyme-linked immunoassay; N/A, not applicable; X, collected.

^a^Right side collected for bilateral tissues and/or CNS structures specified for biodistribution or protein analysis while left side retained for histology/microscopy/molecular localization. Brain sampling followed the study specific procedure, with sections from the left side of brain trimmed to match the list of structures and areas collected from the right side for frozen samples for biodistribution and MSD-ELISA GFP analysis. **See Supplemental Figure 15 for representative images.**

^b^Collected in 10% neutral-buffered formalin for at least two but no more than three overnights and then process to paraffin block.

^c^n=4 from each region collected and frozen for biodistribution and protein analysis. For histopathology, at least n=4 each were collected from cervical, thoracic, and lumbar regions, and at least n=4 from the sacral regions (extraforaminal sacral DRG frequently collocated with transverse sections of caudal spinal nerve roots/cauda equina).

^d^Collected in modified Davidson’s fixative and stored in 10% neutral-buffered formalin per CRO standard operating procedure.

**Supplemental Table 3.** **Summary of Serum Titer Results for Each Dose Group.**

| **Dose Group 1 (Vehicle Control-Intrathecal-LP)** | | | | | | | | | | | | | | | | | | | | | | |
| --- | --- | --- | --- | --- | --- | --- | --- | --- | --- | --- | --- | --- | --- | --- | --- | --- | --- | --- | --- | --- | --- | --- |
| **Study Day^a^** | | | | **P0001** | | | | | | **P0002** | | | **P0003** | | | | | **P0004** | | | | |
| –15 | | | | (<20) Neg Titer | | | | | | 160 | | | 20480 | | | | | 320 | | | | |
| 15 | | | | 20 | | | | | | 160 | | | 163840 | | | | | 160 | | | | |
| 22 | | | | 20 | | | | | | 640 | | | 163840 | | | | | (<20) Neg Titer | | | | |
| 28 | | | | 20 | | | | | | 160 | | | 40960 | | | | | (<20) Neg Titer | | | | |
| **Dose Group 2 (Vehicle Control-ICM)** | | | | | | | | | | | | | | | | | | | | | | |
| **Study Day^a^** | | | **P0101** | | | | | **P0102** | | | | | | | | **P0103** | | | | | | **P0104** |
| –15 | | | 320 | | | | | 80 | | | | | | | | (<20) Neg Titer | | | | | | 2560 |
| 15 | | | 1280 | | | | | 20 | | | | | | | | (<20) Neg Titer | | | | | | 10240 |
| 22 | | | 640 | | | | | (<20) Neg Titer | | | | | | | | (<20) Neg Titer | | | | | | 5120 |
| 28 | | | 640 | | | | | (<20) Neg Titer | | | | | | | | (<20) Neg Titer | | | | | | 10240 |
| **Dose Group 3 (Intrathecal-LP, 1.0×10^13^ vg/ animal)** | | | | | | | | | | | | | | | | | | | | | | |
| **Study Day^a^** | | **P0201** | | | | | **P0202** | | | | | | | | **P0203** | | | | | | **P0204** | |
| –15 | | 160 | | | | | (<20) Neg Titer | | | | | | | | (<20) Neg Titer | | | | | | 20 | |
| 15 | | 163840 | | | | | 40960 | | | | | | | | 20480 | | | | | | 81920 | |
| 22 | | 81920 | | | | | 81920 | | | | | | | | 10240 | | | | | | 163840 | |
| 28 | | 40960 | | | | | 163840 | | | | | | | | 5120 | | | | | | 163840 | |
| **Dose Group 4 (Intrathecal-LP, 3.0×10^13^ vg/ animal)** | | | | | | | | | | | | | | | | | | | | | | |
| **Study Day^a^** | | | | | **P0301** | | | | **P0302** | | | **P0303** | | | | | **P0304** | | | | | |
| –15 | | | | | 320 | | | | 20 | | | 160 | | | | | (<20) Neg Titer | | | | | |
| 15 | | | | | 81920 | | | | 327680 | | | 327680 | | | | | 1310700 | | | | | |
| 22 | | | | | 81920 | | | | 327680 | | | 163840 | | | | | 327680 | | | | | |
| 28 | | | | | 81920 | | | | 163840 | | | 163840 | | | | | 327680 | | | | | |
| **Dose Group 5 (ICM, 1.0×10^13^ vg/ animal)** | | | | | | | | | | | | | | | | | | | | | | |
| **Study Day^a^** | | **P0401** | | | | | **P0402** | | | | **P0403** | | | | | | | | **P0404** | | | |
| –15 | | 640 | | | | | 20 | | | | (<20) Neg Titer | | | | | | | | (<20) Neg Titer | | | |
| 15 | | 163840 | | | | | 327680 | | | | 20480 | | | | | | | | 163840 | | | |
| 22 | | 163840 | | | | | 327680 | | | | 20480 | | | | | | | | 163840 | | | |
| 28 | | 163840 | | | | | 327680 | | | | 40960 | | | | | | | | 163840 | | | |
| **Dose Group 6 (ICM, 3.0×10^13^ vg/ animal)** | | | | | | | | | | | | | | | | | | | | | | |
| **Study Day^a^** | **P0501** | | | | | **P0502** | | | | | | | | **P0503** | | | | | | **P0504** | | |
| –15 | 160 | | | | | (<20) Neg Titer | | | | | | | | 40 | | | | | | (<20) Neg Titer | | |
| 15 | 327680 | | | | | 327680 | | | | | | | | 1310700 | | | | | | 40960 | | |
| 22 | 327680 | | | | | 163840 | | | | | | | | 655360 | | | | | | 163840 | | |
| 28 | 655360 | | | | | 81920 | | | | | | | | 655360 | | | | | | 40960 | | |

Abbreviations: (<20), titer <20; ICM, intracisterna magna; LP, lumbar puncture; Neg Titer, negative titer; vg, vector genome.

^a^Study Day –15 was provided because at the time of collection the individual animals had not been assigned to a specific dosing cohort.

### Supplemental Table 4. Summary of CSF Titer Results for Each Dose Group (Study Day Versus Animal).

| **Dose Group 1 (Vehicle Control-Intrathecal-LP)** | | | | |
| --- | --- | --- | --- | --- |
| **Study Day** | **P0001** | **P0002** | **P0003** | **P0004** |
| 1 | NI | NI | 160 | NI |
| 15 | NI | NI | 160 | NI |
| 28 | NI | NI | 160 | NI |
| **Dose Group 2 (Vehicle Control-ICM)** | | | | |
| **Study Day** | **P0101** | **P0102** | **P0103** | **P0104** |
| 1 | NI | NI | NI | 320 |
| 15 | NI | NI | NI | 40 |
| 28 | NI | NI | NI | 40 |
| **Dose Group 3 (Intrathecal-LP, 1.0×10^13^ vg/animal)** | | | | |
| **Study Day** | **P0201** | **P0202** | **P0203** | **P0204** |
| 1  15  28 | NI  1280  5120 | NI  80  10240 | NI  (<20) Neg Titer  NI | NI  160  80 |
| **Dose Group 4 (Intrathecal-LP, 3.0×10^13^ vg/ animal)** | | | | |
| **Study Day** | **P0301** | **P0302** | **P0303** | **P0304** |
| 1 | NI | NI | NI | NI |
| 15 | 80 | 1280 | 320 | 5120 |
| 28 | 640 | 20480 | 2560 | 5120 |
| **Dose Group 5 (ICM, 1.0×10^13^ vg/ animal)** | | | | |
| **Study Day** | **P0401** | **P0402** | **P0403** | **P0404** |
| 1 | NI | NI | NI | NI |
| 15 | 640 | 1280 | 160 | 320 |
| 28 | 20480 | 20480 | 2560 | 5120 |
| **Dose Group 6 (ICM, 3.0×10^13^ vg/ animal)** | | | | |
| **Study Day** | **P0501** | **P0502** | **P0503** | **P0504** |
| 1 | NI | NI | NI | NI |
| 15 | 5120 | 320 | 640 | 80 |
| 28 | 2560 | 2560 | 2560 | 320 |

Abbreviations: (<20), titer <20; CSF, cerebrospinal fluid; ICM, intracisterna magna; LP, lumbar puncture; NI, negative immunodepletion; Neg Titer, negative titer; vg, vector genome.

**Supplemental Table 5. Summary Statistics of scAAV9-CB-GFP DNA in Group 1 (Vehicle) Intrathecal-LP-Dosed Animals at 28 Days****.**

|  | **scAAV9-CB-GFP DNA concentration (copies/diploid genome)** | | | | | |  |
| --- | --- | --- | --- | --- | --- | --- | --- |
|  | **Median** | **Minimum** | **Maximum** | **Mean** | **SD** | **CV** | **N** |
| Prefrontal cortex | 0 | 0 | 0 | 0 | N/A | N/A | 4 |
| Temporal cortex-entorhinal cortex | 0 | 0 | 0 | 0 | N/A | N/A | 4 |
| Caudate nucleus | 0 | 0 | 0 | 0 | N/A | N/A | 4 |
| Putamen (basal ganglia) | 0 | 0 | 0 | 0 | N/A | N/A | 4 |
| Cingulate gyrus | 0 | 0 | 0 | 0 | N/A | N/A | 4 |
| Corpus callosum | 0 | 0 | 0 | 0 | N/A | N/A | 4 |
| Temporal cortex-auditory cortex | 0 | 0 | 0 | 0 | N/A | N/A | 4 |
| Parietal cortex | 0 | 0 | 0 | 0 | N/A | N/A | 4 |
| Thalamus | 0 | 0 | 0 | 0 | N/A | N/A | 4 |
| Hypothalamus | 0 | 0 | 0 | 0 | N/A | N/A | 4 |
| Hippocampus | 0 | 0 | 0 | 0 | N/A | N/A | 4 |
| Amygdala | 0 | 0 | 0 | 0 | N/A | N/A | 4 |
| Substantia nigra | 0 | 0 | 0 | 0 | N/A | N/A | 4 |
| Pons | 0 | 0 | 0 | 0 | N/A | N/A | 4 |
| Cerebellum | 0 | 0 | 0 | 0 | N/A | N/A | 4 |
| Occipital cortex: primary visual cortex | 0 | 0 | 0 | 0 | N/A | N/A | 4 |
| Deep cerebellar nuclei | 0 | 0 | 0 | 0 | N/A | N/A | 4 |
| Heart | 0 | 0 | 0 | 0 | N/A | N/A | 4 |
| Spinal cord (Cervical) | 0 | 0 | 0 | 0 | N/A | N/A | 4 |
| Spinal cord (thoracic) | 0 | 0 | 0 | 0 | N/A | N/A | 4 |
| Spinal cord (lumbar) | 0 | 0 | 0 | 0 | N/A | N/A | 4 |
| Spinal cord (sacral) | 0 | 0 | 0 | 0 | N/A | N/A | 4 |
| Trigeminal ganglia/nerve | 0 | 0 | 0 | 0 | N/A | N/A | 4 |
| Liver | 0 | 0 | 0 | 0 | N/A | N/A | 4 |
| Lung | 0 | 0 | 0 | 0 | N/A | N/A | 4 |
| Kidney  (right pole) | 0 | 0 | 0 | 0 | N/A | N/A | 4 |
| Ovary | 0 | 0 | 0 | 0 | N/A | N/A | 4 |
| Spleen | 0 | 0 | 0 | 0 | N/A | N/A | 4 |
| Mandibular | 0 | 0 | 0 | 0 | N/A | N/A | 4 |
| Muscle, biceps femoris | 0 | 0 | 0 | 0 | N/A | N/A | 4 |
| Muscle, diaphragm | 0 | 0 | 0 | 0 | N/A | N/A | 4 |
| Pancreas | 0 | 0 | 0 | 0 | N/A | N/A | 4 |
| Eye | 0 | 0 | 0 | 0 | N/A | N/A | 4 |
| Optic nerve | 0 | 0 | 0 | 0 | N/A | N/A | 4 |
| DRG cervical | 0 | 0 | 0 | 0 | N/A | N/A | 4 |
| DRG thoracic | 0 | 0 | 0 | 0 | N/A | N/A | 4 |
| DRG lumbar | 0 | 0 | 0 | 0 | N/A | N/A | 4 |
| DRG sacral | 0 | 0 | 0 | 0 | N/A | N/A | 4 |
| Blood pellet | 0 | 0 | 0 | 0 | N/A | N/A | 4 |

Abbreviations: CV, coefficient of variation; DRG, dorsal root ganglion; LP, lumbar puncture; N/A, not applicable; scAAV9-CB-GFP, self-complementary adeno-associated virus serotype 9–chicken β-actin promoter–green fluorescent protein; SD, standard deviation.

**Supplemental Table 6. Summary Statistics of scAAV9-CB-GFP DNA in Group 2 (Vehicle) ICM-Dosed Animals at 28 Days.**

|  | **scAAV9-CB-GFP DNA concentration (copies/diploid genome)** | | | | | |  |
| --- | --- | --- | --- | --- | --- | --- | --- |
|  | **Median** | **Minimum** | **Maximum** | **Mean** | **SD** | **CV** | **N** |
| Prefrontal cortex | 0 | 0 | 0 | 0 | N/A | N/A | 4 |
| Temporal cortex-entorhinal cortex | 0 | 0 | 0 | 0 | N/A | N/A | 4 |
| Caudate nucleus | 0 | 0 | 0 | 0 | N/A | N/A | 4 |
| Putamen (basal ganglia) | 0 | 0 | 0 | 0 | N/A | N/A | 4 |
| Cingulate gyrus | 0 | 0 | 0 | 0 | N/A | N/A | 4 |
| Corpus callosum | 0 | 0 | 0 | 0 | N/A | N/A | 4 |
| Temporal cortex-auditory cortex | 0 | 0 | 0 | 0 | N/A | N/A | 4 |
| Parietal cortex | 0 | 0 | 0 | 0 | N/A | N/A | 4 |
| Thalamus | 0 | 0 | 0 | 0 | N/A | N/A | 4 |
| Hypothalamus | 0 | 0 | 0 | 0 | N/A | N/A | 4 |
| Hippocampus | 0 | 0 | 0 | 0 | N/A | N/A | 4 |
| Amygdala | 0 | 0 | 0 | 0 | N/A | N/A | 4 |
| Substantia nigra | 0 | 0 | 0 | 0 | N/A | N/A | 4 |
| Pons | 0 | 0 | 0 | 0 | N/A | N/A | 4 |
| Cerebellum | 0 | 0 | 0 | 0 | N/A | N/A | 4 |
| Occipital cortex: primary visual cortex | 0 | 0 | 0 | 0 | N/A | N/A | 4 |
| Deep cerebellar nuclei | 0 | 0 | 0 | 0 | N/A | N/A | 4 |
| Heart | 0 | 0 | 0 | 0 | N/A | N/A | 4 |
| Spinal cord (cervical) | 0 | 0 | 0 | 0 | N/A | N/A | 4 |
| Spinal cord (thoracic) | 0 | 0 | 0 | 0 | N/A | N/A | 4 |
| Spinal cord (lumbar) | 0 | 0 | 0 | 0 | N/A | N/A | 4 |
| Spinal cord (sacral) | 0 | 0 | 0 | 0 | N/A | N/A | 4 |
| Trigeminal ganglia/nerve | 0 | 0 | 0 | 0 | N/A | N/A | 4 |
| Liver | 0 | 0 | 0 | 0 | N/A | N/A | 4 |
| Lung | 0 | 0 | 0 | 0 | N/A | N/A | 4 |
| Kidney  (right pole) | 0 | 0 | 0 | 0 | N/A | N/A | 4 |
| Ovary | 0 | 0 | 0 | 0 | N/A | N/A | 4 |
| Spleen | 0 | 0 | 0 | 0 | N/A | N/A | 4 |
| Mandibular | 0 | 0 | 0 | 0 | N/A | N/A | 4 |
| Muscle, biceps femoris | 0 | 0 | 0 | 0 | N/A | N/A | 4 |
| Muscle, diaphragm | 0 | 0 | 0 | 0 | N/A | N/A | 4 |
| Pancreas | 0 | 0 | 0 | 0 | N/A | N/A | 4 |
| Eye | 0 | 0 | 0 | 0 | N/A | N/A | 4 |
| Optic nerve | 0 | 0 | 0 | 0 | N/A | N/A | 4 |
| DRG cervical | 0 | 0 | 0 | 0 | N/A | N/A | 4 |
| DRG thoracic | 0 | 0 | 0 | 0 | N/A | N/A | 4 |
| DRG lumbar | 0 | 0 | 0 | 0 | N/A | N/A | 4 |
| DRG sacral | 0 | 0 | 0 | 0 | N/A | N/A | 4 |
| Blood pellet | 0 | 0 | 0 | 0 | N/A | N/A | 4 |

Abbreviations: CV, coefficient of variation; DRG, dorsal root ganglion; ICM, intracisterna magna; N/A, not applicable; scAAV9-CB-GFP, self-complementary adeno-associated virus serotype 9–chicken β-actin promoter–green fluorescent protein; SD, standard deviation.

**Supplemental Table 7. Summary Statistics of scAAV9-CB-GFP DNA in Group 3 (1.0×10^13^ vg/animal) Intrathecal-LP-Dosed Animals at 28 Days.**

|  | **scAAV9-CB-GFP DNA concentration (copies/diploid genome)** | | | | | |  |
| --- | --- | --- | --- | --- | --- | --- | --- |
|  | **Median** | **Minimum** | **Maximum** | **Mean** | **SD** | **CV** | **N** |
| Prefrontal cortex | 0.21 | 0.02 | 0.95 | 0.34 | 0.43 | 125 | 4 |
| Temporal cortex-entorhinal cortex | 0.11 | 0.00 | 1.09 | 0.33 | 0.52 | 157 | 4 |
| Caudate nucleus | 0.00 | 0.00 | 0.04 | 0.01 | 0.02 | 200 | 4 |
| Putamen (basal ganglia) | 0.02 | 0.00 | 0.06 | 0.02 | 0.03 | 121 | 4 |
| Cingulate gyrus | 0.05 | 0.00 | 0.12 | 0.06 | 0.05 | 89 | 4 |
| Corpus callosum | 0.00 | 0.00 | 0.00 | 0.00 | 0.00 | N/A | 4 |
| Temporal cortex-auditory cortex | 0.03 | 0.00 | 0.59 | 0.16 | 0.29 | 174 | 4 |
| Parietal cortex | 0.05 | 0.00 | 0.60 | 0.18 | 0.29 | 163 | 4 |
| Thalamus | 0.02 | 0.00 | 0.06 | 0.03 | 0.03 | 120 | 4 |
| Hypothalamus | 0.03 | 0.00 | 0.99 | 0.26 | 0.48 | 183 | 4 |
| Hippocampus | 0.04 | 0.00 | 0.22 | 0.07 | 0.10 | 138 | 4 |
| Amygdala | 0.04 | 0.03 | 2.08 | 0.55 | 1.02 | 185 | 4 |
| Substantia nigra | 0.00 | 0.00 | 0.09 | 0.02 | 0.04 | 200 | 4 |
| Pons | 0.04 | 0.03 | 0.35 | 0.11 | 0.15 | 138 | 4 |
| Cerebellum | 0.00 | 0.00 | 0.08 | 0.02 | 0.04 | 189 | 4 |
| Occipital cortex: primary visual cortex | 0.15 | 0.00 | 0.57 | 0.22 | 0.26 | 122 | 4 |
| Deep cerebellar nuclei | 0.07 | 0.00 | 0.15 | 0.07 | 0.08 | 116 | 4 |
| Heart | 0.28 | 0.20 | 0.39 | 0.29 | 0.09 | 32.0 | 4 |
| Spinal cord (cervical) | 0.75 | 0.03 | 1.46 | 0.75 | 0.82 | 110 | 4 |
| Spinal cord (thoracic) | 1.67 | 0.10 | 3.14 | 1.65 | 1.26 | 76.6 | 4 |
| Spinal cord (lumbar) | 1.75 | 0.22 | 1.83 | 1.39 | 0.78 | 56.3 | 4 |
| Spinal cord (sacral) | 1.72 | 0.79 | 2.46 | 1.67 | 0.68 | 41.0 | 4 |
| Trigeminal ganglia/nerve | 0.07 | 0.02 | 1.85 | 0.51 | 0.90 | 177 | 4 |
| Liver | 12.46 | 8.48 | 36.52 | 17.48 | 13.04 | 74.6 | 4 |
| Lung | 0.16 | 0.11 | 0.30 | 0.18 | 0.09 | 48.6 | 4 |
| Kidney  (right pole) | 0.13 | 0.08 | 0.28 | 0.15 | 0.09 | 56.7 | 4 |
| Ovary | 0.02 | 0.01 | 0.05 | 0.03 | 0.02 | 60.1 | 4 |
| Spleen | 0.39 | 0.12 | 1.92 | 0.70 | 0.82 | 116 | 4 |
| Mandibular | 0.62 | 0.27 | 1.30 | 0.70 | 0.44 | 62.5 | 4 |
| Muscle, biceps femoris | 0.05 | 0.04 | 0.08 | 0.06 | 0.02 | 36.1 | 4 |
| Muscle, diaphragm | 0.12 | 0.04 | 0.17 | 0.11 | 0.06 | 50.3 | 4 |
| Pancreas | 0.04 | 0.00 | 0.14 | 0.06 | 0.06 | 109 | 4 |
| Eye | 0.07 | 0.02 | 0.08 | 0.06 | 0.03 | 48.6 | 4 |
| Optic nerve | 0.27 | 0.00 | 3.17 | 0.93 | 1.52 | 164 | 4 |
| DRG cervical | 2.07 | 0.63 | 4.79 | 2.39 | 1.85 | 77.4 | 4 |
| DRG thoracic | 1.48 | 0.33 | 1.56 | 1.21 | 0.59 | 48.6 | 4 |
| DRG lumbar | 1.55 | 0.64 | 5.73 | 2.37 | 2.30 | 97.4 | 4 |
| DRG sacral | 4.32 | 1.40 | 10.06 | 5.03 | 3.70 | 73.5 | 4 |
| Blood pellet | 1.56 | 0.16 | 3.28 | 1.64 | 1.28 | 78.1 | 4 |

Abbreviations: CV, coefficient of variation; DRG, dorsal root ganglion; LP, lumbar puncture; N/A, not applicable; scAAV9-CB-GFP, self-complementary adeno-associated virus serotype 9–chicken β-actin promoter–green fluorescent protein; SD, standard deviation; vg, vector genome.

**Supplemental Table 8. Summary Statistics of scAAV9-CB-GFP DNA in Group 4 (3.0×10^13^ vg/animal) Intrathecal-LP-Dosed Animals at 28 Days.**

|  | **scAAV9-CB-GFP DNA concentration (copies/diploid genome)** | | | | | |  |
| --- | --- | --- | --- | --- | --- | --- | --- |
|  | **Median** | **Minimum** | **Maximum** | **Mean** | **SD** | **CV** | **N** |
| Prefrontal cortex | 0.71 | 0.09 | 1.58 | 0.77 | 0.64 | 83.4 | 4 |
| Temporal cortex-entorhinal cortex | 0.10 | 0.00 | 1.09 | 0.32 | 0.51 | 160 | 4 |
| Caudate nucleus | 0.05 | 0.00 | 0.09 | 0.05 | 0.04 | 79.7 | 4 |
| Putamen (basal ganglia) | 0.06 | 0.00 | 0.09 | 0.06 | 0.04 | 74.2 | 4 |
| Cingulate gyrus | 0.10 | 0.05 | 0.55 | 0.20 | 0.24 | 118 | 4 |
| Corpus callosum | 0.04 | 0.03 | 0.18 | 0.07 | 0.07 | 98.3 | 4 |
| Temporal cortex-auditory cortex | 0.60 | 0.08 | 2.26 | 0.89 | 0.98 | 111 | 4 |
| Parietal cortex | 0.41 | 0.03 | 2.94 | 0.95 | 1.34 | 142 | 4 |
| Thalamus | 0.07 | 0.06 | 0.09 | 0.07 | 0.01 | 18.5 | 4 |
| Hypothalamus | 0.07 | 0.00 | 0.45 | 0.15 | 0.20 | 138 | 4 |
| Hippocampus | 0.46 | 0.09 | 2.32 | 0.83 | 1.05 | 126 | 4 |
| Amygdala | 0.11 | 0.04 | 0.40 | 0.16 | 0.16 | 100 | 4 |
| Substantia nigra | 0.07 | 0.00 | 0.11 | 0.06 | 0.05 | 72.4 | 4 |
| Pons | 0.28 | 0.09 | 0.75 | 0.35 | 0.31 | 90.6 | 4 |
| Cerebellum | 0.08 | 0.01 | 2.61 | 0.70 | 1.28 | 183 | 4 |
| Occipital cortex: primary visual cortex | 0.45 | 0.03 | 0.72 | 0.41 | 0.28 | 68.8 | 4 |
| Deep cerebellar nuclei | 0.16 | 0.03 | 1.83 | 0.55 | 0.86 | 158 | 4 |
| Heart | 1.17 | 0.82 | 2.06 | 1.31 | 0.54 | 40.9 | 4 |
| Spinal cord (cervical) | 1.65 | 0.81 | 2.99 | 1.77 | 0.90 | 50.8 | 4 |
| Spinal cord (thoracic) | 3.94 | 1.37 | 4.04 | 3.32 | 1.30 | 39.3 | 4 |
| Spinal cord (lumbar) | 3.69 | 1.12 | 4.89 | 3.35 | 1.62 | 48.4 | 4 |
| Spinal cord (sacral) | 3.88 | 1.20 | 5.64 | 3.65 | 1.83 | 50.2 | 4 |
| Trigeminal ganglia/nerve | 5.82 | 0.71 | 8.74 | 5.27 | 3.35 | 63.5 | 4 |
| Liver | 72.12 | 4.86 | 87.12 | 59.06 | 37.86 | 64.1 | 4 |
| Lung | 0.71 | 0.42 | 1.10 | 0.74 | 0.29 | 40.0 | 4 |
| Kidney  (right pole) | 0.42 | 0.39 | 0.84 | 0.52 | 0.21 | 41.4 | 4 |
| Ovary | 0.10 | 0.07 | 0.13 | 0.10 | 0.03 | 30.7 | 4 |
| Spleen | 0.62 | 0.50 | 0.77 | 0.63 | 0.13 | 21.1 | 4 |
| Mandibular | 3.14 | 0.96 | 4.52 | 2.94 | 1.49 | 50.7 | 4 |
| Muscle, biceps femoris | 0.29 | 0.22 | 0.40 | 0.30 | 0.08 | 26.3 | 4 |
| Muscle, diaphragm | 0.53 | 0.32 | 10.01 | 2.85 | 4.77 | 168 | 4 |
| Pancreas | 0.17 | 0.11 | 0.36 | 0.20 | 0.11 | 55.8 | 4 |
| Eye | 0.22 | 0.07 | 0.38 | 0.22 | 0.13 | 55.9 | 4 |
| Optic nerve | 0.64 | 0.04 | 5.00 | 1.58 | 2.31 | 146 | 4 |
| DRG cervical | 3.95 | 2.40 | 11.94 | 5.56 | 4.32 | 77.7 | 4 |
| DRG thoracic | 2.44 | 1.69 | 5.94 | 3.13 | 1.94 | 61.9 | 4 |
| DRG lumbar | 3.19 | 0.59 | 3.94 | 2.73 | 1.52 | 55.6 | 4 |
| DRG sacral | 5.93 | 2.47 | 11.30 | 6.41 | 3.80 | 59.3 | 4 |
| Blood pellet | 6.91 | 4.47 | 20.53 | 9.71 | 7.35 | 75.7 | 4 |

Abbreviations: CV, coefficient of variation; DRG, dorsal root ganglion; LP, lumbar puncture; scAAV9-CB-GFP, self-complementary adeno-associated virus serotype 9–chicken β-actin promoter–green fluorescent protein; SD, standard deviation; vg, vector genome.

**Supplemental Table 9. Summary Statistics of scAAV9-CB-GFP DNA in Group 5 (1.0×10^13^ vg/animal) ICM-Dosed Animals at 28 Days.**

|  | **scAAV9-CB-GFP DNA concentration (copies/diploid genome)** | | | | | |  |
| --- | --- | --- | --- | --- | --- | --- | --- |
|  | **Median** | **Minimum** | **Maximum** | **Mean** | **SD** | **CV** | **N** |
| Prefrontal cortex | 0.05 | 0.04 | 0.09 | 0.06 | 0.02 | 40.3 | 4 |
| Temporal cortex-entorhinal cortex | 0.66 | 0.00 | 1.86 | 0.79 | 0.88 | 111 | 4 |
| Caudate nucleus | 0.00 | 0.00 | 0.03 | 0.01 | 0.02 | 200 | 4 |
| Putamen (basal ganglia) | 0.01 | 0.00 | 0.03 | 0.01 | 0.02 | 116 | 4 |
| Cingulate gyrus | 0.00 | 0.00 | 0.53 | 0.13 | 0.27 | 200 | 4 |
| Corpus callosum | 0.00 | 0.00 | 0.00 | 0.00 | 0.00 | N/A | 4 |
| Temporal cortex-auditory cortex | 0.16 | 0.04 | 0.27 | 0.15 | 0.09 | 60.7 | 4 |
| Parietal cortex | 0.06 | 0.00 | 1.32 | 0.36 | 0.64 | 180 | 4 |
| Thalamus | 0.00 | 0.00 | 0.00 | 0.00 | 0.00 | N/A | 4 |
| Hypothalamus | 0.00 | 0.00 | 0.04 | 0.01 | 0.02 | 200 | 4 |
| Hippocampus | 0.03 | 0.00 | 0.34 | 0.10 | 0.16 | 159 | 4 |
| Amygdala | 0.02 | 0.00 | 0.69 | 0.18 | 0.34 | 183 | 4 |
| Substantia nigra | 0.28 | 0.00 | 0.83 | 0.35 | 0.37 | 107 | 4 |
| Pons | 0.03 | 0.00 | 0.07 | 0.03 | 0.04 | 116 | 4 |
| Cerebellum | 0.01 | 0.00 | 0.05 | 0.02 | 0.02 | 121 | 4 |
| Occipital cortex: primary visual cortex | 0.03 | 0.00 | 0.11 | 0.04 | 0.05 | 110 | 4 |
| Deep cerebellar nuclei | 0.01 | 0.00 | 0.14 | 0.04 | 0.07 | 175 | 4 |
| Heart | 0.12 | 0.07 | 0.14 | 0.11 | 0.03 | 26.3 | 4 |
| Spinal cord (cervical) | 0.30 | 0.13 | 1.26 | 0.50 | 0.52 | 103 | 4 |
| Spinal cord (thoracic) | 1.32 | 0.54 | 3.95 | 1.78 | 1.59 | 89.2 | 4 |
| Spinal cord (lumbar) | 1.47 | 0.10 | 3.33 | 1.59 | 1.36 | 85.6 | 4 |
| Spinal cord (sacral) | 0.28 | 0.08 | 25.89 | 6.64 | 12.84 | 193 | 4 |
| Trigeminal ganglia/nerve | 3.28 | 0.90 | 7.00 | 3.61 | 2.75 | 76.1 | 4 |
| Liver | 10.57 | 3.54 | 15.77 | 10.11 | 6.06 | 60.0 | 4 |
| Lung | 0.15 | 0.06 | 0.21 | 0.14 | 0.07 | 52.7 | 4 |
| Kidney  (right pole) | 0.09 | 0.06 | 0.14 | 0.10 | 0.04 | 37.8 | 4 |
| Ovary | 0.02 | 0.01 | 0.02 | 0.02 | 0.01 | 42.9 | 4 |
| Spleen | 0.22 | 0.20 | 0.36 | 0.25 | 0.07 | 29.8 | 4 |
| Mandibular | 5.67 | 0.70 | 13.91 | 6.49 | 6.24 | 96.1 | 4 |
| Muscle, biceps femoris | 0.04 | 0.00 | 0.09 | 0.04 | 0.04 | 88.6 | 4 |
| Muscle, diaphragm | 0.05 | 0.05 | 1.75 | 0.47 | 0.85 | 179 | 4 |
| Pancreas | 0.03 | 0.02 | 0.06 | 0.03 | 0.02 | 62.9 | 4 |
| Eye | 0.05 | 0.02 | 0.19 | 0.08 | 0.08 | 104 | 4 |
| Optic nerve | 0.53 | 0.02 | 1.46 | 0.64 | 0.60 | 94.6 | 4 |
| DRG cervical | 2.83 | 1.87 | 8.22 | 3.94 | 2.98 | 75.6 | 4 |
| DRG thoracic | 3.04 | 2.09 | 10.05 | 4.56 | 3.71 | 81.4 | 4 |
| DRG lumbar | 7.38 | 3.78 | 10.71 | 7.31 | 3.05 | 41.7 | 4 |
| DRG sacral | 0.41 | 0.06 | 20.24 | 5.28 | 9.98 | 189 | 4 |
| Blood pellet | 1.63 | 1.24 | 1.82 | 1.58 | 0.25 | 15.6 | 4 |

Abbreviations: CV, coefficient of variation; DRG, dorsal root ganglion; ICM, intracisterna magna; N/A, not applicable; scAAV9-CB-GFP, self-complementary adeno-associated virus serotype 9–chicken β-actin promoter–green fluorescent protein; SD, standard deviation; vg, vector genomes.

**Supplemental Table 10. Summary Statistics of scAAV9-CB-GFP DNA in Group 6 (3.0×10^13^ vg/animal) ICM-Dosed Animals at 28 Days.**

|  | **scAAV9-CB-GFP DNA concentration (copies/diploid genome)** | | | | | |  |
| --- | --- | --- | --- | --- | --- | --- | --- |
|  | **Median** | **Minimum** | **Maximum** | **Mean** | **SD** | **CV** | **N** |
| Prefrontal cortex | 0.18 | 0.05 | 0.42 | 0.21 | 0.15 | 74.2 | 4 |
| Temporal cortex-entorhinal cortex | 0.15 | 0.13 | 0.16 | 0.14 | 0.01 | 9.47 | 4 |
| Caudate nucleus | 0.08 | 0.00 | 0.18 | 0.08 | 0.08 | 90.9 | 4 |
| Putamen (basal ganglia) | 0.10 | 0.04 | 0.21 | 0.12 | 0.07 | 62.0 | 4 |
| Cingulate gyrus | 0.14 | 0.08 | 1.12 | 0.37 | 0.50 | 135 | 4 |
| Corpus callosum | 0.37 | 0.04 | 1.12 | 0.47 | 0.53 | 112 | 4 |
| Temporal cortex-auditory cortex | 0.64 | 0.08 | 1.95 | 0.82 | 0.88 | 107 | 4 |
| Parietal cortex | 0.30 | 0.03 | 0.74 | 0.34 | 0.31 | 91.4 | 4 |
| Thalamus | 0.09 | 0.03 | 0.17 | 0.09 | 0.06 | 62.0 | 4 |
| Hypothalamus | 0.17 | 0.07 | 0.27 | 0.17 | 0.11 | 65.7 | 4 |
| Hippocampus | 0.26 | 0.10 | 0.74 | 0.34 | 0.29 | 86.2 | 4 |
| Amygdala | 0.19 | 0.08 | 2.35 | 0.70 | 1.10 | 156 | 4 |
| Substantia nigra | 0.12 | 0.04 | 0.17 | 0.11 | 0.06 | 48.4 | 4 |
| Pons | 0.15 | 0.07 | 0.29 | 0.16 | 0.09 | 57.5 | 4 |
| Cerebellum | 0.02 | 0.00 | 0.03 | 0.02 | 0.01 | 53.9 | 4 |
| Occipital cortex: primary visual cortex | 0.13 | 0.07 | 0.58 | 0.23 | 0.24 | 105 | 4 |
| Deep cerebellar nuclei | 0.07 | 0.01 | 0.09 | 0.06 | 0.03 | 60.0 | 4 |
| Heart | 0.64 | 0.47 | 0.84 | 0.65 | 0.15 | 23.7 | 4 |
| Spinal cord (cervical) | 1.13 | 0.34 | 1.85 | 1.12 | 0.67 | 60.2 | 4 |
| Spinal cord (thoracic) | 3.39 | 0.50 | 6.62 | 3.47 | 2.58 | 74.1 | 4 |
| Spinal cord (lumbar) | 3.80 | 2.74 | 6.61 | 4.24 | 1.66 | 39.1 | 4 |
| Spinal cord (sacral) | 9.18 | 1.21 | 14.07 | 8.41 | 5.62 | 66.8 | 4 |
| Trigeminal ganglia/nerve | 1.57 | 0.39 | 2.20 | 1.43 | 0.86 | 60.0 | 4 |
| Liver | 81.32 | 33.62 | 110.88 | 76.78 | 36.80 | 47.9 | 4 |
| Lung | 0.45 | 0.21 | 0.60 | 0.43 | 0.19 | 45.1 | 4 |
| Kidney  (right pole) | 0.31 | 0.26 | 0.38 | 0.32 | 0.06 | 17.4 | 4 |
| Ovary | 0.06 | 0.04 | 0.10 | 0.07 | 0.03 | 43.1 | 4 |
| Spleen | 0.37 | 0.31 | 2.34 | 0.85 | 1.00 | 118 | 4 |
| Mandibular | 1.97 | 1.14 | 6.90 | 2.99 | 2.68 | 89.5 | 4 |
| Muscle, biceps femoris | 0.20 | 0.12 | 0.21 | 0.18 | 0.04 | 24.3 | 4 |
| Muscle, diaphragm | 0.35 | 0.11 | 0.54 | 0.34 | 0.19 | 57.0 | 4 |
| Pancreas | 0.19 | 0.03 | 0.48 | 0.22 | 0.20 | 89.2 | 4 |
| Eye | 0.19 | 0.11 | 0.27 | 0.19 | 0.07 | 34.4 | 4 |
| Optic nerve | 0.72 | 0.05 | 2.22 | 0.93 | 0.93 | 100 | 4 |
| DRG cervical | 1.77 | 1.51 | 3.08 | 2.03 | 0.71 | 35.0 | 4 |
| DRG thoracic | 3.72 | 2.40 | 9.69 | 4.88 | 3.37 | 69.0 | 4 |
| DRG lumbar | 6.73 | 3.90 | 12.45 | 7.45 | 3.61 | 48.5 | 4 |
| DRG sacral | 2.79 | 2.24 | 21.12 | 7.23 | 9.26 | 128 | 4 |
| Blood pellet | 2.93 | 2.36 | 19.32 | 6.88 | 8.30 | 121 | 4 |

Abbreviations: CV, coefficient of variation; DRG, dorsal root ganglion; ICM, intracisterna magna; scAAV9-CB-GFP, self-complementary adeno-associated virus serotype 9–chicken β-actin promoter–green fluorescent protein; SD, standard deviation; vg, vector genomes.

###### Supplemental Table 11. Rank Order of Tissue Concentrations of scAAV9-CB-GFP DNA at 28 Days Based on Median Tissue Concentrations.

|  | **Intrathecal-LP (1.0×10^13^)** | **Intrathecal-LP (3.0×10^13^)** | **ICM**  **(1.0×10^13^)** | **ICM**  **(3.0×10^13^)** |
| --- | --- | --- | --- | --- |
| Prefrontal cortex | 15 | 15 | 24 | 27 |
| Temporal cortex-entorhinal cortex | 20 | 32 | 10 | 30 |
| Caudate nucleus | 38 | 38 | 37 | 36 |
| Putamen (basal ganglia) | 35 | 37 | 32 | 34 |
| Cingulate gyrus | 24 | 31 | 36 | 31 |
| Corpus callosum | N/A | 39 | N/A | 18 |
| Temporal cortex-auditory cortex | 32 | 18 | 17 | 15 |
| Parietal cortex | 25 | 23 | 21 | 21 |
| Thalamus | 34 | 34 | N/A | 35 |
| Hypothalamus | 31 | 36 | 35 | 28 |
| Hippocampus | 30 | 20 | 27 | 22 |
| Amygdala | 27 | 29 | 30 | 24 |
| Substantia nigra | 37 | 35 | 14 | 33 |
| Pons | 28 | 25 | 28 | 29 |
| Cerebellum | 36 | 33 | 34 | 39 |
| Occipital cortex: primary visual cortex | 17 | 21 | 26 | 32 |
| Deep cerebellar nuclei | 23 | 28 | 33 | 37 |
| Heart | 13 | 13 | 19 | 14 |
| Spinal cord (cervical) | 10 | 12 | 13 | 12 |
| Spinal cord (thoracic) | 6 | 6 | 9 | 6 |
| Spinal cord (lumbar) | 4 | 8 | 8 | 4 |
| Spinal cord (sacral) | 5 | 7 | 15 | 2 |
| Trigeminal ganglia/nerve | 21 | 4 | 4 | 11 |
| Liver | 1 | 1 | 1 | 1 |
| Lung | 16 | 14 | 18 | 16 |
| Kidney (right pole) | 18 | 22 | 20 | 20 |
| Ovary | 33 | 30 | 31 | 38 |
| Spleen | 12 | 17 | 16 | 17 |
| Mandibular | 11 | 10 | 3 | 9 |
| Muscle, biceps femoris | 26 | 24 | 25 | 23 |
| Muscle, diaphragm | 19 | 19 | 23 | 19 |
| Pancreas | 29 | 27 | 29 | 26 |
| Eye | 22 | 26 | 22 | 25 |
| Optic nerve | 14 | 16 | 11 | 13 |
| DRG cervical | 3 | 5 | 6 | 10 |
| DRG thoracic | 9 | 11 | 5 | 5 |
| DRG lumbar | 8 | 9 | 2 | 3 |
| DRG sacral | 2 | 3 | 12 | 8 |
| Blood pellet | 7 | 2 | 7 | 7 |

Abbreviations: DRG, dorsal root ganglion; ICM, intracisterna magna; LP, lumbar puncture; N/A, not applicable; scAAV9-CB-GFP, self-complementary adeno-associated virus serotype 9–chicken β-actin promoter–green fluorescent protein.

###### Supplemental Table 12. Relative Median Concentrations of scAAV9-CB-GFP DNA in Animals at 28 Days Post-Dose.

|  | **DNA concentration**  **(copies/diploid genome)** | |  | **DNA concentration**  **(copies/diploid genome)** | |  |
| --- | --- | --- | --- | --- | --- | --- |
|  | **Intrathecal-LP (1.0×10^13^ vg/animal)** | **Intrathecal-LP (3.0×10^13^ vg/animal)** | **Ratio** | **ICM**  **(1.0×10^13^ vg/animal)** | **ICM**  **(3.0×10^13^ vg/animal)** | **Ratio** |
| Prefrontal cortex | 0.21 | 0.71 | 3.42 | 0.05 | 0.18 | 3.58 |
| Temporal cortex-entorhinal cortex | 0.11 | 0.10 | 0.89 | 0.66 | 0.15 | 0.18 |
| Caudate nucleus | 0.00 | 0.05 | NC | 0.00 | 0.08 | 10.76 |
| Putamen (basal ganglia) | 0.02 | 0.06 | 3.52 | 0.01 | 0.10 | 8.45 |
| Cingulate gyrus | 0.05 | 0.10 | 1.96 | 0.00 | 0.14 | 2.79 |
| Corpus callosum | 0.00 | 0.04 |  | 0.00 | 0.37 |  |
| Temporal cortex-auditory cortex | 0.03 | 0.60 | 18.29 | 0.16 | 0.64 | 5.33 |
| Parietal cortex | 0.05 | 0.41 | 8.07 | 0.06 | 0.30 | 0.96 |
| Thalamus | 0.02 | 0.07 | 3.68 | 0.00 | 0.09 |  |
| Hypothalamus | 0.03 | 0.07 | 2.14 | 0.00 | 0.17 | 18.49 |
| Hippocampus | 0.04 | 0.46 | 13.13 | 0.03 | 0.26 | 3.41 |
| Amygdala | 0.04 | 0.11 | 2.38 | 0.02 | 0.19 | 3.82 |
| Substantia nigra | 0.00 | 0.07 | NC | 0.28 | 0.12 | 0.33 |
| Pons | 0.04 | 0.28 | 7.35 | 0.03 | 0.15 | 4.90 |
| Cerebellum | 0.00 | 0.08 | 45.43 | 0.01 | 0.02 | 1.12 |
| Occipital cortex: primary visual cortex | 0.15 | 0.45 | 3.03 | 0.03 | 0.13 | 5.25 |
| Deep cerebellar nuclei | 0.07 | 0.16 | 2.33 | 0.01 | 0.07 | 1.47 |
| Heart | 0.28 | 1.17 | 4.14 | 0.12 | 0.64 | 5.89 |
| Spinal cord (cervical) | 0.75 | 1.65 | 2.18 | 0.30 | 1.13 | 2.23 |
| Spinal cord (thoracic) | 1.67 | 3.94 | 2.35 | 1.32 | 3.39 | 1.95 |
| Spinal cord (lumbar) | 1.75 | 3.69 | 2.11 | 1.47 | 3.80 | 2.66 |
| Spinal cord (sacral) | 1.72 | 3.88 | 2.26 | 0.28 | 9.18 | 1.27 |
| Trigeminal ganglia/nerve | 0.07 | 5.82 | 78.39 | 3.28 | 1.57 | 0.40 |
| Liver | 12.46 | 72.12 | 5.79 | 10.57 | 81.32 | 7.59 |
| Lung | 0.16 | 0.71 | 4.53 | 0.15 | 0.45 | 3.10 |
| Kidney (right pole) | 0.13 | 0.42 | 3.25 | 0.09 | 0.31 | 3.26 |
| Ovary | 0.02 | 0.10 | 4.33 | 0.02 | 0.06 | 3.93 |
| Spleen | 0.39 | 0.62 | 1.60 | 0.22 | 0.37 | 3.39 |
| Mandibular | 0.62 | 3.14 | 5.08 | 5.67 | 1.97 | 0.46 |
| Muscle, biceps femoris | 0.05 | 0.29 | 5.74 | 0.04 | 0.20 | 4.16 |
| Muscle, diaphragm | 0.12 | 0.53 | 4.39 | 0.05 | 0.35 | 0.72 |
| Pancreas | 0.04 | 0.17 | 4.60 | 0.03 | 0.19 | 6.44 |
| Eye | 0.07 | 0.22 | 3.15 | 0.05 | 0.19 | 2.40 |
| Optic nerve | 0.27 | 0.64 | 2.40 | 0.53 | 0.72 | 1.45 |
| DRG cervical | 2.07 | 3.95 | 1.91 | 2.83 | 1.77 | 0.52 |
| DRG thoracic | 1.48 | 2.44 | 1.65 | 3.04 | 3.72 | 1.07 |
| DRG lumbar | 1.55 | 3.19 | 2.06 | 7.38 | 6.73 | 1.02 |
| DRG sacral | 4.32 | 5.93 | 1.37 | 0.41 | 2.79 | 1.37 |
| Blood pellet | 1.56 | 6.91 | 4.42 | 1.63 | 2.93 | 4.36 |
|  |  | **Median** | 3.33 |  | **Median** | 2.79 |
|  |  | **Min** | 0.89 |  | **Min** | 0.18 |
|  |  | **Max** | 78.39 |  | **Max** | 18.49 |

Abbreviations: DRG, dorsal root ganglion; ICM, intracisterna magna; LP, lumbar puncture; Max, maximum; Min, minimum; NC, not calculable; scAAV9-CB-GFP, self-complementary adeno-associated virus serotype 9–chicken β-actin promoter–green fluorescent protein; vg, vector genomes.

**Supplemental Table 13. Individual Animal Data for scAAV9-CB-GFP Vector Genome ddPCR Quantitation in Intrathecal-LP Control Animals (Group 1).**

|  | **Average CNV of animal** | | | |
| --- | --- | --- | --- | --- |
| **Group 1 (Intrathecal-LP, vehicle)** | **P0001** | **P0002** | **P0003** | **P0004** |
| r. Prefrontal cortex | 0.0000 | 0.0000 | 0.0000 | 0.0000 |
| r. Temporal cortex-entorhinal cortex | 0.0011 | 0.0000 | 0.0000 | 0.0007 |
| r. Caudate nucleus | 0.0000 | 0.0000 | 0.0000 | 0.0000 |
| r. Putamen (basal ganglia) | 0.0000 | 0.0000 | 0.0000 | 0.0000 |
| Cingulate gyrus | 0.0000 | 0.0003 | 0.0000 | 0.0000 |
| Corpus callosum | 0.0000 | 0.0000 | 0.0000 | 0.0004 |
| r. Temporal cortex-auditory cortex | 0.0000 | 0.0003 | 0.0004 | 0.0009 |
| r. Parietal cortex | 0.0000 | 0.0000 | 0.0000 | 0.0000 |
| r. Thalamus | 0.0000 | 0.0000 | 0.0000 | 0.0000 |
| r. Hypothalamus | 0.0000 | 0.0000 | 0.0004 | 0.0000 |
| r. Hippocampus | 0.0000 | 0.0000 | 0.0000 | 0.0000 |
| r. Amygdala | 0.0000 | 0.0000 | 0.0006 | 0.0004 |
| r. Substantia nigra | 0.0000 | 0.0000 | 0.0003 | 0.0000 |
| r. Pons | 0.0000 | 0.0000 | 0.0000 | 0.0000 |
| r. Cerebellum | 0.0000 | 0.0000 | 0.0000 | 0.0000 |
| r. Occipital cortex: primary visual cortex | 0.0005 | 0.0000 | 0.0000 | 0.0000 |
| r. Deep cerebellar nuclei | 0.0000 | 0.0000 | 0.0000 | 0.0000 |
| Heart | 0.0006 | 0.0000 | 0.0000 | 0.0000 |
| Spinal cord (cervical) | 0.0000 | 0.0000 | 0.0000 | 0.0000 |
| Spinal cord (thoracic) | 0.0000 | 0.0000 | 0.0000 | 0.0000 |
| Spinal cord (lumbar) | 0.0000 | 0.0000 | 0.0000 | 0.0000 |
| Spinal cord (sacral) | 0.0000 | 0.0000 | 0.0000 | 0.0000 |
| r. Trigeminal ganglia/nerve | 0.0000 | 0.0000 | 0.0000 | 0.0003 |
| Liver | 0.0000 | 0.0000 | 0.0000 | 0.0000 |
| Lung | 0.0000 | 0.0000 | 0.0000 | 0.0000 |
| Kidney (right pole) | 0.0000 | 0.0000 | 0.0000 | 0.0000 |
| r. Ovary | 0.0000 | 0.0000 | 0.0000 | 0.0000 |
| Spleen | 0.0000 | 0.0000 | 0.0000 | 0.0000 |
| r. Mandibular | 0.0000 | 0.0000 | 0.0000 | 0.0000 |
| Muscle, biceps femoris | 0.0318^b^ | 0.0000 | 0.0022 | 0.0003 |
| Muscle, diaphragm | 0.0010 | 0.0000 | 0.0000 | 0.0001 |
| Pancreas | 0.0000 | 0.0000 | 0.0000 | 0.0000 |
| r. Eye | 0.0000 | 0.0000 | 0.0000 | 0.0000 |
| r. Optic nerve | 0.0000 | 0.0000 | 0.0000 | 0.0000 |
| DRG cervical^c^ |  | 0.0000 | 0.0000 | 0.0000 |
| DRG thoracic | 0.0005 | 0.0000 | 0.0000 | 0.0000 |
| DRG lumbar | 0.0000 | 0.0000 | 0.0000 | 0.0000 |
| DRG sacral | 0.0000 | 0.0000 | 0.0000 | 0.0000 |
| Blood pellet | 0.0000 | 0.0190 | 0.0000 | 0.0000 |

Abbreviations: CNV, copy number variation or the concentration (or ratio) of the target gene (GFP)/concentration of the *CFTR* reference gene; ddPCR, droplet digital polymerase chain reaction; DRG, dorsal root ganglion; LP, lumbar puncture; scAAV9-CB-GFP, self-complementary adeno-associated virus serotype 9–chicken β-actin promoter–green fluorescent protein.

^a^All samples from Group 1 are clean of green fluorescent protein vector genome, and all are below limit of quantification.

^b^Sample P0001 muscle, biceps femoris was tested twice, and both runs gave a low droplet count in two of three wells. Thus, the result was calculated from two working wells of the two runs.

^c^P0001 DRG cervical sample amount was not enough for analysis.

**Supplemental Table 14. Individual Animal Data for scAAV9-CB-GFP Vector Genome ddPCR Quantitation in ICM Control Animals (Group 2).**

|  | **Average CNV of animal** | | | |
| --- | --- | --- | --- | --- |
| **Group 2 (ICM, vehicle)** | **P0101** | **P0102** | **P0103** | **P0104** |
| r. Prefrontal cortex | 0.0000 | 0.0000 | 0.0000 | 0.0000 |
| r. Temporal cortex-entorhinal cortex | 0.0000 | 0.0000 | 0.0000 | 0.0000 |
| r. Caudate nucleus | 0.0000 | 0.0000 | 0.0000 | 0.0000 |
| r. Putamen (basal ganglia) | 0.0000 | 0.0000 | 0.0000 | 0.0000 |
| Cingulate gyrus | 0.0000 | 0.0006 | 0.0000 | 0.0000 |
| Corpus callosum | 0.0000 | 0.0000 | 0.0000 | 0.0000 |
| r. Temporal cortex-auditory cortex | 0.0000 | 0.0000 | 0.0003 | 0.0000 |
| r. Parietal cortex | 0.0000 | 0.0000 | 0.0000 | 0.0000 |
| r. Thalamus | 0.0000 | 0.0000 | 0.0000 | 0.0000 |
| r. Hypothalamus | 0.0000 | 0.0000 | 0.0000 | 0.0000 |
| r. Hippocampus | 0.0000 | 0.0000 | 0.0000 | 0.0000 |
| r. Amygdala | 0.0000 | 0.0000 | 0.0044 | 0.0000 |
| r. Substantia nigra | 0.0000 | 0.0003 | 0.0000 | 0.0000 |
| r. Pons | 0.0000 | 0.0000 | 0.0000 | 0.0000 |
| r. Cerebellum | 0.0000 | 0.0000 | 0.0000 | 0.0000 |
| r. Occipital cortex: primary visual cortex | 0.0000 | 0.0000 | 0.0002 | 0.0000 |
| r. Deep cerebellar nuclei | 0.0000 | 0.0000 | 0.0000 | 0.0000 |
| Heart | 0.0003 | 0.0000 | 0.0000 | 0.0000 |
| Spinal cord (cervical) | 0.0002 | 0.0002 | 0.0000 | 0.0000 |
| Spinal cord (thoracic) | 0.0000 | 0.0000 | 0.0000 | 0.0000 |
| Spinal cord (lumbar) | 0.0002 | 0.0000 | 0.0001 | 0.0000 |
| Spinal cord (sacral) | 0.0000 | 0.0000 | 0.0000 | 0.0000 |
| r. Trigeminal ganglia/nerve | 0.0000 | 0.0000 | 0.0000 | 0.0000 |
| Liver | 0.0000 | 0.0000 | 0.0000 | 0.0000 |
| Lung | 0.0000 | 0.0000 | 0.0000 | 0.0000 |
| Kidney (right pole) | 0.0000 | 0.0000 | 0.0000 | 0.0000 |
| r. Ovary | 0.0000 | 0.0000 | 0.0000 | 0.0000 |
| Spleen | 0.0000 | 0.0000 | 0.0000 | 0.0000 |
| r. Mandibular | 0.0000 | 0.0000 | 0.0000 | 0.0000 |
| Muscle, biceps femoris | 0.0000 | 0.0000 | 0.0000 | 0.0000 |
| Muscle, diaphragm | 0.0000 | 0.0000 | 0.0000 | 0.0000 |
| Pancreas | 0.0000 | 0.0000 | 0.0000 | 0.0000 |
| r. Eye | 0.0000 | 0.0000 | 0.0000 | 0.0000 |
| r. Optic nerve | 0.0000 | 0.0000 | 0.0000 | 0.0000 |
| DRG cervical | 0.0000 | 0.0000 | 0.0000 | 0.0000 |
| DRG thoracic | 0.0000 | 0.0000 | 0.0000 | 0.0000 |
| DRG lumbar | 0.0000 | 0.0000 | 0.0000 | 0.0000 |
| DRG sacral | 0.0000 | 0.0000 | 0.0000 | 0.0006 |
| Blood pellet | 0.0000 | 0.0000 | 0.0000 | 0.0035 |

Abbreviations: CNV, copy number variation or the concentration (or ratio) of the target gene (GFP)/concentration of the *CFTR* reference gene; ddPCR, droplet digital polymerase chain reaction; DRG, dorsal root ganglion; ICM, intracisterna magna; scAAV9-CB-GFP, self-complementary adeno-associated virus serotype 9–chicken β-actin promoter–green fluorescent protein.

**Supplemental Table 15. Individual Animal Data for scAAV9-CB-GFP Vector Genome ddPCR Quantitation in Intrathecal-LP-treated Animals at 1.0×10^13^ (Group 3).**

|  | **Average CNV of animal** | | | |
| --- | --- | --- | --- | --- |
| **Group 3  (Intrathecal-LP, dosed 1.0×10^13^ vg/animal)** | **P0201** | **P0202** | **P0203** | **P0204** |
| r. Prefrontal cortex | 0.9495 | 0.3564 | 0.0564 | 0.0174 |
| r. Temporal cortex-entorhinal cortex | 0.1922 | 1.0945 | 0.0286 | BLOQ |
| r. Caudate nucleus | BLOQ | BLOQ | 0.0389 | BLOQ |
| r. Putamen (basal ganglia) | 0.0583 | BLOQ | 0.0368 | BLOQ |
| Cingulate gyrus | 0.1230 | 0.0485 | 0.0549 | BLOQ |
| Corpus callosum | BLOQ | BLOQ | BLOQ | BLOQ |
| r. Temporal cortex-auditory cortex | 0.5891 | 0.0652 | BLOQ | BLOQ |
| r. Parietal cortex | 0.0875 | 0.6027 | BLOQ | 0.0143 |
| r. Thalamus | BLOQ | 0.0401 | 0.0606 | BLOQ |
| r. Hypothalamus | 0.0317 | 0.9852 | 0.0347 | BLOQ |
| r. Hippocampus | 0.2202 | 0.0468 | BLOQ | 0.0234 |
| r. Amygdala | 0.0475 | 2.0750 | 0.0408 | 0.0323 |
| r. Substantia nigra | BLOQ | 0.0851 | BLOQ | BLOQ |
| r. Pons | 0.3450 | 0.0403 | 0.0304 | 0.0347 |
| r. Cerebellum | BLOQ | 0.0812 | 0.0037 | BLOQ |
| r. Occipital cortex: primary visual cortex | 0.2710 | 0.5697 | 0.0278 | BLOQ |
| r. Deep cerebellar nuclei | 0.1383 | 0.1533 | BLOQ | BLOQ |
| Heart | 0.2005 | 0.3917 | 0.3433^a^ | 0.2228 |
| Spinal cord (cervical) | 1.4635 | 1.4611 | 0.0467 | 0.0286 |
| Spinal cord (thoracic) | 1.9625 | 3.1362 | 1.3864 | 0.1014 |
| Spinal cord (lumbar) | 1.7220 | 1.8322 | 1.7795 | 0.2174 |
| Spinal cord (sacral) | 2.4587 | 0.7916 | 1.6486 | 1.7834 |
| r. Trigeminal ganglia/nerve | 1.8537 | 0.0928 | 0.0557 | 0.0241 |
| Liver | 8.4772 | 36.5169^a^ | 15.3240^a^ | 9.5990 |
| Lung | 0.1156 | 0.3000 | 0.1127 | 0.1995 |
| Kidney (right pole) | 0.0751 | 0.2777 | 0.1313 | 0.1288 |
| r. Ovary | 0.0116 | 0.0485 | 0.0183 | 0.0289 |
| Spleen | 0.1238 | 1.9186 | 0.4423 | 0.3344 |
| r. Mandibular | 0.2687 | 0.7107 | 1.2991 | 0.5253 |
| Muscle, biceps femoris | 0.0407 | 0.0835 | 0.0599 | 0.0408 |
| Muscle, diaphragm | 0.0394 | 0.1398 | 0.1734 | 0.1034 |
| Pancreas | 0.0046 | 0.0323 | 0.1427 | 0.0416 |
| r. Eye | 0.0173 | 0.0824 | 0.0670 | 0.0737 |
| r. Optic nerve | 0.5226 | 3.1746 | BLOQ | 0.0148 |
| DRG cervical | 2.8520 | 1.2906 | 4.7890 | 0.6311 |
| DRG thoracic | 1.4569 | 1.5596 | 1.5059 | 0.3307 |
| DRG lumbar | 1.9098 | 0.6382 | 5.7345 | 1.1865 |
| DRG sacral | 10.0642 | 5.1663 | 3.4770 | 1.3967 |
| Blood pellet | 0.1566 | 3.2823 | 1.6433 | 1.4824 |

Abbreviations: BLOQ, below limit of quantification; CNV, copy number variation or the concentration (or ratio) of the target gene (GFP)/concentration of the *CFTR* reference gene; ddPCR, droplet digital polymerase chain reaction; DRG, dorsal root ganglion; LP, lumbar puncture; scAAV9-CB-GFP, self-complementary adeno-associated virus serotype 9–chicken β-actin promoter–green fluorescent protein; vg, vector genomes.

^a^Results reported from repeat testing.

**Supplemental Table 16. Individual Animal Data for scAAV9-CB-GFP Vector Genome ddPCR Quantitation in Intrathecal-LP-treated Animals at 3.0×10^13^ (Group 4).**

|  | **Average CNV of animal** | | | |
| --- | --- | --- | --- | --- |
| **Group 4  (Intrathecal-LP, dosed 3.0×10^13^ vg/animal)** | **P0301** | **P0302** | **P0303** | **P0304** |
| r. Prefrontal cortex | 0.4698 | 1.5770 | 0.0891 | 0.9413 |
| r. Temporal cortex-entorhinal cortex | 0.0961 | 0.1007 | BLOQ | 1.0904 |
| r. Caudate nucleus | 0.0911 | 0.0406 | BLOQ | 0.0592 |
| r. Putamen (basal ganglia) | 0.0922 | 0.0494 | BLOQ | 0.0800 |
| Cingulate gyrus | 0.1261 | 0.5538 | 0.0473 | 0.0762 |
| Corpus callosum | 0.0564 | 0.0306 | 0.0251 | 0.1766 |
| r. Temporal cortex-auditory cortex | 0.2930 | 2.2631 | 0.0847 | 0.8997 |
| r. Parietal cortex | 0.3307 | 2.9375 | 0.0293 | 0.4913 |
| r. Thalamus | 0.0913 | 0.0698 | 0.0586 | 0.0776 |
| r. Hypothalamus | 0.0847 | 0.0577 | BLOQ | 0.4485 |
| r. Hippocampus | 0.8047 | 2.3240 | 0.0903 | 0.1169 |
| r. Amygdala | 0.3997 | 0.0364 | 0.0930 | 0.1173 |
| r. Substantia nigra | 0.1107 | 0.0792 | BLOQ | 0.0675 |
| r. Pons | 0.7479 | 0.1080 | 0.0862 | 0.4435 |
| r. Cerebellum | 0.0917 | 2.6109 | 0.0127 | 0.0764 |
| r. Occipital cortex: primary visual cortex | 0.4329 | 0.7177^a^ | 0.0305 | 0.4733 |
| r. Deep cerebellar nuclei | 0.1654 | 1.8330 | 0.0268 | 0.1568 |
| Heart | 2.0646 | 0.8239 | 1.0816 | 1.2648 |
| Spinal cord (cervical) | 2.9914 | 1.6266 | 0.8143 | 1.6651 |
| Spinal cord (thoracic) | 3.8851 | 4.0010 | 1.3693 | 4.0408 |
| Spinal cord (lumbar) | 4.0579 | 3.3165 | 1.1188 | 4.8898 |
| Spinal cord (sacral) | 3.8740 | 3.8839 | 1.2016 | 5.6402 |
| r. Trigeminal ganglia/nerve | 5.5802 | 8.7407 | 6.0601 | 0.7078 |
| Liver | 82.9586 | 61.2867 | 87.1194 | 4.8644 |
| Lung | 1.0980 | 0.6012 | 0.4159 | 0.8271 |
| Kidney (right pole) | 0.8397 | 0.3922 | 0.4392 | 0.4049 |
| r. Ovary | 0.1226 | 0.0818 | 0.0665 | 0.1293 |
| Spleen | 0.7127 | 0.5002 | 0.5328 | 0.7707 |
| r. Mandibular | 2.8821 | 3.3946 | 0.9552 | 4.5248 |
| Muscle, biceps femoris | 0.3272 | 0.2509 | 0.2221 | 0.3968 |
| Muscle, diaphragm | 0.3174 | 0.6012 | 10.0059 | 0.4672 |
| Pancreas | 0.3580 | 0.1309^a^ | 0.2087 | 0.1092 |
| r. Eye | 0.2173 | 0.3798 | 0.0731 | 0.2258 |
| r. Optic nerve | 0.9629 | 4.9974 | 0.0353 | 0.3270 |
| DRG cervical | 2.3952 | 3.8692 | 4.0284 | 11.9361 |
| DRG thoracic | 2.8536 | 5.9448 | 2.0342 | 1.6905 |
| DRG lumbar | 0.5900 | 3.9398 | 2.7230 | 3.6623 |
| DRG sacral | 2.4660 | 7.2205 | 4.6401 | 11.3007 |
| Blood pellet | 7.8610 | 20.5327 | 5.9682 | 4.4654 |

Abbreviations: BLOQ, below limit of quantification; CNV, copy number variation or the concentration (or ratio) of the target gene (GFP)/concentration of the *CFTR* reference genes; ddPCR, droplet digital polymerase chain reaction; DRG, dorsal root ganglion; LP, lumbar puncture; scAAV9-CB-GFP, self-complementary adeno-associated virus serotype 9–chicken β-actin promoter–green fluorescent protein; vg, vector genomes.

^a^Results reported from repeat testing.

**Supplemental Table 17. Individual Animal Data for scAAV9-CB-GFP Vector Genome ddPCR Quantitation in ICM-treated Animals at 1.0×10^13^ (Group 5).**

|  | **Average CNV of animal** | | | |
| --- | --- | --- | --- | --- |
| **Group 5  (ICM, dosed 1.0×10^13^ vg/animal)** | **P0401** | **P0402** | **P0403** | **P0404** |
| r. Prefrontal cortex | 0.0902 | 0.0396 | 0.0573 | 0.0428 |
| r. Temporal cortex-entorhinal cortex | 0.1371 | 1.8580 | BLOQ | 1.1794 |
| r. Caudate nucleus | BLOQ | BLOQ | BLOQ | 0.0307 |
| r. Putamen (basal ganglia) | BLOQ | 0.0265 | BLOQ | 0.0280 |
| Cingulate gyrus | 0.5307 | BLOQ | BLOQ | BLOQ |
| Corpus callosum | BLOQ | BLOQ | BLOQ | BLOQ |
| r. Temporal cortex-auditory cortex | 0.1445 | 0.2676 | 0.0390 | 0.1674 |
| r. Parietal cortex | 1.3248 | 0.0394 | BLOQ | 0.0719 |
| r. Thalamus | BLOQ | BLOQ | BLOQ | BLOQ |
| r. Hypothalamus | BLOQ | BLOQ | BLOQ | 0.0372 |
| r. Hippocampus | 0.3362 | 0.0333 | BLOQ | 0.0299 |
| r. Amygdala | 0.6895 | BLOQ | BLOQ | 0.0484 |
| r. Substantia nigra | 0.8298 | BLOQ | 0.1161 | 0.4535 |
| r. Pons | 0.0626 | BLOQ | BLOQ | 0.0722 |
| r. Cerebellum | 0.0084 | 0.0454 | 0.0039 | 0.0070 |
| r. Occipital cortex: primary visual cortex | 0.0525 | 0.0147 | BLOQ | 0.1072 |
| r. Deep cerebellar nuclei | 0.1428 | BLOQ | BLOQ | 0.0155 |
| Heart | 0.0692 | 0.1225 | 0.1115 | 0.1362 |
| Spinal cord (cervical) | 1.2619 | 0.1315 | 0.2567 | 0.3515 |
| Spinal cord (thoracic) | 3.9535 | 1.9925 | 0.5426 | 0.6433 |
| Spinal cord (lumbar) | 3.3310 | 1.0921 | 0.0997 | 1.8453 |
| Spinal cord (sacral) | 25.8917 | 0.0831 | 0.2050 | 0.3619 |
| r. Trigeminal ganglia/nerve | 4.6246 | 1.9333 | 0.8968 | 6.9998 |
| Liver | 14.7327 | 15.7683 | 6.4045 | 3.5399 |
| Lung | 0.0992 | 0.2062 | 0.0550 | 0.1911 |
| Kidney (right pole) | 0.0749 | 0.1414 | 0.0604 | 0.1149 |
| r. Ovary | 0.0099 | 0.0201 | 0.0115 | 0.0249 |
| Spleen | 0.2351 | 0.3600 | 0.1995 | 0.2070 |
| r. Mandibular | 0.7049 | 13.9120^b^ | 2.0130 | 9.3292 |
| Muscle, biceps femoris | BLOQ | 0.0935 | 0.0446 | 0.0358 |
| Muscle, diaphragm | 1.7450 | 0.0483 | 0.0489 | 0.0532 |
| Pancreas | 0.0178 | 0.0348 | 0.0647 | 0.0201 |
| r. Eye | 0.0179 | 0.1941 | 0.0817 | 0.0222 |
| r. Optic nerve | 0.6232 | 0.4441 | 1.4581 | 0.0229 |
| DRG cervical | 1.8717 | 3.7188 | 1.9478 | 8.2217 |
| DRG thoracic | 2.0904 | 2.6234 | 10.0533 | 3.4612 |
| DRG lumbar | 3.7752 | 10.7115 | 8.7568 | 6.0020 |
| DRG sacral | 20.2388 | 0.0628 | 0.0975 | 0.7155 |
| Blood pellet | 1.8220 | 1.6488 | 1.6042 | 1.2380 |

Abbreviations: BLOQ, below limit of quantification; CNV, copy number variation or the concentration (or ratio) of the target gene (GFP)/concentration of the *CFTR* reference gene; ddPCR, droplet digital polymerase chain reaction; DRG, dorsal root ganglion; ICM, intracisterna magna; scAAV9-CB-GFP, self-complementary adeno-associated virus serotype 9–chicken β-actin promoter–green fluorescent protein; vg, vector genomes.

^a^All samples from Group 2 are clean of the GFP vector genome, all below LOQ

^b^Results reported from repeat analysis.

**Supplemental Table 18. Individual Animal Data for scAAV9-CB-GFP Vector Genome ddPCR Quantitation in ICM-treated Animals at 3.0×10^13^ (Group 6).**

|  | **Average CNV of animal** | | | |
| --- | --- | --- | --- | --- |
| **Group 6  (ICM, dosed 3×10^13^** **vg/animal)** | **P0501** | **P0502** | **P0503** | **P0504** |
| r. Prefrontal cortex | 0.4168 | 0.0540 | 0.1622 | 0.1890 |
| r. Temporal cortex-entorhinal cortex | 0.1532 | 0.1264 | 0.1381 | 0.1552 |
| r. Caudate nucleus | 0.0918 | BLOQ | 0.0588 | 0.1796 |
| r. Putamen (basal ganglia) | 0.1043 | 0.0415 | 0.1019 | 0.2129 |
| Cingulate gyrus | 1.1188 | 0.1067 | 0.1800 | 0.0759 |
| Corpus callosum | 0.6933 | 0.0406 | 0.0351 | 1.1180 |
| r. Temporal cortex-auditory cortex | 1.1052 | 0.1655 | 0.0770 | 1.9484 |
| r. Parietal cortex | 0.4337 | 0.0281 | 0.1717 | 0.7426 |
| r. Thalamus | 0.0675 | 0.0335 | 0.1044 | 0.1689 |
| r. Hypothalamus | 0.0683 | 0.2689 | 0.0800 | 0.2706 |
| r. Hippocampus | 0.7414 | 0.1435 | 0.1007 | 0.3761 |
| r. Amygdala | 2.3518 | 0.0774 | 0.2006 | 0.1893 |
| r. Substantia nigra | 0.0972 | 0.0445 | 0.1692 | 0.1449 |
| r. Pons | 0.1227 | 0.0656 | 0.2874 | 0.1842 |
| r. Cerebellum | 0.0269 | 0.0047 | 0.0174 | 0.0234 |
| r. Occipital cortex: primary visual cortex | 0.1818 | 0.0662 | 0.0881 | 0.5801 |
| r. Deep cerebellar nuclei | 0.0866 | 0.0080 | 0.0624 | 0.0751 |
| Heart | 0.4748 | 0.5981 | 0.6759 | 0.8413 |
| Spinal cord (cervical) | 1.8513 | 0.8057 | 0.3447 | 1.4639 |
| Spinal cord (thoracic) | 4.1330 | 0.4968 | 6.6211 | 2.6426 |
| Spinal cord (lumbar) | 3.8024 | 2.7435 | 6.6057 | 3.7987 |
| Spinal cord (sacral) | 11.3787 | 1.2135 | 14.0692 | 6.9721 |
| r. Trigeminal ganglia/nerve | 1.0678 | 2.1965 | 2.0657 | 0.3885 |
| Liver | 103.6373 | 59.0015 | 33.6182 | 110.8759 |
| Lung | 0.5790 | 0.3208 | 0.2108 | 0.6015 |
| Kidney (right pole) | 0.2840 | 0.2627 | 0.3453 | 0.3833 |
| r. Ovary | 0.1036 | 0.0418 | 0.0689 | 0.0469 |
| Spleen | 0.4134 | 0.3127 | 0.3231 | 2.3425 |
| r. Mandibular | 1.3653 | 2.5690 | 6.8984 | 1.1354 |
| Muscle, biceps femoris | 0.2112 | 0.1190 | 0.2127 | 0.1798 |
| Muscle, diaphragm | 0.4548 | 0.1124 | 0.2516 | 0.5390 |
| Pancreas | 0.4807 | 0.0344 | 0.1075 | 0.2628 |
| r. Eye | 0.2696 | 0.1965 | 0.1836 | 0.1099 |
| r. Optic nerve | 2.2151 | 0.8897 | 0.0516 | 0.5472 |
| DRG cervical | 3.0819 | 1.7438 | 1.7888 | 1.5118 |
| DRG thoracic | 2.7003 | 4.7475 | 9.6906 | 2.3972 |
| DRG lumbar | 3.9016 | 6.2240 | 7.2308 | 12.4546 |
| DRG sacral | 2.6180 | 2.2354 | 2.9535 | 21.1183 |
| Blood pellet | 19.3202 | 2.3640 | 3.0634 | 2.7882 |

Abbreviations: BLOW, below limit of quantification; CNV, copy number variation or the concentration (or ratio) of the target gene (GFP)/concentration of the *CFTR* reference gene; ddPCR, droplet digital polymerase chain reaction; DRG, dorsal root ganglion; ICM, intracisterna magna; scAAV9-CB-GFP, self-complementary adeno-associated virus serotype 9–chicken β-actin promoter–green fluorescent protein; vg, vector genomes.

^a^All samples from Group 2 are clean of the GFP vector genome, all below LOQ.

**Supplemental Table 19. Summary Statistics of GFP Protein Expression in Group 1 (Vehicle) Intrathecal-LP-Dosed Animals at 28 Days.**

|  | **GFP protein concentration (pg/mg of protein)** | | | | | |  |
| --- | --- | --- | --- | --- | --- | --- | --- |
|  | **Median** | **Minimum** | **Maximum** | **Mean** | **SD** | **CV** | **N** |
| Prefrontal cortex | 0 | 0 | 0 | 0 | N/A | N/A | 4 |
| Temporal cortex-entorhinal cortex | 0 | 0 | 0 | 0 | N/A | N/A | 4 |
| Caudate nucleus | 0 | 0 | 0 | 0 | N/A | N/A | 4 |
| Putamen (basal ganglia) | 0 | 0 | 0 | 0 | N/A | N/A | 4 |
| Cingulate gyrus | 0 | 0 | 0 | 0 | N/A | N/A | 4 |
| Corpus callosum | 0 | 0 | 0 | 0 | N/A | N/A | 4 |
| Temporal cortex-auditory cortex | 0 | 0 | 0 | 0 | N/A | N/A | 4 |
| Parietal cortex | 0 | 0 | 0 | 0 | N/A | N/A | 4 |
| Thalamus | 0 | 0 | 0 | 0 | N/A | N/A | 4 |
| Hypothalamus | 0 | 0 | 0 | 0 | N/A | N/A | 4 |
| Hippocampus | 0 | 0 | 0 | 0 | N/A | N/A | 4 |
| Amygdala | 0 | 0 | 0 | 0 | N/A | N/A | 4 |
| Substantia nigra | 0 | 0 | 0 | 0 | N/A | N/A | 4 |
| Pons | 0 | 0 | 0 | 0 | N/A | N/A | 4 |
| Cerebellum | 0 | 0 | 0 | 0 | N/A | N/A | 4 |
| Occipital cortex: primary visual cortex | 0 | 0 | 0 | 0 | N/A | N/A | 4 |
| Deep cerebellar nuclei | 0 | 0 | 0 | 0 | N/A | N/A | 4 |
| Heart | 0 | 0 | 0 | 0 | N/A | N/A | 4 |
| Spinal cord (cervical) | 0 | 0 | 0 | 0 | N/A | N/A | 4 |
| Spinal cord (thoracic) | 0 | 0 | 0 | 0 | N/A | N/A | 4 |
| Spinal cord (lumbar) | 0 | 0 | 0 | 0 | N/A | N/A | 4 |
| Spinal cord (sacral) | 0 | 0 | 0 | 0 | N/A | N/A | 4 |
| Trigeminal ganglia/nerve | 0 | 0 | 0 | 0 | N/A | N/A | 4 |
| Liver | 0 | 0 | 0 | 0 | N/A | N/A | 4 |
| Lung | 0 | 0 | 0 | 0 | N/A | N/A | 4 |
| Kidney  (right pole) | 0 | 0 | 0 | 0 | N/A | N/A | 4 |
| Spleen | 0 | 0 | 0 | 0 | N/A | N/A | 4 |
| Mandibular | 0 | 0 | 0 | 0 | N/A | N/A | 4 |
| Muscle, biceps femoris | 0 | 0 | 0 | 0 | N/A | N/A | 4 |
| Muscle, diaphragm | 0 | 0 | 0 | 0 | N/A | N/A | 4 |
| Pancreas | 0 | 0 | 0 | 0 | N/A | N/A | 4 |
| Eye | 0 | 0 | 0 | 0 | N/A | N/A | 4 |
| Optic nerve | 0 | 0 | 0 | 0 | N/A | N/A | 4 |
| DRG cervical | 0 | 0 | 0 | 0 | N/A | N/A | 4 |
| DRG thoracic | 0 | 0 | 0 | 0 | N/A | N/A | 4 |
| DRG lumbar | 0 | 0 | 0 | 0 | N/A | N/A | 4 |
| DRG sacral | 0 | 0 | 0 | 0 | N/A | N/A | 4 |

Abbreviations: CV, coefficient of variation; DRG, dorsal root ganglion; GFP, green fluorescent protein; LP, lumbar puncture; N/A, not applicable; pg, picogram; SD, standard deviation.

**Supplemental Table 20. Summary Statistics of GFP Protein Expression in Group 2 (Vehicle) ICM-Dosed Animals at 28 Days.**

|  | **GFP protein concentration (pg/mg of protein)** | | | | | |  |
| --- | --- | --- | --- | --- | --- | --- | --- |
|  | **Median** | **Minimum** | **Maximum** | **Mean** | **SD** | **CV** | **N** |
| Prefrontal cortex | 0 | 0 | 0 | 0 | N/A | N/A | 4 |
| Temporal cortex-entorhinal cortex | 0 | 0 | 0 | 0 | N/A | N/A | 4 |
| Caudate nucleus | 0 | 0 | 0 | 0 | N/A | N/A | 4 |
| Putamen (basal ganglia) | 0 | 0 | 0 | 0 | N/A | N/A | 4 |
| Cingulate gyrus | 0 | 0 | 0 | 0 | N/A | N/A | 4 |
| Corpus callosum | 0 | 0 | 0 | 0 | N/A | N/A | 4 |
| Temporal cortex-auditory cortex | 0 | 0 | 0 | 0 | N/A | N/A | 4 |
| Parietal cortex | 0 | 0 | 0 | 0 | N/A | N/A | 4 |
| Thalamus | 0 | 0 | 0 | 0 | N/A | N/A | 4 |
| Hypothalamus | 0 | 0 | 0 | 0 | N/A | N/A | 4 |
| Hippocampus | 0 | 0 | 0 | 0 | N/A | N/A | 4 |
| Amygdala | 0 | 0 | 0 | 0 | N/A | N/A | 4 |
| Substantia nigra | 0 | 0 | 0 | 0 | N/A | N/A | 4 |
| Pons | 0 | 0 | 0 | 0 | N/A | N/A | 4 |
| Cerebellum | 0 | 0 | 0 | 0 | N/A | N/A | 4 |
| Occipital cortex: primary visual cortex | 0 | 0 | 0 | 0 | N/A | N/A | 4 |
| Deep cerebellar nuclei | 0 | 0 | 0 | 0 | N/A | N/A | 4 |
| Heart | 0 | 0 | 0 | 0 | N/A | N/A | 4 |
| Spinal cord (cervical) | 0 | 0 | 0 | 0 | N/A | N/A | 4 |
| Spinal cord (thoracic) | 0 | 0 | 0 | 0 | N/A | N/A | 4 |
| Spinal cord (lumbar) | 0 | 0 | 0 | 0 | N/A | N/A | 4 |
| Spinal cord (sacral) | 0 | 0 | 0 | 0 | N/A | N/A | 4 |
| Trigeminal ganglia/nerve | 0 | 0 | 0 | 0 | N/A | N/A | 4 |
| Liver | 0 | 0 | 0 | 0 | N/A | N/A | 4 |
| Lung | 0 | 0 | 0 | 0 | N/A | N/A | 4 |
| Kidney (right pole) | 0 | 0 | 0 | 0 | N/A | N/A | 4 |
| Spleen | 0 | 0 | 0 | 0 | N/A | N/A | 4 |
| Mandibular | 0 | 0 | 0 | 0 | N/A | N/A | 4 |
| Muscle, biceps femoris | 0 | 0 | 0 | 0 | N/A | N/A | 4 |
| Muscle, diaphragm | 0 | 0 | 0 | 0 | N/A | N/A | 4 |
| Pancreas | 0 | 0 | 0 | 0 | N/A | N/A | 4 |
| Eye | 0 | 0 | 0 | 0 | N/A | N/A | 4 |
| Optic nerve | 0 | 0 | 0 | 0 | N/A | N/A | 4 |
| DRG cervical | 0 | 0 | 0 | 0 | N/A | N/A | 4 |
| DRG thoracic | 0 | 0 | 0 | 0 | N/A | N/A | 4 |
| DRG lumbar | 0 | 0 | 0 | 0 | N/A | N/A | 4 |
| DRG sacral | 0 | 0 | 0 | 0 | N/A | N/A | 4 |

Abbreviations: CV, coefficient of variation; DRG, dorsal root ganglion; GFP, green fluorescent protein; ICM, intracisterna magna; N/A, not applicable; pg, picogram; SD, standard deviation.

**Supplemental Table 21. Summary Statistics of GFP Protein Expression in Group 3 (1.0×10^13^ vg/animal) Intrathecal-LP-Dosed Animals at 28 Days.**

|  | **GFP protein concentration (pg/mg of protein)** | | | | | |  |
| --- | --- | --- | --- | --- | --- | --- | --- |
|  | **Median** | **Minimum** | **Maximum** | **Mean** | **SD** | **CV** | **N** |
| Prefrontal cortex | 57.44 | 27.17 | 92.19 | 58.56 | 32.70 | 55.85 | 4 |
| Temporal cortex-entorhinal cortex | 82.56 | 0.00 | 112.30 | 69.36 | 51.32 | 73.99 | 4 |
| Caudate nucleus | 0.00 | 0.00 | 28.47 | 7.12 | 14.23 | 200.00 | 4 |
| Putamen (basal ganglia) | 40.57 | 29.48 | 69.13 | 44.94 | 17.49 | 38.91 | 4 |
| Cingulate gyrus | 105.20 | 38.09 | 126.21 | 93.68 | 40.54 | 43.28 | 4 |
| Corpus callosum | 50.22 | 0.00 | 107.42 | 51.97 | 60.07 | 115.60 | 4 |
| Temporal cortex-auditory cortex | 48.30 | 42.22 | 167.09 | 76.48 | 60.50 | 79.11 | 4 |
| Parietal cortex | 154.10 | 120.29 | 184.84 | 153.33 | 28.48 | 18.57 | 4 |
| Thalamus | 45.84 | 0.00 | 103.02 | 48.68 | 43.53 | 89.44 | 4 |
| Hypothalamus | 140.26 | 56.01 | 194.40 | 132.73 | 57.17 | 43.07 | 4 |
| Hippocampus | 80.40 | 0.00 | 754.68 | 228.87 | 352.59 | 154.06 | 4 |
| Amygdala | 218.78 | 84.90 | 459.57 | 245.51 | 159.15 | 64.82 | 4 |
| Substantia nigra | 73.15 | 38.32 | 443.08 | 156.92 | 193.14 | 123.08 | 4 |
| Pons | 339.24 | 104.53 | 648.70 | 357.93 | 240.85 | 67.29 | 4 |
| Cerebellum | 156.51 | 35.72 | 405.78 | 188.63 | 155.58 | 82.48 | 4 |
| Occipital cortex: primary visual cortex | 213.74 | 45.94 | 391.56 | 216.25 | 147.49 | 68.20 | 4 |
| Deep cerebellar nuclei | 266.43 | 33.24 | 1785.45 | 587.89 | 806.30 | 137.15 | 4 |
| Heart | 4520.06 | 155.28 | 12934.27 | 5532.42 | 6398.85 | 115.66 | 4 |
| Spinal cord (cervical) | 959.41 | 187.79 | 5057.01 | 1790.90 | 2212.05 | 123.52 | 4 |
| Spinal cord (thoracic) | 2711.49 | 316.91 | 4081.62 | 2455.38 | 1761.02 | 71.72 | 4 |
| Spinal cord (lumbar) | 2545.40 | 151.31 | 6127.13 | 2842.31 | 2969.61 | 104.48 | 4 |
| Spinal cord (sacral) | 1886.39 | 83.59 | 7627.83 | 2871.05 | 3525.50 | 122.79 | 4 |
| Trigeminal ganglia/nerve | 280.67 | 36.10 | 609.34 | 301.70 | 273.53 | 90.66 | 4 |
| Liver | 1625.08 | 88.48 | 3389.46 | 1682.02 | 1842.33 | 109.53 | 4 |
| Lung | 14.63 | 0.00 | 243.49 | 68.19 | 117.68 | 172.57 | 4 |
| Kidney  (right pole) | 113.33 | 40.36 | 312.29 | 144.83 | 117.10 | 80.86 | 4 |
| Spleen | 36.98 | 0.00 | 775.24 | 212.30 | 376.91 | 177.54 | 4 |
| Mandibular | 70.27 | 0.00 | 560.14 | 175.17 | 265.06 | 151.31 | 4 |
| Muscle, biceps femoris | 12951.30 | 6402.15 | 21104.95 | 13352.43 | 6231.79 | 46.67 | 4 |
| Muscle, diaphragm | 10579.10 | 178.00 | 14712.10 | 9012.07 | 6236.73 | 69.20 | 4 |
| Pancreas | 244.14 | 33.11 | 9790.18 | 2577.90 | 4809.23 | 186.56 | 4 |
| Eye | 316.94 | 0.00 | 1197.48 | 457.84 | 565.24 | 123.46 | 4 |
| Optic nerve | 231.24 | 0.00 | 303.07 | 191.39 | 133.80 | 69.91 | 4 |
| DRG cervical | 544.25 | 51.85 | 1638.99 | 694.83 | 686.43 | 98.79 | 4 |
| DRG thoracic | 432.44 | 66.40 | 3084.29 | 1003.89 | 1401.20 | 139.58 | 4 |
| DRG lumbar | 494.07 | 128.43 | 3802.83 | 1229.85 | 1727.03 | 140.43 | 4 |
| DRG sacral | 492.84 | 33.87 | 5823.03 | 1710.65 | 2760.02 | 161.34 | 4 |

Abbreviations: CV, coefficient of variation; DRG, dorsal root ganglion; GFP, green fluorescent protein; LP, lumbar puncture; pg, picogram; SD, standard deviation; vg, vector genomes.

**Supplemental Table 22. Summary Statistics of GFP Protein Expression in Group 4 (3.0×10^13^ vg/animal) Intrathecal-LP-Dosed Animals at 28 Days.**

|  | **GFP protein concentration (pg/mg of protein)** | | | | | |  |
| --- | --- | --- | --- | --- | --- | --- | --- |
|  | **Median** | **Minimum** | **Maximum** | **Mean** | **SD** | **CV** | **N** |
| Prefrontal cortex | 137.11 | 91.02 | 249.65 | 153.72 | 69.36 | 45.12 | 4 |
| Temporal cortex-entorhinal cortex | 255.51 | 100.37 | 385.18 | 249.14 | 120.41 | 48.33 | 4 |
| Caudate nucleus | 58.30 | 44.85 | 116.72 | 69.54 | 32.14 | 46.22 | 4 |
| Putamen (basal ganglia) | 111.06 | 62.03 | 134.76 | 104.73 | 31.27 | 29.85 | 4 |
| Cingulate gyrus | 158.96 | 108.18 | 247.13 | 168.30 | 64.83 | 38.52 | 4 |
| Corpus callosum | 149.41 | 87.28 | 183.92 | 142.50 | 40.61 | 28.50 | 4 |
| Temporal cortex-auditory cortex | 271.19 | 161.51 | 440.66 | 286.14 | 117.73 | 41.14 | 4 |
| Parietal cortex | 311.78 | 88.87 | 429.35 | 285.44 | 164.29 | 57.56 | 4 |
| Thalamus | 172.50 | 145.26 | 240.32 | 182.65 | 40.75 | 22.31 | 4 |
| Hypothalamus | 249.22 | 227.56 | 549.43 | 318.86 | 154.71 | 48.52 | 4 |
| Hippocampus | 320.55 | 214.53 | 505.65 | 340.32 | 121.45 | 35.69 | 4 |
| Amygdala | 991.28 | 372.78 | 1336.87 | 923.05 | 426.76 | 46.23 | 4 |
| Substantia nigra | 402.75 | 208.53 | 1668.59 | 670.66 | 673.50 | 100.42 | 4 |
| Pons | 796.80 | 580.14 | 990.40 | 791.03 | 199.57 | 25.23 | 4 |
| Cerebellum | 712.40 | 376.39 | 868.15 | 667.33 | 207.76 | 31.13 | 4 |
| Occipital cortex: primary visual cortex | 288.83 | 127.61 | 1473.40 | 544.67 | 626.39 | 115.00 | 4 |
| Deep cerebellar nuclei | 967.68 | 235.41 | 1495.64 | 916.60 | 666.38 | 72.70 | 4 |
| Heart | 6665.81 | 1751.93 | 18169.13 | 8313.17 | 7156.83 | 86.09 | 4 |
| Spinal cord (cervical) | 1515.04 | 765.82 | 2087.25 | 1470.79 | 590.44 | 40.14 | 4 |
| Spinal cord (thoracic) | 1642.98 | 1582.93 | 2132.64 | 1750.38 | 259.94 | 14.85 | 4 |
| Spinal cord (lumbar) | 1249.84 | 776.25 | 1430.90 | 1176.71 | 296.24 | 25.18 | 4 |
| Spinal cord (sacral) | 1128.71 | 264.19 | 11844.10 | 3591.43 | 5517.18 | 153.62 | 4 |
| Trigeminal ganglia/nerve | 616.67 | 127.74 | 1244.30 | 651.34 | 536.28 | 82.34 | 4 |
| Liver | 4149.96 | 245.12 | 7691.11 | 4059.04 | 3285.71 | 80.95 | 4 |
| Lung | 198.52 | 94.89 | 360.83 | 213.19 | 119.70 | 56.15 | 4 |
| Kidney  (right pole) | 212.94 | 79.76 | 364.52 | 217.54 | 135.34 | 62.21 | 4 |
| Spleen | 85.53 | 44.11 | 112.25 | 81.86 | 33.80 | 41.30 | 4 |
| Mandibular | 92.30 | 76.31 | 235.95 | 124.22 | 74.89 | 60.29 | 4 |
| Muscle, biceps femoris | 7450.60 | 6325.06 | 16407.99 | 9408.56 | 4754.19 | 50.53 | 4 |
| Muscle, diaphragm | 10224.39 | 6098.65 | 16837.10 | 10846.13 | 4558.77 | 42.03 | 4 |
| Pancreas | 1647.56 | 107.78 | 8693.14 | 3024.01 | 3862.95 | 127.74 | 4 |
| Eye | 453.11 | 106.64 | 662.93 | 418.95 | 285.73 | 68.20 | 4 |
| Optic nerve | 433.79 | 95.52 | 1042.49 | 501.40 | 439.96 | 87.75 | 4 |
| DRG cervical | 1111.67 | 916.86 | 1565.34 | 1176.38 | 306.89 | 26.09 | 4 |
| DRG thoracic | 977.92 | 745.08 | 1363.25 | 1016.04 | 317.21 | 31.22 | 4 |
| DRG lumbar | 809.85 | 33.05 | 1190.04 | 710.70 | 505.10 | 71.07 | 4 |
| DRG sacral | 580.53 | 145.73 | 1547.73 | 713.63 | 600.20 | 84.11 | 4 |

Abbreviations: CV, coefficient of variation; DRG, dorsal root ganglion; GFP, green fluorescent protein; LP, lumbar puncture; pg, picogram; SD, standard deviation; vg, vector genomes.

**Supplemental Table 23. Summary Statistics of GFP Protein Expression in Group 5 (1.0×10^13^ vg/animal) ICM-Dosed Animals at 28 Days.**

|  | **GFP protein concentration (pg/mg of protein)** | | | | | |  |
| --- | --- | --- | --- | --- | --- | --- | --- |
|  | **Median** | **Minimum** | **Maximum** | **Mean** | **SD** | **CV** | **N** |
| Prefrontal cortex | 33.19 | 28.74 | 125.18 | 55.08 | 46.79 | 84.95 | 4 |
| Temporal cortex-entorhinal cortex | 36.68 | 25.42 | 539.09 | 159.47 | 253.17 | 158.76 | 4 |
| Caudate nucleus | 18.53 | 10.59 | 175.53 | 55.80 | 79.93 | 143.25 | 4 |
| Putamen (basal ganglia) | 23.52 | 21.02 | 24.09 | 23.04 | 1.41 | 6.10 | 4 |
| Cingulate gyrus | 43.04 | 27.29 | 325.16 | 109.63 | 143.88 | 131.24 | 4 |
| Corpus callosum | 37.28 | 15.33 | 2288.41 | 594.57 | 1129.27 | 189.93 | 4 |
| Temporal cortex-auditory cortex | 44.32 | 26.55 | 395.96 | 127.79 | 179.22 | 140.25 | 4 |
| Parietal cortex | 57.04 | 33.61 | 1491.96 | 409.91 | 721.46 | 176.00 | 4 |
| Thalamus | 35.78 | 30.09 | 44.21 | 36.47 | 6.62 | 18.15 | 4 |
| Hypothalamus | 61.39 | 33.40 | 205.34 | 90.38 | 78.01 | 86.31 | 4 |
| Hippocampus | 36.63 | 26.52 | 51.62 | 37.85 | 10.44 | 27.57 | 4 |
| Amygdala | 138.37 | 25.54 | 2413.15 | 678.86 | 1159.73 | 170.84 | 4 |
| Substantia nigra | 77.70 | 48.41 | 198.20 | 100.50 | 66.78 | 66.45 | 4 |
| Pons | 175.90 | 108.60 | 405.00 | 216.35 | 134.60 | 62.22 | 4 |
| Cerebellum | 74.72 | 31.00 | 290.89 | 117.83 | 119.07 | 101.05 | 4 |
| Occipital cortex: primary visual cortex | 102.63 | 44.22 | 388.25 | 159.43 | 157.28 | 98.65 | 4 |
| Deep cerebellar nuclei | 197.81 | 55.68 | 4458.40 | 1227.43 | 2155.50 | 175.61 | 4 |
| Heart | 946.08 | 689.65 | 2501.72 | 1270.89 | 831.26 | 65.41 | 4 |
| Spinal cord (cervical) | 2111.12 | 410.56 | 5605.88 | 2559.67 | 2186.86 | 85.44 | 4 |
| Spinal cord (thoracic) | 1655.40 | 491.13 | 5020.31 | 2205.56 | 2116.95 | 95.98 | 4 |
| Spinal cord (lumbar) | 975.68 | 339.94 | 2897.53 | 1297.21 | 1188.67 | 91.63 | 4 |
| Spinal cord (sacral) | 548.70 | 185.25 | 8872.87 | 2538.88 | 4226.32 | 166.46 | 4 |
| Trigeminal ganglia/nerve | 502.92 | 139.29 | 2502.31 | 911.86 | 1075.75 | 117.97 | 4 |
| Liver | 2542.28 | 91.36 | 5694.51 | 2717.61 | 2667.68 | 98.16 | 4 |
| Lung | 87.19 | 32.06 | 117.93 | 81.09 | 37.50 | 46.24 | 4 |
| Kidney  (right pole) | 82.50 | 38.65 | 122.92 | 81.64 | 38.92 | 47.67 | 4 |
| Spleen | 194.92 | 24.32 | 820.39 | 308.64 | 357.31 | 115.77 | 4 |
| Mandibular | 120.65 | 43.04 | 439.65 | 181.00 | 178.99 | 98.89 | 4 |
| Muscle, biceps femoris | 12758.10 | 363.70 | 19349.98 | 11307.47 | 7957.84 | 70.38 | 4 |
| Muscle, diaphragm | 10171.29 | 1656.53 | 13573.96 | 8893.27 | 5123.88 | 57.62 | 4 |
| Pancreas | 5840.27 | 3430.71 | 6057.30 | 5109.42 | 1457.86 | 28.53 | 4 |
| Eye | 93.62 | 19.57 | 2371.79 | 644.65 | 1152.88 | 178.84 | 4 |
| Optic nerve | 531.98 | 58.32 | 1282.22 | 601.12 | 546.20 | 90.86 | 4 |
| DRG cervical | 2438.46 | 539.20 | 16688.75 | 5526.22 | 7495.51 | 135.64 | 4 |
| DRG thoracic | 3186.24 | 481.23 | 5496.27 | 3087.50 | 2124.67 | 68.82 | 4 |
| DRG lumbar | 1071.25 | 350.04 | 2623.12 | 1278.91 | 1062.78 | 83.10 | 4 |
| DRG sacral | 994.12 | 87.65 | 2470.71 | 1136.65 | 1094.22 | 96.27 | 4 |

Abbreviations: CV, coefficient of variation; DRG, dorsal root ganglion; GFP, green fluorescent protein; ICM, intracisterna magna; pg, picogram; SD, standard deviation; vg, vector genomes.

**Supplemental Table 24. Summary Statistics of GFP Protein Expression in Group 6 (3.0×10^13^ vg/animal) ICM-Dosed Animals at 28 Days.**

|  | **GFP protein concentration (pg/mg of protein)** | | | | | |  |
| --- | --- | --- | --- | --- | --- | --- | --- |
|  | **Median** | **Minimum** | **Maximum** | **Mean** | **SD** | **CV** | **N** |
| Prefrontal cortex | 110.28 | 50.24 | 535.04 | 201.46 | 226.62 | 112.49 | 4 |
| Temporal cortex-entorhinal cortex | 117.23 | 37.01 | 697.10 | 242.14 | 308.73 | 127.50 | 4 |
| Caudate nucleus | 54.74 | 18.76 | 186.17 | 78.60 | 74.39 | 94.64 | 4 |
| Putamen (basal ganglia) | 66.04 | 37.36 | 206.63 | 94.02 | 78.37 | 83.36 | 4 |
| Cingulate gyrus | 138.42 | 62.90 | 334.33 | 168.52 | 118.86 | 70.53 | 4 |
| Corpus callosum | 111.88 | 71.99 | 313.98 | 152.43 | 112.45 | 73.77 | 4 |
| Temporal cortex-auditory cortex | 116.23 | 83.73 | 499.69 | 203.97 | 198.06 | 97.10 | 4 |
| Parietal cortex | 213.44 | 72.16 | 457.45 | 239.13 | 160.32 | 67.04 | 4 |
| Thalamus | 101.70 | 67.72 | 388.41 | 164.88 | 149.88 | 90.90 | 4 |
| Hypothalamus | 122.90 | 50.95 | 483.36 | 195.03 | 199.72 | 102.40 | 4 |
| Hippocampus | 138.44 | 110.06 | 3116.40 | 875.84 | 1493.80 | 170.56 | 4 |
| Amygdala | 412.46 | 147.13 | 2217.69 | 797.44 | 966.03 | 121.14 | 4 |
| Substantia nigra | 239.45 | 122.97 | 604.64 | 301.63 | 209.53 | 69.46 | 4 |
| Pons | 740.71 | 239.90 | 1531.71 | 813.26 | 617.96 | 75.99 | 4 |
| Cerebellum | 483.81 | 64.99 | 1181.85 | 553.61 | 521.36 | 94.17 | 4 |
| Occipital cortex: primary visual cortex | 237.48 | 104.96 | 2686.84 | 816.69 | 1248.49 | 152.87 | 4 |
| Deep cerebellar nuclei | 662.71 | 205.05 | 1058.76 | 647.31 | 387.38 | 59.84 | 4 |
| Heart | 3329.16 | 920.88 | 6953.07 | 3633.07 | 2595.53 | 71.44 | 4 |
| Spinal cord (cervical) | 5775.95 | 875.67 | 10011.78 | 5609.84 | 4555.80 | 81.21 | 4 |
| Spinal cord (thoracic) | 1178.26 | 615.56 | 13533.14 | 4126.31 | 6279.25 | 152.18 | 4 |
| Spinal cord (lumbar) | 1065.01 | 263.37 | 12950.41 | 3835.95 | 6095.92 | 158.92 | 4 |
| Spinal cord (sacral) | 1335.85 | 311.75 | 17547.83 | 5132.82 | 8291.60 | 161.54 | 4 |
| Trigeminal ganglia/nerve | 868.82 | 195.58 | 1352.54 | 821.44 | 541.56 | 65.93 | 4 |
| Liver | 5589.01 | 1062.75 | 18792.80 | 7758.39 | 8214.83 | 105.88 | 4 |
| Lung | 200.30 | 67.52 | 804.24 | 318.09 | 332.05 | 104.39 | 4 |
| Kidney  (right pole) | 986.52 | 214.62 | 5936.13 | 2030.95 | 2696.80 | 132.79 | 4 |
| Spleen | 69.64 | 18.69 | 3200.61 | 839.65 | 1574.23 | 187.49 | 4 |
| Mandibular | 221.19 | 60.78 | 2595.34 | 774.62 | 1222.35 | 157.80 | 4 |
| Muscle, biceps femoris | 9081.03 | 4633.60 | 24544.13 | 11834.95 | 8970.45 | 75.80 | 4 |
| Muscle, diaphragm | 9191.35 | 4242.27 | 22233.70 | 11214.67 | 7924.10 | 70.66 | 4 |
| Pancreas | 2941.06 | 712.59 | 8466.73 | 3765.36 | 3414.67 | 90.69 | 4 |
| Eye | 418.34 | 54.58 | 1683.38 | 643.66 | 769.99 | 119.63 | 4 |
| Optic nerve | 381.82 | 178.56 | 799.36 | 435.39 | 302.70 | 69.52 | 4 |
| DRG cervical | 1631.84 | 544.31 | 32693.17 | 9125.29 | 15726.97 | 172.34 | 4 |
| DRG thoracic | 1832.38 | 311.90 | 11506.10 | 3870.69 | 5164.19 | 133.42 | 4 |
| DRG lumbar | 1393.82 | 308.98 | 8518.05 | 2903.67 | 3804.28 | 131.02 | 4 |
| DRG sacral | 1069.44 | 191.19 | 19236.71 | 5391.69 | 9242.23 | 171.42 | 4 |

Abbreviations: CV, coefficient of variation; DRG, dorsal root ganglion; GFP, green fluorescent protein; ICM, intracisterna magna; NA, not applicable; pg, picogram; SD, standard deviation; vg, vector genomes.

###### Supplemental Table 25. Rank Order of GFP Protein Concentrations in Tissues at 28 Days Based on Median Tissue Concentrations.

|  | **Intrathecal-LP (1.0×10^13^)** | **Intrathecal-LP (3.0×10^13^)** | **ICM**  **(1.0×10^13^)** | **ICM**  **(3.0×10^13^)** |
| --- | --- | --- | --- | --- |
| Prefrontal cortex | 30 | 33 | 35 | 33 |
| Temporal cortex-entorhinal cortex | 26 | 26 | 32 | 30 |
| Caudate nucleus | 37 | 37 | 37 | 37 |
| Putamen (basal ganglia) | 34 | 34 | 36 | 36 |
| Cingulate gyrus | 25 | 31 | 30 | 28 |
| Corpus callosum | 31 | 32 | 31 | 32 |
| Temporal cortex-auditory cortex | 32 | 25 | 29 | 31 |
| Parietal cortex | 22 | 23 | 28 | 25 |
| Thalamus | 33 | 30 | 34 | 34 |
| Hypothalamus | 23 | 27 | 27 | 29 |
| Hippocampus | 27 | 22 | 33 | 27 |
| Amygdala | 19 | 11 | 19 | 20 |
| Substantia nigra | 28 | 21 | 25 | 22 |
| Pons | 13 | 15 | 18 | 16 |
| Cerebellum | 21 | 16 | 26 | 18 |
| Occipital cortex: primary visual cortex | 20 | 24 | 21 | 23 |
| Deep cerebellar nuclei | 16 | 13 | 16 | 17 |
| Heart | 3 | 3 | 12 | 5 |
| Spinal cord (cervical) | 8 | 7 | 7 | 3 |
| Spinal cord (thoracic) | 4 | 6 | 8 | 11 |
| Spinal cord (lumbar) | 5 | 8 | 11 | 13 |
| Spinal cord (sacral) | 6 | 9 | 13 | 10 |
| Trigeminal ganglia/nerve | 15 | 17 | 15 | 15 |
| Liver | 7 | 4 | 5 | 4 |
| Lung | 36 | 29 | 23 | 26 |
| Kidney (right pole) | 24 | 28 | 24 | 14 |
| Spleen | 35 | 36 | 17 | 35 |
| Mandibular | 29 | 35 | 20 | 24 |
| Muscle, biceps femoris | 1 | 2 | 1 | 2 |
| Muscle, diaphragm | 2 | 1 | 2 | 1 |
| Pancreas | 17 | 5 | 3 | 6 |
| Eye | 14 | 19 | 22 | 19 |
| Optic nerve | 18 | 20 | 14 | 21 |
| DRG cervical | 9 | 10 | 6 | 8 |
| DRG thoracic | 12 | 12 | 4 | 7 |
| DRG lumbar | 10 | 14 | 9 | 9 |
| DRG sacral | 11 | 18 | 10 | 12 |

Abbreviations: DRG, dorsal root ganglion; GFP, green fluorescent protein; ICM, intracisterna magna; LP, lumbar puncture.

###### Supplemental Table 26. Relative Median GFP Protein Concentrations of Animals at 4 Weeks Post-Dose.

|  | **GFP concentration (pg/mg of protein)** | |  | **GFP concentration (pg/mg of protein)** | |  |
| --- | --- | --- | --- | --- | --- | --- |
|  | **Intrathecal-LP (1.0×10^13^ vg/animal)** | **Intrathecal-LP (3.0×10^13^ vg/animal)** | **Ratio** | **ICM**  **(1.0×10^13^ vg/animal)** | **ICM**  **(3.0×10^13^ vg/animal)** | **Ratio** |
| Prefrontal cortex | 57.44 | 137.11 | 2.39 | 33.19 | 110.28 | 3.32 |
| Temporal cortex-entorhinal cortex | 82.56 | 255.51 | 3.09 | 36.68 | 117.23 | 3.20 |
| Caudate nucleus | 0.00 | 58.30 |  | 18.53 | 54.74 |  |
| Putamen (basal ganglia) | 40.57 | 111.06 | 2.74 | 23.52 | 66.04 | 2.81 |
| Cingulate gyrus | 105.20 | 158.96 | 1.51 | 43.04 | 138.42 |  |
| Corpus callosum | 50.22 | 149.41 |  | 37.28 | 111.88 |  |
| Temporal cortex-auditory cortex | 48.30 | 271.19 | 5.62 | 44.32 | 116.23 | 2.62 |
| Parietal cortex | 154.10 | 311.78 | 2.02 | 57.04 | 213.44 | 3.74 |
| Thalamus | 45.84 | 172.50 | 3.76 | 35.78 | 101.70 |  |
| Hypothalamus | 140.26 | 249.22 | 1.78 | 61.39 | 122.90 |  |
| Hippocampus | 80.40 | 320.55 | 3.99 | 36.63 | 138.44 | 3.78 |
| Amygdala | 218.78 | 991.28 | 4.53 | 138.37 | 412.46 | 2.98 |
| Substantia nigra | 73.15 | 402.75 |  | 77.70 | 239.45 | 3.08 |
| Pons | 339.24 | 796.80 | 2.35 | 175.90 | 740.71 | 4.21 |
| Cerebellum | 156.51 | 712.40 | 4.55 | 74.72 | 483.81 | 6.48 |
| Occipital cortex: primary visual cortex | 213.74 | 288.83 | 1.35 | 102.63 | 237.48 | 2.31 |
| Deep cerebellar nuclei | 266.43 | 967.68 | 3.63 | 197.81 | 662.71 | 3.35 |
| Heart | 4520.06 | 6665.81 | 1.47 | 946.08 | 3329.16 | 3.52 |
| Spinal cord (cervical) | 959.41 | 1515.04 | 1.58 | 2111.12 | 5775.95 | 2.74 |
| Spinal cord (thoracic) | 2711.49 | 1642.98 | 0.61 | 1655.40 | 1178.26 | 0.71 |
| Spinal cord (lumbar) | 2545.40 | 1249.84 | 0.49 | 975.68 | 1065.01 | 1.09 |
| Spinal cord (sacral) | 1886.39 | 1128.71 | 0.60 | 548.70 | 1335.85 | 2.43 |
| Trigeminal ganglia/nerve | 280.67 | 616.67 | 2.20 | 502.92 | 868.82 | 1.73 |
| Liver | 1625.08 | 4149.96 | 2.55 | 2542.28 | 5589.01 | 2.20 |
| Lung | 14.63 | 198.52 | 13.56 | 87.19 | 200.30 | 2.30 |
| Kidney  (right pole) | 113.33 | 212.94 | 1.88 | 82.50 | 986.52 | 11.96 |
| Spleen | 36.98 | 85.53 | 2.31 | 194.92 | 69.64 | 0.36 |
| Mandibular | 70.27 | 92.30 | 1.31 | 120.65 | 221.19 | 1.83 |
| Muscle, biceps femoris | 12951.30 | 7450.60 | 0.58 | 12758.10 | 9081.03 | 0.71 |
| Muscle, diaphragm | 10579.10 | 10224.39 | 0.97 | 10171.29 | 9191.35 | 0.90 |
| Pancreas | 244.14 | 1647.56 | 6.75 | 5840.27 | 2941.06 | 0.50 |
| Eye | 316.94 | 453.11 | 1.43 | 93.62 | 418.34 | 4.47 |
| Optic nerve | 231.24 | 433.79 | 1.88 | 531.98 | 381.82 | 0.72 |
| DRG cervical | 544.25 | 1111.67 | 2.04 | 2438.46 | 1631.84 | 0.67 |
| DRG thoracic | 432.44 | 977.92 | 2.26 | 3186.24 | 1832.38 | 0.58 |
| DRG lumbar | 494.07 | 809.85 | 1.64 | 1071.25 | 1393.82 | 1.30 |
| DRG sacral | 492.84 | 580.53 | 1.18 | 994.12 | 1069.44 | 1.08 |
|  |  | **Median** | 2.03 |  | **Median** | 2.37 |
|  |  | **Min** | 0.49 |  | **Min** | 0.36 |
|  |  | **Max** | 13.56 |  | **Max** | 11.96 |

Abbreviations: DRG, dorsal root ganglion; GFP, green fluorescent protein; ICM, intracisterna magna; LP, lumbar puncture; pg, picogram; vg, vector genomes.

**Supplemental Table 27. Individual Animal Data for scAAV9-CB-GFP Protein Expression Quantitation in Intrathecal-LP Control Animals (Group 1).**

|  | **GFP pg/mg of protein** | | | |
| --- | --- | --- | --- | --- |
| **Group 1 (intrathecal-LP, vehicle)** | **001** | **002** | **003** | **004** |
| Brain - Prefrontal cortex | BLOQ | BLOQ | BLOQ | BLOQ |
| Brain - temporal cortex | BLOQ | BLOQ | BLOQ | BLOQ |
| Brain - caudate nucleus | BLOQ | BLOQ | BLOQ | BLOQ |
| Brain - putamen | BLOQ | BLOQ | BLOQ | BLOQ |
| Brain - cingulate gyrus | BLOQ | BLOQ | BLOQ | BLOQ |
| Brain - corpus callosum | BLOQ | BLOQ | BLOQ | BLOQ |
| Brain - temporal cortex auditory | BLOQ | BLOQ | BLOQ | BLOQ |
| Brain - parietal cortex | BLOQ | BLOQ | BLOQ | BLOQ |
| Brain - thalamus | BLOQ | BLOQ | BLOQ | BLOQ |
| Brain - hypothalamus | BLOQ | BLOQ | BLOQ | BLOQ |
| Brain - hippocampus | BLOQ | BLOQ | BLOQ | BLOQ |
| Brain - amygdala | BLOQ | BLOQ | BLOQ | BLOQ |
| Brain - substantia nigra | BLOQ | BLOQ | BLOQ | BLOQ |
| Brain - pons | BLOQ | BLOQ | BLOQ | BLOQ |
| Brain - cerebellum | BLOQ | BLOQ | BLOQ | BLOQ |
| Brain - occipital cortex primary visual | BLOQ | BLOQ | BLOQ | BLOQ |
| Brain - deep cerebellar nuclei | BLOQ | BLOQ | BLOQ | BLOQ |
| Heart | BLOQ | BLOQ | BLOQ | BLOQ |
| Spinal cord (cervical) | BLOQ | BLOQ | BLOQ | BLOQ |
| Spinal cord (thoracic) | BLOQ | BLOQ | BLOQ | BLOQ |
| Spinal cord (lumbar) | BLOQ | BLOQ | BLOQ | BLOQ |
| Spinal cord (sacral) | BLOQ | BLOQ | BLOQ | BLOQ |
| Trigeminal ganglion nerve | BLOQ | BLOQ | BLOQ | BLOQ |
| Liver | BLOQ | BLOQ | BLOQ | BLOQ |
| Lung | BLOQ | BLOQ | BLOQ | BLOQ |
| Kidney | BLOQ | BLOQ | BLOQ | BLOQ |
| Spleen | BLOQ | BLOQ | BLOQ | BLOQ |
| Mand LN | BLOQ | BLOQ | BLOQ | BLOQ |
| Muscle, biceps femoris | BLOQ | BLOQ | BLOQ | BLOQ |
| Muscle, diaphragm | BLOQ | BLOQ | BLOQ | BLOQ |
| Pancreas | BLOQ | BLOQ | BLOQ | BLOQ |
| Eye | BLOQ | BLOQ | BLOQ | BLOQ |
| Optic nerve | BLOQ | BLOQ | BLOQ | BLOQ |
| DRG cervical | BLOQ | BLOQ | BLOQ | BLOQ |
| DRG thoracic | BLOQ | BLOQ | BLOQ | BLOQ |
| DRG lumbar | BLOQ | BLOQ | BLOQ | BLOQ |
| DRG sacral | BLOQ | BLOQ | BLOQ | BLOQ |

Abbreviations: BLOQ, below limit of quantification; DRG, dorsal root ganglion; GFP, green fluorescent protein; LP, lumbar puncture; Mand LN, mandibular lymph node; pg, picogram; scAAV9-CB-GFP, self-complementary adeno-associated virus serotype 9–chicken β-actin promoter–green fluorescent protein.

**Supplemental Table 28. Individual Animal Data for scAAV9-CB-GFP Protein Expression Quantitation in ICM Control Animals (Group 2).**

|  | **GFP pg/mg of protein** | | | |
| --- | --- | --- | --- | --- |
| **Group 2 (ICM, vehicle)** | **101** | **102** | **103** | **104** |
| Brain - Prefrontal cortex | BLOQ | BLOQ | BLOQ | BLOQ |
| Brain - temporal cortex | BLOQ | BLOQ | BLOQ | BLOQ |
| Brain - caudate nucleus | BLOQ | BLOQ | BLOQ | BLOQ |
| Brain - putamen | BLOQ | BLOQ | BLOQ | BLOQ |
| Brain - cingulate gyrus | BLOQ | BLOQ | BLOQ | BLOQ |
| Brain - corpus callosum | BLOQ | BLOQ | BLOQ | BLOQ |
| Brain - temporal cortex auditory | BLOQ | BLOQ | BLOQ | BLOQ |
| Brain - parietal cortex | BLOQ | BLOQ | BLOQ | BLOQ |
| Brain - thalamus | BLOQ | BLOQ | BLOQ | BLOQ |
| Brain - hypothalamus | BLOQ | BLOQ | BLOQ | BLOQ |
| Brain - hippocampus | BLOQ | BLOQ | BLOQ | BLOQ |
| Brain - amygdala | BLOQ | BLOQ | BLOQ | BLOQ |
| Brain - substantia nigra | BLOQ | BLOQ | BLOQ | BLOQ |
| Brain - pons | BLOQ | BLOQ | BLOQ | BLOQ |
| Brain - cerebellum | BLOQ | BLOQ | BLOQ | BLOQ |
| Brain - occipital cortex primary visual | BLOQ | BLOQ | BLOQ | BLOQ |
| Brain - deep cerebellar nuclei | BLOQ | BLOQ | BLOQ | BLOQ |
| Heart | BLOQ | BLOQ | BLOQ | BLOQ |
| Spinal cord (cervical) | BLOQ | BLOQ | BLOQ | BLOQ |
| Spinal cord (thoracic) | BLOQ | BLOQ | BLOQ | BLOQ |
| Spinal cord (lumbar) | BLOQ | BLOQ | BLOQ | BLOQ |
| Spinal cord (sacral) | BLOQ | BLOQ | BLOQ | BLOQ |
| Trigeminal ganglion nerve | BLOQ | BLOQ | BLOQ | BLOQ |
| Liver | BLOQ | BLOQ | BLOQ | BLOQ |
| Lung | BLOQ | BLOQ | BLOQ | BLOQ |
| Kidney | BLOQ | BLOQ | BLOQ | BLOQ |
| Spleen | BLOQ | BLOQ | BLOQ | BLOQ |
| Mand LN | BLOQ | BLOQ | BLOQ | BLOQ |
| Muscle, biceps femoris | BLOQ | BLOQ | BLOQ | BLOQ |
| Muscle, diaphragm | BLOQ | BLOQ | BLOQ | BLOQ |
| Pancreas | BLOQ | BLOQ | BLOQ | BLOQ |
| Eye | BLOQ | BLOQ | BLOQ | BLOQ |
| Optic nerve | BLOQ | BLOQ | BLOQ | BLOQ |
| DRG cervical | BLOQ | BLOQ | BLOQ | BLOQ |
| DRG thoracic | BLOQ | BLOQ | BLOQ | BLOQ |
| DRG lumbar | BLOQ | BLOQ | BLOQ | BLOQ |
| DRG sacral | BLOQ | BLOQ | BLOQ | BLOQ |

Abbreviations: BLOQ, below limit of quantification; DRG, dorsal root ganglion; GFP, green fluorescent protein; ICM, intracisterna magna; Mand LN, mandibular lymph node; pg, picogram; scAAV9-CB-GFP, self-complementary adeno-associated virus serotype 9–chicken β-actin promoter–green fluorescent protein.

**Supplemental Table 29. Individual Animal Data for scAAV9-CB-GFP Protein Expression Quantitation in Intrathecal-LP-treated Animals at 1.0×10^13^ (Group 3).**

|  | **GFP pg/mg of protein** | | | |
| --- | --- | --- | --- | --- |
| **Group 3 (intrathecal-LP, dosed 1.0×10^13^ vg/animal)** | **201** | **202** | **203** | **204** |
| Brain - Prefrontal cortex | 34.10391 | 80.77747 | 92.19108 | 27.16518 |
| Brain - temporal cortex | 103.7348 | 112.2996 | 61.38952 | BLOQ |
| Brain - caudate nucleus | BLOQ | BLOQ | 28.46743 | BLOQ |
| Brain - putamen | 35.32458 | 45.81729 | 69.13442 | 29.48446 |
| Brain - cingulate gyrus | 126.2143 | 89.13426 | 121.265 | 38.08863 |
| Brain - corpus callosum | BLOQ | 107.4249 | 100.4433 | BLOQ |
| Brain - temporal cortex auditory | 167.0905 | 46.44482 | 50.14828 | 42.21936 |
| Brain - parietal cortex | 184.8402 | 167.2739 | 140.922 | 120.2893 |
| Brain - thalamus | BLOQ | 59.00122 | 103.0218 | 32.67705 |
| Brain - hypothalamus | 139.4509 | 194.4018 | 141.074 | 56.01159 |
| Brain - hippocampus | 82.08944 | 754.6823 | 78.70869 | BLOQ |
| Brain - amygdala | 257.095 | 459.5696 | 180.4552 | 84.90486 |
| Brain - substantia nigra | 42.15537 | 443.0814 | 104.1389 | 38.32124 |
| Brain - pons | 104.5318 | 450.0675 | 648.7015 | 228.4088 |
| Brain - cerebellum | 405.78 | 153.4847 | 159.5259 | 35.71874 |
| Brain - occipital cortex primary visual | 266.2106 | 391.5579 | 161.2739 | 45.93865 |
| Brain - deep cerebellar nuclei | 235.3546 | 1785.445 | 297.5108 | 33.23526 |
| Heart | 213.9633 | 12934.27 | 8826.166 | 155.2807 |
| Spinal cord (cervical) | 787.1841 | 5057.007 | 1131.631 | 187.7895 |
| Spinal cord (thoracic) | 1722.895 | 4081.623 | 3700.093 | 316.9124 |
| Spinal cord (lumbar) | 514.6541 | 6127.127 | 4576.148 | 151.3098 |
| Spinal cord (sacral) | 312.928 | 3459.857 | 7627.827 | 83.59389 |
| Trigeminal ganglion nerve | 109.8124 | 451.5283 | 609.3379 | 36.10414 |
| Liver | 88.63656 | 3161.522 | 3389.463 | 88.47709 |
| Lung | BLOQ | 29.2698 | 243.4939 | BLOQ |
| Kidney | 103.4023 | 123.2596 | 312.2917 | 40.35675 |
| Spleen | BLOQ | 73.95814 | 775.2389 | BLOQ |
| Mand LN | BLOQ | 140.5498 | 560.142 | BLOQ |
| Muscle, biceps femoris | 21104.95 | 14923.09 | 10979.51 | 6402.154 |
| Muscle, diaphragm | 177.9983 | 11369.03 | 9789.169 | 14712.1 |
| Pancreas | 256.035 | 232.2516 | 9790.179 | 33.11463 |
| Eye | 32.24188 | 1197.484 | 601.6479 | BLOQ |
| Optic nerve | 303.0714 | 204.5555 | 257.9313 | BLOQ |
| DRG cervical | 366.2738 | 722.2174 | 1638.989 | 51.84749 |
| DRG thoracic | 309.8069 | 555.0791 | 3084.286 | 66.40054 |
| DRG lumbar | 367.9383 | 620.2035 | 3802.832 | 128.4333 |
| DRG sacral | 206.6596 | 779.0231 | 5823.028 | 33.87306 |

Abbreviations: BLOQ, below limit of quantification; DRG, dorsal root ganglion; GFP, green fluorescent protein; LP, lumbar puncture; Mand LN, mandibular lymph node; pg, picogram; scAAV9-CB-GFP, self-complementary adeno-associated virus serotype 9–chicken β-actin promoter–green fluorescent protein, vg, vector genomes.

**Supplemental Table 30. Individual Animal Data for scAAV9-CB-GFP Protein Expression Quantitation in Intrathecal-LP-treated Animals at 3.0×10^13^ (Group 4).**

|  | **GFP pg/mg of protein** | | | |
| --- | --- | --- | --- | --- |
| **Group 4  (Intrathecal-LP, dosed 3.0×10^13^ vg/animal)** | **P0301** | **P0302** | **P0303** | **P0304** |
| Brain - Prefrontal cortex | 249.6474 | 156.4518 | 91.01781 | 117.7776 |
| Brain - temporal cortex | 218.2424 | 385.1818 | 100.374 | 292.7767 |
| Brain - caudate nucleus | 60.67858 | 55.92727 | 44.85486 | 116.7185 |
| Brain - putamen | 119.0354 | 134.7647 | 62.03007 | 103.0844 |
| Brain - cingulate gyrus | 247.1274 | 195.0413 | 122.87 | 108.181 |
| Brain - corpus callosum | 156.0165 | 183.9236 | 87.27756 | 142.7975 |
| Brain - temporal cortex auditory | 440.6551 | 241.8119 | 300.5776 | 161.5066 |
| Brain - parietal cortex | 429.3507 | 412.3631 | 88.87021 | 211.1901 |
| Brain - thalamus | 240.3188 | 145.2624 | 167.3553 | 177.6529 |
| Brain - hypothalamus | 227.5621 | 549.4298 | 266.6284 | 231.8081 |
| Brain - hippocampus | 308.0505 | 505.6494 | 214.5254 | 333.0522 |
| Brain - amygdala | 1168.794 | 1336.87 | 813.7749 | 372.7753 |
| Brain - substantia nigra | 465.3513 | 208.5313 | 1668.589 | 340.1511 |
| Brain - pons | 580.1389 | 929.4578 | 664.1327 | 990.399 |
| Brain - cerebellum | 697.246 | 868.1457 | 376.3925 | 727.5484 |
| Brain - occipital cortex primary visual | 358.5808 | 1473.396 | 127.6143 | 219.0819 |
| Brain - deep cerebellar nuclei | 454.0262 | 1481.33 | 235.4066 | 1495.636 |
| Heart | 8671.226 | 4660.387 | 18169.13 | 1751.934 |
| Spinal cord (cervical) | 1227.87 | 2087.251 | 765.8204 | 1802.213 |
| Spinal cord (thoracic) | 1590.661 | 2132.642 | 1582.931 | 1695.304 |
| Spinal cord (lumbar) | 776.2477 | 1132.374 | 1367.308 | 1430.901 |
| Spinal cord (sacral) | 1055.567 | 11844.1 | 1201.858 | 264.1928 |
| Trigeminal ganglion nerve | 1244.301 | 959.1894 | 274.1432 | 127.7435 |
| Liver | 2627.925 | 5671.988 | 7691.112 | 245.1229 |
| Lung | 140.3618 | 360.8256 | 256.6826 | 94.8879 |
| Kidney | 297.5697 | 364.5201 | 128.3196 | 79.76043 |
| Spleen | 112.2508 | 108.4768 | 62.59241 | 44.11288 |
| Mand LN | 90.27852 | 235.9479 | 94.32916 | 76.30599 |
| Muscle, biceps femoris | 6325.057 | 16407.99 | 8356.033 | 6545.161 |
| Muscle, diaphragm | 6098.651 | 11478.12 | 16837.1 | 8970.657 |
| Pancreas | 1238.626 | 2056.501 | 8693.135 | 107.781 |
| Eye | 659.914 | 662.925 | 246.3056 | 106.6425 |
| Optic nerve | 672.5795 | 1042.485 | 95.5225 | 195.0047 |
| DRG cervical | 1278.293 | 1565.34 | 916.8556 | 945.0382 |
| DRG thoracic | 748.8163 | 1207.025 | 1363.251 | 745.0768 |
| DRG lumbar | 978.3012 | 1190.04 | 33.0501 | 641.4041 |
| DRG sacral | 1547.732 | 464.2252 | 696.8324 | 145.7268 |

Abbreviations: DRG, dorsal root ganglion; GFP, green fluorescent protein; LP, lumbar puncture; Mand LN, mandibular lymph node; pg, picogram; scAAV9-CB-GFP, self-complementary adeno-associated virus serotype 9–chicken β-actin promoter–green fluorescent protein; vg, vector genomes.

**Supplemental Table 31. Individual Animal Data for scAAV9-CB-GFP Protein Expression Quantitation in ICM-treated Animals at 1.0×10^13^ (Group 5).**

|  | **GFP pg/mg of protein** | | | |
| --- | --- | --- | --- | --- |
| **Group 5  (ICM, dosed 1.0×1013 vg/animal)** | **P0401** | **P0402** | **P0403** | **P0404** |
| Brain - Prefrontal cortex | 125.182 | 32.30861 | 34.07724 | 28.7396 |
| Brain - temporal cortex | 539.0901 | 41.68146 | 31.67192 | 25.42107 |
| Brain - caudate nucleus | 175.5299 | 10.59315 | 20.77727 | 16.28613 |
| Brain - putamen | 24.08631 | 23.90539 | 23.14422 | 21.02119 |
| Brain - cingulate gyrus | 325.1637 | 41.92018 | 44.15854 | 27.29357 |
| Brain - corpus callosum | 2288.405 | 37.74139 | 36.81228 | 15.33209 |
| Brain - temporal cortex auditory | 395.9644 | 26.54655 | 32.86316 | 55.7771 |
| Brain - parietal cortex | 1491.964 | 51.98357 | 62.0883 | 33.61278 |
| Brain - thalamus | 39.655 | 30.09346 | 44.20518 | 31.91072 |
| Brain - hypothalamus | 205.338 | 33.40136 | 53.90442 | 68.87222 |
| Brain - hippocampus | 51.62271 | 26.52374 | 38.33403 | 34.92517 |
| Brain - amygdala | 2413.146 | 25.538 | 228.0017 | 48.73901 |
| Brain - substantia nigra | 198.201 | 48.40878 | 71.37282 | 84.03192 |
| Brain - pons | 405.0002 | 131.8316 | 219.973 | 108.6024 |
| Brain - cerebellum | 290.8877 | 31.00205 | 100.4626 | 48.9712 |
| Brain - occipital cortex primary visual | 388.2505 | 44.21926 | 70.01535 | 135.2376 |
| Brain - deep cerebellar nuclei | 4458.4 | 55.68012 | 253.0896 | 142.5363 |
| Heart | 878.2521 | 1013.914 | 2501.72 | 689.6544 |
| Spinal cord (cervical) | 5605.879 | 410.5586 | 2263.804 | 1958.428 |
| Spinal cord (thoracic) | 5020.314 | 661.1627 | 2649.63 | 491.1306 |
| Spinal cord (lumbar) | 2897.526 | 339.9407 | 1502.3 | 449.0592 |
| Spinal cord (sacral) | 8872.874 | 596.8566 | 500.5506 | 185.2536 |
| Trigeminal ganglion nerve | 2502.308 | 429.2704 | 576.5663 | 139.2916 |
| Liver | 5694.506 | 879.6044 | 4204.957 | 91.35812 |
| Lung | 117.9262 | 101.0195 | 73.3523 | 32.05598 |
| Kidney | 104.7573 | 60.24009 | 122.9203 | 38.65379 |
| Spleen | 820.3863 | 109.9822 | 279.8545 | 24.32062 |
| Mand LN | 439.6452 | 82.62256 | 158.681 | 43.03878 |
| Muscle, biceps femoris | 363.697 | 11944.14 | 19349.98 | 13572.06 |
| Muscle, diaphragm | 1656.532 | 9391.396 | 10951.19 | 13573.96 |
| Pancreas | 3430.706 | 6057.296 | 5840.269 | BLOQ |
| Eye | 19.56947 | 2371.79 | 150.0655 | 37.17385 |
| Optic nerve | 1282.218 | 280.0623 | 783.8919 | 58.32481 |
| DRG cervical | 16688.75 | 539.2022 | 2379.91 | 2497.003 |
| DRG thoracic | 3867.563 | 2504.917 | 5496.269 | 481.2349 |
| DRG lumbar | 1633.643 | 508.8558 | 2623.122 | 350.0355 |
| DRG sacral | 2470.712 | 414.8189 | 1573.418 | 87.65066 |

Abbreviations: BLOQ, below limit of quantification; DRG, dorsal root ganglion; GFP, green fluorescent protein; ICM, intracisterna magna; Mand LN, mandibular lymph node; pg, picogram; scAAV9-CB-GFP, self-complementary adeno-associated virus serotype 9–chicken β-actin promoter–green fluorescent protein; vg, vector genomes.

**Supplemental Table 32. Individual Animal Data for scAAV9-CB-GFP Protein Expression Quantitation in ICM-treated Animals at 3.0×10^13^ (Group 6).**

|  | **GFP pg/mg of protein** | | | |
| --- | --- | --- | --- | --- |
| **Group 6  (ICM, dosed 3×10^13^** **vg/animal)** | **P0501** | **P0502** | **P0503** | **P0504** |
| Brain - Prefrontal cortex | 69.64744 | 50.24038 | 150.9193 | 535.041 |
| Brain - temporal cortex | 63.92072 | 37.01057 | 170.5462 | 697.0968 |
| Brain - caudate nucleus | 42.27121 | 18.75569 | 67.20687 | 186.1674 |
| Brain - putamen | 44.02028 | 37.36085 | 88.06455 | 206.6349 |
| Brain - cingulate gyrus | 107.3865 | 62.90226 | 169.4634 | 334.3277 |
| Brain - corpus callosum | 71.99289 | 79.62708 | 144.125 | 313.9777 |
| Brain - temporal cortex auditory | 130.0048 | 83.73358 | 102.4588 | 499.6909 |
| Brain - parietal cortex | 202.374 | 72.16461 | 224.5107 | 457.451 |
| Brain - thalamus | 101.1713 | 67.71868 | 102.2292 | 388.4067 |
| Brain - hypothalamus | 71.09878 | 50.95496 | 174.7044 | 483.3559 |
| Brain - hippocampus | 126.6863 | 110.0587 | 150.2002 | 3116.401 |
| Brain - amygdala | 234.663 | 147.1321 | 590.2639 | 2217.692 |
| Brain - substantia nigra | 250.2834 | 122.9739 | 228.6136 | 604.6436 |
| Brain - pons | 239.9039 | 359.8069 | 1531.71 | 1121.615 |
| Brain - cerebellum | 777.3063 | 64.99272 | 190.3053 | 1181.849 |
| Brain - occipital cortex primary visual | 104.9603 | 213.2176 | 261.7385 | 2686.843 |
| Brain - deep cerebellar nuclei | 868.6648 | 205.0526 | 456.7534 | 1058.759 |
| Heart | 920.8833 | 2421.625 | 4236.703 | 6953.074 |
| Spinal cord (cervical) | 2580.533 | 875.6663 | 10011.78 | 8971.361 |
| Spinal cord (thoracic) | 964.978 | 615.5592 | 13533.14 | 1391.547 |
| Spinal cord (lumbar) | 685.6804 | 263.3706 | 12950.41 | 1444.341 |
| Spinal cord (sacral) | 1189.636 | 311.7544 | 17547.83 | 1482.066 |
| Trigeminal ganglion nerve | 1186.307 | 195.5754 | 551.3425 | 1352.537 |
| Liver | 1062.746 | 1952.536 | 18792.8 | 9225.493 |
| Lung | 156.1719 | 67.52287 | 244.422 | 804.2416 |
| Kidney | 214.6229 | 5936.131 | 249.2802 | 1723.765 |
| Spleen | 51.92289 | 18.68848 | 87.35749 | 3200.613 |
| Mand LN | 60.78255 | 70.68404 | 371.6893 | 2595.337 |
| Muscle, biceps femoris | 24544.13 | 4633.596 | 11616.39 | 6545.667 |
| Muscle, diaphragm | 6938.214 | 4242.272 | 22233.7 | 11444.49 |
| Pancreas | 1892.574 | 712.5947 | 3989.536 | 8466.732 |
| Eye | 54.58269 | 65.41853 | 771.2613 | 1683.379 |
| Optic nerve | 569.838 | 193.8 | 178.5595 | 799.3638 |
| DRG cervical | 2193.6 | 544.3082 | 32693.17 | 1070.082 |
| DRG thoracic | 1227.172 | 311.8986 | 11506.1 | 2437.592 |
| DRG lumbar | 844.008 | 308.9828 | 8518.048 | 1943.629 |
| DRG sacral | 784.1121 | 191.1862 | 19236.71 | 1354.771 |

Abbreviations: DRG, dorsal root ganglion; GFP, green fluorescent protein; ICM, intracisterna magna; Mand LN, mandibular lymph node; pg, picogram; scAAV9-CB-GFP, self-complementary adeno-associated virus serotype 9–chicken β-actin promoter–green fluorescent protein; vg, vector genomes.

**Supplemental Table 33. Macroscopic and Microscopic Observations from Individual Animals.**

| **Animal Number: P0001 Group/Subgroup: 1/1 Sex: F Fate Status: Terminal Sacrifice**  **Date of Fate: 12 Oct 20 Phase of Fate: Dosing Phase Wk/Day of Fate: 4/28 TBW(g): 1600.0** | | | | |
| --- | --- | --- | --- | --- |
| Organ Name: None | | | | |
| Macroscopic Observation(s)  Colon: Discolored; mucosa; multiple, indistinct; red; collected  Spleen: Raised area; mid region; single, up to 5 mm3; white; collected | | | | Microscopic Observation(s)  Brain: Infiltrate, mononuclear cell; minimal; perivascular, neuropil/choroid plexus/neuropil: slide 45, 47, 49 neuropil: slide 44  Colon: Congestion, agonal; minimal  Ganglion, Dorsal Root, Sacral: Autophagy; minimal  Ganglion, Dorsal Root, Sacral: Infiltrate, mononuclear cell; minimal  Intrathecal Injection Site, Cisterna Magna: MISSING  Intrathecal Injection Site, Cisterna Magna: TISSUE  COMMENT: Not available per histo comment  Kidney: Infiltrate, mononuclear cell; slight  Liver: Infiltrate, mononuclear cell; minimal  Ovary, Left: Immature; Present  Spinal Cord, Cervical: Gliosis, white matter; minimal; focal; dorsal funiculus white matter  Spinal Cord, Thoracic: Degeneration, axon, funiculus; minimal; focal; ventral funiculus white matter  Thyroid: Ectopic tissue, lymph node; Present  Thyroid: Ectopic tissue, thymus; Present |
| The following tissues were examined macroscopically and were unremarkable: Adrenal; Animal; Aorta; Brain; Cecum; Cervix; Duodenum; Esophagus; Eye, Left; Femur; GALT/Peyer's Patch; Gall Bladder; Ganglion, Cervical Dorsal Root; Ganglion, Dorsal Root, Sacral; Ganglion, Lumbar Dorsal Root; Ganglion, Superior Cervical; Ganglion, Thoracic Dorsal Root; Heart; Ileum; Intrathecal Injection Site, Cisterna Magna; Intrathecal Injection Site, Sacral Spinal Cord; Jejunum; Kidney; Liver; Lung; Lymph Node, Mandibular; Lymph Node, Mesenteric; Mammary Gland; Mandibular Salivary Gland; Marrow, Femur; Marrow, Sternum; Muscle, Biceps Femoris; Nerve, Optic, Left; Nerve, Radial; Nerve, Sciatic; Nerve, Sural; Nerve, Tibial; Nerve, Ulnar; Ovary, Left; Pancreas; Parathyroid; Pituitary; Rectum; Skin/Subcutis; Spinal Cord; Spinal Nerve Roots, Cauda Equina; Sternum; Stomach; Thymus; Thyroid; Tongue; Trachea; Urinary Bladder; Uterus; Vagina | | | | |
| The following tissues were examined microscopically and were unremarkable: Adrenal, Cortex; Adrenal, Medulla; Aorta; Cecum; Duodenum; Eye, Left; Femur; GALT/Peyer's Patch; Ganglion, Cervical Dorsal Root; Ganglion, Lumbar Dorsal Root; Ganglion, Superior Cervical; Ganglion, Thoracic Dorsal Root; Heart; Intrathecal Injection Site, Sacral Spinal Cord; Lung; Lymph Node, Mandibular; Marrow, Femur; Marrow, Sternum; Muscle, Biceps Femoris; Nerve, Radial; Nerve, Sciatic; Nerve, Sural; Nerve, Tibial; Nerve, Ulnar; Pancreas; Parathyroid; Spinal Cord, Lumbar; Spinal Nerve Roots, Cauda Equina; Spleen; Sternum; Stomach; Thymus | | | | |
| ‑‑‑‑‑‑‑‑‑‑‑‑‑‑‑‑‑‑‑‑‑‑‑‑‑‑‑‑‑‑‑‑‑‑‑‑‑‑‑‑‑‑‑‑‑‑‑‑‑‑‑‑‑‑‑‑‑‑‑‑‑‑‑‑‑‑‑‑‑‑‑‑‑‑‑‑‑‑‑‑‑‑‑‑ | | | | |
| **Animal Number: P0002 Group/Subgroup: 1/1 Sex: F Fate Status: Terminal Sacrifice**  **Date of Fate: 12 Oct 20 Phase of Fate: Dosing Phase Wk/Day of Fate: 4/28 TBW(g): 1400.0** | | | | |
| Organ Name: None | | | | |
| Macroscopic Observation(s)  None | Microscopic Observation(s)  Cecum: Parasite, protozoa, increased; slight  Ganglion, Dorsal Root, Sacral: Infiltrate, mononuclear cell; minimal / slide 41  Ganglion, Lumbar Dorsal Root: Infiltrate, mononuclear cell; minimal  Ganglion, Superior Cervical: Vacuolation, neuron; minimal  Intrathecal Injection Site, Cisterna Magna: MISSING  Intrathecal Injection Site, Cisterna Magna: TISSUE  COMMENT: Not available per histo comment  Lung: Infiltrate, mononuclear cell; minimal; bronchioles; terminal  Lymph Node, Mandibular: Lymphocytes, increased; slight; follicle, bilateral  Nerve, Ulnar: Infiltrate, mononuclear cell; minimal; focal  Ovary, Left: Immature; Present | | | |
| The following tissues were examined macroscopically and were unremarkable:  Adrenal; Animal; Aorta; Brain; Cecum; Cervix; Colon; Duodenum; Esophagus; Eye, Left; Femur; GALT/Peyer's Patch; Gall Bladder; Ganglion, Cervical Dorsal Root; Ganglion, Dorsal Root, Sacral; Ganglion, Lumbar Dorsal Root; Ganglion, Superior Cervical; Ganglion, Thoracic Dorsal Root; Heart; Ileum; Intrathecal Injection Site, Cisterna Magna; Intrathecal Injection Site, Sacral Spinal Cord;  Jejunum; Kidney; Liver; Lung; Lymph Node, Mandibular; Lymph Node, Mesenteric; Mammary Gland;  Mandibular Salivary Gland; Marrow, Femur; Marrow, Sternum; Muscle, Biceps Femoris; Nerve, Optic,  Left; Nerve, Radial; Nerve, Sciatic; Nerve, Sural; Nerve, Tibial; Nerve, Ulnar; Ovary, Left;  Pancreas; Parathyroid; Pituitary; Rectum; Skin/Subcutis; Spinal Cord; Spinal Nerve Roots, Cauda Equina; Spleen; Sternum; Stomach; Thymus; Thyroid; Tongue; Trachea; Urinary Bladder; Uterus; Vagina | | | | |
| The following tissues were examined microscopically and were unremarkable:  Adrenal, Cortex; Adrenal, Medulla; Aorta; Brain; Duodenum; Eye, Left; Femur; GALT/Peyer's Patch; Ganglion, Cervical Dorsal Root; Ganglion, Thoracic Dorsal Root; Heart; Intrathecal Injection Site, Sacral Spinal Cord; Kidney; Liver; Marrow, Femur; Marrow, Sternum; Muscle, Biceps Femoris; Nerve, Radial; Nerve, Sciatic; Nerve, Sural; Nerve, Tibial; Pancreas; Parathyroid; Spinal Cord, Cervical; Spinal Cord, Lumbar; Spinal Cord, Thoracic; Spinal Nerve Roots, Cauda Equina; Spleen; Sternum; Stomach; Thymus; Thyroid | | | | |
| ‑‑‑‑‑‑‑‑‑‑‑‑‑‑‑‑‑‑‑‑‑‑‑‑‑‑‑‑‑‑‑‑‑‑‑‑‑‑‑‑‑‑‑‑‑‑‑‑‑‑‑‑‑‑‑‑‑‑‑‑‑‑‑‑‑‑‑‑‑‑‑‑‑‑‑‑‑‑‑‑‑‑‑‑ | | | | |
| **Animal Number: P0003 Group/Subgroup: 1/2 Sex: F Fate Status: Terminal Sacrifice**  **Date of Fate: 15 Oct 20 Phase of Fate: Dosing Phase Wk/Day of Fate: 4/28 TBW(g): 1600.0** | | | | |
| Organ Name: None | | | | |
| Macroscopic Observation(s  None | | | | Microscopic Observation(s)  Ganglion, Lumbar Dorsal Root: Autophagy; minimal  Intrathecal Injection Site, Cisterna Magna: Degeneration; minimal; dorsal, lateral  Lung: Infiltrate, macrophages, alveolus; minimal  Ovary, Left: Immature; Present  Spinal Cord, Cervical: Degeneration, axon, funiculus; minimal; focal; dorsal funiculus white matter  Spinal Cord, Thoracic: Degeneration, axon, funiculus; minimal; focal; lateral funiculus white matter  Spleen: Macrophages, increased, red pulp; moderate  Thyroid: Ectopic tissue, thymus; Present |
| The following tissues were examined macroscopically and were unremarkable: Adrenal; Animal; Aorta; Brain; Cecum; Cervix; Colon; Duodenum; Esophagus; Eye, Left; Femur; GALT/Peyer's Patch; Gall Bladder; Ganglion, Cervical Dorsal Root; Ganglion, Dorsal Root, Sacral; Ganglion, Lumbar Dorsal Root; Ganglion, Superior Cervical; Ganglion, Thoracic Dorsal Root; Heart; Ileum; Intrathecal Injection Site, Cisterna Magna; Intrathecal Injection Site, Sacral Spinal Cord; Jejunum; Kidney; Liver; Lung; Lymph Node, Mandibular; Lymph Node, Mesenteric; Mammary Gland; Mandibular Salivary Gland; Marrow, Femur; Marrow, Sternum; Muscle, Biceps Femoris; Nerve, Optic, Left; Nerve, Radial; Nerve, Sciatic; Nerve, Sural; Nerve, Tibial; Nerve, Ulnar; Ovary, Left; Pancreas; Parathyroid; Pituitary; Rectum; Skin/Subcutis; Spinal Cord; Spinal Nerve Roots, Cauda Equina; Spleen; Sternum; Stomach; Thymus; Thyroid; Tongue; Trachea; Urinary Bladder; Uterus; Vagina | | | | |
| The following tissues were examined microscopically and were unremarkable: Adrenal, Cortex; Adrenal, Medulla; Aorta; Brain; Cecum; Duodenum; Eye, Left; Femur; GALT/Peyer's Patch; Ganglion, Cervical Dorsal Root; Ganglion, Dorsal Root, Sacral; Ganglion, Superior Cervical; Ganglion, Thoracic Dorsal Root; Heart; Intrathecal Injection Site, Sacral Spinal Cord; Kidney; Liver; Lymph Node, Mandibular; Marrow, Femur; Marrow, Sternum; Muscle, Biceps Femoris; Nerve, Radial; Nerve, Sciatic; Nerve, Sural; Nerve, Tibial; Nerve, Ulnar; Pancreas; Parathyroid; Spinal Cord, Lumbar; Spinal Nerve Roots, Cauda Equina; Sternum; Stomach; Thymus | | | | |
| ‑‑‑‑‑‑‑‑‑‑‑‑‑‑‑‑‑‑‑‑‑‑‑‑‑‑‑‑‑‑‑‑‑‑‑‑‑‑‑‑‑‑‑‑‑‑‑‑‑‑‑‑‑‑‑‑‑‑‑‑‑‑‑‑‑‑‑‑‑‑‑‑‑‑‑‑‑‑‑‑‑‑‑‑ | | | | |
| **Animal Number: P0004 Group/Subgroup: 1/2 Sex: F Fate Status: Terminal Sacrifice**  **Date of Fate: 15 Oct 20 Phase of Fate: Dosing Phase Wk/Day of Fate: 4/28 TBW(g): 1400.0** | | | | |
| Organ Name: None | | | | |
| Macroscopic Observation(s)  Colon: discolored; mucosa; multiple, up to 2mm2; red; collected | | | | Microscopic Observation(s)  Colon: Discolored; mucosa; multiple, up to 2 mm2; red; collected  Ganglion, Lumbar Dorsal Root: Vacuolation, neuron; minimal  Heart: Infiltrate, mononuclear cell; minimal  Intrathecal Injection Site, Cisterna Magna: MISSING  Intrathecal Injection Site, Cisterna Magna: TISSUE COMMENT: Not available per histo comment  Nerve, Tibial: Degeneration, axon; minimal  Ovary, Left: Immature; Present  Spinal Cord, Lumbar: Degeneration, axon, funiculus; minimal; focal; lateral funiculus white matter |
| The following tissues were examined macroscopically and were unremarkable: Adrenal; Animal; Aorta; Brain; Cecum; Cervix; Duodenum; Esophagus; Eye, Left; Femur; GALT/Peyer's Patch; Gall Bladder; Ganglion, Cervical Dorsal Root; Ganglion, Dorsal Root, Sacral; Ganglion, Lumbar Dorsal Root; Ganglion, Superior Cervical; Ganglion, Thoracic Dorsal Root; Heart; Ileum; Intrathecal Injection Site, Cisterna Magna; Intrathecal Injection Site, Sacral Spinal Cord; Jejunum; Kidney; Liver; Lung; Lymph Node, Mandibular; Lymph Node, Mesenteric; Mammary Gland; Mandibular Salivary Gland; Marrow, Femur; Marrow, Sternum; Muscle, Biceps Femoris; Nerve, Optic, Left; Nerve, Radial; Nerve, Sciatic; Nerve, Sural; Nerve, Tibial; Nerve, Ulnar; Ovary, Left; Pancreas; Parathyroid; Pituitary; Rectum; Skin/Subcutis; Spinal Cord; Spinal Nerve Roots, Cauda Equina; Spleen; Sternum; Stomach; Thymus; Thyroid; Tongue; Trachea; Urinary Bladder; Uterus; Vagina | | | | |
| The following tissues were examined microscopically and were unremarkable: Adrenal, Cortex; Adrenal, Medulla; Aorta; Brain; Cecum; Colon; Duodenum; Eye, Left; Femur; GALT/Peyer's Patch; Ganglion, Cervical Dorsal Root; Ganglion, Dorsal Root, Sacral; Ganglion, Superior Cervical; Ganglion, Thoracic Dorsal Root; Intrathecal Injection Site, Sacral Spinal Cord; Kidney; Liver; Lung; Lymph Node, Mandibular; Marrow, Femur; Marrow, Sternum; Muscle, Biceps Femoris; Nerve, Radial; Nerve, Sciatic; Nerve, Sural; Nerve, Ulnar; Pancreas; Parathyroid; Spinal Cord, Cervical; Spinal Cord, Thoracic; Spinal Nerve Roots, Cauda Equina; Spleen; Sternum; Stomach; Thymus; Thyroid | | | | |
| ‑‑‑‑‑‑‑‑‑‑‑‑‑‑‑‑‑‑‑‑‑‑‑‑‑‑‑‑‑‑‑‑‑‑‑‑‑‑‑‑‑‑‑‑‑‑‑‑‑‑‑‑‑‑‑‑‑‑‑‑‑‑‑‑‑‑‑‑‑‑‑‑‑‑‑‑‑‑‑‑‑‑‑‑ | | | | |
| **Animal Number: P0101 Group/Subgroup: 2/1 Sex: F Fate Status: Terminal Sacrifice**  **Date of Fate: 12 Oct 20 Phase of Fate: Dosing Phase Wk/Day of Fate: 4/28 TBW(g): 1900.0** | | | | |
| Organ Name: None | | | | |
| Macroscopic Observation(s)  None | | | | Microscopic Observation(s)  Ganglion, Cervical Dorsal Root: Autophagy; minimal  Ganglion, Superior Cervical: Vacuolation, neuron; minimal  Intrathecal Injection Site, Sacral Spinal Cord: MISSING  Intrathecal Injection Site, Sacral Spinal Cord: TISSUE COMMENT: Not available per histo comment  Kidney: Fibrosis, glomerular capsule; slight  Kidney: Infiltrate, mononuclear cell; minimal  Liver: Infiltrate, mononuclear cell; minimal  Lung: Infiltrate, macrophages, alveolus; minimal  Lymph Node, Mandibular: Lymphocytes, increased; minimal; follicle, unilateral  Ovary, Left: Immature; Present |
| The following tissues were examined macroscopically and were unremarkable: Adrenal; Animal; Aorta; Brain; Cecum; Cervix; Colon; Duodenum; Esophagus; Eye, Left; Femur; GALT/Peyer's Patch; Gall Bladder; Ganglion, Cervical Dorsal Root; Ganglion, Dorsal Root, Sacral; Ganglion, Lumbar Dorsal Root; Ganglion, Superior Cervical; Ganglion, Thoracic Dorsal Root; Heart; Ileum; Intrathecal Injection Site, Cisterna Magna; Intrathecal Injection Site, Sacral Spinal Cord; Jejunum; Kidney; Liver; Lung; Lymph Node, Mandibular; Lymph Node, Mesenteric; Mammary Gland; Mandibular Salivary Gland; Marrow, Femur; Marrow, Sternum; Muscle, Biceps Femoris; Nerve, Optic, Left; Nerve, Radial; Nerve, Sciatic; Nerve, Sural; Nerve, Tibial; Nerve, Ulnar; Ovary, Left; Pancreas; Parathyroid; Pituitary; Rectum; Skin/Subcutis; Spinal Cord; Spinal Nerve Roots, Cauda Equina; Spleen; Sternum; Stomach; Thymus; Thyroid; Tongue; Trachea; Urinary Bladder; Uterus; Vagina | | | | |
| The following tissues were examined microscopically and were unremarkable: Adrenal, Cortex; Adrenal, Medulla; Aorta; Brain; Cecum; Duodenum; Eye, Left; Femur; GALT/Peyer's Patch; Ganglion, Dorsal Root, Sacral; Ganglion, Lumbar Dorsal Root; Ganglion, Thoracic Dorsal Root; Heart; Intrathecal Injection Site, Cisterna Magna; Marrow, Femur; Marrow, Sternum; Muscle, Biceps Femoris; Nerve, Radial; Nerve, Sciatic; Nerve, Sural; Nerve, Tibial; Nerve, Ulnar; Pancreas; Parathyroid; Spinal Cord, Cervical; Spinal Cord, Lumbar; Spinal Cord, Thoracic; Spinal Nerve Roots, Cauda Equina; Spleen; Sternum; Stomach; Thymus; Thyroid | | | | |
| ‑‑‑‑‑‑‑‑‑‑‑‑‑‑‑‑‑‑‑‑‑‑‑‑‑‑‑‑‑‑‑‑‑‑‑‑‑‑‑‑‑‑‑‑‑‑‑‑‑‑‑‑‑‑‑‑‑‑‑‑‑‑‑‑‑‑‑‑‑‑‑‑‑‑‑‑‑‑‑‑‑‑‑‑ | | | | |
| **Animal Number: P0102 Group/Subgroup: 2/1 Sex: F Fate Status: Terminal Sacrifice**  **Date of Fate: 12 Oct 20 Phase of Fate: Dosing Phase Wk/Day of Fate: 4/28 TBW(g): 1800.0** | | | | |
| Organ Name: None | | | | |
| Macroscopic Observation(s)  Colon: Discolored; mucosa; multiple, up to 2 mm2; red; collected  Lung: Discolored; lobe; caudal; left; single, over 10 mm2; red; collected  Lung: Discolored; lobe; cranial; left; few, up to 10 mm2; red; collected | | | | Microscopic Observation(s)  Ganglion, Dorsal Root, Sacral: Infiltrate, mononuclear cell; minimal  Intrathecal Injection Site, Sacral Spinal Cord: MISSING  Intrathecal Injection Site, Sacral Spinal Cord: TISSUE COMMENT: Not available per histo comment  Liver: Infiltrate, mononuclear cell; minimal  Lymph Node, Mandibular: Lymphocytes, increased; minimal; follicle, unilateral  Ovary, Left: Immature; Present  Spinal Cord, Lumbar: Degeneration, axon, funiculus; minimal; focal; lateral funiculus white matter  Thyroid: Ectopic tissue, thymus; Present |
| The following tissues were examined macroscopically and were unremarkable: Adrenal; Animal; Aorta; Brain; Cecum; Cervix; Duodenum; Esophagus; Eye, Left; Femur; GALT/Peyer's Patch; Gall Bladder; Ganglion, Cervical Dorsal Root; Ganglion, Dorsal Root, Sacral; Ganglion, Lumbar Dorsal Root; Ganglion, Superior Cervical; Ganglion, Thoracic Dorsal Root; Heart; Ileum; Intrathecal Injection Site, Cisterna Magna; Intrathecal Injection Site, Sacral Spinal Cord; Jejunum; Kidney; Liver; Lymph Node, Mandibular; Lymph Node, Mesenteric; Mammary Gland; Mandibular Salivary Gland; Marrow, Femur; Marrow, Sternum; Muscle, Biceps Femoris; Nerve, Optic, Left; Nerve, Radial; Nerve, Sciatic; Nerve, Sural; Nerve, Tibial; Nerve, Ulnar; Ovary, Left; Pancreas; Parathyroid; Pituitary; Rectum; Skin/Subcutis; Spinal Cord; Spinal Nerve Roots, Cauda Equina; Spleen; Sternum; Stomach; Thymus; Thyroid; Tongue; Trachea; Urinary Bladder; Uterus; Vagina | | | | |
| The following tissues were examined microscopically and were unremarkable: Adrenal, Cortex; Adrenal, Medulla; Aorta; Brain; Cecum; Colon; Duodenum; Eye, Left; Femur; GALT/Peyer's Patch; Ganglion, Cervical Dorsal Root; Ganglion, Lumbar Dorsal Root; Ganglion, Superior Cervical; Ganglion, Thoracic Dorsal Root; Heart; Intrathecal Injection Site, Cisterna Magna; Kidney; Lung; Marrow, Femur; Marrow, Sternum; Muscle, Biceps Femoris; Nerve, Radial; Nerve, Sciatic; Nerve, Sural; Nerve, Tibial; Nerve, Ulnar; Pancreas; Parathyroid; Spinal Cord, Cervical; Spinal Cord, Thoracic; Spinal Nerve Roots, Cauda Equina; Spleen; Sternum; Stomach; Thymus | | | | |
| ‑‑‑‑‑‑‑‑‑‑‑‑‑‑‑‑‑‑‑‑‑‑‑‑‑‑‑‑‑‑‑‑‑‑‑‑‑‑‑‑‑‑‑‑‑‑‑‑‑‑‑‑‑‑‑‑‑‑‑‑‑‑‑‑‑‑‑‑‑‑‑‑‑‑‑‑‑‑‑‑‑‑‑‑ | | | | |
| **Animal Number: P0103 Group/Subgroup: 2/2 Sex: F Fate Status: Terminal Sacrifice**  **Date of Fate: 15 Oct 20 Phase of Fate: Dosing Phase Wk/Day of Fate: 4/28 TBW(g): 1300.0** | | | | |
| Organ Name: None | | | | |
| Macroscopic Observation(s)  Cecum: Discolored; mucosa; single, up to 5 mm2; red; collected/present on ileocecal junction  Colon: Discolored; mucosa; multiple, pinpoint; red; collected | | | | Microscopic Observation(s)  Cecum: Congestion/hemorrhage, agonal; minimal  Cecum: Parasite, protozoa, increased; slight  Intrathecal Injection Site, Cisterna Magna: Degeneration; moderate; focally extensive; lateral/The degeneration is unilateral in the dorsal portion of the lateral funiculus to include a unilateral dorsal (posterior) horn of the grey matter (consistent with a procedure related event).  Intrathecal Injection Site, Cisterna Magna: Gliosis, grey matter; moderate; focally extensive, unilateral; dorsal horn  Intrathecal Injection Site, Cisterna Magna: TISSUE COMMENT: The axon degeneration is unilateral in the dorsal portion of the lateral funiculus to include unilateral degeneration of the dorsal (posterior) horn of the grey matter (consistent with a procedure related event).  Lymph Node, Mandibular: Lymphocytes, increased; minimal; bilateral  Ovary, Left: Immature; Present  Spinal Cord, Cervical: Degeneration, axon, funiculus; slight; lateral & ventral  Spinal Cord, Lumbar: Degeneration, axon, funiculus; slight; lateral, ventral  Spinal Cord, Thoracic: Degeneration, axon, funiculus; slight; lateral, ventral |
| The following tissues were examined macroscopically and were unremarkable: Adrenal; Animal; Aorta; Brain; Cervix; Duodenum; Esophagus; Eye, Left; Femur; GALT/Peyer's Patch; Gall Bladder; Ganglion, Cervical Dorsal Root; Ganglion, Dorsal Root, Sacral; Ganglion, Lumbar Dorsal Root; Ganglion, Superior Cervical; Ganglion, Thoracic Dorsal Root; Heart; Ileum; Intrathecal Injection Site, Cisterna Magna; Intrathecal Injection Site, Sacral Spinal Cord; Jejunum; Kidney; Liver; Lung; Lymph Node, Mandibular; Lymph Node, Mesenteric; Mammary Gland; Mandibular Salivary Gland; Marrow, Femur; Marrow, Sternum; Muscle, Biceps Femoris; Nerve, Optic, Left; Nerve, Radial; Nerve, Sciatic; Nerve, Sural; Nerve, Tibial; Nerve, Ulnar; Ovary, Left; Pancreas; Parathyroid; Pituitary; Rectum; Skin/Subcutis; Spinal Cord; Spinal Nerve Roots, Cauda Equina; Spleen; Sternum; Stomach; Thymus; Thyroid; Tongue; Trachea; Urinary Bladder; Uterus; Vagina | | | | |
| The following tissues were examined microscopically and were unremarkable: Adrenal, Cortex; Adrenal, Medulla; Aorta; Brain; Colon; Duodenum; Eye, Left; Femur; GALT/Peyer's Patch; Ganglion, Cervical Dorsal Root; Ganglion, Dorsal Root, Sacral; Ganglion, Lumbar Dorsal Root; Ganglion, Superior Cervical; Ganglion, Thoracic Dorsal Root; Heart; Intrathecal Injection Site, Sacral Spinal Cord; Kidney; Liver; Lung; Marrow, Femur; Marrow, Sternum; Muscle, Biceps Femoris; Nerve, Radial; Nerve, Sciatic; Nerve, Sural; Nerve, Tibial; Nerve, Ulnar; Pancreas; Parathyroid; Spinal Nerve Roots, Cauda Equina; Spleen; Sternum; Stomach; Thymus; Thyroid | | | | |
| ‑‑‑‑‑‑‑‑‑‑‑‑‑‑‑‑‑‑‑‑‑‑‑‑‑‑‑‑‑‑‑‑‑‑‑‑‑‑‑‑‑‑‑‑‑‑‑‑‑‑‑‑‑‑‑‑‑‑‑‑‑‑‑‑‑‑‑‑‑‑‑‑‑‑‑‑‑‑‑‑‑‑‑‑ | | | | |
| **Animal Number: P0104 Group/Subgroup: 2/2 Sex: F Fate Status: Terminal Sacrifice**  **Date of Fate: 15 Oct 20 Phase of Fate: Dosing Phase Wk/Day of Fate: 4/28 TBW(g): 1500.0** | | | | |
| Organ Name: None | | | | |
| Macroscopic Observation(s)  Cecum: Discolored; mucosa; entire; single, up to 5 mm2; red; collected/Ileocecal junction.  Ileum: Discolored; mucosa; single, over 10 mm2; dark red; collected | | | | Microscopic Observation(s)  Brain: Infiltrate, mononuclear cell; minimal; focal; meninges/slide 50  Cecum: Congestion/hemorrhage, agonal; moderate  Ganglion, Lumbar Dorsal Root: Autophagy; minimal  Ganglion, Lumbar Dorsal Root: Infiltrate, mononuclear cell; minimal  Ganglion, Superior Cervical: Infiltrate, mononuclear cell; minimal  Heart: Infiltrate, mononuclear cell; minimal  Intrathecal Injection Site, Sacral Spinal Cord: MISSING  Intrathecal Injection Site, Sacral Spinal Cord: TISSUE COMMENT: Not available per histo comment  Kidney: Infiltrate, mononuclear cell; minimal  Kidney: Inflammation, mononuclear cell; minimal; focal  Lung: Infiltrate, macrophages, alveolus; minimal  Lymph Node, Mandibular: Lymphocytes, increased; slight; bilateral  Nerve, Sciatic: Infiltrate, mononuclear cell; minimal; focal; perineural adventitia  Nerve, Ulnar: Degeneration, axon; minimal  Ovary, Left: Immature; Present |
| The following tissues were examined macroscopically and were unremarkable : Adrenal; Animal; Aorta; Brain; Cervix; Colon; Duodenum; Esophagus; Eye, Left; Femur; GALT/Peyer's Patch; Gall Bladder; Ganglion, Cervical Dorsal Root; Ganglion, Dorsal Root, Sacral; Ganglion, Lumbar Dorsal Root; Ganglion, Superior Cervical; Ganglion, Thoracic Dorsal Root; Heart; Intrathecal Injection Site, Cisterna Magna; Intrathecal Injection Site, Sacral Spinal Cord; Jejunum; Kidney; Liver; Lung; Lymph Node, Mandibular; Lymph Node, Mesenteric; Mammary Gland; Mandibular Salivary Gland; Marrow, Femur; Marrow, Sternum; Muscle, Biceps Femoris; Nerve, Optic, Left; Nerve, Radial; Nerve, Sciatic; Nerve, Sural; Nerve, Tibial; Nerve, Ulnar; Ovary, Left; Pancreas; Parathyroid; Pituitary; Rectum; Skin/Subcutis; Spinal Cord; Spinal Nerve Roots, Cauda Equina; Spleen; Sternum; Stomach; Thymus; Thyroid; Tongue; Trachea; Urinary Bladder; Uterus; Vagina | | | | |
| The following tissues were examined microscopically and were unremarkable: Adrenal, Cortex; Adrenal, Medulla; Aorta; Duodenum; Eye, Left; Femur; GALT/Peyer's Patch; Ganglion, Cervical Dorsal Root; Ganglion, Dorsal Root, Sacral; Ganglion, Thoracic Dorsal Root; Ileum; Intrathecal Injection Site, Cisterna Magna; Liver; Marrow, Femur; Marrow, Sternum; Muscle, Biceps Femoris; Nerve, Radial; Nerve, Sural; Nerve, Tibial; Pancreas; Parathyroid; Spinal Cord, Cervical; Spinal Cord, Lumbar; Spinal Cord, Thoracic; Spinal Nerve Roots, Cauda Equina; Spleen; Sternum; Stomach; Thymus; Thyroid | | | | |
| ‑‑‑‑‑‑‑‑‑‑‑‑‑‑‑‑‑‑‑‑‑‑‑‑‑‑‑‑‑‑‑‑‑‑‑‑‑‑‑‑‑‑‑‑‑‑‑‑‑‑‑‑‑‑‑‑‑‑‑‑‑‑‑‑‑‑‑‑‑‑‑‑‑‑‑‑‑‑‑‑‑‑‑‑ | | | | |
| **Animal Number: P0201 Group/Subgroup: 3/1 Sex: F Fate Status: Terminal Sacrifice**  **Date of Fate: 12 Oct 20 Phase of Fate: Dosing Phase Wk/Day of Fate: 4/28 TBW(g): 1800.0** | | | | |
| Organ Name: None | | | | |
| Macroscopic Observation(s) | | | | Microscopic Observation(s)  Adipose, Brown: Infiltrate, mononuclear cell; minimal; pericardial/slide 26  Brain: Gliosis, white matter; minimal; focal/slide 49, 51  Brain: Infiltrate, mononuclear cell; slight; perivascular, meninges/neuropil/perivascular meninges: slide 44 perivascular meninges/neuropil: slide 46, 47, 48,49, 50, 51, 52, 53  Ganglion, Cervical Dorsal Root: Degeneration, axon, dorsal nerve root/spinal nerve; slight  Ganglion, Cervical Dorsal Root: Degeneration/necrosis, neuron; slight  Ganglion, Cervical Dorsal Root: Infiltrate, mononuclear cell; slight  Ganglion, Cervical Dorsal Root: Inflammation, mononuclear cell, dorsal nerve root/spinal nerve; slight  Ganglion, Cervical Dorsal Root: Inflammation, mononuclear cell, neuron; slight  Ganglion, Cervical Dorsal Root: Satellite glial cell, increased/neuronal cell loss; slight  Ganglion, Cervical Dorsal Root: Vacuolation, neuron; minimal  Ganglion, Cervical Dorsal Root: TISSUE COMMENT: Primarily affects 1 of 4 ganglia.  Ganglion, Dorsal Root, Sacral: Degeneration, axon, dorsal nerve root/spinal nerve; marked  Ganglion, Dorsal Root, Sacral: Degeneration/necrosis, neuron; moderate  Ganglion, Dorsal Root, Sacral: Infiltrate, mononuclear cell; marked  Ganglion, Dorsal Root, Sacral: Inflammation, mononuclear cell, dorsal nerve root/spinal nerve; moderate  Ganglion, Dorsal Root, Sacral: Inflammation, mononuclear cell, neuron; moderate  Ganglion, Dorsal Root, Sacral: Satellite glia; cell, increased/neuronal cell loss; slight  Ganglion, Dorsal Root, Sacral: TISSUE COMMENT: Similar findings in a ganglion present on slide 8E.  Ganglion, Lumbar Dorsal Root: Degeneration/necrosis, neuron; minimal  Ganglion, Lumbar Dorsal Root: Infiltrate, mononuclear cell; slight  Ganglion, Lumbar Dorsal Root: Infiltrate, mononuclear cell, dorsal nerve root/spinal nerve; minimal  Ganglion, Lumbar Dorsal Root: Inflammation, mononuclear cell, neuron; minimal  Ganglion, Lumbar Dorsal Root: Satellite glial cell, increased/neuronal cell loss; minimal  Ganglion, Lumbar Dorsal Root: Vacuolation, neuron; minimal  Ganglion, Superior Cervical: Infiltrate, mononuclear cell; minimal  Ganglion, Superior Cervical: Vacuolation, neuron; minimal  Ganglion, Thoracic Dorsal Root: Infiltrate, mononuclear cell; minimal  Heart: Infiltrate, mononuclear cell; slight  Intrathecal Injection Site, Cisterna Magna: MISSING  Intrathecal Injection Site, Cisterna Magna: TISSUE COMMENT: Not available per histo comment  Intrathecal Injection Site, Sacral Spinal Cord: Degeneration, axon, funiculus; slight; dorsal  Intrathecal Injection Site, Sacral Spinal Cord: Degeneration, axon, nerve roots/cauda equina; moderate  Intrathecal Injection Site, Sacral Spinal Cord: Gliosis, nerve roots/cauda equina; slight  Intrathecal Injection Site, Sacral Spinal Cord: Infiltrate, mononuclear cell; minimal; focal; meninges  Kidney: Infiltrate, mononuclear cell; minimal  Muscle, Biceps Femoris: Infiltrate, mononuclear cell; minimal  Nerve, Radial: BOTH MISSING  Nerve, Sciatic: Degeneration, axon; slight  Nerve, Sural: BOTH MISSING  Nerve, Tibial: Degeneration, axon; moderate  Nerve, Ulnar: Degeneration, axon; minimal  Ovary, Left: Immature; Present  Spinal Cord, Cervical: Degeneration, axon, funiculus; slight; dorsal funiculus white matter  Spinal Cord, Cervical: Gliosis, white matter; minimal; multifocal  Spinal Cord, Lumbar: Degeneration, axon, funiculus; moderate; dorsal and ventral/One ventral site is more ventrolateral.  Spinal Cord, Lumbar: Degeneration, axon, nerve root; marked  Spinal Cord, Lumbar: Gliosis, grey matter; slight; multifocal  Spinal Cord, Lumbar: Gliosis, white matter; minimal; focal  Spinal Cord, Lumbar: Infiltrate, mononuclear cell; minimal; focal; meninges  Spinal Cord, Thoracic: Degeneration, axon, funiculus; slight; dorsal funiculus white matter  Spinal Cord, Thoracic: Gliosis, grey matter; minimal; focal  Spinal Cord, Thoracic: Gliosis, white matter; slight; multifocal  Spinal Nerve Roots, Cauda Equina: Degeneration, axon; slight  Spinal Nerve Roots, Cauda Equina: Infiltrate, mononuclear cell; minimal; perivascular, adventitia  Spinal Nerve Roots, Cauda Equina: Inflammation, mononuclear cell; slight  Spinal Nerve Roots, Cauda Equina: TISSUE COMMENT: For clarity and greater specificity, spinal nerve roots are further defined to include the "caudal" spinal nerve roots.  Spleen: Pigment; slight/Consistent with hemosiderin |
| The following tissues were examined macroscopically and were unremarkable: Adrenal; Animal; Aorta; Brain; Cecum; Cervix; Colon; Duodenum; Esophagus; Eye, Left; Femur; GALT/Peyer's Patch; Gall Bladder; Ganglion, Cervical Dorsal Root; Ganglion, Dorsal Root, Sacral; Ganglion, Lumbar Dorsal Root; Ganglion, Superior Cervical; Ganglion, Thoracic Dorsal Root; Heart; Ileum; Intrathecal Injection Site, Cisterna Magna; Intrathecal Injection Site, Sacral Spinal Cord; Jejunum; Kidney; Liver; Lung; Lymph Node, Mandibular; Lymph Node, Mesenteric; Mammary Gland; Mandibular Salivary Gland; Marrow, Femur; Marrow, Sternum; Muscle, Biceps Femoris; Nerve, Optic, Left; Nerve, Radial; Nerve, Sciatic; Nerve, Sural; Nerve, Tibial; Nerve, Ulnar; Ovary, Left; Pancreas; Parathyroid; Pituitary; Rectum; Skin/Subcutis; Spinal Cord; Spinal Nerve Roots, Cauda Equina; Spleen; Sternum; Stomach; Thymus; Thyroid; Tongue; Trachea; Urinary Bladder; Uterus; Vagina | | | | |
| The following tissues were examined microscopically and were unremarkable: Adrenal, Cortex; Adrenal, Medulla; Aorta; Cecum; Duodenum; Eye, Left; Femur; GALT/Peyer's Patch; Liver; Lung; Lymph Node, Mandibular; Marrow, Femur; Marrow, Sternum; Pancreas; Parathyroid; Sternum; Stomach; Thymus; Thyroid | | | | |
| ‑‑‑‑‑‑‑‑‑‑‑‑‑‑‑‑‑‑‑‑‑‑‑‑‑‑‑‑‑‑‑‑‑‑‑‑‑‑‑‑‑‑‑‑‑‑‑‑‑‑‑‑‑‑‑‑‑‑‑‑‑‑‑‑‑‑‑‑‑‑‑‑‑‑‑‑‑‑‑‑‑‑‑‑ | | | | |
| **Animal Number: P0202 Group/Subgroup: 3/1 Sex: F Fate Status: Terminal Sacrifice**  **Date of Fate: 12 Oct 20 Phase of Fate: Dosing Phase Wk/Day of Fate: 4/28 TBW(g): 1900.0** | | | | |
| Organ Name: None | | | | |
| Macroscopic Observation(s)  None | | | | Microscopic Observation(s)  Brain: Gliosis, white matter; minimal; multifocal; white matter/slide 45, 46, 47, 50, 51 Gliosis in 51 surrounds a degenerative neuron.  Brain: Infiltrate, mononuclear cell; minimal; perivascular, meninges/neuropil/meninges/neuropil: slide 44, 45, 46, 50, 51 meninges: 47  Brain: Pigment; minimal; focally extensive/slide 51  Ganglion, Cervical Dorsal Root: Degeneration, axon, dorsal nerve root/spinal nerve; minimal  Ganglion, Cervical Dorsal Root: Degeneration/necrosis, neuron; slight  Ganglion, Cervical Dorsal Root: Infiltrate, mononuclear cell; slight  Ganglion, Cervical Dorsal Root: Inflammation, mononuclear cell, dorsal nerve root/spinal nerve; minimal  Ganglion, Cervical Dorsal Root: Inflammation, mononuclear cell, neuron; slight  Ganglion, Cervical Dorsal Root: Satellite glial cell, increased/neuronal cell loss; minimal  Ganglion, Dorsal Root, Sacral: Degeneration, axon, dorsal nerve root/spinal nerve; moderate  Ganglion, Dorsal Root, Sacral: Degeneration/necrosis, neuron; moderate  Ganglion, Dorsal Root, Sacral: Infiltrate, mononuclear cell; moderate; ganglia & adventitia  Ganglion, Dorsal Root, Sacral: Inflammation, mononuclear cell, dorsal nerve root/spinal nerve; moderate  Ganglion, Dorsal Root, Sacral: Inflammation, mononuclear cell, neuron; moderate  Ganglion, Dorsal Root, Sacral: Satellite glial cell, increased/neuronal cell loss; slight  Ganglion, Lumbar Dorsal Root: Autophagy; minimal  Ganglion, Lumbar Dorsal Root: Degeneration/necrosis, neuron; slight  Ganglion, Lumbar Dorsal Root: Infiltrate, mononuclear cell; slight  Ganglion, Lumbar Dorsal Root: Infiltrate, mononuclear cell, dorsal nerve root/spinal nerve; minimal  Ganglion, Lumbar Dorsal Root: Inflammation, mononuclear cell, neuron; slight  Ganglion, Thoracic Dorsal Root: Degeneration/necrosis, neuron; minimal  Ganglion, Thoracic Dorsal Root: Infiltrate, mononuclear cell; slight; periganglia, adventitia, nerve root/spinal nerve  Ganglion, Thoracic Dorsal Root: Inflammation, mononuclear cell, neuron; minimal  Heart: Inflammation, mononuclear cell; moderate  Intrathecal Injection Site, Cisterna Magna: MISSING  Intrathecal Injection Site, Cisterna Magna: TISSUE COMMENT: Not available per histo comment  Intrathecal Injection Site, Sacral Spinal Cord: Degeneration, axon, funiculus; minimal; dorsal  Intrathecal Injection Site, Sacral Spinal Cord: Degeneration, axon, nerve roots/cauda equina; slight  Intrathecal Injection Site, Sacral Spinal Cord: Gliosis, grey matter; slight/In one instance surrounds a neuron.  Intrathecal Injection Site, Sacral Spinal Cord: Gliosis, nerve roots/cauda equina; slight  Intrathecal Injection Site, Sacral Spinal Cord: Gliosis, white matter; slight  Liver: Hyperplasia, oval cell; minimal  Liver: Infiltrate, mononuclear cell; minimal  Lymph Node, Mandibular: Lymphocytes, increased; slight; bilateral  Muscle, Biceps Femoris: Infiltrate, mononuclear cell; minimal  Nerve, Radial: Degeneration, axon; minimal  Nerve, Ulnar: Degeneration, axon; slight  Ovary, Left: Immature; Present  Pancreas: Infiltrate, mononuclear cell, endocrine; minimal  Pancreas: Infiltrate, mononuclear cell, exocrine; slight  Parathyroid: Infiltrate, mononuclear cell; minimal  Spinal Cord, Cervical: Degeneration, axon, funiculus; minimal; dorsal funiculus white matter  Spinal Cord, Cervical: Gliosis, white matter; slight; multifocal  Spinal Cord, Cervical: Infiltrate, mononuclear cell, nerve roots; minimal  Spinal Cord, Lumbar: Gliosis, white matter; slight; multifocal  Spinal Cord, Thoracic: Degeneration, axon, funiculus; minimal; dorsal and lateral  Spinal Cord, Thoracic: Gliosis, white matter; minimal; multifocal  Spinal Nerve Roots, Cauda Equina: Degeneration, axon; slight  Spinal Nerve Roots, Cauda Equina: Gliosis; slight  Spinal Nerve Roots, Cauda Equina: Infiltrate, mononuclear cell; slight; perivascular, adventitia |
| The following tissues were examined macroscopically and were unremarkable: Adrenal; Animal; Aorta; Brain; Cecum; Cervix; Colon; Duodenum; Esophagus; Eye, Left; Femur; GALT/Peyer's Patch; Gall Bladder; Ganglion, Cervical Dorsal Root; Ganglion, Dorsal Root, Sacral; Ganglion, Lumbar Dorsal Root; Ganglion, Superior Cervical; Ganglion, Thoracic Dorsal Root; Heart; Ileum; Intrathecal Injection Site, Cisterna Magna; Intrathecal Injection Site, Sacral Spinal Cord; Jejunum; Kidney; Liver; Lung; Lymph Node, Mandibular; Lymph Node, Mesenteric; Mammary Gland; Mandibular Salivary Gland; Marrow, Femur; Marrow, Sternum; Muscle, Biceps Femoris; Nerve, Optic, Left; Nerve, Radial; Nerve, Sciatic; Nerve, Sural; Nerve, Tibial; Nerve, Ulnar; Ovary, Left; Pancreas; Parathyroid; Pituitary; Rectum; Skin/Subcutis; Spinal Cord; Spinal Nerve Roots, Cauda Equina; Spleen; Sternum; Stomach; Thymus; Thyroid; Tongue; Trachea; Urinary Bladder; Uterus; Vagina | | | | |
| The following tissues were examined microscopically and were unremarkable: Adrenal, Cortex; Adrenal, Medulla; Aorta; Cecum; Duodenum; Eye, Left; Femur; GALT/Peyer's Patch; Ganglion, Superior Cervical; Kidney; Lung; Marrow, Femur; Marrow, Sternum; Nerve, Sciatic; Nerve, Sural; Nerve, Tibial; Spleen; Sternum; Stomach; Thymus; Thyroid | | | | |
| ‑‑‑‑‑‑‑‑‑‑‑‑‑‑‑‑‑‑‑‑‑‑‑‑‑‑‑‑‑‑‑‑‑‑‑‑‑‑‑‑‑‑‑‑‑‑‑‑‑‑‑‑‑‑‑‑‑‑‑‑‑‑‑‑‑‑‑‑‑‑‑‑‑‑‑‑‑‑‑‑‑‑‑‑ | | | | |
| **Animal Number: P0203 Group/Subgroup: 3/2 Sex: F Fate Status: Terminal Sacrifice**  **Date of Fate: 15 Oct 20 Phase of Fate: Dosing Phase Wk/Day of Fate: 4/28 TBW(g): 1500.0** | | | | |
| Organ Name: None | | | | |
| Macroscopic Observation(s)  Cecum: Discolored; mucosa; single, up to 5 mm2; red; collected/present on ileocecal junction | | | | Microscopic Observation(s)  Cecum: Congestion/hemorrhage, agonal; slight  Cecum: Parasite, protozoa, increased; minimal  Ganglion, Cervical Dorsal Root: Infiltrate, mononuclear cell; minimal  Ganglion, Dorsal Root, Sacral: Degeneration/necrosis, neuron; minimal  Ganglion, Dorsal Root, Sacral: Infiltrate, mononuclear cell; slight  Ganglion, Dorsal Root, Sacral: Inflammation, mononuclear cell, neuron; minimal  Ganglion, Dorsal Root, Sacral: Vacuolation, neuron; minimal  Ganglion, Lumbar Dorsal Root: Degeneration/necrosis, neuron; minimal  Ganglion, Lumbar Dorsal Root: Infiltrate, mononuclear cell; minimal  Ganglion, Lumbar Dorsal Root: Inflammation, mononuclear cell, neuron; minimal  Ganglion, Superior Cervical: Infiltrate, mononuclear cell; minimal  Ganglion, Thoracic Dorsal Root: Autophagy; minimal  Heart: Infiltrate, mononuclear cell; minimal  Intrathecal Injection Site, Cisterna Magna: MISSING  Intrathecal Injection Site, Cisterna Magna: TISSUE COMMENT: Not available per histo comment  Intrathecal Injection Site, Sacral Spinal Cord: Degeneration, axon, funiculus; minimal; lateral  Kidney: Infiltrate, mononuclear cell; minimal  Liver: Infiltrate, mononuclear cell; slight; portal and random  Nerve, Sciatic: Infiltrate, mononuclear cell; minimal  Ovary, Left: Immature; Present  Parathyroid: Fibrosis, interstitial; minimal  Spinal Cord, Cervical: Degeneration, axon, funiculus; slight; lateral & ventral  Spinal Cord, Lumbar: Degeneration, axon, funiculus; minimal; lateral  Spinal Cord, Thoracic: Degeneration, axon, funiculus; slight; lateral, ventral |
| The following tissues were examined macroscopically and were unremarkable: Adrenal; Animal; Aorta; Brain; Cervix; Colon; Duodenum; Esophagus; Eye, Left; Femur; GALT/Peyer's Patch; Gall Bladder; Ganglion, Cervical Dorsal Root; Ganglion, Dorsal Root, Sacral; Ganglion, Lumbar Dorsal Root; Ganglion, Superior Cervical; Ganglion, Thoracic Dorsal Root; Heart; Ileum; Intrathecal Injection Site, Cisterna Magna; Intrathecal Injection Site, Sacral Spinal Cord; Jejunum; Kidney; Liver; Lung; Lymph Node, Mandibular; Lymph Node, Mesenteric; Mammary Gland; Mandibular Salivary Gland; Marrow, Femur; Marrow, Sternum; Muscle, Biceps Femoris; Nerve, Optic, Left; Nerve, Radial; Nerve, Sciatic; Nerve, Sural; Nerve, Tibial; Nerve, Ulnar; Ovary, Left; Pancreas; Parathyroid; Pituitary; Rectum; Skin/Subcutis; Spinal Cord; Spinal Nerve Roots, Cauda Equina; Spleen; Sternum; Stomach; Thymus; Thyroid; Tongue; Trachea; Urinary Bladder; Uterus; Vagina | | | | |
| The following tissues were examined microscopically and were unremarkable: Adrenal, Cortex; Adrenal, Medulla; Aorta; Brain; Duodenum; Eye, Left; Femur; GALT/Peyer's Patch; Lung; Lymph Node, Mandibular; Marrow, Femur; Marrow, Sternum; Muscle, Biceps Femoris; Nerve, Radial; Nerve, Sural; Nerve, Tibial; Nerve, Ulnar; Pancreas; Spinal Nerve Roots, Cauda Equina; Spleen; Sternum; Stomach; Thymus; Thyroid | | | | |
| ‑‑‑‑‑‑‑‑‑‑‑‑‑‑‑‑‑‑‑‑‑‑‑‑‑‑‑‑‑‑‑‑‑‑‑‑‑‑‑‑‑‑‑‑‑‑‑‑‑‑‑‑‑‑‑‑‑‑‑‑‑‑‑‑‑‑‑‑‑‑‑‑‑‑‑‑‑‑‑‑‑‑‑‑ | | | | |
| **Animal Number: P0204 Group/Subgroup: 3/2 Sex: F Fate Status: Terminal Sacrifice**  **Date of Fate: 15 Oct 20 Phase of Fate: Dosing Phase Wk/Day of Fate: 4/28 TBW(g): 1400.0** | | | | |
| Organ Name: None | | | | |
| Macroscopic Observation(s)  Colon: Discolored; mucosa; multiple, up to 2 mm2; red; collected  Ileum: Discolored; mucosa; single, up to 5 mm2; red; collected/Ileocecal junction | | Microscopic Observation(s)  Adipose, Brown: Infiltrate, mononuclear cell; minimal; pericardial/slide 25  Brain: Gliosis, white matter; slight; multifocal/slide 45, 47, 48  Brain: Infiltrate, mononuclear cell; minimal; perivascular, neuropil/slide 45, 47, 48  Cecum: Parasite, protozoa, increased; minimal  Colon: Inflammation, vessel; slight  Ganglion, Cervical Dorsal Root: Infiltrate, mononuclear cell; minimal  Ganglion, Dorsal Root, Sacral: Degeneration/necrosis, neuron; slight  Ganglion, Dorsal Root, Sacral: Infiltrate, mononuclear cell; slight; ganglia & adventitia  Ganglion, Dorsal Root, Sacral: Inflammation, mononuclear cell, neuron; slight  Ganglion, Dorsal Root, Sacral: TISSUE COMMENT: Also on slide 8E and 41.  Ganglion, Lumbar Dorsal Root: Infiltrate, mononuclear cell; slight  Ganglion, Thoracic Dorsal Root: Degeneration/necrosis, neuron; minimal; focal  Ganglion, Thoracic Dorsal Root: Inflammation, mononuclear cell, neuron; minimal; focal  Heart: Infiltrate, mononuclear cell; slight  Ileum: Congestion/hemorrhage; minimal; focal  Ileum: Inflammation, neutrophilic; slight; focal  Ileum: Inflammation, vessel; slight  Ileum: Parasite, protozoa, increased; slight  Ileum: TISSUE COMMENT: This is the ileocecal junction.  Kidney: Inflammation, mononuclear cell; slight; unilateral/Vascular/perivascular & interstitial.  Liver: Inflammation, mononuclear cell; slight; perivascular, portal/Large portal areas.  Lymph Node, Mandibular: Lymphocytes, increased; minimal; unilateral  Ovary, Left: Immature; Present  Pancreas: Atrophy; minimal; focal  Pancreas: Infiltrate, mononuclear cell, endocrine; minimal  Pancreas: Inflammation, vessel; slight  Spinal Cord, Cervical: Degeneration, axon, funiculus; minimal; lateral & ventral  Spinal Cord, Cervical: Gliosis, white matter; minimal; focal | | |
| The following tissues were examined macroscopically and were unremarkable: Adrenal; Animal; Aorta; Brain; Cecum; Cervix; Duodenum; Esophagus; Eye, Left; Femur; GALT/Peyer's Patch; Gall Bladder; Ganglion, Cervical Dorsal Root; Ganglion, Dorsal Root, Sacral; Ganglion, Lumbar Dorsal Root; Ganglion, Superior Cervical; Ganglion, Thoracic Dorsal Root; Heart; Intrathecal Injection Site, Cisterna Magna; Intrathecal Injection Site, Sacral Spinal Cord; Jejunum; Kidney; Liver; Lung; Lymph Node, Mandibular; Lymph Node, Mesenteric; Mammary Gland; Mandibular Salivary Gland; Marrow, Femur; Marrow, Sternum; Muscle, Biceps Femoris; Nerve, Optic, Left; Nerve, Radial; Nerve, Sciatic; Nerve, Sural; Nerve, Tibial; Nerve, Ulnar; Ovary, Left; Pancreas; Parathyroid; Pituitary; Rectum; Skin/Subcutis; Spinal Cord; Spinal Nerve Roots, Cauda Equina; Spleen; Sternum; Stomach; Thymus; Thyroid; Tongue; Trachea; Urinary Bladder; Uterus; Vagina | | | | |
| The following tissues were examined microscopically and were unremarkable: Adrenal, Cortex; Adrenal, Medulla; Aorta; Duodenum; Eye, Left; Femur; GALT/Peyer's Patch; Ganglion, Superior Cervical; Intrathecal Injection Site, Cisterna Magna; Intrathecal Injection Site, Sacral Spinal Cord; Lung; Marrow, Femur; Marrow, Sternum; Muscle, Biceps Femoris; Nerve, Radial; Nerve, Sciatic; Nerve, Sural; Nerve, Tibial; Nerve, Ulnar; Parathyroid; Spinal Cord, Lumbar; Spinal Cord, Thoracic; Spinal Nerve Roots, Cauda Equina; Spleen; Sternum; Stomach; Thymus; Thyroid | | | | |
| ‑‑‑‑‑‑‑‑‑‑‑‑‑‑‑‑‑‑‑‑‑‑‑‑‑‑‑‑‑‑‑‑‑‑‑‑‑‑‑‑‑‑‑‑‑‑‑‑‑‑‑‑‑‑‑‑‑‑‑‑‑‑‑‑‑‑‑‑‑‑‑‑‑‑‑‑‑‑‑‑‑‑‑‑ | | | | |
| **Animal Number: P0301 Group/Subgroup: 4/1 Sex: F Fate Status: Terminal Sacrifice**  **Date of Fate: 12 Oct 20 Phase of Fate: Dosing Phase Wk/Day of Fate: 4/28 TBW(g): 1300.0** | | | | |
| Organ Name: None | | | | |
| Macroscopic Observation(s)  None | | | | Microscopic Observation(s)  Adipose, Brown: Infiltrate, mononuclear cell; slight; periaorta/slide 9A  Brain: Gliosis, white matter; minimal; multifocal; white matter/slide 46, 47, 51  Brain: Infiltrate, mononuclear cell; minimal; perivascular, meninges/meninges: slide 47, 51  Cecum: Parasite, nematode; Present  Ganglion, Cervical Dorsal Root: Infiltrate, mononuclear cell; minimal  Ganglion, Dorsal Root, Sacral: Degeneration, axon, dorsal nerve root/spinal nerve; moderate  Ganglion, Dorsal Root, Sacral: Degeneration/necrosis, neuron; moderate  Ganglion, Dorsal Root, Sacral: Inflammation, mononuclear cell, dorsal nerve root/spinal nerve; slight  Ganglion, Dorsal Root, Sacral: Inflammation, mononuclear cell, neuron; slight  Ganglion, Dorsal Root, Sacral: TISSUE COMMENT: Most pronounced in 1 of 4 ganglia on primary slide and on slide 41 (cauda equina slide).  Ganglion, Lumbar Dorsal Root: Degeneration/necrosis, neuron; minimal  Ganglion, Lumbar Dorsal Root: Infiltrate, mononuclear cell; minimal  Ganglion, Lumbar Dorsal Root: Inflammation, mononuclear cell, neuron; minimal  Ganglion, Lumbar Dorsal Root: Vacuolation, neuron; slight  Ganglion, Thoracic Dorsal Root: Infiltrate, mononuclear cell; minimal  Heart: Inflammation, mononuclear cell; slight  Intrathecal Injection Site, Cisterna Magna: MISSING  Intrathecal Injection Site, Cisterna Magna: TISSUE COMMENT: Not available per histo comment  Intrathecal Injection Site, Sacral Spinal Cord: Degeneration, axon, funiculus; minimal; dorsal  Intrathecal Injection Site, Sacral Spinal Cord: Degeneration, axon, nerve roots/cauda equina; slight  Intrathecal Injection Site, Sacral Spinal Cord: Gliosis, grey matter; slight; multifocal  Intrathecal Injection Site, Sacral Spinal Cord: Gliosis, nerve roots/cauda equina; minimal  Kidney: Infiltrate, mononuclear cell; minimal  Liver: Infiltrate, mononuclear cell; minimal; portal  Lung: Infiltrate, macrophages, alveolus; minimal  Lymph Node, Mandibular: Lymphocytes, increased; slight; bilateral  Muscle, Biceps Femoris: Infiltrate, mononuclear cell; minimal  Ovary, Left: Immature; Present  Pancreas: Infiltrate, mononuclear cell, endocrine; minimal  Pancreas: Infiltrate, mononuclear cell, exocrine; slight  Parathyroid: Infiltrate, mononuclear cell; minimal  Spinal Cord, Cervical: Degeneration, axon, funiculus; minimal; dorsal funiculus white matter  Spinal Cord, Cervical: Gliosis, white matter; minimal; focal  Spinal Cord, Lumbar: Gliosis, white matter; minimal; focal  Spinal Cord, Lumbar: Infiltrate, mononuclear cell; minimal; meninges/Near the location where the dorsal nerve root exits.  Spinal Cord, Thoracic: Degeneration, axon, funiculus; minimal; dorsal funiculus white matter  Spinal Cord, Thoracic: Gliosis, white matter; minimal; multifocal  Spinal Nerve Roots, Cauda Equina: Degeneration, axon; slight  Spinal Nerve Roots, Cauda Equina: Gliosis; minimal  Spinal Nerve Roots, Cauda Equina: Infiltrate, mononuclear cell; slight  Thyroid: Ectopic tissue, thymus; Present |
| The following tissues were examined macroscopically and were unremarkable: Adrenal; Animal; Aorta; Brain; Cecum; Cervix; Colon; Duodenum; Esophagus; Eye, Left; Femur; GALT/Peyer's Patch; Gall Bladder; Ganglion, Cervical Dorsal Root; Ganglion, Dorsal Root, Sacral; Ganglion, Lumbar Dorsal Root; Ganglion, Superior Cervical; Ganglion, Thoracic Dorsal Root; Heart; Ileum; Intrathecal Injection Site, Cisterna Magna; Intrathecal Injection Site, Sacral Spinal Cord; Jejunum; Kidney; Liver; Lung; Lymph Node, Mandibular; Lymph Node, Mesenteric; Mammary Gland; Mandibular Salivary Gland; Marrow, Femur; Marrow, Sternum; Muscle, Biceps Femoris; Nerve, Optic, Left; Nerve, Radial; Nerve, Sciatic; Nerve, Sural; Nerve, Tibial; Nerve, Ulnar; Ovary, Left; Pancreas; Parathyroid; Pituitary; Rectum; Skin/Subcutis; Spinal Cord; Spinal Nerve Roots, Cauda Equina; Spleen; Sternum; Stomach; Thymus; Thyroid; Tongue; Trachea; Urinary Bladder; Uterus; Vagina | | | | |
| The following tissues were examined microscopically and were unremarkable: Adrenal, Cortex; Adrenal, Medulla; Aorta; Duodenum; Eye, Left; Femur; GALT/Peyer's Patch; Ganglion, Superior Cervical; Marrow, Femur; Marrow, Sternum; Nerve, Radial; Nerve, Sciatic; Nerve, Sural; Nerve, Tibial; Nerve, Ulnar; Spleen; Sternum; Stomach; Thymus | | | | |
| ‑‑‑‑‑‑‑‑‑‑‑‑‑‑‑‑‑‑‑‑‑‑‑‑‑‑‑‑‑‑‑‑‑‑‑‑‑‑‑‑‑‑‑‑‑‑‑‑‑‑‑‑‑‑‑‑‑‑‑‑‑‑‑‑‑‑‑‑‑‑‑‑‑‑‑‑‑‑‑‑‑‑‑‑ | | | | |
| **Animal Number: P0302 Group/Subgroup: 4/1 Sex: F Fate Status: Terminal Sacrifice**  **Date of Fate: 12 Oct 20 Phase of Fate: Dosing Phase Wk/Day of Fate: 4/28 TBW(g): 1800.0** | | | | |
| Organ Name None | | | | |
| Macroscopic Observation(s)  None | | | | Microscopic Observation(s)  Adipose, Brown: Infiltrate, mononuclear cell; minimal; pericardial/slide 25, 26  Brain: Gliosis, grey matter; slight/These are often in the cortex subjacent to the  meninges (surface). slide 47, 48  Brain: Gliosis, white matter; minimal; white matter/slide 45, 47, 51, 52  Brain: Hematopoiesis, extramedullary, choroid plexus; minimal; focal  Brain: Infiltrate, mononuclear cell; slight; perivascular, meninges/neuropil/meninges: slide 45, 46, 47, 51meninges/neuropil: 44, 49  Cecum: Parasite, nematode; Present  Ganglion, Cervical Dorsal Root: Degeneration, axon, dorsal nerve root/spinal nerve; marked  Ganglion, Cervical Dorsal Root: Degeneration/necrosis, neuron; moderate  Ganglion, Cervical Dorsal Root: Infiltrate, mononuclear cell; moderate  Ganglion, Cervical Dorsal Root: Inflammation, mononuclear cell, dorsal nerve root/spinal nerve; moderate  Ganglion, Cervical Dorsal Root: Inflammation, mononuclear cell, neuron; moderate  Ganglion, Cervical Dorsal Root: Vacuolation, neuron; minimal  Ganglion, Dorsal Root, Sacral: Degeneration, axon, dorsal nerve root/spinal nerve; moderate  Ganglion, Dorsal Root, Sacral: Degeneration/necrosis, neuron; moderate  Ganglion, Dorsal Root, Sacral: Infiltrate, mononuclear cell; moderate  Ganglion, Dorsal Root, Sacral: Inflammation, cell, dorsal nerve root/spinal nerve; slight  Ganglion, Dorsal Root, Sacral: Inflammation, mononuclear cell, neuron; moderate  Ganglion, Dorsal Root, Sacral: TISSUE COMMENT: Also see slide 41 (cauda equina)  Ganglion, Lumbar Dorsal Root: Degeneration/necrosis, neuron; minimal  Ganglion, Lumbar Dorsal Root: Infiltrate, mononuclear cell; slight  Ganglion, Lumbar Dorsal Root: Infiltrate, mononuclear cell, dorsal nerve root/spinal nerve; minimal  Ganglion, Lumbar Dorsal Root: Inflammation, mononuclear cell, neuron; minimal  Ganglion, Superior Cervical: Infiltrate, mononuclear cell; minimal  Ganglion, Thoracic Dorsal Root: Degeneration, axon, dorsal nerve root/spinal nerve; moderate  Ganglion, Thoracic Dorsal Root: Degeneration/necrosis, neuron; slight  Ganglion, Thoracic Dorsal Root: Infiltrate, mononuclear cell; moderate  Ganglion, Thoracic Dorsal Root: Inflammation, mononuclear cell, dorsal nerve root/spinal nerve; slight  Ganglion, Thoracic Dorsal Root: Inflammation, mononuclear cell, neuron; slight  Heart: Inflammation, mononuclear cell; slight  Intrathecal Injection Site, Cisterna Magna: MISSING  Intrathecal Injection Site, Cisterna Magna: TISSUE COMMENT: Not available per histo comment  Intrathecal Injection Site, Sacral Spinal Cord: Degeneration, axon, funiculus; slight; dorsal  Intrathecal Injection Site, Sacral Spinal Cord: Degeneration, axon, nerve roots/cauda equina; moderate; dorsal  Intrathecal Injection Site, Sacral Spinal Cord: Gliosis, grey matter; slight; multifocal  Intrathecal Injection Site, Sacral Spinal Cord: Gliosis, nerve roots/cauda equina; moderate; dorsal  Intrathecal Injection Site, Sacral Spinal Cord:  Gliosis, white matter; minimal; focal  Intrathecal Injection Site, Sacral Spinal Cord:  Infiltrate, mononuclear cell; slight; meninges  Kidney: Infiltrate, mononuclear cell; slight  Liver: Hyperplasia, oval cell; minimal  Liver: Infiltrate, mononuclear cell; slight; portal  Liver: Necrosis, hepatocyte, increased; minimal  Lymph Node, Mandibular: Lymphocytes, increased; minimal; unilateral  Muscle, Biceps Femoris: Infiltrate, mononuclear cell; minimal  Nerve, Radial: Degeneration, axon; slight  Nerve, Sciatic: Infiltrate, mononuclear cell; minimal  Nerve, Sural: Degeneration, axon; slight  Nerve, Tibial: Degeneration, axon; minimal  Nerve, Ulnar: Degeneration, axon; moderate  Ovary, Left: Immature; Present  Pancreas: Infiltrate, mononuclear cell, endocrine; slight  Pancreas: Infiltrate, mononuclear cell, exocrine; slight  Spinal Cord, Cervical: Degeneration, axon, funiculus; moderate; dorsal funiculus white matter  Spinal Cord, Cervical: Degeneration, axon, nerve root; minimal  Spinal Cord, Cervical: Gliosis, grey matter; minimal  Spinal Cord, Cervical: Gliosis, nerve root; minimal  Spinal Cord, Lumbar: Degeneration, axon, funiculus;  slight; dorsal funiculus white matter  Spinal Cord, Lumbar: Gliosis, white matter; minimal; multifocal  Spinal Cord, Thoracic: Degeneration, axon, funiculus; slight; dorsal funiculus white matter  Spinal Cord, Thoracic: Gliosis, nerve root; minimal  Spinal Cord, Thoracic: Gliosis, white matter; minimal; multifocal  Spinal Nerve Roots, Cauda Equina: Degeneration, axon; slight  Spinal Nerve Roots, Cauda Equina: Gliosis; slight  Spinal Nerve Roots, Cauda Equina: Infiltrate, mononuclear cell; slight  Stomach: Infiltrate, mononuclear cell, ganglia/nerve; minimal |
| The following tissues were examined macroscopically and were unremarkable: Adrenal; Animal; Aorta; Brain; Cecum; Cervix; Colon; Duodenum; Esophagus; Eye, Left; Femur; GALT/Peyer's Patch; Gall Bladder; Ganglion, Cervical Dorsal Root; Ganglion, Dorsal Root, Sacral; Ganglion, Lumbar Dorsal Root; Ganglion, Superior Cervical; Ganglion, Thoracic Dorsal Root; Heart; Ileum; Intrathecal Injection Site, Cisterna Magna; Intrathecal Injection Site, Sacral Spinal Cord; Jejunum; Kidney; Liver; Lung; Lymph Node, Mandibular; Lymph Node, Mesenteric; Mammary Gland; Mandibular Salivary Gland; Marrow, Femur; Marrow, Sternum; Muscle, Biceps Femoris; Nerve, Optic, Left; Nerve, Radial; Nerve, Sciatic; Nerve, Sural; Nerve, Tibial; Nerve, Ulnar; Ovary, Left; Pancreas; Parathyroid; Pituitary; Rectum; Skin/Subcutis; Spinal Cord; Spinal Nerve Roots, Cauda Equina; Spleen; Sternum; Stomach; Thymus; Thyroid; Tongue; Trachea; Urinary Bladder; Uterus; Vagina | | | | |
| The following tissues were examined microscopically and were unremarkable: Adrenal, Cortex; Adrenal, Medulla; Aorta; Duodenum; Eye, Left; Femur; GALT/Peyer's Patch; Lung; Marrow, Femur; Marrow, Sternum; Parathyroid; Spleen; Sternum; Thymus; Thyroid | | | | |
| ‑‑‑‑‑‑‑‑‑‑‑‑‑‑‑‑‑‑‑‑‑‑‑‑‑‑‑‑‑‑‑‑‑‑‑‑‑‑‑‑‑‑‑‑‑‑‑‑‑‑‑‑‑‑‑‑‑‑‑‑‑‑‑‑‑‑‑‑‑‑‑‑‑‑‑‑‑‑‑‑‑‑‑‑ | | | | |
| **Animal Number: P0303 Group/Subgroup: 4/2 Sex: F Fate Status: Terminal Sacrifice**  **Date of Fate: 15 Oct 20 Phase of Fate: Dosing Phase Wk/Day of Fate: 4/28 TBW(g): 1500.0** | | | | |
| Organ Name None | | | | |
| Macroscopic Observation(s)  Cecum: Discolored; mucosa; single, up to 5 mm2; red; collected/present on ileocecal junction  Lymph Node, Other: Large; renal; bilateral; present; collected | | | | Microscopic Observation(s)  Adipose, Brown: Infiltrate, mononuclear cell; minimal; periaorta  Brain: Infiltrate, mononuclear cell; minimal; meninges/meninges: slide 44, 46, 49, 53  Cecum: Congestion/hemorrhage, agonal; minimal  Cecum: Parasite, nematode; Present  Ganglion, Cervical Dorsal Root: Infiltrate, mononuclear cell; minimal  Ganglion, Dorsal Root, Sacral: Autophagy; minimal  Ganglion, Dorsal Root, Sacral: Degeneration/necrosis, neuron; minimal  Ganglion, Dorsal Root, Sacral: Inflammation, mononuclear cell, neuron; minimal  Ganglion, Lumbar Dorsal Root: Autophagy; minimal  Ganglion, Lumbar Dorsal Root: Infiltrate, mononuclear cell; minimal  Ganglion, Superior Cervical: Infiltrate, mononuclear cell; minimal  Ganglion, Thoracic Dorsal Root: Infiltrate, mononuclear cell; minimal  Ganglion, Thoracic Dorsal Root: Inflammation, mononuclear cell, muscle; slight  Heart: Infiltrate, mononuclear cell; slight  Intrathecal Injection Site, Cisterna Magna: MISSING  Intrathecal Injection Site, Cisterna Magna: TISSUE COMMENT: Not available per histo comment  Intrathecal Injection Site, Sacral Spinal Cord: Degeneration, axon, funiculus; minimal; dorsal  Intrathecal Injection Site, Sacral Spinal Cord: Infiltrate, mononuclear cell; minimal; meninges  Kidney: Degeneration/necrosis, tubule; minimal  Kidney: Infiltrate, mononuclear cell; slight; juxtaglomerular apparatus  Liver: Hyperplasia, oval cell; minimal  Liver: Infiltrate, mononuclear cell; slight; portal/random  Liver: Necrosis, hepatocyte, increased; minimal  Lymph Node, Mandibular: Lymphocytes, increased; minimal; follicle  Ovary, Left: Immature; Present  Spinal Cord, Cervical: Degeneration, axon, funiculus; minimal; dorsal, lateral/The lateral focus is characterized by a markedly swollen axon just within the grey matter.  Spinal Cord, Cervical: Gliosis, white matter; minimal; focal  Spinal Cord, Lumbar: Degeneration, axon, funiculus; minimal; dorsal funiculus white matter  Spinal Cord, Thoracic: Degeneration, axon, funiculus; minimal; dorsal funiculus white matter |
| The following tissues were examined macroscopically and were unremarkable: Adrenal; Animal; Aorta; Brain; Cervix; Colon; Duodenum; Esophagus; Eye, Left; Femur; GALT/Peyer's Patch; Gall Bladder; Ganglion, Cervical Dorsal Root; Ganglion, Dorsal Root, Sacral; Ganglion, Lumbar Dorsal Root; Ganglion, Superior Cervical; Ganglion, Thoracic Dorsal Root; Heart; Ileum; Intrathecal Injection Site, Cisterna Magna; Intrathecal Injection Site, Sacral Spinal Cord; Jejunum; Kidney; Liver; Lung; Lymph Node, Mandibular; Lymph Node, Mesenteric; Mammary Gland; Mandibular Salivary Gland; Marrow, Femur; Marrow, Sternum; Muscle, Biceps Femoris; Nerve, Optic, Left; Nerve, Radial; Nerve, Sciatic; Nerve, Sural; Nerve, Tibial; Nerve, Ulnar; Ovary, Left; Pancreas; Parathyroid; Pituitary; Rectum; Skin/Subcutis; Spinal Cord; Spinal Nerve Roots, Cauda Equina; Spleen; Sternum; Stomach; Thymus; Thyroid; Tongue; Trachea; Urinary Bladder; Uterus; Vagina | | | | |
| The following tissues were examined microscopically and were unremarkable: Adrenal, Cortex; Adrenal, Medulla; Aorta; Duodenum; Eye, Left; Femur; GALT/Peyer's Patch; Lung; Lymph Node, Other; Marrow, Femur; Marrow, Sternum; Muscle, Biceps Femoris; Nerve, Radial; Nerve, Sciatic; Nerve, Sural; Nerve, Tibial; Nerve, Ulnar; Pancreas; Parathyroid; Spinal Nerve Roots, Cauda Equina; Spleen; Sternum; Stomach; Thymus; Thyroid | | | | |
| ‑‑‑‑‑‑‑‑‑‑‑‑‑‑‑‑‑‑‑‑‑‑‑‑‑‑‑‑‑‑‑‑‑‑‑‑‑‑‑‑‑‑‑‑‑‑‑‑‑‑‑‑‑‑‑‑‑‑‑‑‑‑‑‑‑‑‑‑‑‑‑‑‑‑‑‑‑‑‑‑‑‑‑‑ | | | | |
| **Animal Number: P0304 Group/Subgroup: 4/2 Sex: F Fate Status: Terminal Sacrifice**  **Date of Fate: 15 Oct 20 Phase of Fate: Dosing Phase Wk/Day of Fate: 4/28 TBW(g): 1500.0** | | | | |
| Organ Name: None | | | | |
| Macroscopic Observation(s)  None | | | | Microscopic Observation(s)  Adipose, Brown: Infiltrate, mononuclear cell; moderate; periaorta, perisuperior cervical ganglia/slide 9A, 40 Brain: Gliosis, grey matter; slight; multifocal; all regions/slide 46, 47, 48, 50  Brain: Gliosis, white matter; slight; multifocal; regions/slide 45, 47, 48, 49, 50, 51, 52  Brain: Infiltrate, mononuclear cell; slight; perivascular, meninges/neuropil/slide 44, 45, 46, 47, 48, 49, 51  Ganglion, Cervical Dorsal Root: Autophagy; minimal  Ganglion, Cervical Dorsal Root: Degeneration/necrosis, neuron; minimal  Ganglion, Cervical Dorsal Root: Infiltrate, mononuclear cell; minimal  Ganglion, Cervical Dorsal Root: Inflammation, mononuclear cell, neuron; minimal  Ganglion, Cervical Dorsal Root: Satellite glial cell, increased/neuronal cell loss; minimal  Ganglion, Cervical Dorsal Root: Vacuolation, neuron; minimal  Ganglion, Dorsal Root, Sacral: Autophagy; minimal  Ganglion, Dorsal Root, Sacral: Degeneration, axon, dorsal nerve root/spinal nerve; slight  Ganglion, Dorsal Root, Sacral: Degeneration/necrosis, neuron; slight  Ganglion, Dorsal Root, Sacral: Infiltrate, mononuclear cell; slight  Ganglion, Dorsal Root, Sacral: Infiltrate, mononuclear cell, dorsal nerve root/spinal nerve; minimal  Ganglion, Dorsal Root, Sacral: Inflammation, mononuclear cell, dorsal nerve root/spinal nerve; slight  Ganglion, Dorsal Root, Sacral: Inflammation, mononuclear cell, neuron; slight  Ganglion, Dorsal Root, Sacral: Inflammation, vessel; slight  Ganglion, Lumbar Dorsal Root: Autophagy; minimal  Ganglion, Lumbar Dorsal Root: Degeneration/necrosis, neuron; minimal  Ganglion, Lumbar Dorsal Root: Infiltrate, mononuclear cell; slight  Ganglion, Lumbar Dorsal Root: Inflammation, mononuclear cell, neuron; minimal  Ganglion, Thoracic Dorsal Root: Autophagy; minimal  Ganglion, Thoracic Dorsal Root: Degeneration/necrosis, neuron; minimal  Ganglion, Thoracic Dorsal Root: Inflammation, mononuclear cell, neuron; minimal  Heart: Inflammation, mononuclear cell; moderate  Intrathecal Injection Site, Cisterna Magna: Degeneration; slight; dorsal  Intrathecal Injection Site, Cisterna Magna: Gliosis, white matter; minimal; multifocal  Intrathecal Injection Site, Cisterna Magna: Infiltrate, mononuclear cell; minimal; meninges  Intrathecal Injection Site, Sacral Spinal Cord: Degeneration, axon, nerve roots/cauda equina; moderate  Kidney: Degeneration/necrosis, tubule; moderate  Kidney: Glomerulonephritis; moderate  Kidney: Hyperplasia/hypertrophy, Bowman's capsule; slight  Kidney: Inclusion, basophilic, tubule cell; Present/Consistent with cytomegalovirus.  Liver: Infiltrate, mononuclear cell; minimal; portal  Lung: Infiltrate, macrophages, alveolus; minimal  Lymph Node, Mandibular: Lymphocytes, increased; slight; follicle  Muscle, Biceps Femoris: Infiltrate, mononuclear cell; minimal  Nerve, Radial: Degeneration, axon; minimal  Nerve, Sciatic: Infiltrate, mononuclear cell; minimal  Nerve, Ulnar: Degeneration, axon; slight  Nerve, Ulnar: Infiltrate, mononuclear cell; minimal  Ovary, Left: Immature; Present  Pancreas: Infiltrate, mononuclear cell, endocrine; minimal  Pancreas: Infiltrate, mononuclear cell, exocrine; minimal  Spinal Cord, Cervical: Degeneration, axon, funiculus; slight; dorsal funiculus white matter  Spinal Cord, Cervical: Gliosis, white matter; minimal; multifocal  Spinal Cord, Lumbar: Degeneration, axon, funiculus; slight; dorsal funiculus white matter  Spinal Cord, Lumbar: Gliosis, grey matter; slight; multifocal  Spinal Cord, Lumbar: Gliosis, white matter; moderate; multifocal  Spinal Cord, Lumbar: Infiltrate, mononuclear cell; slight; meninges  Spinal Cord, Lumbar: Infiltrate, mononuclear cell, nerve roots; minimal  Spinal Cord, Thoracic: Degeneration, axon, funiculus; minimal; dorsal funiculus white matter  Spinal Cord, Thoracic: Gliosis, white matter; slight; multifocal  Spinal Nerve Roots, Cauda Equina: Degeneration, axon; moderate  Spinal Nerve Roots, Cauda Equina: Gliosis; slight  Spleen: Lymphocytes, increased; slight  Thyroid: Ectopic tissue, thymus; Present  Thyroid: Infiltrate, mononuclear cell; minimal; unilateral |
| The following tissues were examined macroscopically and were unremarkable: Adrenal; Animal; Aorta; Brain; Cecum; Cervix; Colon; Duodenum; Esophagus; Eye, Left; Femur; GALT/Peyer's Patch; Gall Bladder; Ganglion, Cervical Dorsal Root; Ganglion, Dorsal Root, Sacral; Ganglion, Lumbar Dorsal Root; Ganglion, Superior Cervical; Ganglion, Thoracic Dorsal Root; Heart; Ileum; Intrathecal Injection Site, Cisterna Magna; Intrathecal Injection Site, Sacral Spinal Cord; Jejunum; Kidney; Liver; Lung; Lymph Node, Mandibular; Lymph Node, Mesenteric; Mammary Gland; Mandibular Salivary Gland; Marrow, Femur; Marrow, Sternum; Muscle, Biceps Femoris; Nerve, Optic, Left; Nerve, Radial; Nerve, Sciatic; Nerve, Sural; Nerve, Tibial; Nerve, Ulnar; Ovary, Left; Pancreas; Parathyroid; Pituitary; Rectum; Skin/Subcutis; Spinal Cord; Spinal Nerve Roots, Cauda Equina; Spleen; Sternum; Stomach; Thymus; Thyroid; Tongue; Trachea; Urinary Bladder; Uterus; Vagina | | | | |
| The following tissues were examined microscopically and were unremarkable: Adrenal, Cortex; Adrenal, Medulla; Aorta; Cecum; Duodenum; Eye, Left; Femur; GALT/Peyer's Patch; Ganglion, Superior Cervical; Marrow, Femur; Marrow, Sternum; Nerve, Sural; Nerve, Tibial; Parathyroid; Sternum; Stomach; Thymus | | | | |
| ‑‑‑‑‑‑‑‑‑‑‑‑‑‑‑‑‑‑‑‑‑‑‑‑‑‑‑‑‑‑‑‑‑‑‑‑‑‑‑‑‑‑‑‑‑‑‑‑‑‑‑‑‑‑‑‑‑‑‑‑‑‑‑‑‑‑‑‑‑‑‑‑‑‑‑‑‑‑‑‑‑‑‑‑ | | | | |
| **Animal Number: P0401 Group/Subgroup: 5/1 Sex: F Fate Status: Terminal Sacrifice**  **Date of Fate: 12 Oct 20 Phase of Fate: Dosing Phase Wk/Day of Fate: 4/28 TBW(g): 1300.0** | | | | |
| Organ Name: None | | | | |
| Macroscopic Observation(s)  None | | | | Microscopic Observation(s)  Adipose, Brown: Infiltrate, mononuclear cell; minimal; pericardial/slide 25R  Adrenal, Medulla: Infiltrate, mononuclear cell; minimal; unilateral  Brain: Gliosis, white matter; minimal; focal/slide 51  Brain: Infiltrate, mononuclear cell; slight; perivascular, meninges/neuropil/meninges: slide 44, 47, 48, 52 neuropil: slide 45, 46  Eye, Left: Infiltrate, mononuclear cell; minimal; focal; limbus  Ganglion, Cervical Dorsal Root: Degeneration/necrosis, neuron; slight  Ganglion, Cervical Dorsal Root: Infiltrate, mononuclear cell; minimal  Ganglion, Cervical Dorsal Root: Inflammation, mononuclear cell, neuron; slight  Ganglion, Dorsal Root, Sacral: Degeneration, axon, dorsal nerve root/spinal nerve; moderate  Ganglion, Dorsal Root, Sacral: Degeneration/necrosis, neuron; moderate  Ganglion, Dorsal Root, Sacral: Infiltrate, mononuclear cell; moderate; ganglia & adventitia  Ganglion, Dorsal Root, Sacral: Inflammation, mononuclear cell, dorsal nerve root/spinal nerve; moderate  Ganglion, Dorsal Root, Sacral: Inflammation, mononuclear cell, neuron; moderate  Ganglion, Lumbar Dorsal Root: Degeneration/necrosis, neuron; moderate  Ganglion, Lumbar Dorsal Root: Infiltrate, mononuclear cell; slight  Ganglion, Lumbar Dorsal Root: Infiltrate, mononuclear cell, dorsal nerve root/spinal nerve; slight  Ganglion, Lumbar Dorsal Root: Inflammation, mononuclear cell, neuron; slight  Ganglion, Superior Cervical: Vacuolation, neuron; minimal  Ganglion, Thoracic Dorsal Root: Infiltrate, mononuclear cell; minimal  Heart: Infiltrate, mononuclear cell; minimal  Intrathecal Injection Site, Cisterna Magna: Degeneration; slight; dorsal, lateral, ventral/Most prominent in dorsal and ventral funiculi.  Intrathecal Injection Site, Cisterna Magna: Gliosis, white matter; minimal; multifocal  Intrathecal Injection Site, Sacral Spinal Cord: MISSING  Intrathecal Injection Site, Sacral Spinal Cord: TISSUE COMMENT: Not available per histo comment  Kidney: Inclusion, basophilic, tubule cell; Present/Slide 17. Consistent with cytomegalovirus a common background finding in juvenile monkeys.  Kidney: Infiltrate, mononuclear cell; slight  Liver: Infiltrate, mononuclear cell; minimal; portal  Lymph Node, Mandibular: Lymphocytes, increased; slight; bilateral  Nerve, Sciatic: Degeneration, axon; minimal  Nerve, Sciatic: Infiltrate, mononuclear cell; minimal  Nerve, Sural: BOTH MISSING  Nerve, Tibial: Degeneration, axon; minimal  Ovary, Left: Immature; Present  Pancreas: Hemorrhage; slight; focal  Parathyroid: MISSING  Spinal Cord, Cervical: Degeneration, axon, funiculus; moderate; dorsal, lateral, & ventral/Most prominent in ventral and dorsal funiculi.  Spinal Cord, Cervical: Gliosis, white matter; minimal; multifocal  Spinal Cord, Lumbar: Degeneration, axon, funiculus; slight; dorsal, lateral, and ventral/Most prominent in dorsal funiculi.  Spinal Cord, Lumbar: Gliosis, white matter; minimal; multifocal  Spinal Cord, Thoracic: Degeneration, axon, funiculus; slight; dorsal, lateral, & ventral/Most prominent in dorsal funiculus.  Spinal Cord, Thoracic: Gliosis, grey matter; minimal; multifocal  Spinal Nerve Roots, Cauda Equina: Degeneration, axon; slight  Spinal Nerve Roots, Cauda Equina: Gliosis; slight  Thyroid: BOTH MISSING |
| The following tissues were examined macroscopically and were unremarkable: Adrenal; Animal; Aorta; Brain; Cecum; Cervix; Colon; Duodenum; Esophagus; Eye, Left; Femur; GALT/Peyer's Patch; Gall Bladder; Ganglion, Cervical Dorsal Root; Ganglion, Dorsal Root, Sacral; Ganglion, Lumbar Dorsal Root; Ganglion, Superior Cervical; Ganglion, Thoracic Dorsal Root; Heart; Ileum; Intrathecal Injection Site, Cisterna Magna; Intrathecal Injection Site, Sacral Spinal Cord; Jejunum; Kidney; Liver; Lung; Lymph Node, Mandibular; Lymph Node, Mesenteric; Mammary Gland; Mandibular Salivary Gland; Marrow, Femur; Marrow, Sternum; Muscle, Biceps Femoris; Nerve, Optic, Left; Nerve, Radial; Nerve, Sciatic; Nerve, Sural; Nerve, Tibial; Nerve, Ulnar; Ovary, Left; Pancreas; Parathyroid; Pituitary; Rectum; Skin/Subcutis; Spinal Cord; Spinal Nerve Roots, Cauda Equina; Spleen; Sternum; Stomach; Thymus; Thyroid; Tongue; Trachea; Urinary Bladder; Uterus; Vagina | | | | |
| The following tissues were examined microscopically and were unremarkable: Adrenal, Cortex; Aorta; Cecum; Duodenum; Femur; GALT/Peyer's Patch; Lung; Marrow, Femur; Marrow, Sternum; Muscle, Biceps Femoris; Nerve, Radial; Nerve, Ulnar; Spleen; Sternum; Stomach; Thymus | | | | |
| ‑‑‑‑‑‑‑‑‑‑‑‑‑‑‑‑‑‑‑‑‑‑‑‑‑‑‑‑‑‑‑‑‑‑‑‑‑‑‑‑‑‑‑‑‑‑‑‑‑‑‑‑‑‑‑‑‑‑‑‑‑‑‑‑‑‑‑‑‑‑‑‑‑‑‑‑‑‑‑‑‑‑‑‑ | | | | |
| **Animal Number: P0402 Group/Subgroup: 5/1 Sex: F Fate Status: Terminal Sacrifice**  **Date of Fate: 12 Oct 20 Phase of Fate: Dosing Phase Wk/Day of Fate: 4/28 TBW(g): 1500.0** | | | | |
| Organ Name: None | | | | |
| Macroscopic Observation(s)  Colon: Discolored; mucosa; multiple, up to 2 mm2; red; collected Brain: Infiltrate, mononuclear cell; minimal; | | | | Microscopic Observation(s)  Brain: Infiltrate, mononuclear cell; minimal; meninges/slide 48, 51  Eye, Left: Infiltrate, mononuclear cell; minimal; focal; ciliary body  Ganglion, Dorsal Root, Sacral: Autophagy; minimal  Ganglion, Dorsal Root, Sacral: Degeneration, axon, dorsal nerve root/spinal nerve; minimal; focal  Ganglion, Dorsal Root, Sacral: Infiltrate, mononuclear cell; minimal  Ganglion, Dorsal Root, Sacral: Satellite glial cell, increased/neuronal cell loss; minimal; focal  Ganglion, Lumbar Dorsal Root: Autophagy; minimal  Ganglion, Lumbar Dorsal Root: Degeneration/necrosis, neuron; minimal  Ganglion, Lumbar Dorsal Root: Inflammation, mononuclear cell, neuron; minimal  Ganglion, Lumbar Dorsal Root: Satellite glial cell,  increased/neuronal cell loss; minimal  Ganglion, Superior Cervical: BOTH MISSING  Ganglion, Thoracic Dorsal Root: Autophagy; minimal  Heart: Inflammation, mononuclear cell; slight  Intrathecal Injection Site, Cisterna Magna: Gliosis, grey matter; minimal; multifocal  Intrathecal Injection Site, Cisterna Magna: Infiltrate, mononuclear cell; minimal; meninges  Intrathecal Injection Site, Sacral Spinal Cord: MISSING  Intrathecal Injection Site, Sacral Spinal Cord: TISSUE COMMENT: Not available per histo comment  Muscle, Biceps Femoris: Infiltrate, mononuclear cell; minimal  Ovary, Left: Immature; Present  Pancreas: Infiltrate, mononuclear cell, endocrine; minimal  Pancreas: Infiltrate, mononuclear cell, exocrine; slight  Parathyroid: Infiltrate, mononuclear cell; slight  Spinal Cord, Lumbar: Gliosis, white matter; minimal; multifocal |
| The following tissues were examined macroscopically and were unremarkable: Adrenal; Animal; Aorta; Brain; Cecum; Cervix; Duodenum; Esophagus; Eye, Left; Femur; GALT/Peyer's Patch; Gall Bladder; Ganglion, Cervical Dorsal Root; Ganglion, Dorsal Root, Sacral; Ganglion, Lumbar Dorsal Root; Ganglion, Superior Cervical; Ganglion, Thoracic Dorsal Root; Heart; Ileum; Intrathecal Injection Site, Cisterna Magna; Intrathecal Injection Site, Sacral Spinal Cord; Jejunum; Kidney; Liver; Lung; Lymph Node, Mandibular; Lymph Node, Mesenteric; Mammary Gland; Mandibular Salivary Gland; Marrow, Femur; Marrow, Sternum; Muscle, Biceps Femoris; Nerve, Optic, Left; Nerve, Radial; Nerve, Sciatic; Nerve, Sural; Nerve, Tibial; Nerve, Ulnar; Ovary, Left; Pancreas; Parathyroid; Pituitary; Rectum; Skin/Subcutis; Spinal Cord; Spinal Nerve Roots, Cauda Equina; Spleen; Sternum; Stomach; Thymus; Thyroid; Tongue; Trachea; Urinary Bladder; Uterus; Vagina | | | | |
| The following tissues were examined microscopically and were unremarkable: Adrenal, Cortex; Adrenal, Medulla; Aorta; Cecum; Colon; Duodenum; Femur; GALT/Peyer's Patch; Ganglion, Cervical Dorsal Root; Kidney; Liver; Lung; Lymph Node, Mandibular; Marrow, Femur; Marrow, Sternum; Nerve, Radial; Nerve, Sciatic; Nerve, Sural; Nerve, Tibial; Nerve, Ulnar; Spinal Cord, Cervical; Spinal Cord, Thoracic; Spinal Nerve Roots, Cauda Equina; Spleen; Sternum; Stomach; Thymus; Thyroid | | | | |
| ‑‑‑‑‑‑‑‑‑‑‑‑‑‑‑‑‑‑‑‑‑‑‑‑‑‑‑‑‑‑‑‑‑‑‑‑‑‑‑‑‑‑‑‑‑‑‑‑‑‑‑‑‑‑‑‑‑‑‑‑‑‑‑‑‑‑‑‑‑‑‑‑‑‑‑‑‑‑‑‑‑‑‑‑ | | | | |
| **Animal Number: P0403 Group/Subgroup: 5/2 Sex: F Fate Status: Terminal Sacrifice**  **Date of Fate: 15 Oct 20 Phase of Fate: Dosing Phase Wk/Day of Fate: 4/28 TBW(g): 1700.0** | | | | |
| Organ Name: None | | | | |
| Macroscopic Observation(s)  Cecum: Discolored; mucosa; single, up to 5 mm2; red; collected/present on ileocecal junction  Colon: Discolored; mucosa; multiple, pinpoint; red; collected | | | | Microscopic Observation(s)  Brain: Infiltrate, mononuclear cell; minimal; meninges/slide 48, 49  Cecum: Congestion/hemorrhage, agonal; minimal  Colon: Degeneration, ganglia; slight; focal  Colon: Degeneration, tunica muscularis; moderate; focal  Colon: TISSUE COMMENT: The muscle and ganglion degeneration are associated with each other.  Ganglion, Lumbar Dorsal Root: Infiltrate, mononuclear cell; minimal  Ganglion, Superior Cervical: Infiltrate, mononuclear cell; minimal  Heart: Infiltrate, mononuclear cell; minimal  Intrathecal Injection Site, Sacral Spinal Cord: MISSING  Intrathecal Injection Site, Sacral Spinal Cord: TISSUE COMMENT: Not available per histo comment  Liver: Infiltrate, mononuclear cell; minimal; portal/random  Ovary, Left: Immature; Present  Pancreas: Infiltrate, mononuclear cell, exocrine; minimal  Spinal Cord, Lumbar: Degeneration, axon, funiculus; minimal; focal; dorsal funiculus white matter  Spinal Cord, Thoracic: Gliosis, white matter; minimal; focal  Thyroid: Ectopic tissue, thymus; Present |
| The following tissues were examined macroscopically and were unremarkable: Adrenal; Animal; Aorta; Brain; Cervix; Duodenum; Esophagus; Eye, Left; Femur; GALT/Peyer's Patch; Gall Bladder; Ganglion, Cervical Dorsal Root; Ganglion, Dorsal Root, Sacral; Ganglion, Lumbar Dorsal Root; Ganglion, Superior Cervical; Ganglion, Thoracic Dorsal Root; Heart; Ileum; Intrathecal Injection Site, Cisterna Magna; Intrathecal Injection Site, Sacral Spinal Cord; Jejunum; Kidney; Liver; Lung; Lymph Node, Mandibular; Lymph Node, Mesenteric; Mammary Gland; Mandibular Salivary Gland; Marrow, Femur; Marrow, Sternum; Muscle, Biceps Femoris; Nerve, Optic, Left; Nerve, Radial; Nerve, Sciatic; Nerve, Sural; Nerve, Tibial; Nerve, Ulnar; Ovary, Left; Pancreas; Parathyroid; Pituitary; Rectum; Skin/Subcutis; Spinal Cord; Spinal Nerve Roots, Cauda Equina; Spleen; Sternum; Stomach; Thymus; Thyroid; Tongue; Trachea; Urinary Bladder; Uterus; Vagina | | | | |
| The following tissues were examined microscopically and were unremarkable: Adrenal, Cortex; Adrenal, Medulla; Aorta; Duodenum; Eye, Left; Femur; GALT/Peyer's Patch; Ganglion, Cervical Dorsal Root; Ganglion, Dorsal Root, Sacral; Ganglion, Thoracic Dorsal Root; Intrathecal Injection Site, Cisterna Magna; Kidney; Lung; Lymph Node, Mandibular; Marrow, Femur; Marrow, Sternum; Muscle, Biceps Femoris; Nerve, Radial; Nerve, Sciatic; Nerve, Sural; Nerve, Tibial; Nerve, Ulnar; Parathyroid; Spinal Cord, Cervical; Spinal Nerve Roots, Cauda Equina; Spleen; Sternum; Stomach; Thymus | | | | |
| ‑‑‑‑‑‑‑‑‑‑‑‑‑‑‑‑‑‑‑‑‑‑‑‑‑‑‑‑‑‑‑‑‑‑‑‑‑‑‑‑‑‑‑‑‑‑‑‑‑‑‑‑‑‑‑‑‑‑‑‑‑‑‑‑‑‑‑‑‑‑‑‑‑‑‑‑‑‑‑‑‑‑‑‑ | | | | |
| **Animal Number: P0404 Group/Subgroup: 5/2 Sex: F Fate Status: Terminal Sacrifice**  **Date of Fate: 15 Oct 20 Phase of Fate: Dosing Phase Wk/Day of Fate: 4/28 TBW(g): 1600.0** | | | | |
| Organ Name: None | | | | |
| Macroscopic Observation(s)  Muscle, Skeletal, Other: Thickened; entire; collected/Proximal portion of the semispinalis capitis towards the head collected. | | | | Microscopic Observation(s)  Adipose, Brown: Infiltrate, mononuclear cell; minimal; periaorta/slide 9A  Brain: Degeneration, axon; minimal; focal/slide 51  Brain: Gliosis, white matter; minimal; multifocal/slide 49, 51, 53  Brain: Infiltrate, mononuclear cell; minimal; perivascular, neuropil/choroid plexus/meninges: slide 46, 47 neuropil: 45, 49, 51, 53  Ganglion, Cervical Dorsal Root: Degeneration/necrosis, neuron; minimal  Ganglion, Cervical Dorsal Root: Infiltrate, mononuclear cell; minimal  Ganglion, Cervical Dorsal Root: Inflammation, mononuclear cell, neuron; minimal  Ganglion, Dorsal Root, Sacral: Degeneration, axon, dorsal nerve root/spinal nerve; minimal  Ganglion, Dorsal Root, Sacral: Infiltrate, mononuclear cell; moderate  Ganglion, Dorsal Root, Sacral: Infiltrate, mononuclear cell, dorsal nerve root/spinal nerve; minimal  Ganglion, Dorsal Root, Sacral: Inflammation, mononuclear cell, dorsal nerve root/spinal nerve; slight  Ganglion, Lumbar Dorsal Root: Degeneration/necrosis, neuron; slight  Ganglion, Lumbar Dorsal Root: Infiltrate, mononuclear cell; slight  Ganglion, Lumbar Dorsal Root: Infiltrate, mononuclear cell, dorsal nerve root/spinal nerve; minimal  Ganglion, Lumbar Dorsal Root: Inflammation, mononuclear cell, neuron; slight  Ganglion, Superior Cervical: Infiltrate, mononuclear cell; minimal  Ganglion, Thoracic Dorsal Root: Degeneration/necrosis, neuron; minimal; focal  Ganglion, Thoracic Dorsal Root: Infiltrate, mononuclear cell; minimal  Ganglion, Thoracic Dorsal Root: Inflammation, mononuclear cell, neuron; minimal; focal  Heart: Infiltrate, mononuclear cell; minimal  Intrathecal Injection Site, Cisterna Magna: Degeneration; moderate; dorsal, lateral  Intrathecal Injection Site, Cisterna Magna: Gliosis, grey matter; minimal; focal  Intrathecal Injection Site, Cisterna Magna: Gliosis, white matter; minimal; focal  Intrathecal Injection Site, Sacral Spinal Cord: Degeneration, axon, nerve roots/cauda equina; minimal  Liver: Infiltrate, mononuclear cell; slight; portal/random/To include some centrilobular areas.  Lung: Infiltrate, mononuclear cell; minimal; perivascular/terminal bronchioles  Lymph Node, Mandibular: Lymphocytes, increased; slight; follicle, unilateral  Muscle, Biceps Femoris: Infiltrate, mononuclear cell; minimal  Muscle, Skeletal, Other: Degeneration, myofiber; moderate  Muscle, Skeletal, Other: Inflammation granulomatous; moderate  Muscle, Skeletal, Other: Regeneration, myofiber; slight  Muscle, Skeletal, Other: TISSUE COMMENT: Findings are on slide 54  Nerve, Sciatic: Degeneration, axon; slight  Nerve, Sural: Degeneration, axon; minimal  Nerve, Tibial: Degeneration, axon; slight  Nerve, Ulnar: Degeneration, axon; slight  Ovary, Left: Cyst; Present  Ovary, Left: Immature; Present  Pancreas: Infiltrate, mononuclear cell, endocrine; minimal  Pancreas: Infiltrate, mononuclear cell, exocrine; minimal  Parathyroid: Infiltrate, mononuclear cell; slight  Spinal Cord, Cervical: Degeneration, axon, funiculus; moderate; dorsal, lateral/Primarily a lateral orientation.  Spinal Cord, Lumbar: Degeneration, axon, funiculus; slight; dorsal, lateral  Spinal Cord, Lumbar: Degeneration, axon, nerve root; slight  Spinal Cord, Lumbar: Gliosis, white matter; minimal; focal  Spinal Cord, Lumbar: Infiltrate, mononuclear cell; slight; meninges  Spinal Cord, Lumbar: Infiltrate, mononuclear cell, nerve roots; minimal; focal  Spinal Cord, Thoracic: Degeneration, axon, funiculus; moderate; dorsal and lateral  Spinal Cord, Thoracic: Gliosis, white matter; minimal; focal  Spinal Nerve Roots, Cauda Equina: Infiltrate, mononuclear cell; minimal  Stomach: Degeneration/necrosis, vessel; minimal; focal/slide 20W |
| The following tissues were examined macroscopically and were unremarkable: Adrenal; Animal; Aorta; Brain; Cecum; Cervix; Colon; Duodenum; Esophagus; Eye, Left; Femur; GALT/Peyer's Patch; Gall Bladder; Ganglion, Cervical Dorsal Root; Ganglion, Dorsal Root, Sacral; Ganglion, Lumbar Dorsal Root; Ganglion, Superior Cervical; Ganglion, Thoracic Dorsal Root; Heart; Ileum; Intrathecal Injection Site, Cisterna Magna; Intrathecal Injection Site, Sacral Spinal Cord; Jejunum; Kidney; Liver; Lung; Lymph Node, Mandibular; Lymph Node, Mesenteric; Mammary Gland; Mandibular Salivary Gland; Marrow, Femur; Marrow, Sternum; Muscle, Biceps Femoris; Nerve, Optic, Left; Nerve, Radial; Nerve, Sciatic; Nerve, Sural; Nerve, Tibial; Nerve, Ulnar; Ovary, Left; Pancreas; Parathyroid; Pituitary; Rectum; Skin/Subcutis; Spinal Cord; Spinal Nerve Roots, Cauda Equina; Spleen; Sternum; Stomach; Thymus; Thyroid; Tongue; Trachea; Urinary Bladder; Uterus; Vagina | | | | |
| The following tissues were examined microscopically and were unremarkable: Adrenal, Cortex; Adrenal, Medulla; Aorta; Cecum; Duodenum; Eye, Left; Femur; GALT/Peyer's Patch; Kidney; Marrow, Femur; Marrow, Sternum; Nerve, Radial; Spleen; Sternum; Thymus; Thyroid | | | | |
| ‑‑‑‑‑‑‑‑‑‑‑‑‑‑‑‑‑‑‑‑‑‑‑‑‑‑‑‑‑‑‑‑‑‑‑‑‑‑‑‑‑‑‑‑‑‑‑‑‑‑‑‑‑‑‑‑‑‑‑‑‑‑‑‑‑‑‑‑‑‑‑‑‑‑‑‑‑‑‑‑‑‑‑‑ | | | | |
| **Animal Number: P0501 Group/Subgroup: 6/1 Sex: F Fate Status: Terminal Sacrifice**  **Date of Fate: 12 Oct 20 Phase of Fate: Dosing Phase Wk/Day of Fate: 4/28 TBW(g): 1500.0** | | | | |
| Organ Name: None | | | | |
| Macroscopic Observation(s)  Skin/Subcutis: Sore; limb; right hind; single, up to 5 mm2; red; collected | | | | Microscopic Observation(s)  Brain: Gliosis, grey matter; slight; multifocal/slide 45, 46 (extending down from meninges), 47  Brain: Gliosis, white matter; slight; multifocal/slide 44, 45, 47, 49  Brain: Infiltrate, mononuclear cell; slight; perivascular, meninges, neuropil, choroid plexus/meninges: slide 44, 45, 47, 48. 49, 52choroid plexus: slide 46, 51neuropil: 50  Cecum: Parasite, nematode; Present  Ganglion, Cervical Dorsal Root: Degeneration, axon, dorsal nerve root/spinal nerve; minimal  Ganglion, Cervical Dorsal Root: Degeneration/necrosis, neuron; minimal  Ganglion, Cervical Dorsal Root: Infiltrate, mononuclear cell; minimal  Ganglion, Cervical Dorsal Root: Infiltrate, mononuclear cell, dorsal nerve root/spinal nerve; minimal  Ganglion, Cervical Dorsal Root: Inflammation, mononuclear cell, neuron; minimal  Ganglion, Cervical Dorsal Root: Satellite glial cell, increased/neuronal cell loss; minimal  Ganglion, Dorsal Root, Sacral: Autophagy; minimal  Ganglion, Dorsal Root, Sacral: Degeneration, axon, dorsal nerve root/spinal nerve; minimal  Ganglion, Dorsal Root, Sacral: Degeneration/necrosis, neuron; minimal  Ganglion, Dorsal Root, Sacral: Infiltrate, mononuclear cell; slight  Ganglion, Dorsal Root, Sacral: Infiltrate, mononuclear cell, dorsal nerve root/spinal nerve; minimal  Ganglion, Dorsal Root, Sacral: Inflammation, mononuclear cell, neuron; minimal  Ganglion, Lumbar Dorsal Root: Autophagy; minimal  Ganglion, Lumbar Dorsal Root: Infiltrate, mononuclear cell; slight  Ganglion, Superior Cervical: Infiltrate, mononuclear cell; minimal  Ganglion, Superior Cervical: Vacuolation, neuron; minimal  Ganglion, Thoracic Dorsal Root: Autophagy; minimal  Ganglion, Thoracic Dorsal Root: Infiltrate, mononuclear cell; minimal  Heart: Inflammation, mononuclear cell; minimal  Intrathecal Injection Site, Cisterna Magna: Degeneration; slight; dorsal, lateral, ventral  Intrathecal Injection Site, Cisterna Magna: Gliosis, grey matter; slight; multifocal  Intrathecal Injection Site, Cisterna Magna: Gliosis, white matter; minimal; multifocal  Intrathecal Injection Site, Sacral Spinal Cord: MISSING  Intrathecal Injection Site, Sacral Spinal Cord: TISSUE COMMENT: Not available per histo comment  Kidney: Glomerulonephritis; moderate  Liver: Infiltrate, mononuclear cell; slight; portal  Nerve, Sciatic: Degeneration, axon; minimal  Nerve, Sciatic: Infiltrate, mononuclear cell; minimal  Nerve, Sural: BOTH MISSING  Ovary, Left: Immature; Present  Pancreas: Infiltrate, mononuclear cell, endocrine; minimal  Pancreas: Infiltrate, mononuclear cell, exocrine; slight  Spinal Cord, Cervical: Degeneration, axon, funiculus; slight; dorsal, lateral, & ventral  Spinal Cord, Cervical: Gliosis, grey matter; slight  Spinal Cord, Cervical: Gliosis, white matter; minimal  Spinal Cord, Lumbar: Degeneration, axon, funiculus; slight; dorsal, lateral, and ventral  Spinal Cord, Lumbar: Gliosis, grey matter; slight; multifocal  Spinal Cord, Lumbar: Infiltrate, mononuclear cell; minimal; meninges  Spinal Cord, Thoracic: Degeneration, axon, funiculus; slight; lateral, ventral  Spinal Cord, Thoracic: Gliosis, white matter; minimal; multifocal |
| The following tissues were examined macroscopically and were unremarkable: Adrenal; Animal; Aorta; Brain; Cecum; Cervix; Colon; Duodenum; Esophagus; Eye, Left; Femur; GALT/Peyer's Patch; Gall Bladder; Ganglion, Cervical Dorsal Root; Ganglion, Dorsal Root, Sacral; Ganglion, Lumbar Dorsal Root; Ganglion, Superior Cervical; Ganglion, Thoracic Dorsal Root; Heart; Ileum; Intrathecal Injection Site, Cisterna Magna; Intrathecal Injection Site, Sacral Spinal Cord; Jejunum; Kidney; Liver; Lung; Lymph Node, Mandibular; Lymph Node, Mesenteric; Mammary Gland; Mandibular Salivary Gland; Marrow, Femur; Marrow, Sternum; Muscle, Biceps Femoris; Nerve, Optic, Left; Nerve, Radial; Nerve, Sciatic; Nerve, Sural; Nerve, Tibial; Nerve, Ulnar; Ovary, Left; Pancreas; Parathyroid; Pituitary; Rectum; Spinal Cord; Spinal Nerve Roots, Cauda Equina; Spleen; Sternum; Stomach; Thymus; Thyroid; Tongue; Trachea; Urinary Bladder; Uterus; Vagina | | | | |
| The following tissues were examined microscopically and were unremarkable: Adrenal, Cortex; Adrenal, Medulla; Aorta; Duodenum; Eye, Left; Femur; GALT/Peyer's Patch; Lung; Lymph Node, Mandibular; Marrow, Femur; Marrow, Sternum; Muscle, Biceps Femoris; Nerve, Radial; Nerve, Tibial; Nerve, Ulnar; Parathyroid; Skin/Subcutis; Spinal Nerve Roots, Cauda Equina; Spleen; Sternum; Stomach; Thymus; Thyroid | | | | |
| ‑‑‑‑‑‑‑‑‑‑‑‑‑‑‑‑‑‑‑‑‑‑‑‑‑‑‑‑‑‑‑‑‑‑‑‑‑‑‑‑‑‑‑‑‑‑‑‑‑‑‑‑‑‑‑‑‑‑‑‑‑‑‑‑‑‑‑‑‑‑‑‑‑‑‑‑‑‑‑‑‑‑‑‑ | | | | |
| **Animal Number: P0502 Group/Subgroup: 6/1 Sex: F Fate Status: Terminal Sacrifice**  **Date of Fate: 12 Oct 20 Phase of Fate: Dosing Phase Wk/Day of Fate: 4/28 TBW(g): 1500.0** | | | | |
| Organ Name: None | | | | |
| Macroscopic Observation(s)  None | | | | Microscopic Observation(s)  Brain: Infiltrate, mononuclear cell; minimal; meninges/slide 47  Ganglion, Cervical Dorsal Root: Degeneration/necrosis, neuron; minimal  Ganglion, Cervical Dorsal Root: Infiltrate, mononuclear cell; slight  Ganglion, Cervical Dorsal Root: Inflammation, mononuclear cell, neuron; minimal  Ganglion, Dorsal Root, Sacral: Degeneration, axon, dorsal nerve root/spinal nerve; slight  Ganglion, Dorsal Root, Sacral: Degeneration/necrosis, neuron; minimal  Ganglion, Dorsal Root, Sacral: Infiltrate, mononuclear cell; moderate  Ganglion, Dorsal Root, Sacral: Infiltrate, mononuclear cell, dorsal nerve root/spinal nerve; minimal  Ganglion, Dorsal Root, Sacral: Inflammation, mononuclear cell, neuron; minimal  Ganglion, Dorsal Root, Sacral: TISSUE COMMENT: Also see slide 41.  Ganglion, Lumbar Dorsal Root: Degeneration/necrosis, neuron; slight  Ganglion, Lumbar Dorsal Root: Infiltrate, mononuclear cell; slight  Ganglion, Lumbar Dorsal Root: Infiltrate, mononuclear cell, dorsal nerve root/spinal nerve; minimal  Ganglion, Lumbar Dorsal Root: Inflammation, mononuclear cell, neuron; slight  Ganglion, Thoracic Dorsal Root: Infiltrate, mononuclear cell; minimal; ganglia, adventitia  Heart: Inflammation, mononuclear cell; slight  Intrathecal Injection Site, Cisterna Magna: Degeneration; minimal; focal; lateral  Intrathecal Injection Site, Cisterna Magna: Degeneration, neuron; minimal; multifocal  Intrathecal Injection Site, Cisterna Magna:G liosis, grey matter; slight; multifocal/Often present around degenerative neuron.  Intrathecal Injection Site, Sacral Spinal Cord: MISSING  Intrathecal Injection Site, Sacral Spinal Cord: TISSUE COMMENT: Not available per histo comment  Liver: Infiltrate, mononuclear cell; slight; portal/A rare number of affected sites have a mixed cell population.  Lymph Node, Mandibular: Lymphocytes, increased; minimal; unilateral  Nerve, Sciatic: Infiltrate, mononuclear cell; minimal  Ovary, Left: Immature; Present  Pancreas: Infiltrate, mononuclear cell, endocrine; minimal  Pancreas: Infiltrate, mononuclear cell, exocrine; slight  Spinal Cord, Lumbar: Degeneration, axon, funiculus; minimal; dorsal funiculus white matter  Spinal Cord, Lumbar: Gliosis, white matter; minimal; focal  Spinal Cord, Thoracic: Gliosis, white matter; minimal; focal  Thyroid: Infiltrate, mononuclear cell; minimal; focal; unilateral |
| The following tissues were examined macroscopically and were unremarkable: Adrenal; Animal; Aorta; Brain; Cecum; Cervix; Colon; Duodenum; Esophagus; Eye, Left; Femur; GALT/Peyer's Patch; Gall Bladder; Ganglion, Cervical Dorsal Root; Ganglion, Dorsal Root, Sacral; Ganglion, Lumbar Dorsal Root; Ganglion, Superior Cervical; Ganglion, Thoracic Dorsal Root; Heart; Ileum; Intrathecal Injection Site, Cisterna Magna; Intrathecal Injection Site, Sacral Spinal Cord; Jejunum; Kidney; Liver; Lung; Lymph Node, Mandibular; Lymph Node, Mesenteric; Mammary Gland; Mandibular Salivary Gland; Marrow, Femur; Marrow, Sternum; Muscle, Biceps Femoris; Nerve, Optic, Left; Nerve, Radial; Nerve, Sciatic; Nerve, Sural; Nerve, Tibial; Nerve, Ulnar; Ovary, Left; Pancreas; Parathyroid; Pituitary; Rectum; Skin/Subcutis; Spinal Cord; Spinal Nerve Roots, Cauda Equina; Spleen; Sternum; Stomach; Thymus; Thyroid; Tongue; Trachea; Urinary Bladder; Uterus; Vagina | | | | |
| The following tissues were examined microscopically and were unremarkable: Adrenal, Cortex; Adrenal, Medulla; Aorta; Cecum; Duodenum; Eye, Left; Femur; GALT/Peyer's Patch; Ganglion, Superior Cervical; Kidney; Lung; Marrow, Femur; Marrow, Sternum; Muscle, Biceps Femoris; Nerve, Radial; Nerve, Sural; Nerve, Tibial; Nerve, Ulnar; Parathyroid; Spinal Cord, Cervical; Spinal Nerve Roots, Cauda Equina; Spleen; Sternum; Stomach; Thymus | | | | |
| ‑‑‑‑‑‑‑‑‑‑‑‑‑‑‑‑‑‑‑‑‑‑‑‑‑‑‑‑‑‑‑‑‑‑‑‑‑‑‑‑‑‑‑‑‑‑‑‑‑‑‑‑‑‑‑‑‑‑‑‑‑‑‑‑‑‑‑‑‑‑‑‑‑‑‑‑‑‑‑‑‑‑‑‑ | | | | |
| **Animal Number: P0503 Group/Subgroup: 6/2 Sex: F Fate Status: Terminal Sacrifice**  **Date of Fate: 15 Oct 20 Phase of Fate: Dosing Phase Wk/Day of Fate: 4/28 TBW(g): 1600.0** | | | | |
| Organ Name: None | | | | |
| Macroscopic Observation(s)  Cecum: Discolored; mucosa; single, up to 5 mm2; red; collected/present on ileocecal junction  Colon: Discolored; mucosa; multiple, pinpoint; red; collected | | | Microscopic Observation(s)  Adipose, Brown: Infiltrate, mononuclear cell; slight; pericardial/periaorta/Severity based on pericardial portion of the finding.  Brain: Infiltrate, mononuclear cell; minimal; meninges/meninges: slide 45, 48  Cecum: Congestion/hemorrhage, agonal; minimal  Ganglion, Cervical Dorsal Root: Degeneration/necrosis, neuron; minimal  Ganglion, Cervical Dorsal Root: Infiltrate, mononuclear cell; minimal  Ganglion, Cervical Dorsal Root: Inflammation, mononuclear cell, neuron; minimal  Ganglion, Dorsal Root, Sacral: Degeneration, axon, dorsal nerve root/spinal nerve; moderate  Ganglion, Dorsal Root, Sacral: Degeneration/necrosis, neuron; slight  Ganglion, Dorsal Root, Sacral: Infiltrate, mononuclear cell; moderate  Ganglion, Dorsal Root, Sacral: Inflammation, mononuclear cell, dorsal nerve root/spinal nerve; moderate  Ganglion, Dorsal Root, Sacral: Inflammation, mononuclear cell, neuron; slight  Ganglion, Lumbar Dorsal Root: Degeneration/necrosis, neuron; minimal  Ganglion, Lumbar Dorsal Root: Infiltrate, mononuclear cell; slight  Ganglion, Lumbar Dorsal Root: Infiltrate, mononuclear cell, dorsal nerve root/spinal nerve; minimal  Ganglion, Lumbar Dorsal Root: Inflammation, mononuclear cell, neuron; minimal  Ganglion, Superior Cervical: Infiltrate, mononuclear cell; minimal  Ganglion, Thoracic Dorsal Root: Degeneration/necrosis, neuron; minimal  Ganglion, Thoracic Dorsal Root: Infiltrate, mononuclear cell; minimal  Ganglion, Thoracic Dorsal Root: Inflammation, mononuclear cell, neuron; minimal  Heart: Inflammation, mononuclear cell; slight  Intrathecal Injection Site, Cisterna Magna: Degeneration; slight; dorsal, lateral, ventral  Intrathecal Injection Site, Sacral Spinal Cord:  Degeneration, axon, nerve roots/cauda equina; moderate  Liver: Infiltrate, mononuclear cell; slight; portal/random  Muscle, Biceps Femoris: Infiltrate, mononuclear cell; minimal  Nerve, Sciatic: Degeneration, axon; minimal  Nerve, Sciatic: Infiltrate, mononuclear cell; minimal  Nerve, Sural: Degeneration, axon; slight  Nerve, Tibial: Degeneration, axon; moderate  Ovary, Left: Immature; Present  Pancreas: Degeneration; moderate  Pancreas: Inflammation, mononuclear cell, endocrine; slight  Pancreas: Inflammation, mononuclear cell, exocrine; moderate  Parathyroid: Infiltrate, mononuclear cell; minimal  Spinal Cord, Cervical: Degeneration, axon, funiculus; slight; dorsal, lateral, & ventral  Spinal Cord, Cervical: Gliosis, white matter; minimal; focal  Spinal Cord, Lumbar: Degeneration, axon, funiculus; slight; dorsal, lateral, and ventral  Spinal Cord, Lumbar: Degeneration, axon, nerve root; slight  Spinal Nerve Roots, Cauda Equina: Degeneration, axon; moderate  Spinal Nerve Roots, Cauda Equina: Gliosis; slight | |
| The following tissues were examined macroscopically and were unremarkable: Adrenal; Animal; Aorta; Brain; Cervix; Duodenum; Esophagus; Eye, Left; Femur; GALT/Peyer's Patch; Gall Bladder; Ganglion, Cervical Dorsal Root; Ganglion, Dorsal Root, Sacral; Ganglion, Lumbar Dorsal Root; Ganglion, Superior Cervical; Ganglion, Thoracic Dorsal Root; Heart; Ileum; Intrathecal Injection Site, Cisterna Magna; Intrathecal Injection Site, Sacral Spinal Cord; Jejunum; Kidney; Liver; Lung; Lymph Node, Mandibular; Lymph Node, Mesenteric; Mammary Gland; Mandibular Salivary Gland; Marrow, Femur; Marrow, Sternum; Muscle, Biceps Femoris; Nerve, Optic, Left; Nerve, Radial; Nerve, Sciatic; Nerve, Sural; Nerve, Tibial; Nerve, Ulnar; Ovary, Left; Pancreas; Parathyroid; Pituitary; Rectum; Skin/Subcutis; Spinal Cord; Spinal Nerve Roots, Cauda Equina; Spleen; Sternum; Stomach; Thymus; Thyroid; Tongue; Trachea; Urinary Bladder; Uterus; Vagina | | | | |
| The following tissues were examined microscopically and were unremarkable: Adrenal, Cortex; Adrenal, Medulla; Aorta; Colon; Duodenum; Eye, Left; Femur; GALT/Peyer's Patch; Kidney; Lung; Lymph Node, Mandibular; Marrow, Femur; Marrow, Sternum; Nerve, Radial; Nerve, Ulnar; Spinal Cord, Thoracic; Spleen; Sternum; Stomach; Thymus; Thyroid | | | | |
| ‑‑‑‑‑‑‑‑‑‑‑‑‑‑‑‑‑‑‑‑‑‑‑‑‑‑‑‑‑‑‑‑‑‑‑‑‑‑‑‑‑‑‑‑‑‑‑‑‑‑‑‑‑‑‑‑‑‑‑‑‑‑‑‑‑‑‑‑‑‑‑‑‑‑‑‑‑‑‑‑‑‑‑‑‑‑‑‑‑‑‑‑‑‑‑‑‑‑‑‑‑‑‑‑‑‑ | | | | |
| **Animal Number: P0504 Group/Subgroup: 6/2 Sex: F Fate Status: Terminal Sacrifice**  **Date of Fate: 15 Oct 20 Phase of Fate: Dosing Phase Wk/Day of Fate: 4/28 TBW(g): 1700.0** | | | | |
| Organ Name: None | | | | |
| Macroscopic Observation(s)  Cecum: Discolored; mucosa; entire; red; collected/Ileocecal junction  Muscle, Skeletal, Other: Thickened; entire; collected/Proximal portion of semispinalis capitis toward the head collected | | | | Microscopic Observation(s)  Adipose, Brown: Infiltrate, mononuclear cell; slight; periaorta/slide 9A  Brain: Gliosis, white matter; minimal; focal/slide 49  Brain: Infiltrate, mixed cell, white matter; minimal; focal/slide 46  Cecum: Congestion/hemorrhage, agonal; slight  Cecum: Inflammation, vessel; slight; multifocal  Cecum: Parasite, nematode; Present  Duodenum: Inflammation, vessel; slight; focal  Ganglion, Cervical Dorsal Root: Degeneration/necrosis, neuron; minimal  Ganglion, Cervical Dorsal Root: Inflammation, mononuclear cell, neuron; minimal  Ganglion, Dorsal Root, Sacral: Degeneration, axon, dorsal nerve root/spinal nerve; minimal  Ganglion, Dorsal Root, Sacral: Degeneration/necrosis, neuron; slight  Ganglion, Dorsal Root, Sacral: Infiltrate, mononuclear cell; slight  Ganglion, Dorsal Root, Sacral: Infiltrate, mononuclear cell, dorsal nerve root/spinal nerve; slight  Ganglion, Dorsal Root, Sacral: Inflammation, mononuclear cell, neuron; slight  Ganglion, Dorsal Root, Sacral: Vacuolation, neuron; minimal  Ganglion, Lumbar Dorsal Root: Autophagy; minimal  Ganglion, Lumbar Dorsal Root: Degeneration, axon, dorsal nerve root/spinal nerve; slight  Ganglion, Lumbar Dorsal Root: Degeneration/necrosis, neuron; slight  Ganglion, Lumbar Dorsal Root: Infiltrate, mononuclear cell, dorsal nerve root/spinal nerve; minimal  Ganglion, Lumbar Dorsal Root: Inflammation, mononuclear cell, dorsal nerve root/spinal nerve; slight  Ganglion, Lumbar Dorsal Root: Inflammation, mononuclear cell, neuron; slight  Ganglion, Superior Cervical: Infiltrate, mononuclear cell; minimal  Ganglion, Thoracic Dorsal Root: Degeneration/necrosis, neuron; minimal  Ganglion, Thoracic Dorsal Root: Infiltrate, mononuclear cell; minimal  Ganglion, Thoracic Dorsal Root: Inflammation, mononuclear cell, neuron; minimal  Intrathecal Injection Site, Cisterna Magna: Degeneration; moderate; dorsal, lateral/The finding is minimal in the dorsal funiculi, and moderate unilateral finding in the lateral funiculi (considering procedure related).  Intrathecal Injection Site, Sacral Spinal Cord: Degeneration, axon, nerve roots/cauda equina; slight  Kidney: Infiltrate, mononuclear cell; minimal  Liver: Infiltrate, mononuclear cell; slight; portal/centrilobular  Muscle, Biceps Femoris: Infiltrate, mononuclear cell; minimal  Muscle, Skeletal, Other: Degeneration, myofiber; marked  Muscle, Skeletal, Other: Inflammation, mononuclear cell; marked  Muscle, Skeletal, Other: Regeneration, myofiber; moderate  Muscle, Skeletal, Other: TISSUE COMMENT: slide 54  Nerve, Sciatic: Infiltrate, mononuclear cell; minimal  Nerve, Sural: Degeneration, axon; minimal  Nerve, Tibial: Degeneration, axon; minimal  Ovary, Left: Immature; Present  Pancreas: Ectopic tissue, spleen; Present  Pancreas: Infiltrate, mononuclear cell, exocrine; minimal  Parathyroid: Infiltrate, mononuclear cell; moderate  Spinal Cord, Cervical: Degeneration, axon, funiculus; moderate; dorsal, lateral, & ventral/The finding is minimal in the dorsal and ventral funiculi, and moderate unilateral finding in the lateral funiculi (considering procedure related).  Spinal Cord, Cervical: Gliosis, white matter; minimal; multifocal  Spinal Cord, Lumbar: Degeneration, axon, funiculus; moderate; dorsal, lateral, and ventral/The finding is minimal in the ventral, slight in the ventral funiculi, and moderate in the lateral funiculi primarily unilateral (considered procedure related).  Spinal Cord, Lumbar: Degeneration, axon, nerve root; slight  Spinal Cord, Lumbar: Gliosis, grey matter; minimal; focal  Spinal Cord, Lumbar: Gliosis, white matter; minimal; multifocal  Spinal Cord, Thoracic: Degeneration, axon, funiculus; moderate; dorsal and lateral/The finding is slight in the dorsal funiculi, and moderate unilateral finding in the lateral funiculi (considering procedure related).  Spinal Cord, Thoracic: Gliosis, white matter; minimal; focal  Spinal Nerve Roots, Cauda Equina: Degeneration, axon; slight  Spinal Nerve Roots, Cauda Equina: Gliosis; slight  Stomach: Inflammation, vessel; slight; multifocal  Thyroid: Infiltrate, mononuclear cell; minimal  Thyroid: Inflammation, mononuclear cell; slight; unilateral |
| The following tissues were examined macroscopically and were unremarkable: Adrenal; Animal; Aorta; Brain; Cervix; Colon; Duodenum; Esophagus; Eye, Left; Femur; GALT/Peyer's Patch; Gall Bladder; Ganglion, Cervical Dorsal Root; Ganglion, Dorsal Root, Sacral; Ganglion, Lumbar Dorsal Root; Ganglion, Superior Cervical; Ganglion, Thoracic Dorsal Root; Heart; Ileum; Intrathecal Injection Site, Cisterna Magna; Intrathecal Injection Site, Sacral Spinal Cord; Jejunum; Kidney; Liver; Lung; Lymph Node, Mandibular; Lymph Node, Mesenteric; Mammary Gland; Mandibular Salivary Gland; Marrow, Femur; Marrow, Sternum; Muscle, Biceps Femoris; Nerve, Optic, Left; Nerve, Radial; Nerve, Sciatic; Nerve, Sural; Nerve, Tibial; Nerve, Ulnar; Ovary, Left; Pancreas; Parathyroid; Pituitary; Rectum; Skin/Subcutis; Spinal Cord; Spinal Nerve Roots, Cauda Equina; Spleen; Sternum; Stomach; Thymus; Thyroid; Tongue; Trachea; Urinary Bladder; Uterus; Vagina | | | | |
| The following tissues were examined microscopically and were unremarkable: Adrenal, Cortex; Adrenal, Medulla; Aorta; Eye, Left; Femur; GALT/Peyer's Patch; Heart; Lung; Lymph Node, Mandibular; Marrow, Femur; Marrow, Sternum; Nerve, Radial; Nerve, Ulnar; Spleen; Sternum; Thymus | | | | |

**Supplemental Table 34. Green Fluorescent Protein Immunohistochemistry Scores on Spinal, DRG, and Brain Sections from a Control Animal and Animals with 3.0×10^13^ vg/animal of scAAV9-CB-GFP by the Intrathecal-LP and ICM Routes.**

|  | **Control** | **Intrathecal-LP (3.0×10^13^)** | | | | **ICM (3.0×10^13^)** | | | |
| --- | --- | --- | --- | --- | --- | --- | --- | --- | --- |
| Lumbar spinal cord | Neg | Moderate | Moderate | Moderate | Moderate | Moderate | Neg | Mild | Moderate |
| Sacral dorsal root ganglion | Neg | Moderate | Strong | Moderate | Mild | Minimal | Mild | Mild | Strong |
| Prefrontal cortex | Neg | Minimal | Minimal | Minimal | Mild | Minimal | Minimal | Minimal | Minimal |
| Temporal cortex, putamen, cingulate gyrus | Neg | Mild | Mild | Neg | Mild | Mild | Neg | Mild | Minimal |
| Corpus callosum, cingulate gyrus | Neg | Mild | Mild | Minimal | Mild | Minimal | Minimal | Mild | Minimal |
| Temporal cortex | Neg | Mild | Mild | Minimal | Mild | Minimal | Neg | Mild | Minimal |
| Thalamus, hypothalamus, hippocampus, amygdala | Neg | Mild | Mild | Minimal | Mild | Minimal | Minimal | Mild | Minimal |
| Thalamus, hypothalamus, hippocampus, substantia nigra | Neg | Mild | Mild | Minimal | Mild | Mild | n/p | Mild | Minimal |
| Pons, cerebellum | Neg | Mild | Mild | Minimal | Mild | Mild | Minimal | Mild | Moderate |
| Cerebellum deep cerebellar nuclei | Neg | Moderate | Mild | Minimal | Moderate | Mild | Minimal | Mild | Minimal |
| Occipital cortex | Neg | Moderate | Moderate | Minimal | Moderate | Mild | Minimal | Mild | Minimal |
| Cerebellum | Neg | Mild | n/p | Minimal | n/p | Mild | n/p | Mild | Minimal |

Abbreviations: DRG, dorsal root ganglion; ICM, intracisterna magna; LP, lumbar puncture; Neg, negative; n/p, not present; scAAV9-CB-GFP, self-complementary adeno-associated virus serotype 9–chicken β-actin promoter–green fluorescent protein; vg, vector genomes.

**Supplemental Table 35. Green Fluorescent Protein Expression in Systemic Tissue Types.**

| **Expression level** | **Tissue type** |
| --- | --- |
| High | Skeletal muscle myocytes, cardiomyocytes, liver hepatocytes, lymphoid follicular dendritic cells (spleen, lymph node, and GALT/BALT), renal juxtaglomerular apparatus |
| Medium | Brown adipocytes, parathyroid gland, adrenal cortical cells, exocrine pancreas, endocrine pancreas, renal medullary cells |
| Low | Thyroid gland, stomach |
| Minimal | Lung, aorta, adrenal medullary cells, esophageal mucosa, small intestine, large intestine, thymus, lymph node non-follicular cortex and medulla, hepatic portal and sinusoidal cells, white adipocytes, bone marrow, intraocular structures, renal cortical cells, renal glomerulus |

Abbreviations: BALT, bronchus-associated lymphoid tissue; GALT, gut-associated lymphoid tissue.

**SUPPLEMENTAL FIGURES**

**Supplemental Figure 1. Biodistribution of scAAV9-CB-GFP Vector Genomes Normalized to Reference Gene, *CFTR,* in Tissues of Treated Monkeys at 28 Days Post-Injection via the Intrathecal-LP Route.**

NHPs were dosed at 1.0×10^13^ vg/animal. (A) Select brain regions and (B) systemic tissues. For each tissue in each animal, values are averages of three technical replicates and calculated as vg/diploid genome. The median values reported from four replicate animals for all tissues. Values below the limit of quantitation were entered as zero for the purpose of calculating the median.

A.
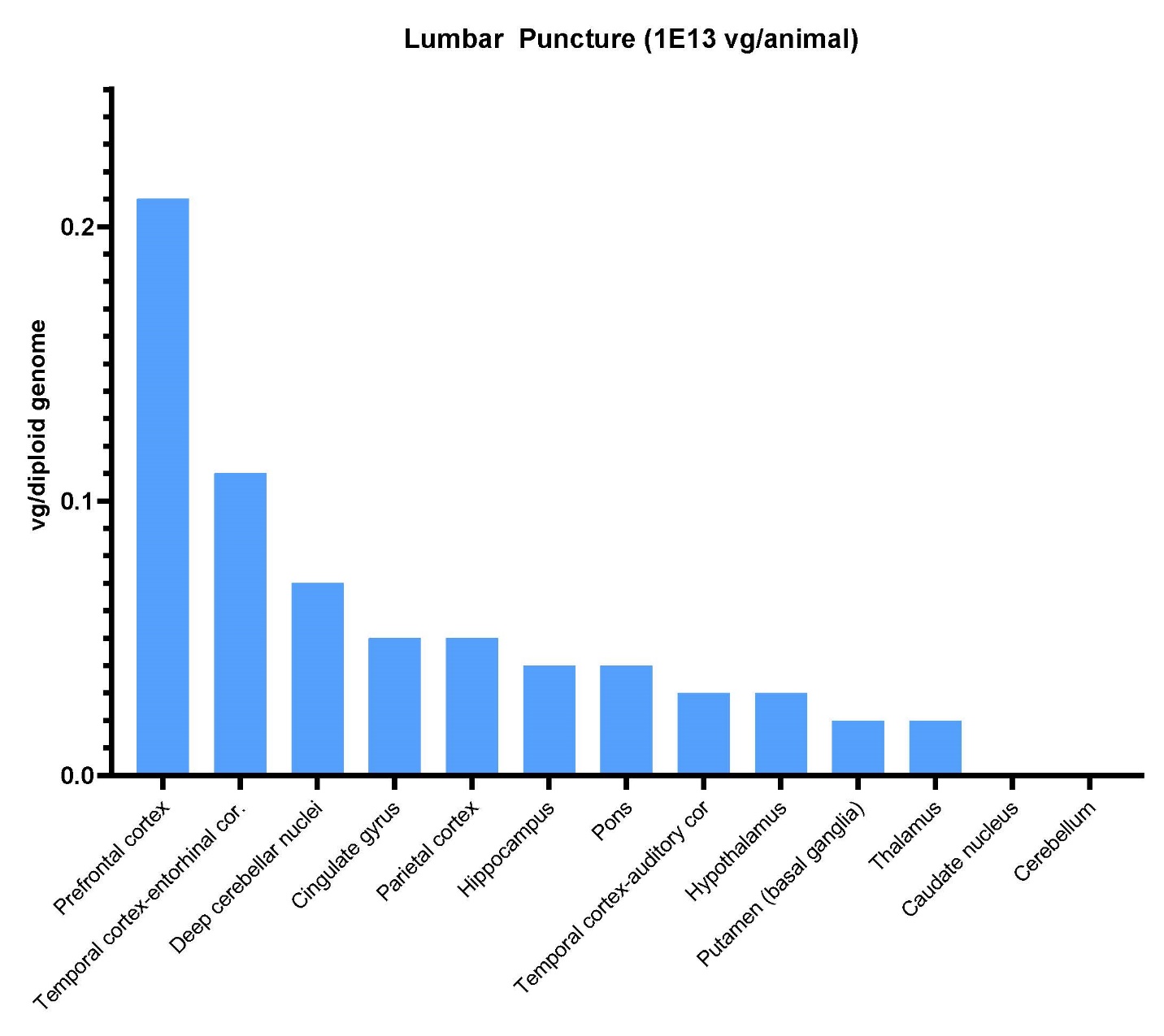

**B.**

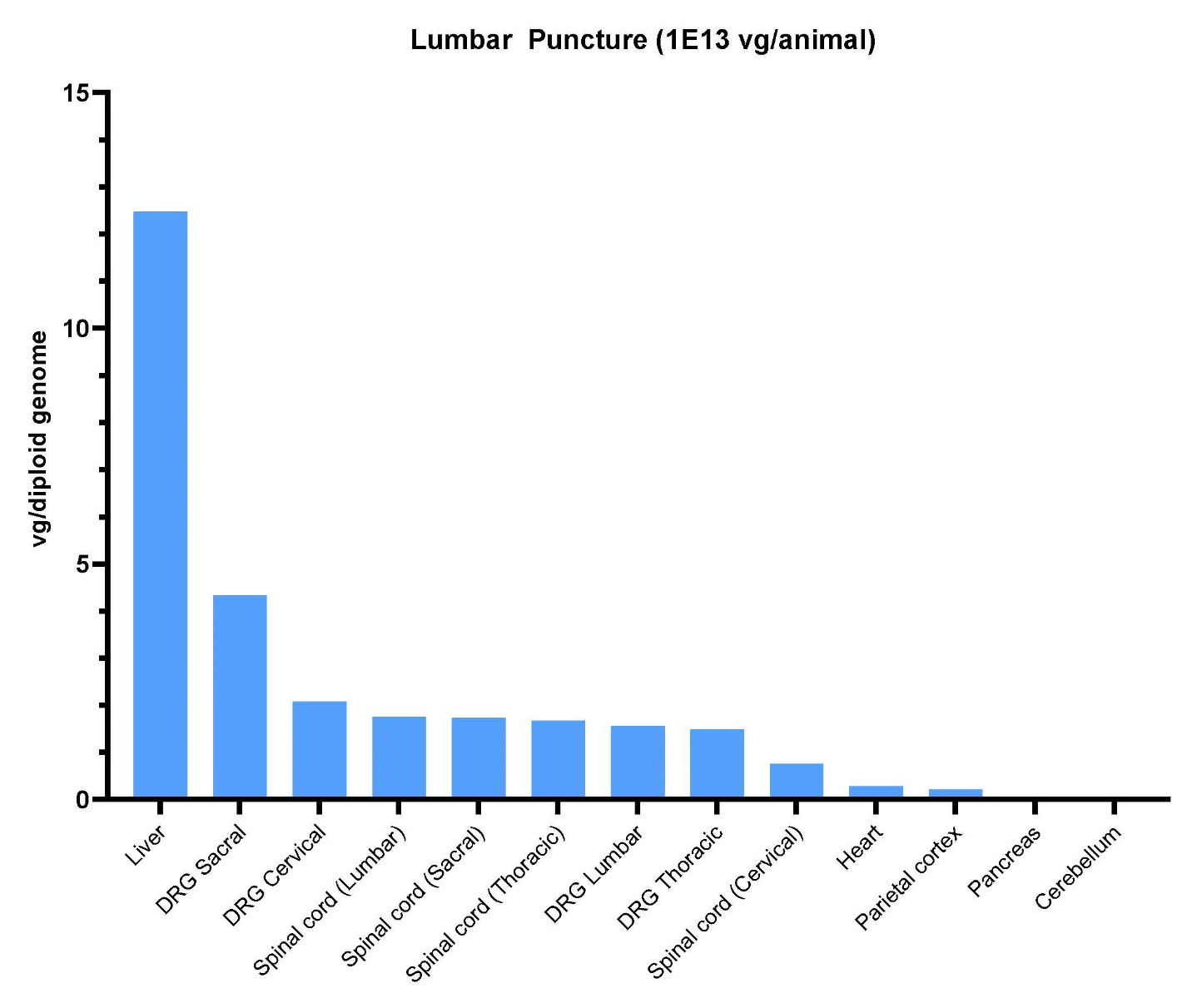

Abbreviations: *CFTR*, cystic fibrosis transmembrane conductance regulator; DRG, dorsal root ganglia; ICM, intracisterna magna; intrathecal-LP, intrathecal infusion by lumbar puncture; NHP, nonhuman primate; scAAV9-CB-GFP, self-complementary adeno-associated virus serotype 9–chicken β-actin promoter–green fluorescent protein; vg, vector genomes.

**Supplemental Figure 2. Biodistribution of scAAV9-CB-GFP Vector Genomes Normalized to Reference Gene, *CFTR,* in Tissues of Treated Monkeys at 28 Days Post-Injection via the ICM Route.**

NHPs were dosed at 1.0×10^13^ vg/animal. (A) Select brain regions and (B) systemic tissues. For each tissue in each animal, values were averages of three technical replicates and calculated as vg/diploid genome. The median values reported from four replicate animals for all tissues. Values below the limit of quantitation were entered as zero for the purpose of calculating the median.

**A.**

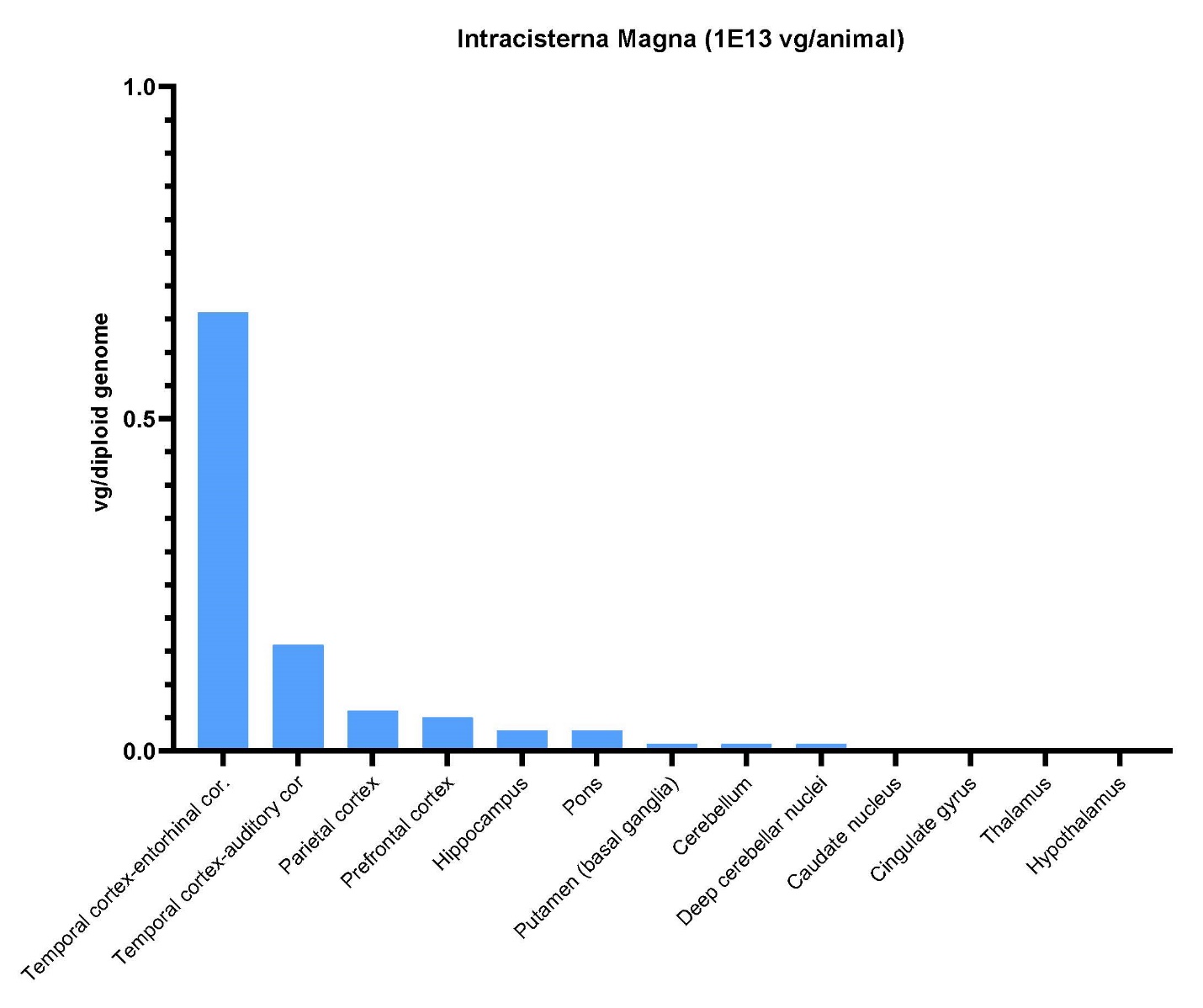

**B.**
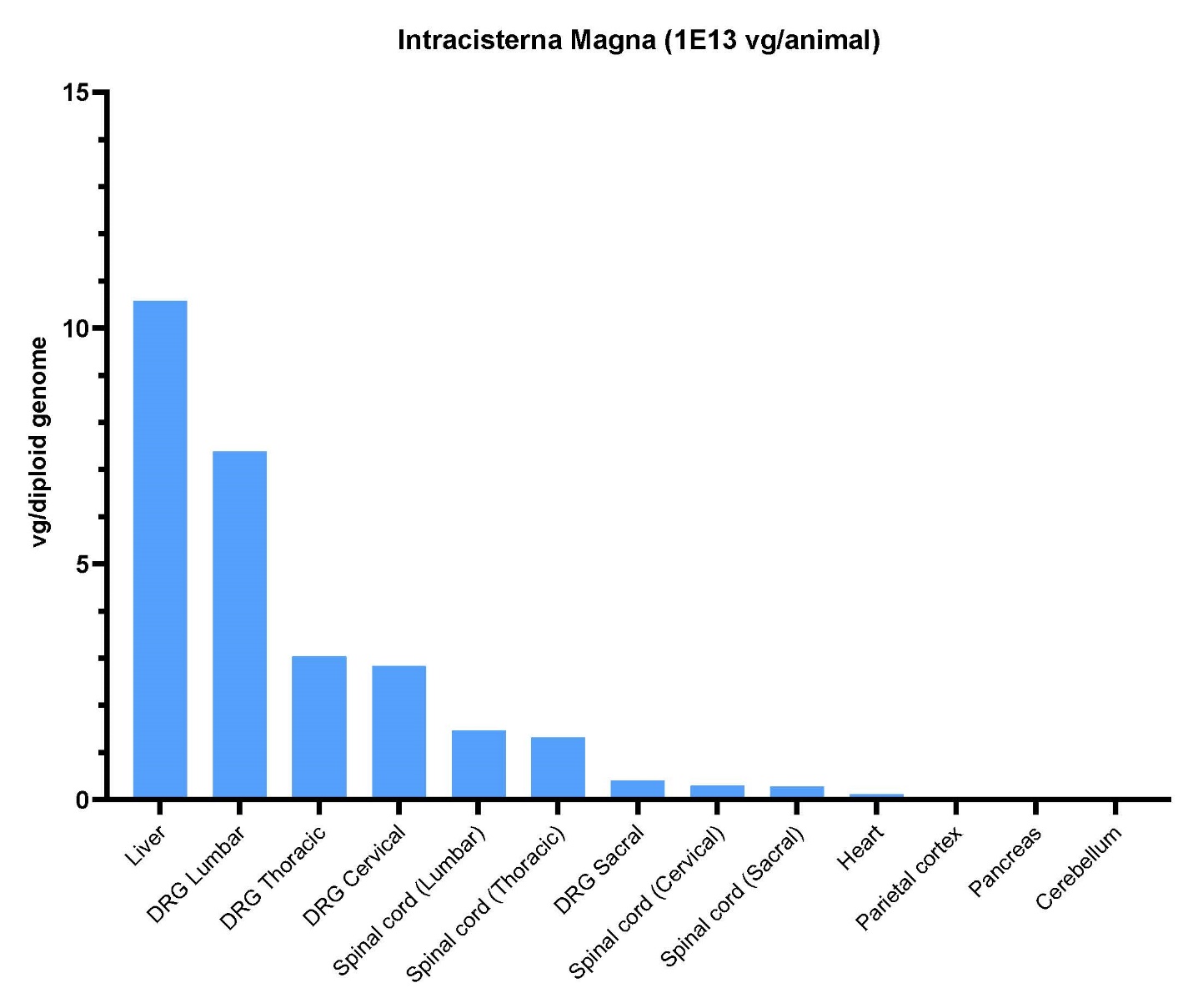

Abbreviations: *CFTR*, cystic fibrosis transmembrane conductance regulator; DRG, dorsal root ganglia; ICM, intracisterna magna; intrathecal-LP, intrathecal infusion by lumbar puncture; NHP, nonhuman primate; scAAV9-CB-GFP, self-complementary adeno-associated virus serotype 9–chicken β-actin promoter–green fluorescent protein; vg, vector genomes.

**Supplemental Figure 3. Biodistribution of scAAV9-CB-GFP Vector Genomes Normalized to Reference Gene, *CFTR,* in Tissues of Treated Monkeys at 28 Days Post-Injection via the ICM Route.**

NHPs were dosed at 3.0×10^13^ vg/animal. (A) Select brain regions and (B) systemic tissues. For each tissue in each animal, values are averages of three technical replicates and calculated as vg/diploid genome. The median values reported from four replicate animals. Values below the limit of quantitation were entered as zero for the purpose of calculating the median.

**A.**

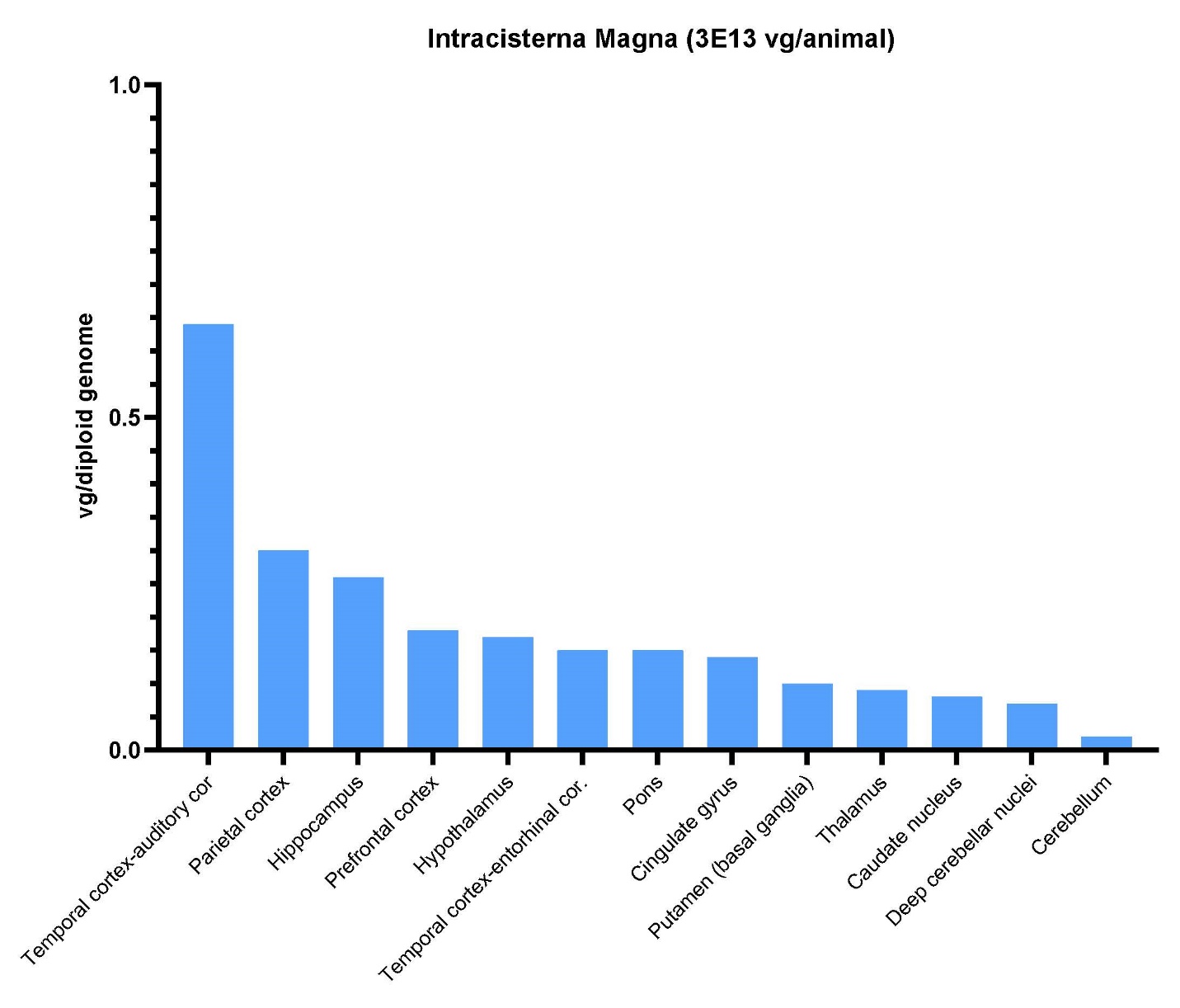

**B.**

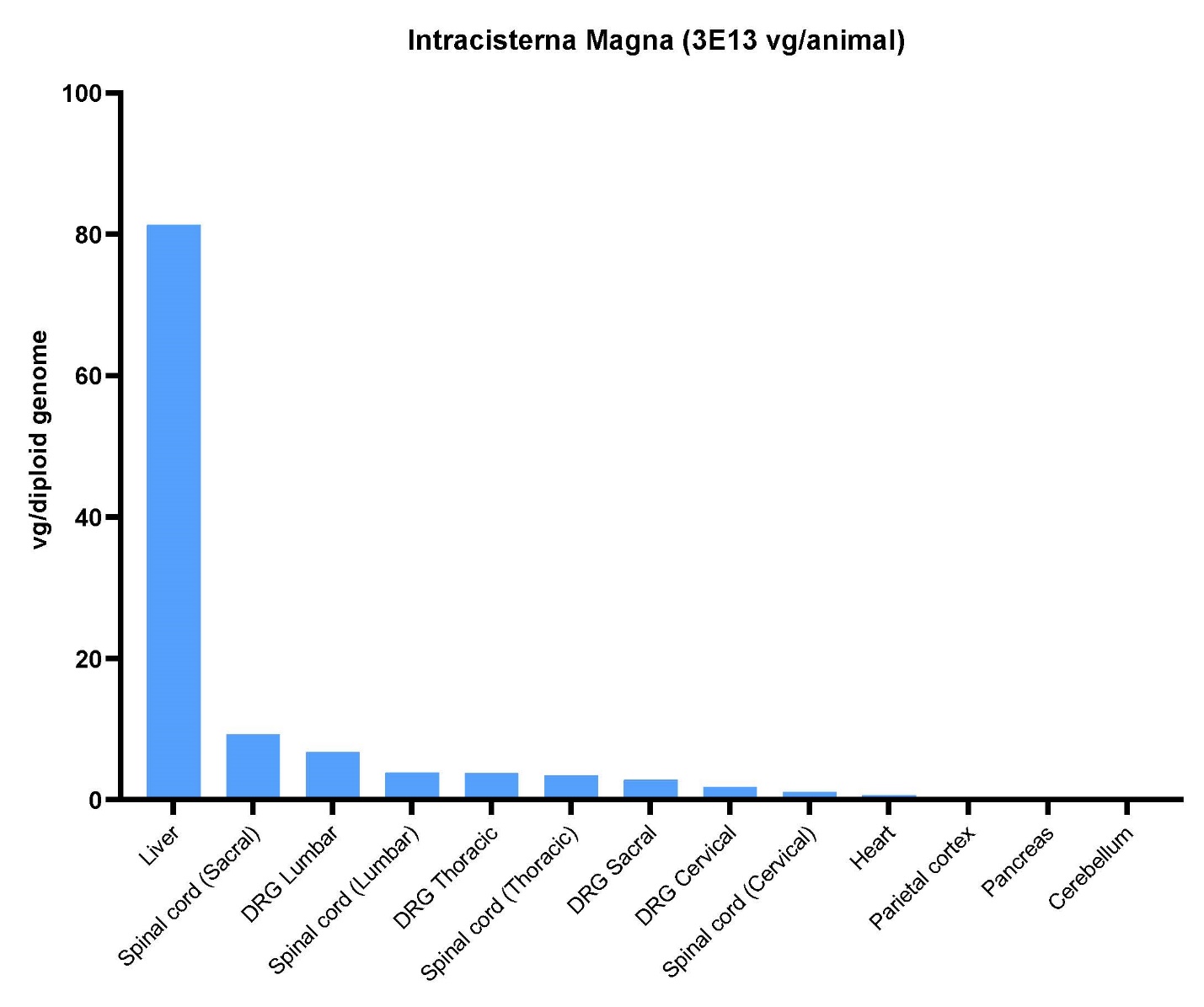

Abbreviations: *CFTR*, cystic fibrosis transmembrane conductance regulator; DRG, dorsal root ganglia; ICM, intracisternal magna; intrathecal-LP, intrathecal infusion by lumbar puncture; NHP, nonhuman primate; scAAV9-CB-GFP, self-complementary adeno-associated virus serotype 9–chicken β-actin promoter–green fluorescent protein; vg, vector genomes.

**Supplemental Figure 4. Biodistribution of GFP Following Intrathecal-LP of scAAV9-CB-GFP.** NHPs received intrathecal-LP scAAV9-CB-GFP at 1.0×10^13^ vg/animal. (A) Select brain regions and (B) systemic tissues. GFP was quantified at 28 days post-injection as measured by ECLIA. For each tissue in each animal, values are averages of two technical replicates and calculated as picogram GFP per mg total protein. The median values reported from four replicate animals for each tissue. Values below the limit of quantitation were entered as zero for the purpose of calculating the median.

**A.**

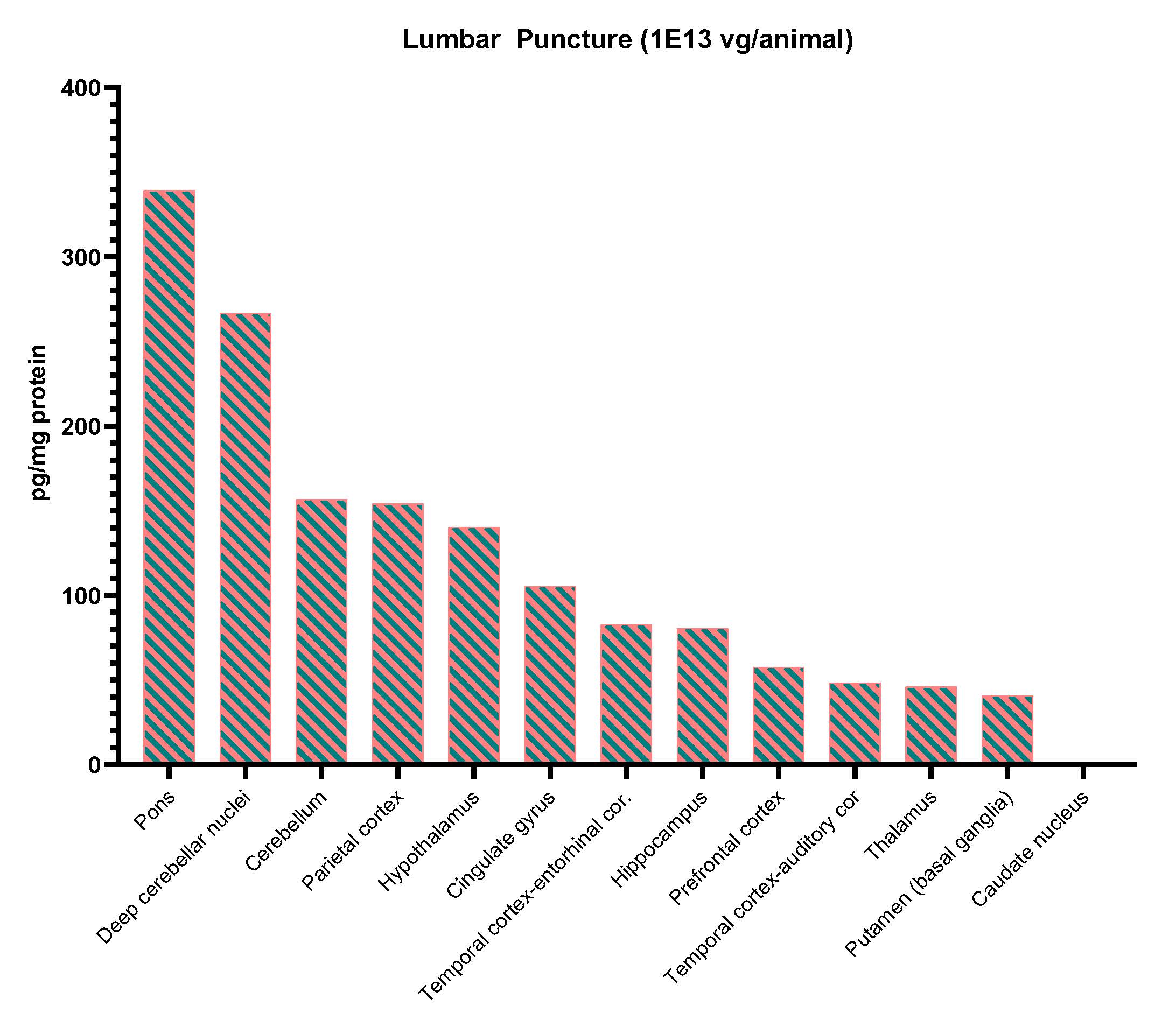

**B.**

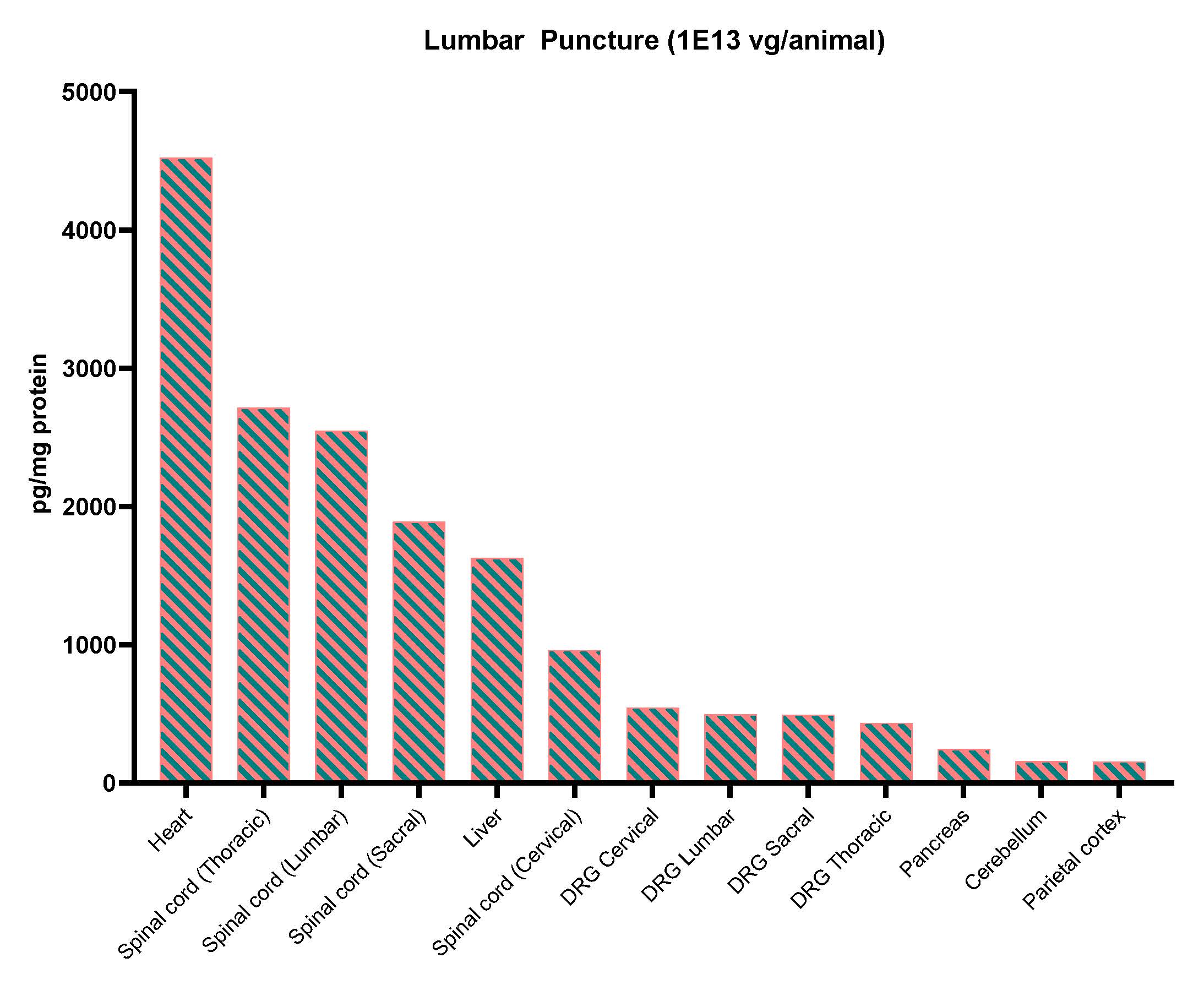

Abbreviations: DRG, dorsal root ganglia; ECLIA, electrochemiluminescence immunosorbent assay; ICM, intracisterna magna; intrathecal-LP, intrathecal infusion by lumbar puncture; NHP, nonhuman primate; scAAV9-CB-GFP, self-complementary adeno-associated virus serotype 9–chicken β-actin promoter–green fluorescent protein; vg, vector genomes.

**Supplemental Figure 5. Biodistribution of GFP Following ICM of scAAV9-CB-GFP.** NHPs received ICM scAAV9-CB-GFP at 1.0×10^13^ vg/animal. (A) Select brain regions and (B) systemic tissues. GFP was quantified at 28 days post-injection as measured by ECLIA. For each tissue in each animal, values are averages of two technical replicates and calculated as picogram GFP per mg total protein. The median values reported from four replicate animals for all tissues. Values below the limit of quantitation were entered as zero for the purpose of calculating the median.

**A.**

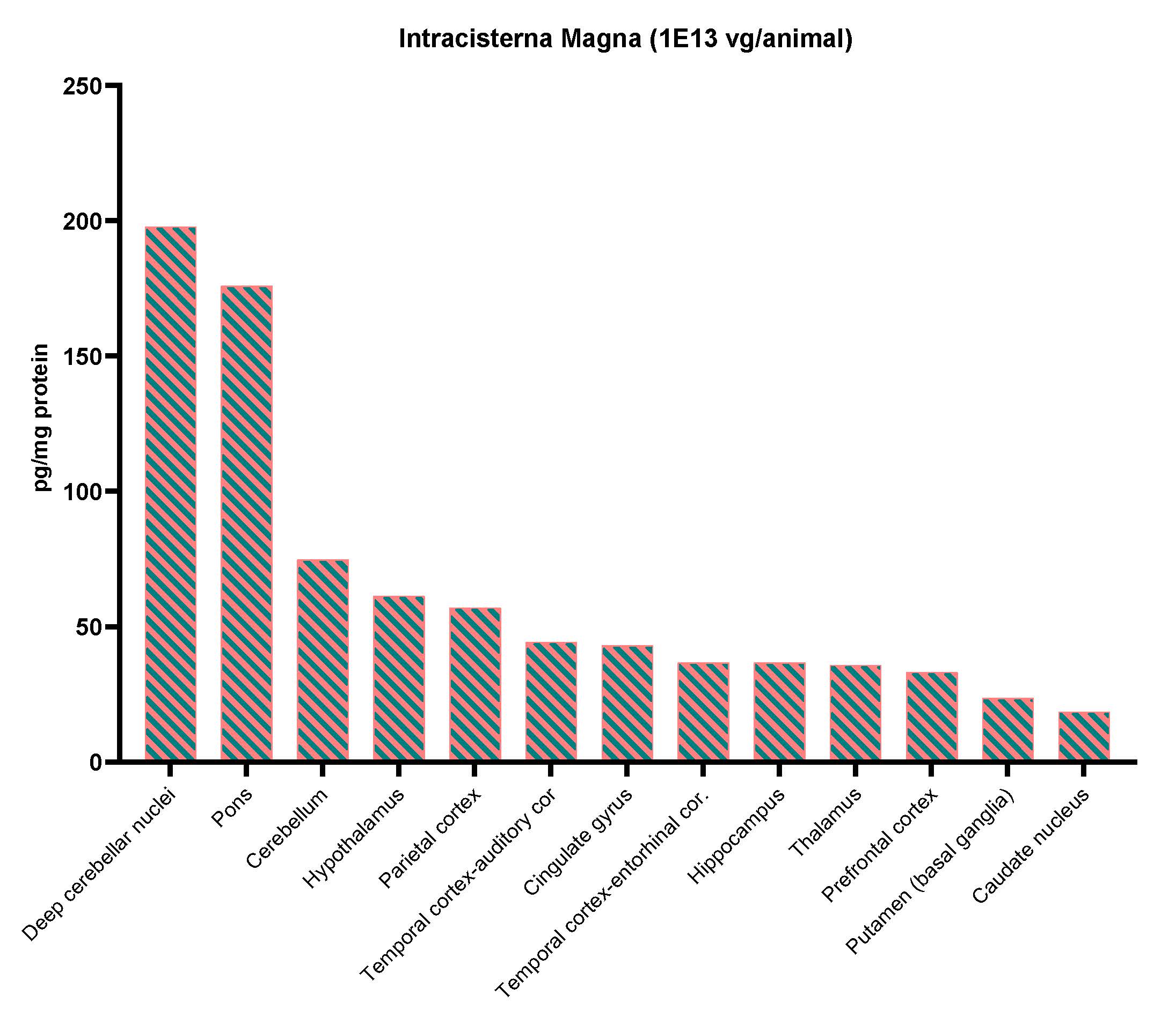

**B.**

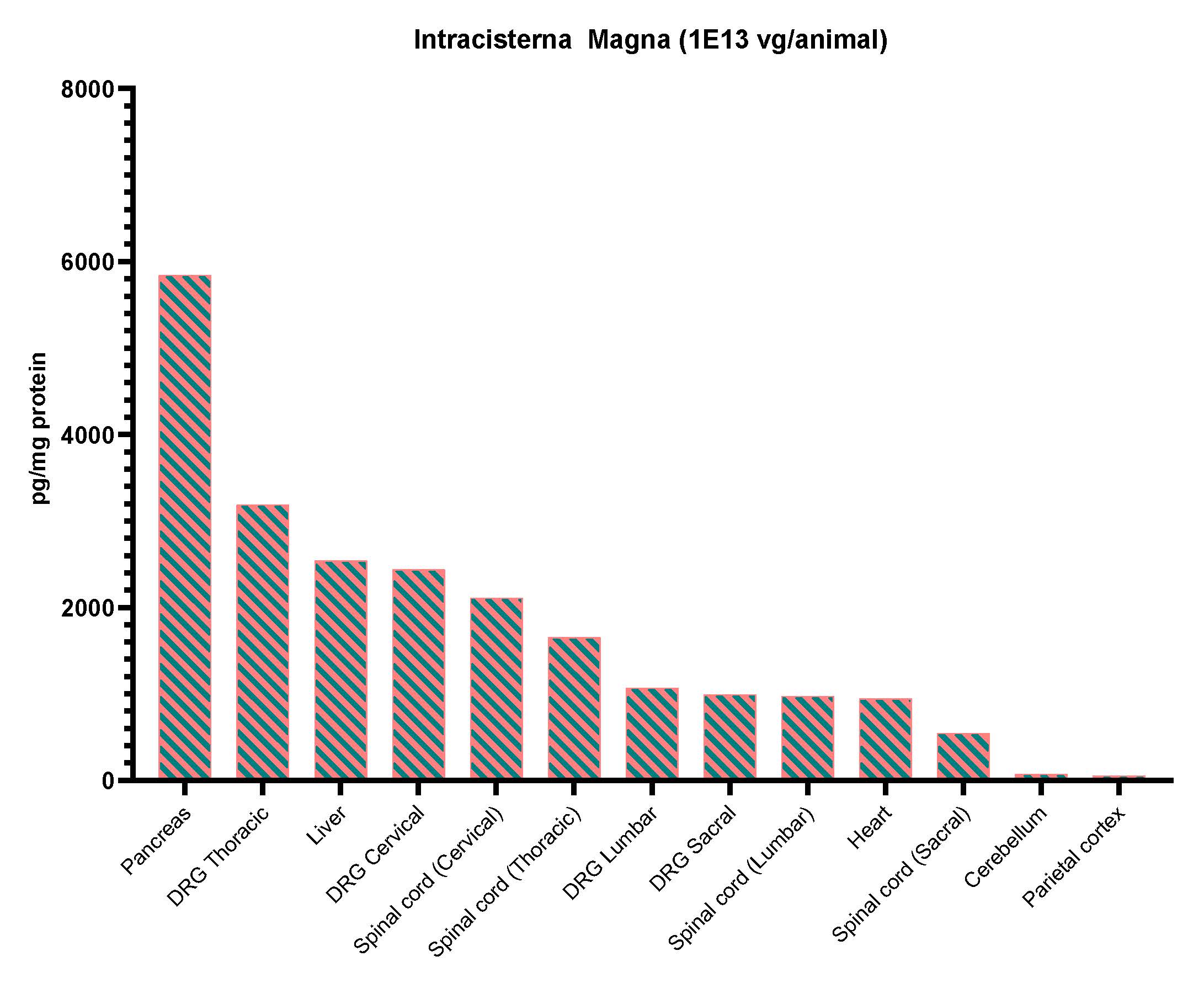

Abbreviations: DRG, dorsal root ganglia; ECLIA, electrochemiluminescence immunosorbent assay; ICM, intracisternal magna; intrathecal-LP, intrathecal infusion by lumbar puncture; NHP, nonhuman primate; scAAV9-CB-GFP, self-complementary adeno-associated virus serotype 9–chicken β-actin promoter–green fluorescent protein; vg, vector genomes.

**Supplemental Figure 6. Biodistribution of GFP Following ICM of scAAV9-CB-GFP.** NHPs received ICM scAAV9-CB-GFP at 3.0×10^13^ vg/animal. (A) Select brain regions and (B) systemic tissues. GFP was quantified at 28 days post-injection as measured by ECLIA. For each tissue in each animal, values are averages of two technical replicates and calculated as picogram GFP per mg total protein. The median values reported from four replicate animals for all tissues. Values below the limit of quantitation were entered as zero for the purpose of calculating the median.

**A.**

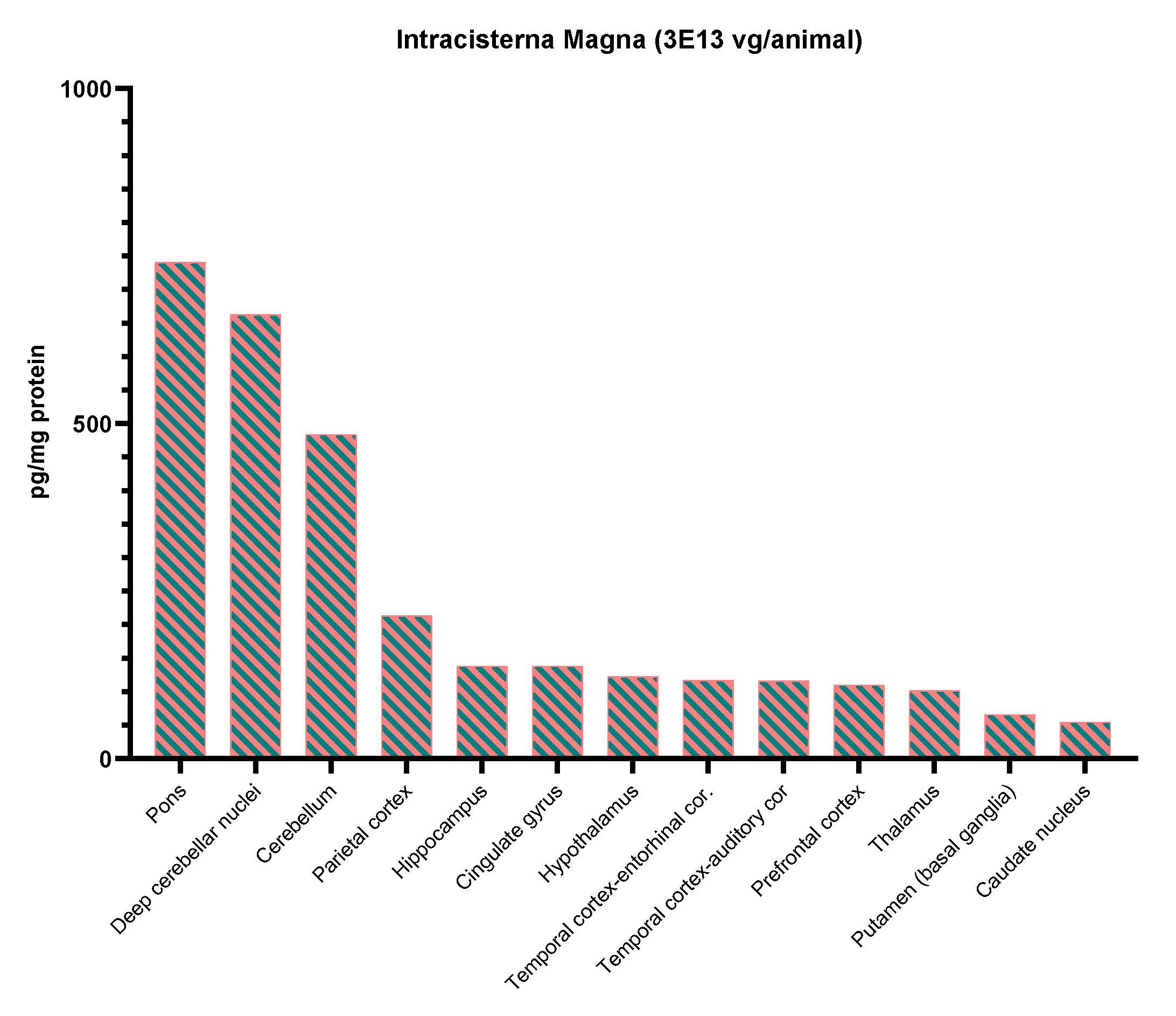

**B.**

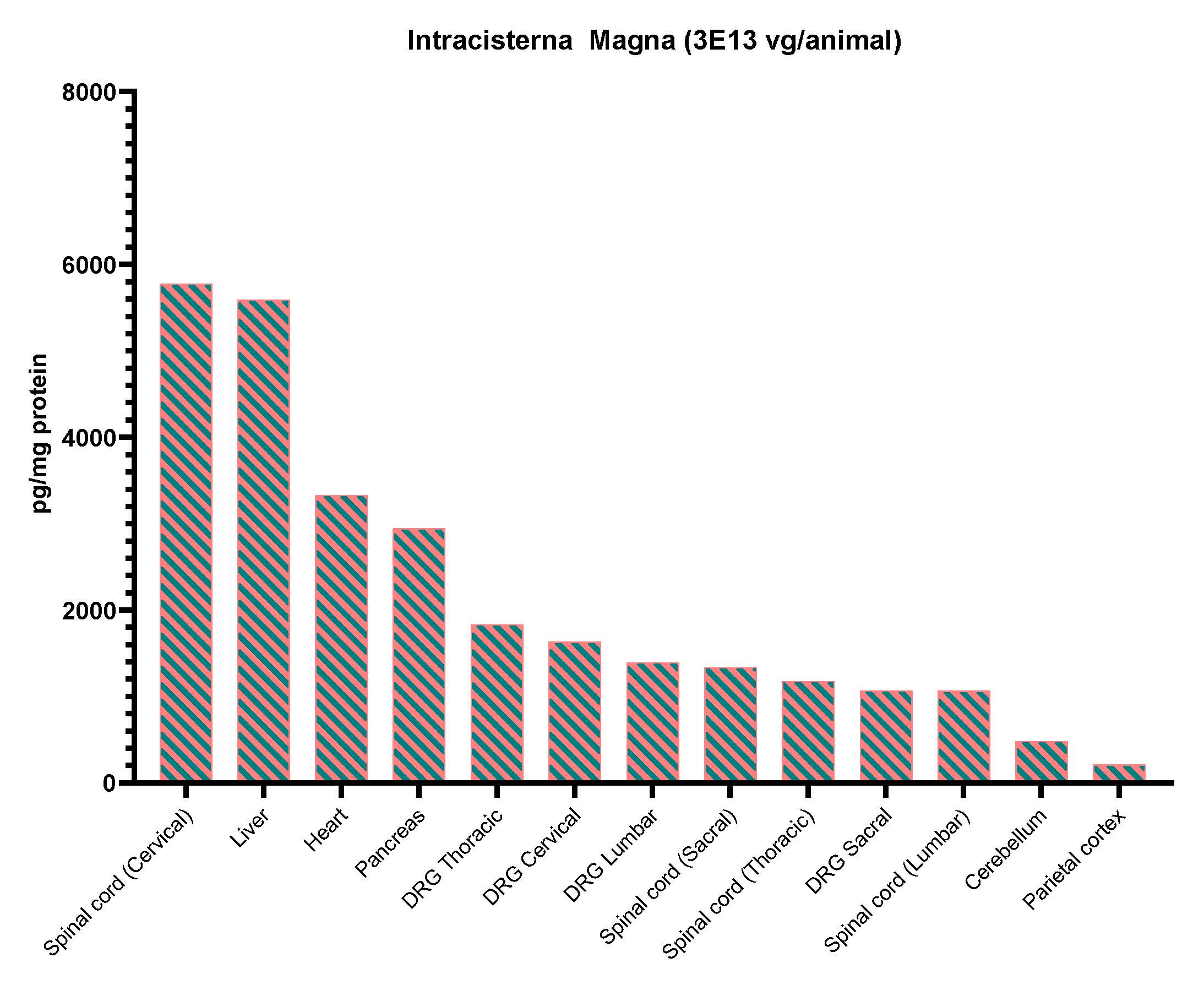

Abbreviations: DRG, dorsal root ganglia; ECLIA, electrochemiluminescence immunosorbent assay; ICM, intracisternal magna; intrathecal-LP, intrathecal infusion by lumbar puncture; NHP, nonhuman primate; scAAV9-CB-GFP, self-complementary adeno-associated virus serotype 9–chicken β-actin promoter–green fluorescent protein; vg, vector genomes.

**Supplemental Figure 7. Quantification of Green Fluorescent Protein Immunohistochemistry Signal by Image Analysis****.**

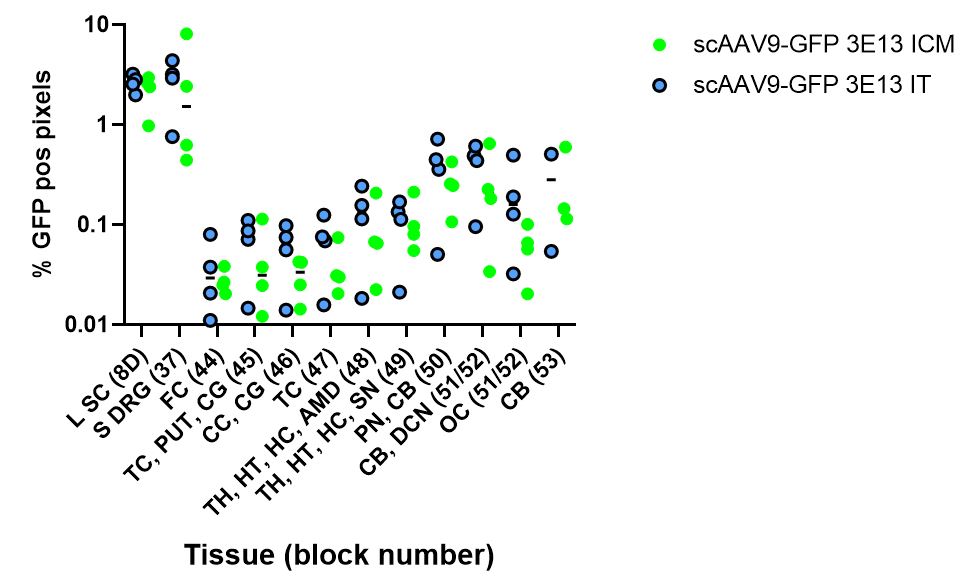

Abbreviations: AMD, amygdala; CB, cerebellum; CC, corpus callosum; CG, cingulate gyrus; DCN, deep cerebellar nuclei; FC, prefrontal cortex; GFP, green fluorescent protein; HC, hippocampus; HT, hypothalamus; ICM, intracisterna magna; IT, intrathecal; LP IT, intrathecal lumbar puncture; L SC, lumbar spinal cord; OC, occipital cortex; pos, positive; PN, pons; PUT, putamen; scAAV9-CB-GFP, self-complementary adeno-associated virus serotype 9–chicken β-actin promoter–green fluorescent protein; S DRG, sacral dorsal root ganglion; SN, substantia nigra; TC, temporal cortex; TH, thalamus.

**Supplemental Figure 8. Immunohistochemistry for Green Fluorescent Protein Performed on Liver (A), Skeletal Muscle (Biceps Femoris) (B), and Heart (C) in Animals Dosed with 3.0×10^13^ vg/animal scAAV9-CB-GFP by the Intrathecal-LP Route.**

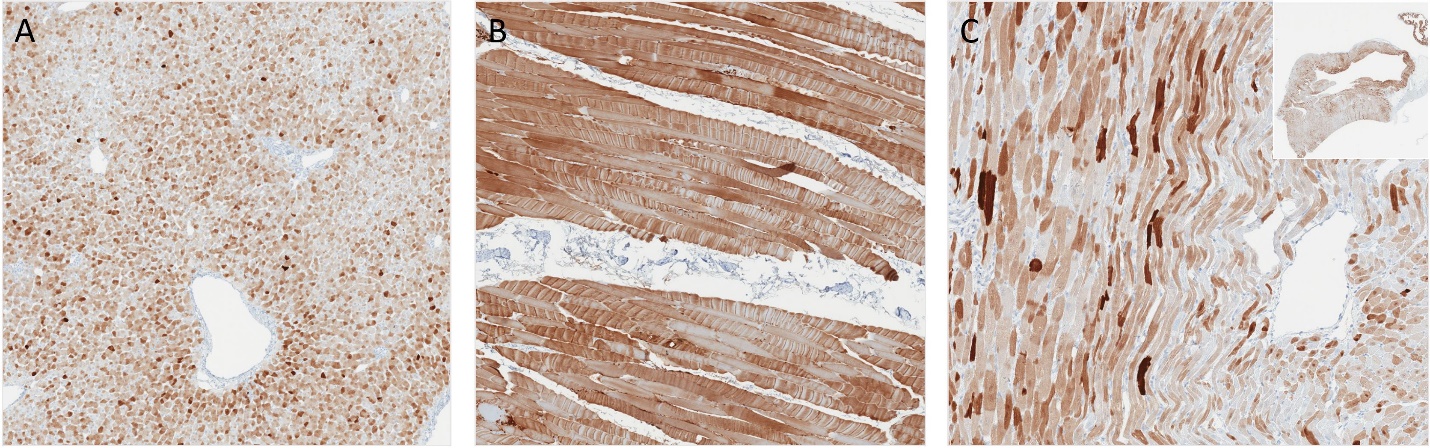

Abbreviations: LP, lumbar puncture; scAAV9-CB-GFP, self-complementary adeno-associated virus serotype 9–chicken β-actin promoter–green fluorescent protein; vg, vector genomes.

**Supplemental Figure 9. Green Fluorescent Protein Immunohistochemistry on Tissue from Control Animals Dosed Via the Intrathecal-LP Route Demonstrating Absence of Non-Specific Signal.**

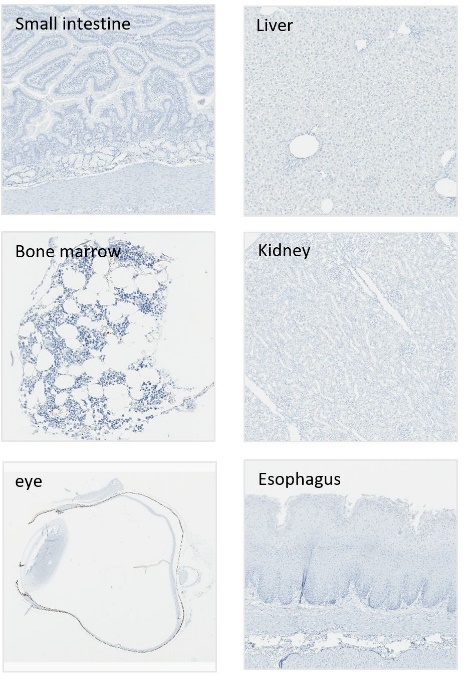

Abbreviations: LP, lumbar puncture.

**Supplemental Figure 10. Green Fluorescent Protein Immunohistochemistry on Tissue from Control Animals Dosed Via the Intrathecal-LP Route Demonstrating Absence of Non-Specific Signal.**

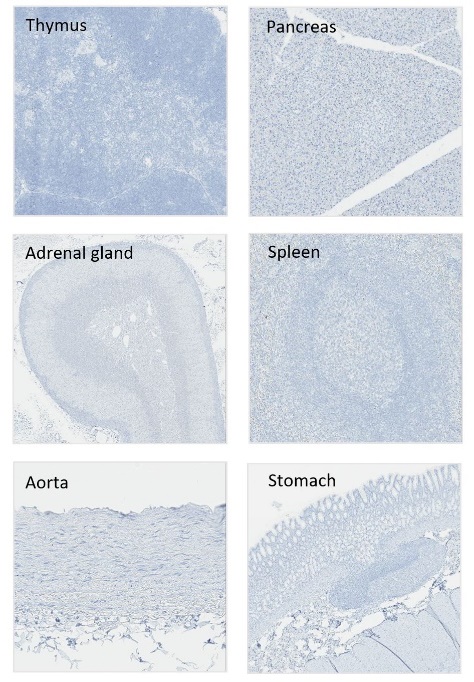

Abbreviations: LP, lumbar puncture.

**Supplemental Figure 11. Green Fluorescent Protein Immunohistochemistry on Tissue from Control Animals Dosed Via the Intrathecal-LP Route Demonstrating Absence of Non-Specific Signal.**

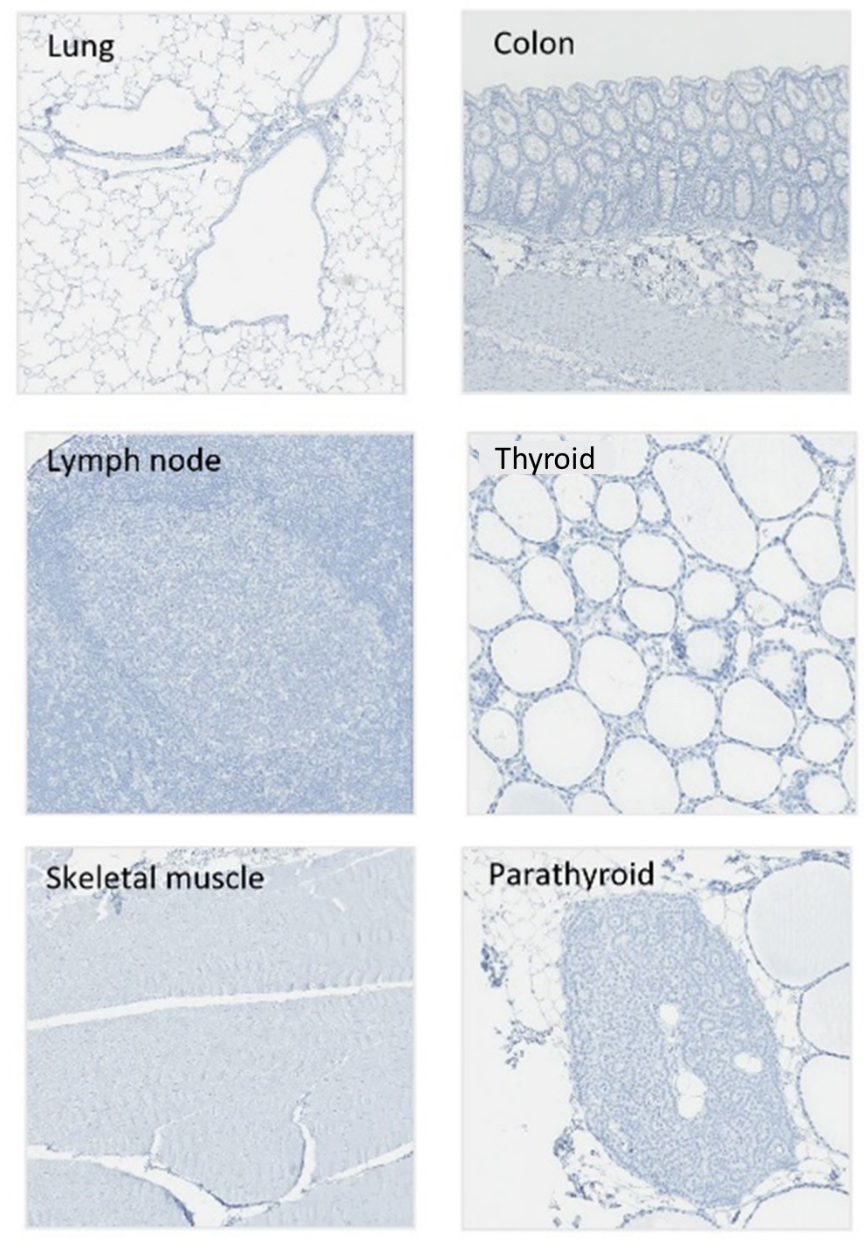

Abbreviations: LP, lumbar puncture.

**Supplemental Figure 12. *In situ* Hybridization for Vector Nucleic Acid Using Anti-Sense Probe in Liver (A), Skeletal Muscle (Biceps Femoris) (B), and Heart (C) in Animals Dosed with 3.0×10^13^ vg/animal scAAV9-CB-GFP by the Intrathecal-LP Route.**

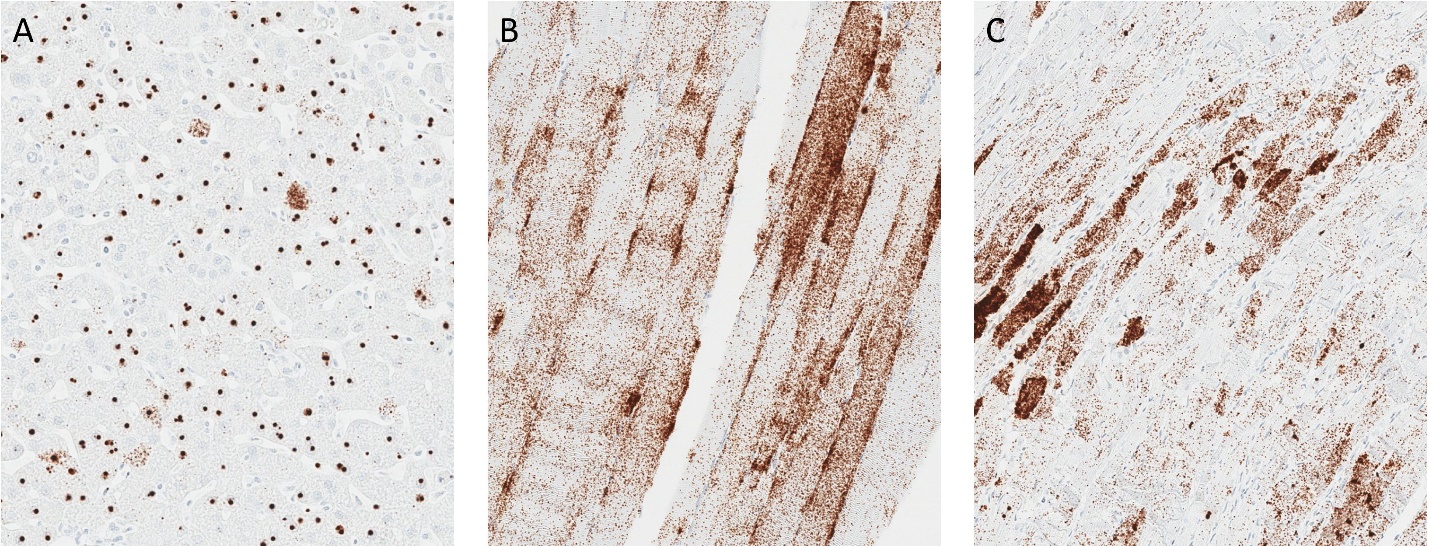

Abbreviations: LP, lumbar puncture; scAAV9-CB-GFP, self-complementary adeno-associated virus serotype 9–chicken β-actin promoter–green fluorescent protein; vg, vector genomes.

**Supplemental Figure 13. Green Fluorescent Protein Immunohistochemistry Performed on Tissues from an Animal Dosed with** **3.0×10^13^ vg scAAV9-CB-GFP by the Intrathecal-LP Route. (A) Thymus, (B) Spleen, (C) Lymph Node, (D) Bone Marrow, (E) Aorta, (F) Brown Adipose Tissue, (G) Adrenal Gland, (H) Thyroid Gland, and (I) Parathyroid Gland.**

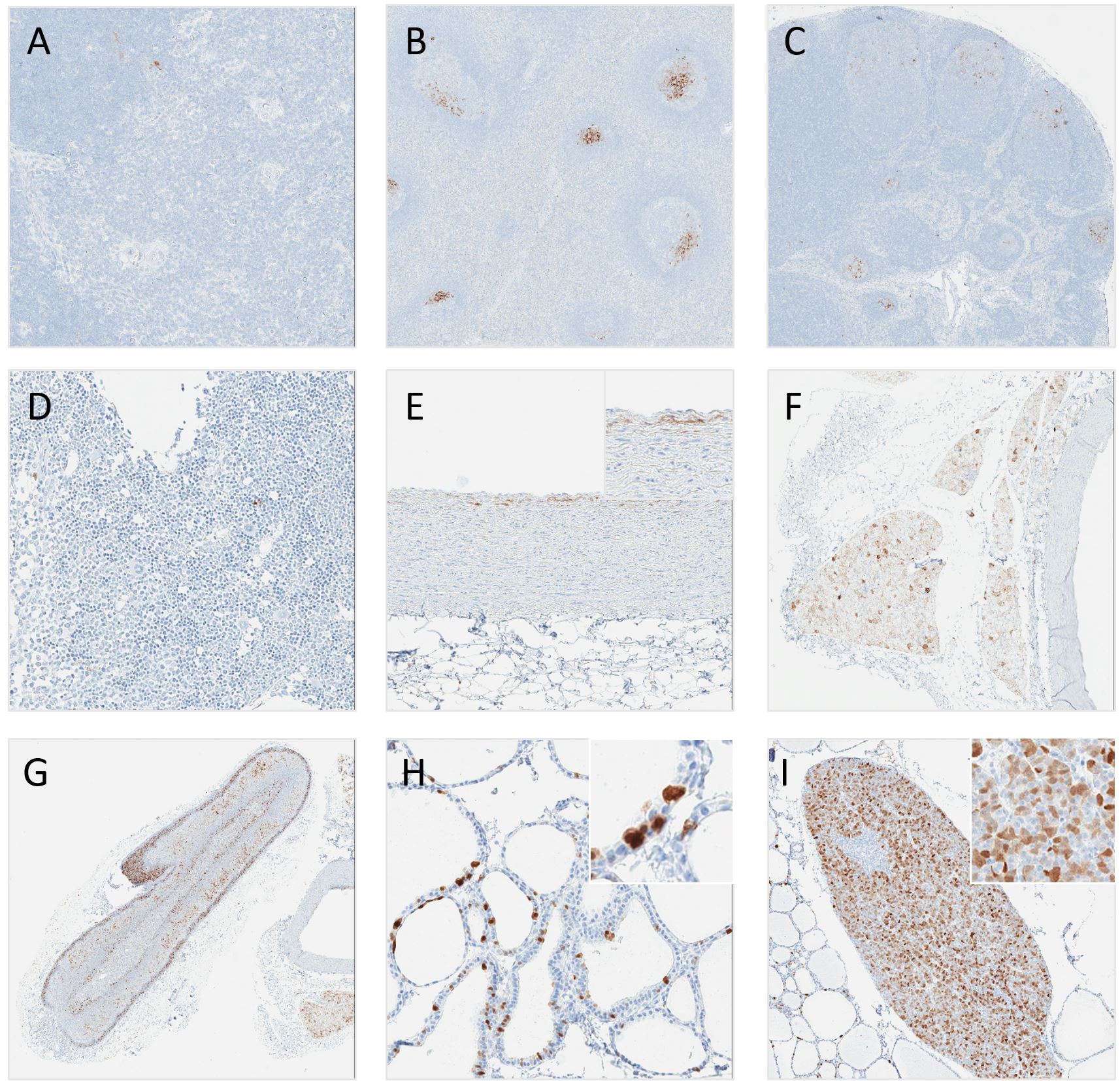

Abbreviations: LP, lumbar puncture; scAAV9-CB-GFP, self-complementary adeno-associated virus serotype 9–chicken β-actin promoter–green fluorescent protein; vg, vector genomes.

**Supplemental Figure 14. Green Fluorescent Protein Immunohistochemistry Performed on Tissues from an Animal Dosed with 3.0×10^13^ vg scAAV9-CB-GFP by the Intrathecal Route. (A) Lung, (B) Bronchiole, (C) Kidney, (D) Esophagus, (E) Stomach, (F) Small Intestine, (G) Colon, (H) Exocrine Pancreas, and (I) Islet of Langerhans.**

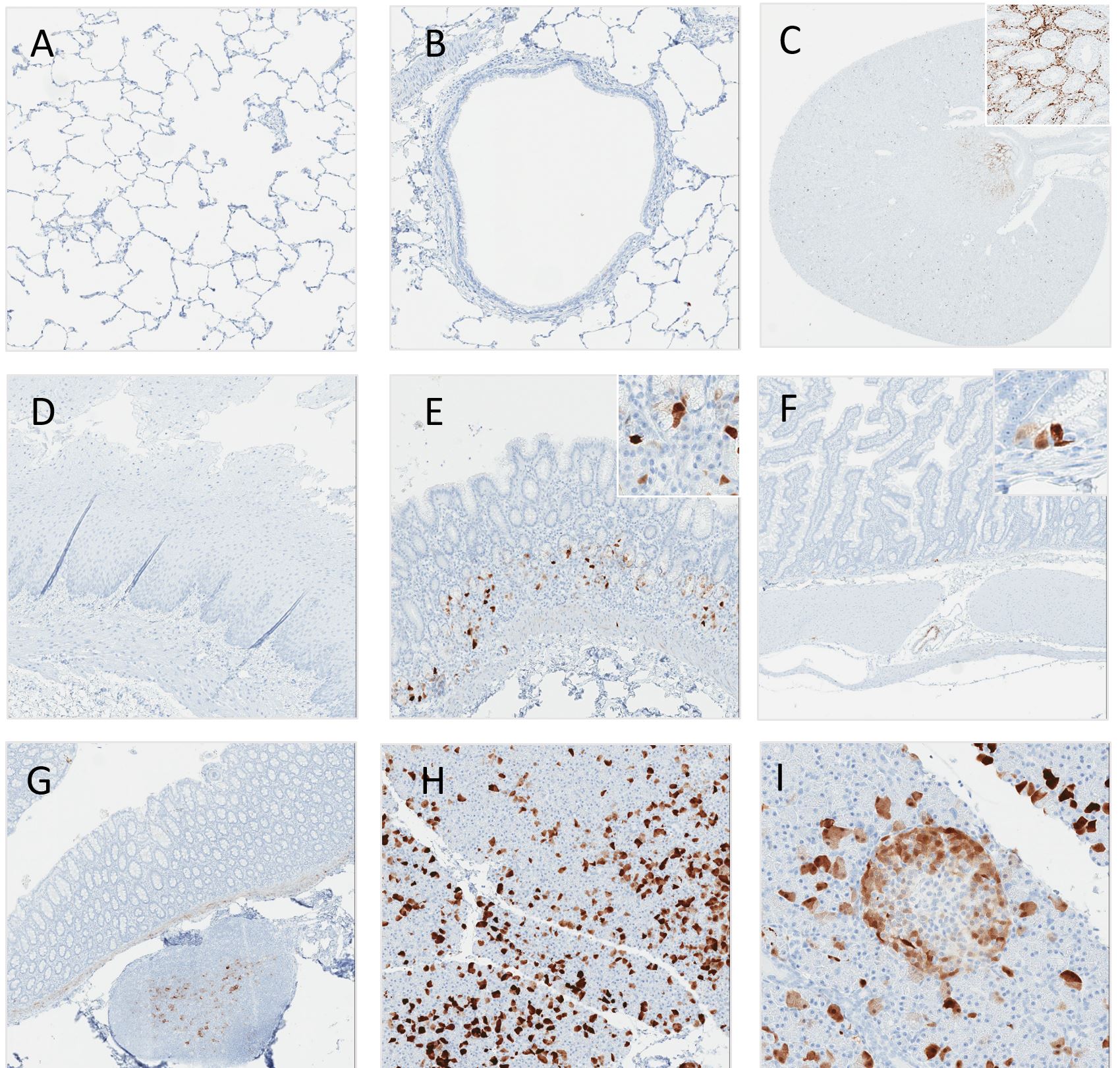

Abbreviations: scAAV9-CB-GFP, self-complementary adeno-associated virus serotype 9–chicken β-actin promoter–green fluorescent protein; vg, vector genomes.

**Supplemental Figure 15a-i. Macroscopic sections of brain sampled for histopathology and biodistribution (ddPCR for CNV and/or ECLIA for GFP protein). Areas sampled for histopathology indicated by the red boxes included prefrontal and parietal cortices, temporal (auditory and entorhinal) cortices, hippocampus, caudate nucleus, putamen, thalamus, hypothalamus, pons, cerebellum, and deep cerebellar nuclei. Frozen samples were collected from the contralateral structures.**

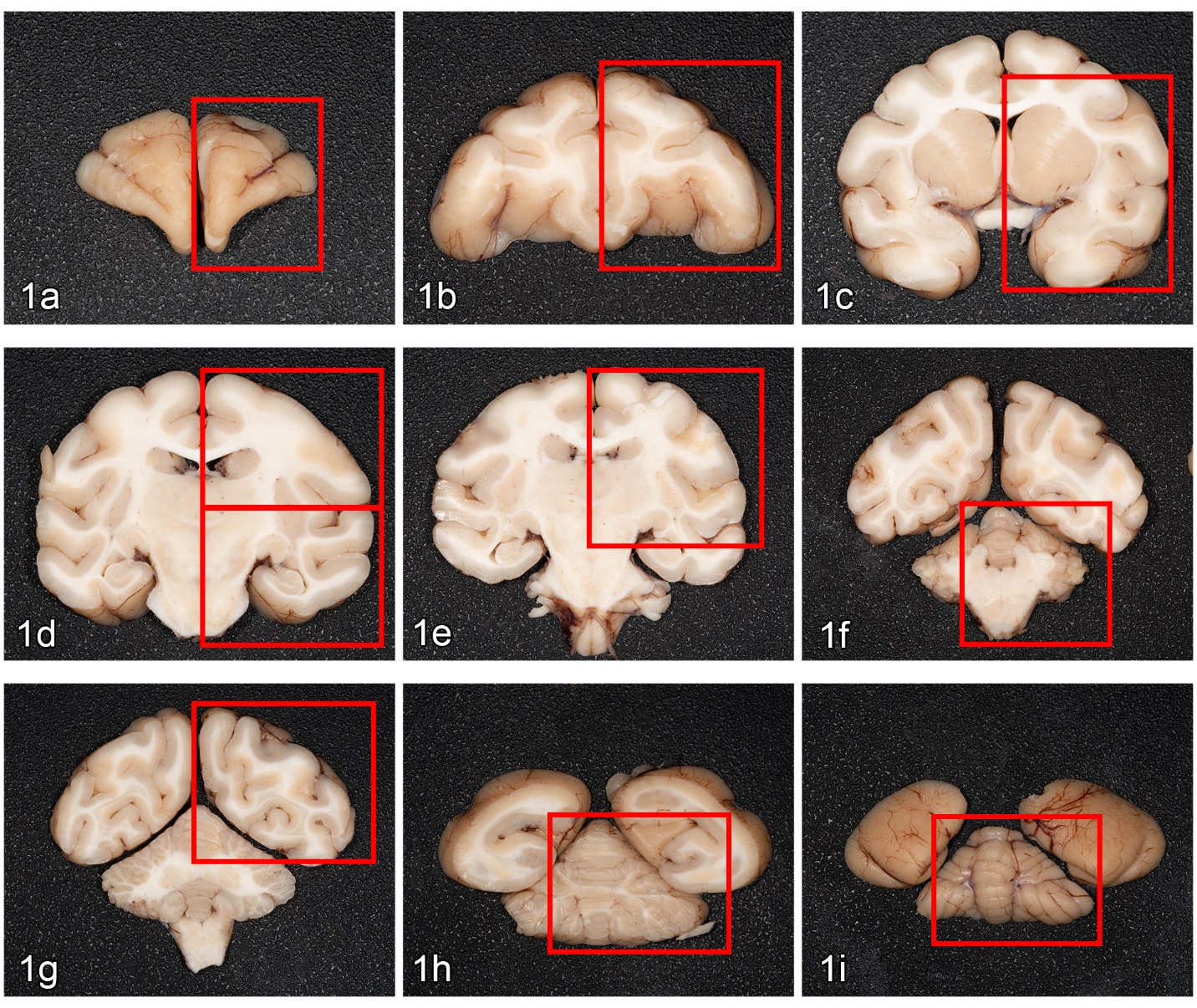
